# Assessment of enzyme active site positioning and tests of catalytic mechanisms through X-ray-derived conformational ensembles

**DOI:** 10.1101/786327

**Authors:** Filip Yabukarski, Justin T Biel, Margaux M Pinney, Tzanko Doukov, Alexander S Powers, James S Fraser, Daniel Herschlag

## Abstract

How enzymes achieve their enormous rate enhancements remains a central question in biology, and our understanding to date has impacted drug development, influenced enzyme design, and deepened our appreciation of evolutionary processes. While enzymes position catalytic and reactant groups in active sites, physics requires that atoms undergo constant motion. Numerous proposals have invoked positioning or motions as central for enzyme function, but a scarcity of experimental data has limited our understanding of positioning and motion, their relative importance, and their changes through the enzyme’s reaction cycle. To examine positioning and motions and test catalytic proposals, we collected “room temperature” X-ray crystallography data for *P. putida* ketosteroid isomerase (KSI), and we obtained conformational ensembles for this and a homologous KSI from multiple PDB crystal structures. Ensemble analyses indicated limited change through KSI’s reaction cycle. Active site positioning was on the 1-1.5 Å scale, and was not exceptional compared to non-catalytic groups. The KSI ensembles provided evidence against catalytic proposals invoking oxyanion hole geometric discrimination between the ground state and transition state or highly precise general base positioning. Instead, increasing *or* decreasing positioning of KSI’s general base reduced catalysis, suggesting optimized Ångstrom-scale conformational heterogeneity that allows KSI to efficiently catalyze multiple reaction steps. Ensemble analyses of surrounding groups for WT and mutant KSIs provided insights into the forces and interactions that allow and limit active site motions. Most generally, this ensemble perspective extends traditional structure–function relationships, providing the basis for a new era of “ensemble–function” interrogation of enzymes.

## Introduction

The central role of enzymes in biology is embodied in the decades of effort spent to deeply investigate the origins of their catalysis (1–10). Enzymes studies now routinely identify the active site groups that interact with substrates and reveal their roles in binding and in facilitating chemical transformations. Nevertheless, these so-called “catalytic groups” alone, outside of the context of a folded enzyme, do not account for the enormous rate enhancements and exquisite specificities exhibited by enzymes (7). Accordingly, classic proposals for enzyme catalysis have invoked the importance of positioning of active site groups within a folded enzyme and of substrates localized and positioned by binding interactions (9, 11–24). While these proposals universally invoke restricted freedom of motion of catalytic groups, the amount of catalysis provided by positioning has been the subject of much conjecture, discussion, and debate (25–33). Conversely, it is also now clear that motions are inherent to enzymes, and that conformational transitions and structural rearrangements are important for enzyme function (17, 34–40). Considering both positioning and motions, it has been recognized that:

> “For catalysis, flexible but not too flexible, as well as rigid but not too rigid, is essential. Specifically, the protein must be rigid enough to maintain the required structure but flexible enough to permit atomic movements as the reaction proceeds.” (5)

The importance of both positioning and motions to enzyme function suggests a nuanced view of enzyme catalysis and underscores the need for direct experimental measurements of positioning and motions within enzymes.

As Feynman noted, “Everything that living things do can be understood in terms of the jigglings and wigglings of atoms” (41). But simply observing motions of active site residues does not tell us how enzymes achieve catalysis. To understand enzymes, we want to know how much an enzyme dampens and alters the motions of catalytic residues. We want to know which increases or decreases in motion increase or decrease the reaction rate, whether the dampening is uniform or whether there are certain preferred coordinates, and what interactions and forces are most responsible for the dampening, so that we can better understand how to design new enzymes. Additionally, to what extent are these residues positioned upon folding of the enzyme, or adjusted and honed as the reaction proceeds? And are active site residues more precisely positioned than analogous residues throughout a folded protein?

To address fundamental questions about how enzymes function, how they evolved, and how to ultimately design new enzymes that rival those of Nature, we need to obtain experimental information about enzyme conformation ensembles—the distribution of enzyme states dictated by their highly-complex multi-dimensional energy landscapes. Observations of well-resolved electron densities from traditional X-ray diffraction data indicate positioning of residues in and around the active site, but do not provide information on the extent and nature of that positioning. Crystallographic B-factors of residues are sometimes used to infer dynamics, but are only indirectly related to intrinsic motion and contain contributions from additional factors such as crystallographic order (42–44). NMR experiments identify groups that experience greater motional freedom and can provide temporal information, but these experiments typically lack information about the directions and extent of these motions (45, 46). Molecular dynamics simulations provide atomic-level models for entire systems, but we currently lack the rigorous experimental tests needed to determine whether or not computational outputs reflect actual physical behavior, which prevents firm mechanistic conclusions from being inferred (47–49).

Fortunately, two X-ray crystallographic approaches have recently emerged that can provide experimentally-derived information about conformational ensembles: high sequence similarity PDB structural ensembles (referred to as “pseudo-ensembles” herein) (50, 51) and multi-conformer models from X-ray data obtained at temperatures above the protein’s glass transition (referred to as “room temperature” or “RT” X-ray diffraction in the literature and herein) (36, 52– 54). These approaches are complementary. Pseudo-ensembles provide information about residues and functional groups that move in concert (i.e., coupled motions) but require dozens of structures. RT X-ray data from single crystals can provide multi-conformer models, allowing ensemble information about new complexes and mutants to be more readily acquired, but do not provide direct information about coupled motions. In addition, RT X-ray studies provide direct information about equilibrium distributions without freezing, which can alter and quench protein motions, and without assuming that different frozen crystals reproduce an equilibrium distribution of states (52, 55–59).

Here we take advantage of the strengths of both approaches: the ability to evaluate correlated side chain rearrangements in and near the active site *via* pseudo-ensembles and the ability to obtain new ensemble-type information of new states from single X-ray datasets at temperatures above the glass transition. In addition, the ability to compare these data aids in evaluating the potential for artifacts from cryo-freezing. Focusing on a model enzyme with very high-resolution data and with analogs representing successive steps along its reaction path has allowed us to obtain insights that would not be possible from static structures, from either ensemble approach alone, or from less extensive or lower resolution data.

We chose the enzyme ketosteroid isomerase (KSI; Figure 1) for this investigation because of our ability to obtain diffraction data of high-resolution, because of the accumulated wealth of structural and mechanistic information for this enzyme, and because of KSI’s use of catalytic strategies common to many enzymes. We obtained ensemble data for KSI from two species, which gave consistent results and allowed us to address unresolved questions from decades of KSI studies: we tested positioning as the origin of the extremely high effectiveness of the KSI general base, and we evaluated a model in which KSI’s oxyanion hole provides geometric discrimination between the reaction’s ground and transition states. We also used our ensembles from two KSI homologs to ask, and answer, more general questions: are catalytic groups more conformationally restricted than non-catalytic residues and do conformational restrictions increase through the reaction cycle? Further, using ensemble data for WT and mutant KSIs, we were able to assess contributions to positioning *vs.* conformational heterogeneity from competing interactions and forces. Most generally, our in-depth analyses of KSI bring an ensemble perspective to bear on traditional structure–function studies and provide the basis for a new era of ensemble–function studies.

**Figure 1.**
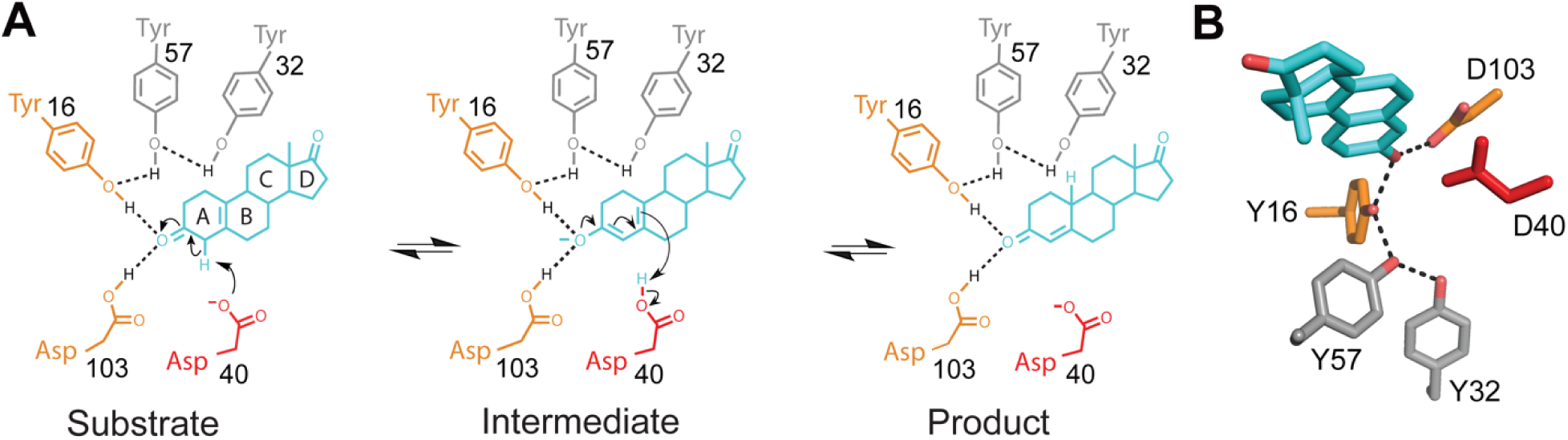
The Ketosteroid Isomerase (KSI) reaction. Reaction mechanism and schematic depiction of the active site (**A**) and its three-dimensional organization (**B**) (PDB 1OH0 (60)). KSI catalyzes double bond isomerization of steroid substrates utilizing a general acid/base D40 (which we refer to herein as a general base, for simplicity), and an oxyanion hole composed of the side chains of Y16 and D103 (protonated); general base and oxyanion hole residues are colored in red and orange, respectively.

## Results

### Limited structural changes throughout the KSI catalytic cycle

Prior analyses of crystal structures for 60 enzymes revealed modest structural changes between Apo and ligand-bound enzyme states, with RMSD < 1 Å on average, and no larger than differences between two Apo forms of the same enzyme in most cases (61). To evaluate the extent to which the KSI structure changes through its catalytic cycle, we took advantage of the 94 crystallographically-independent KSI molecules from the 45 cryo crystal structures available in the Protein Data Bank (PDB, Berman et al., 2000) (Table S1). All these structures, whether Apo, ground-state-analog bound (GSA-bound), or transition-state-analog bound (TSA-bound), were highly similar, as seen visually in Figure 2A and Figure S1B-D and by their similar RMSDs of <1 Å (Figure 2B, top, Figure S2, top, Figure S3A). Two short loops, encompassing residues 62–65 and residues 91–96, exhibit the greatest variation (Figure 2A and Figure S1B-D), and when these loops, representing <10% of KSI’s sequence, are excluded the RMSDs drop by half, with all now falling below 0.5 Å (Figure 2B, bottom, Figure S2, bottom, Figure S3B). To assess possible conformational changes in the reaction cycle, despite the strong similarities across all the structures, we compared the Apo *versus* the TSA-bound structures. The RMSD values were as similar for each group whether compared to an Apo or a TSA-bound structure (Figure 2C), providing no indication of a significant conformational change.

**Figure 2.**
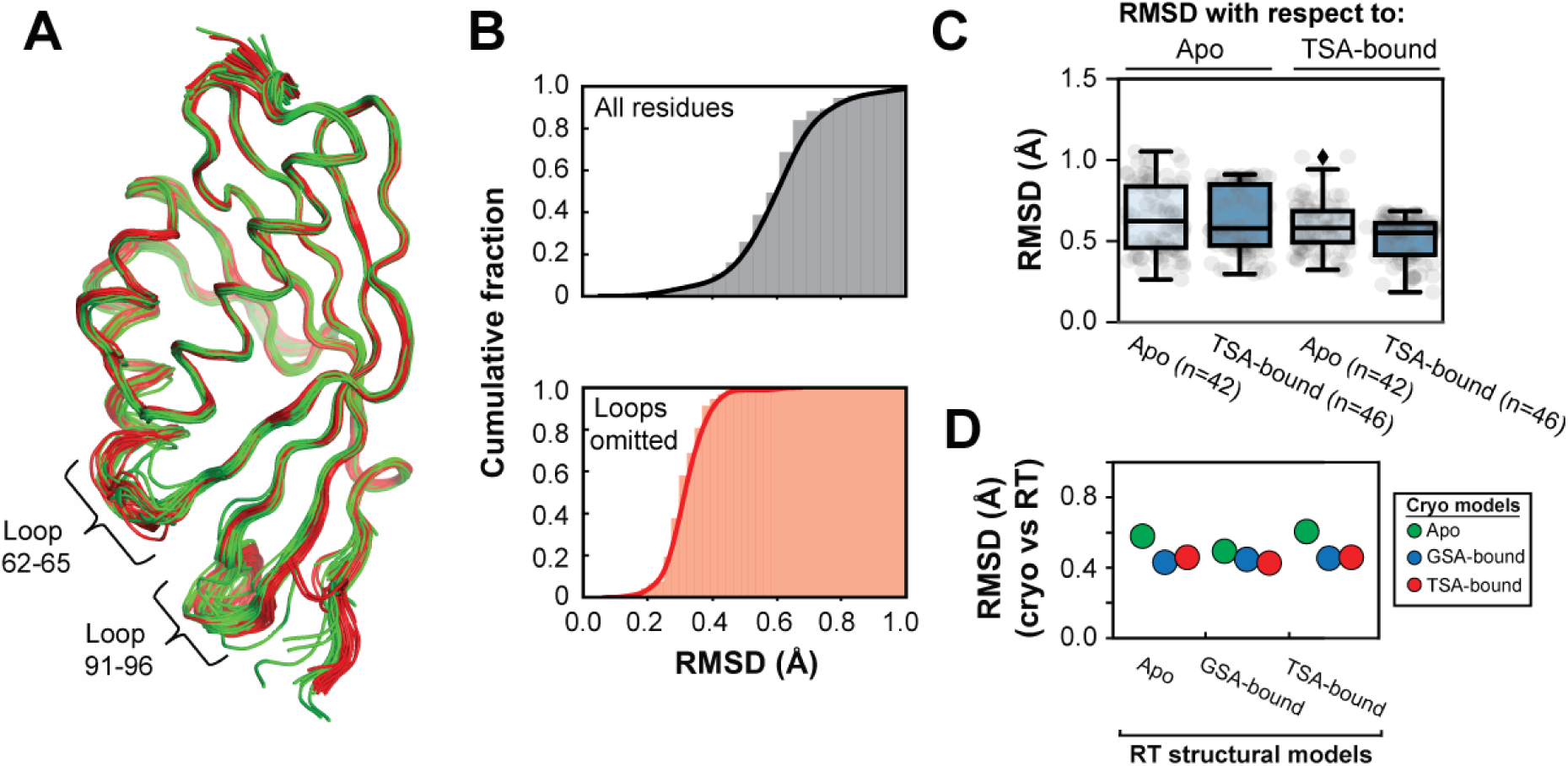
Comparison of KSI crystal structures indicates no structural changes during catalysis. (A) Alignment of all 94 KSI crystallographically-independent molecules from the 45 PDB crystal structures (in ribbon): Apo (green), GSA-bound (blue) and TSA-bound (red). The positions of the two flexible loops are noted. Throughout we evaluate individual monomers of the KSI dimer (see Materials and Methods). (B) Cumulative fraction of backbone RMSDs of all KSI structures *vs.* the highest resolution structure (WT-TSA bound, PDB 1OH0) shown for the entire sequence (top, grey) and excluding the 62–65 and 91–96 loops (bottom, orange) (also see Figures S2 and S3). RMSDs at 0.5 cumulative fraction are 0.6 Å for the entire sequence (top, grey) and 0.3 Å excluding loops (bottom, orange). (C) Alignment of all Apo (n = 42, light blue boxes) and all TSA-bound (n = 46, dark blue boxes) KSI monomers on the highest resolution Apo (PDB 3VSY) and TSA-bound (PDB 1OH0) KSI crystal structures. RMSDs are for the entire sequence and averaged over the two monomers of the KSI dimer. The boxes show the quartiles of the dataset and the whiskers denote the entire distribution. (D) KSI structures obtained from cryo-cooled and RT crystals are highly similar. Backbone RMSDs of Apo, GSA-bound and TSA-bound KSI single-conformation models obtained at RT (280 K, this study; x-axis) *vs.* the highest-resolution cryo model (100 K, from the PDB; y-axis; RMSD with the cryo Apo structure (PDB 3VSY) in green; with the GSA-bound cryo structure (PDB 5KP4) in blue; and with the TSA-bound cryo structure (PDB 1OH0) in red. (See also Figure S4 and Tables S4-5 for details.)

While the KSI cryo X-ray structures furnished 42 Apo states and 46 TSA-bound states to make comparisons and to build conformational pseudo-ensembles for each state, there was only a single GSA-bound structure (Tables S1 and S2). We collected RT X-ray diffraction data for KSI bound to a ground state analog at 280 K and analogous data for Apo KSI and TSA-bound KSI to allow direct comparisons and to remove potential effects from cryo-cooling (Table S3) (52, 55, 56, 59). The resultant structures provided additional support for the absence of conformational changes through the reaction cycle (Figure 2D, Figure S4), allowed us to compare dynamics across states representing KSI’s reaction coordinate (Figure 1A), as described in the following section, and demonstrated that our conclusions were not significantly affected by cryo-cooling.

Overall, the high structural concordance for different KSI states suggests that there are at most small structural differences through its reaction cycle.

### Evaluating the KSI conformational landscape through its catalytic cycle

While there is little change in overall structure across states representing KSI’s reaction coordinate, the extent of conformational heterogeneity (i.e., the diversity of conformations) within each state can change; new interactions that are formed and steric constraints that are introduced can change the enzyme’s conformational landscape without altering the average structure. Whether such changes occur is determined by the relative strengths and shapes of the local potentials that constrain groups, and in this case whether the conformational landscape is dominated by interactions within the protein fold itself or is substantially altered by the added ligand interactions.

We built Apo and TSA-bound pseudo-ensembles for KSI, based on 42 and 46 cryo X-ray structures, respectively (Table S2; Figure 2A and Figure S1). In this approach, each crystal structure (including ones with point mutations and different crystallization conditions) is considered to correspond to a local minimum on the native potential energy surface (50, 51, 63). The degree of motion extracted from pseudo-ensembles has been shown to agree well with estimates of motion from solution NMR (50). The individual KSI structures used ranged in resolution from 1.1 to 2.5 Å (Figure S1A and Table S1) and neither inclusion of only high-resolution structures (≤2.0 Å) nor random omission of structures substantially altered the analyzed ensemble properties (Figures S5-7).

The single GSA-bound KSI cryo structure prevented us from building a ground-state pseudo-ensemble. Thus, to evaluate conformational heterogeneity with a bound ground state analog, we built multi-conformer models from our RT X-ray data, and to provide direct comparisons to Apo- and TSA-bound KSI we built analogous multi-conformer models for these species (Table S3). As the overall KSI structures in each state were indistinguishable at 100 and 280 K, we also collected diffraction data at 250 K, slightly below room temperature but above the average glass transition temperature, where our crystals diffracted to approximately 0.2-0.3 Å higher resolution than at 280 K (Table S3). We used the 250 K data herein as higher-resolution data provide more information for multi-conformer modeling (54).

#### Evaluating and comparing conformational heterogeneity

Above we compared KSI structures overall, via RMSDs. Here we evaluate conformational heterogeneity residue-by-residue and then compare the heterogeneity of each residue across the KSI states. For our pseudo-ensembles, we assayed backbone and side chain positioning *via* Cα and Cβ, respectively, by defining an atomic mean deviation (MDev) parameter (see Materials and Methods). Briefly, for a given atom in a structure, the MDev describes the average displacement of equivalent atoms within the ensemble of structures, with lower and higher values representing smaller and larger positional fluctuations, respectively, corresponding to less or more conformational heterogeneity.

In both the Apo and TSA-bound pseudo-ensembles, MDevs for the backbone (Cα) and side chains (Cβ) were highly similar, below 0.5 Å, with exceptions only in the 62–65 and 91–96 loops, which showed the largest conformational heterogeneity (Figure 3A,B, Figure S9A, S10A, S11A-B). The MDevs for the catalytic residues were below average (dotted line) and on the lower end of observed values, and the MDevs for substrate binding residues were close to the average (Figure 3A). Thus, our ensemble data and analyses indicate that positioning does not substantially increase with transition state-like interactions and suggests that the active site is sufficiently preorganized to efficiently carry out catalysis. Nevertheless, there is not extreme or unusual positioning in the active site (see also “*Determining the extent of positioning of catalytic residues”* below).

**Figure 3.**
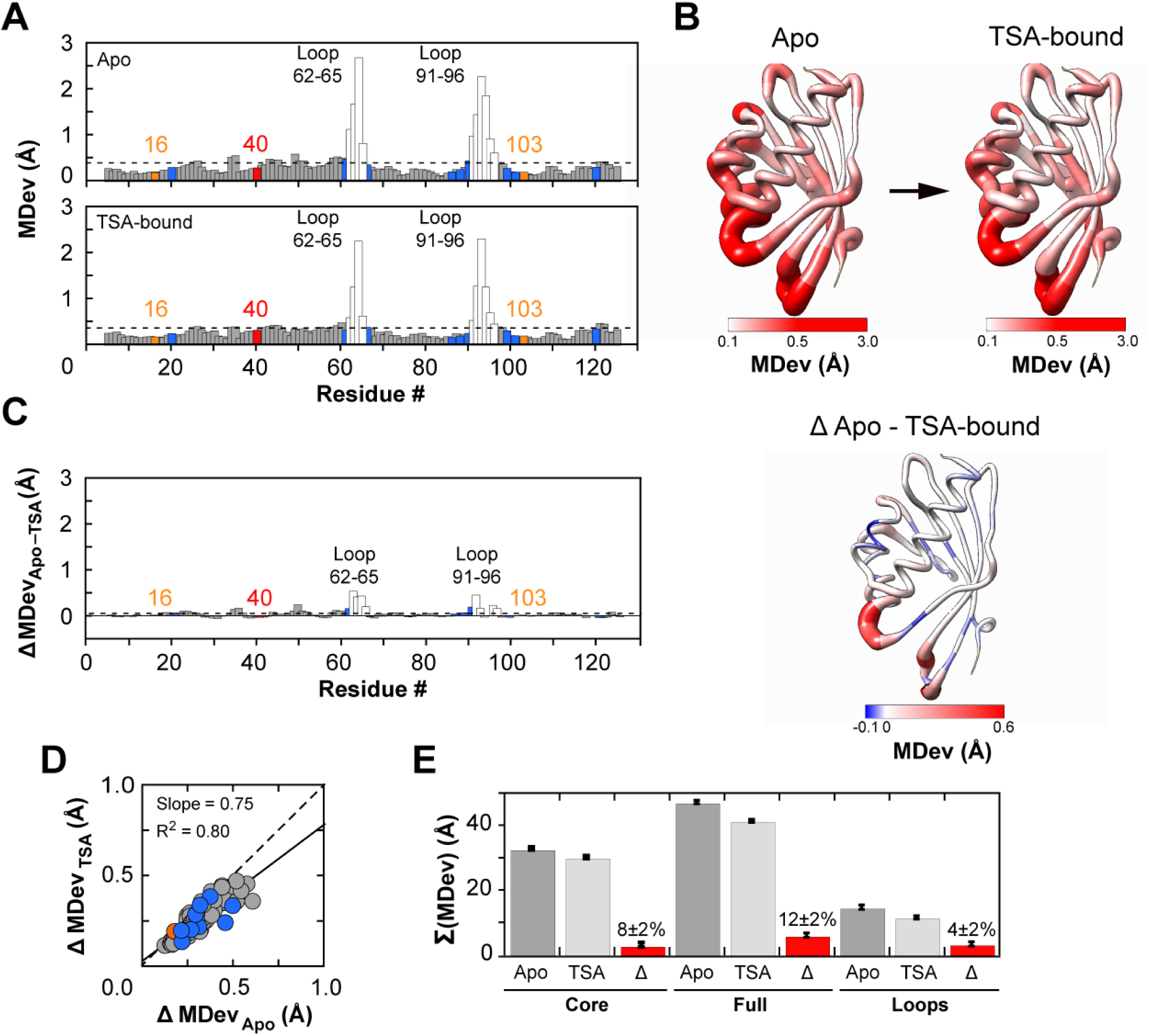
Assessing conformational heterogeneity through the KSI catalytic cycle via pseudo-ensembles. (A) Cα MDevs for KSI Apo (top) and TSA–bound (bottom) states. Dashed lines represent the average MDev. The flexible 62–65 and 91–96 loops are shown as white bars, Y16 and D103 in orange, D40 in red, and binding residues in blue. (B) Worm representation of the MDevs from (A) on the 3D structure of KSI. The thickness of the worm representation follows the color code (from white (MDev ≤ 0.1 Å) to red (MDev ≥ 0.5 Å)). (C) The difference Cα MDev values between the Apo and TSA-bound states (MDevs_Apo-TSA_), such that positive values indicate lower MDevs for the TSA-bound state; the bar plot shows the difference MDev values as a function of the protein sequence (left) and the worm representation shows the difference MDev values represented on the 3D structure of KSI (right) as in panel (B) but with a more sensitive scale to highlight the changes. (D) Correlation plot of Apo and TSA-bound Cα MDevs (excluding loops 62–65 and 91–96). The dashed line of slope 1 represents the expectation for no difference in average MDevs between the two states. Similar results were obtained when loops were included (see Figure S7). (E) Sum of Cα MDevs for Apo (dark grey bars), TSA-bound (light grey bars) and their difference (Δ, red bars). Errors were estimated using bootstrap analysis and error propagation (see Figure S8 and Materials and Methods). Side chain MDevs (using Cβ) gave analogous results (Figures S10 and S11).

To assess changes in conformational heterogeneity through the reaction cycle, we first compared Apo state MDevs to those in the TSA-bound state. The MDev values were similar across the entire structure, as seen qualitatively by comparison of the panels in Figures 3A and 3B and quantitatively in Figure 3C by the difference in MDev between the states and by the strong correlation of the MDev values for the Apo and TSA-bound states (Figure 3D (Cα R^2^ = 0.80), Figure S10D (Cβ R^2^ = 0.76), Figure S7). Nevertheless, the slope of this correlation was less than 1 (R^2^ = 0.75), crudely suggesting an overall dampening of ∼25% in conformational heterogeneity of the enzyme core upon binding of the TSA. A modest dampening in the TSA complex is also supported by smaller MDevs with the TSA bound, with an average reduction (ΔMDev_Apo-TSA_) of 0.05 Å per residue (Figure 3C and Figures S10B, C and S11C, D). We obtained similar results for a homologous KSI (KSI_homolog_) from another organism, despite only 32% sequence identity. For KSI_homolog_, there were fewer but nevertheless a sufficient number of available cryo X-ray structures (n = 42; Figure S12).

To evaluate conformational heterogeneity from our RT multi-conformer models, we calculated crystallographic disorder parameters, (1-S^2^) that report on local conformational heterogeneity by capturing bond vector motions and that agree well with solution NMR measurements (Figure S13) (64). We used (1-S^2^), rather than MDevs, because our RT multi-conformer models contain additional information within each state that is captured by (1-S^2^) and because the limited number of conformational states in multi-conformer models limits the utility of MDev comparisons. (1-S^2^) values range from 1, for a completely unrestrained bond vector, to 0, for completely rigid bond vector. Even though (1-S^2^) and MDev are different measures of heterogeneity, we observed the same reduction in overall heterogeneity of 12% (Figure 3E and Figure S15B). We also obtained information about heterogeneity in the KSI•GSA complex from RT X-ray data, information that cannot be extracted from the single cryo KSI•GSA structure (Table S3, Figure S13-15). Comparisons of the KSI•GSA complex to the Apo and TSA-bound forms revealed that most of the modest reduction in heterogeneity occurs upon formation of the GSA complex (Figure S15).

The 91–96 loop interacts with the substrate and the 62–65 loop interacts with the 91–96 loop but does not interact with the substrate directly. We therefore anticipated that these loops might undergo a conformational dampening upon ligand binding. Nevertheless, the changes are modest (Figure 3C, Figures S9B and S10B), as are the functional effects; mutation of W92, the loop residue making direct substrate contact, increases *K*_M_ only two-fold and decreases *k*_cat_ less than two-fold (65). Further, a substrate with only a single ring and thus lacking the remote ring that is contacted by the 91-96 loop has a *k*_cat_ value within two-fold of the full substrate (66). Thus, the loop is not tightly coupled to the conformational heterogeneity or function of the catalytic residues, and the absence of significant loop effects on catalysis of bound substrates is consistent with the absence of substantial conformational changes or dampening elsewhere in the active site.

In summary, analysis of cryo pseudo-ensembles for KSI from two organisms and RT X-ray data for one of these provide evidence for active site organization resulting predominantly from interactions within the folded protein, with only modest dynamic adaption along the reaction coordinate due to the additional interactions with bound ligands.

### Determining the extent of positioning of catalytic residues

Proposals for the origin of enzymatic power universally invoke positioned catalytic groups in enzyme active sites, relative to their diffusive uncorrelated motions of the same groups free in solution (9, 11–19, 21–23). The analyses above indicate that there are minimal changes in KSI conformational heterogeneity, including the catalytic residues, for species mimicking states along the reaction coordinate, but do not answer the question of how positioned these catalytic groups are; indeed we lack this information for any enzyme.

Given the enormous catalytic potential from positioning—to overcome entropic reaction barriers and enthalpically destabilizing reactants relative to transition states—one might imagine that enzymes have evolved to especially constrain their catalytic groups. For KSI and numerous other enzymes, a potential catalytic mechanism involves a precisely positioned oxyanion hole that distinguishes ground state carbonyl groups from oxyanionic transition states and intermediates (4, 67–75). Conversely, there are cases where motions are clearly required. For example, the *cis*-aconitate intermediate of aconitase swivels from one bound state to another, while attached to aconitase’s FeS cluster via two carboxylate groups, to allow reaction at different positions and thereby the isomerization of citrate to isocitrate (76, 77). While our current knowledge suggests that molecular gymnastics this dramatic are rare, many enzymes, including KSI, proteases and isomerases, use a common set of residues to catalyze more than one reaction step, presumably a result of evolutionary parsimony that necessitates some degree of dynamics (Figure 1A; (78, 79)).

To assess the degree of positioning of KSI’s catalytic residues, we constructed a reduced pseudo-ensemble with 54 KSI molecules, excluding structures from our overall (full) pseudo-ensemble with mutations directly to the residues under analysis and mutations previously identified to alter the positioning of these residues (Table S2). From our RT X-ray data, we created an ensemble by combining the multi-conformer models for the KSI Apo, GSA-bound, and TSA-bound states, given their highly similar overall conformational heterogeneity and their high residue-by-residue similarities (Figures S13, S15). The conformational heterogeneity inferred from the reduced pseudo-ensemble and the RT X-ray ensemble correlated well with the conformational heterogeneity from the full pseudo-ensemble, suggesting that overall ensemble information is retained (for the reduced pseudo-ensemble and the RT X-ray ensemble: R^2^ = 0.98 and R^2^ = 0.84 for Cα and R^2^ = 0.98 and R^2^ = 0.87 for Cβ, respectively; Figures S17-18).

To test whether catalytic groups are particularly constrained, we compared the MDev values for the catalytic atoms of Y16 and D103, the oxyanion hole, and of D40, the general base (Figure 1) with the atoms of chemically similar but non-catalytic residues throughout KSI, and we carried out this comparison with both the reduced pseudo- and the RT-ensembles (Figure 4, Figure S19). The oxyanion hole catalytic groups sit at the lower end of the observed MDev’s, but with values similar to the most constrained non-catalytic groups. The general base oxygen atom of D40 is also not unusually constrained, indeed exhibiting more motion than the equivalent atoms of chemically-similar residues. Further, its noncatalytic oxygen atom has a MDev lower than that for the general base oxygen (Figure 4C-D). These results indicate that catalytic functional groups are not extraordinarily constrained, at least for KSI.

**Figure 4.**
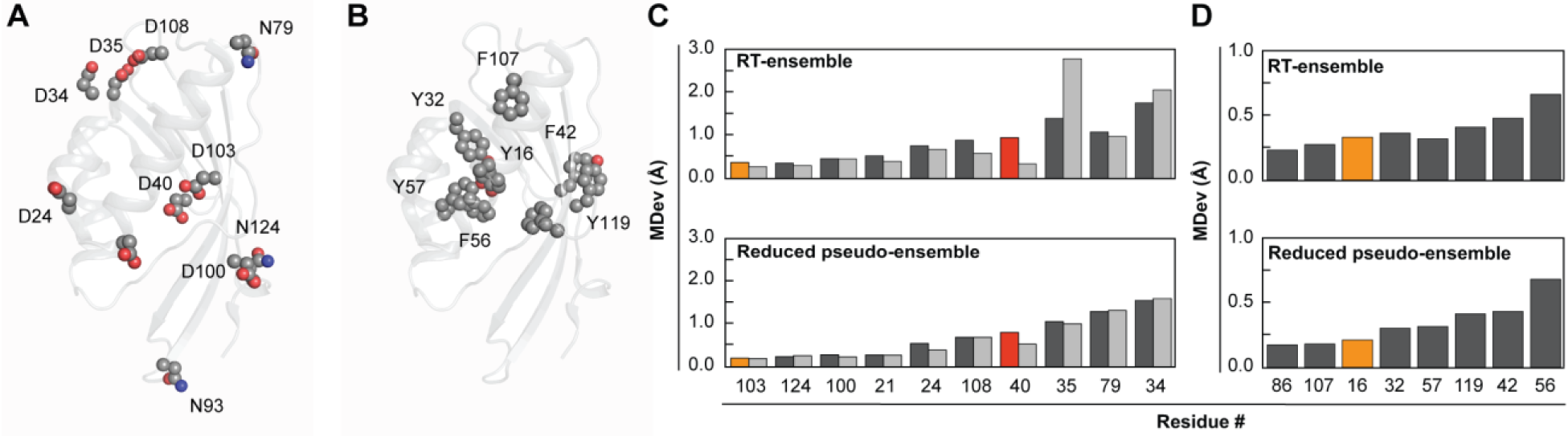
Comparison of positioning of catalytic *vs.* non-catalytic atoms of chemically similar groups. All Asp and Asn residues (A) and Tyr and Phe residues (B) of KSI mapped on the 3D structure (also see Figure S19 for a more complete view and for atom names). MDevs (C) for all Asp and Asn Oδ1 atoms in light grey, O/Nδ2 atoms in dark grey and (D) for all Tyr (OH atoms) and Phe (Cζ atoms), respectively. Y16 and D103 are in orange, and D40 is in red. The upper panels show RT-ensemble results; the lower panels show the results from the reduced pseudo-ensemble. Figure S20 shows analogous results for comparisons of side chain dihedral angles and yield the same conclusions.

We further address side chain positioning and motion in the following sections in the context of testing catalytic mechanisms (immediately below) and of assessing the factors and forces responsible for this positioning (see “*What restricts and permits motions in and around the active site?*”).

### Testing models for KSI catalysis

#### Positioning and general base catalysis

KSI faces a challenge common to many enzymes—a need to efficiently abstract and donate protons and a need to do so at multiple substrate positions. In the face of this challenge, KSI exhibits an effective molarity (EM) of 10^3^–10^5^ M, unusually high for general base catalysis (Figure 5A; (80)). The large rate advantage from the KSI’s general base, D40 (Figure 1), could most simply be accounted for by highly precise positioning of D40, a model we can now evaluate.

**Figure 5.**
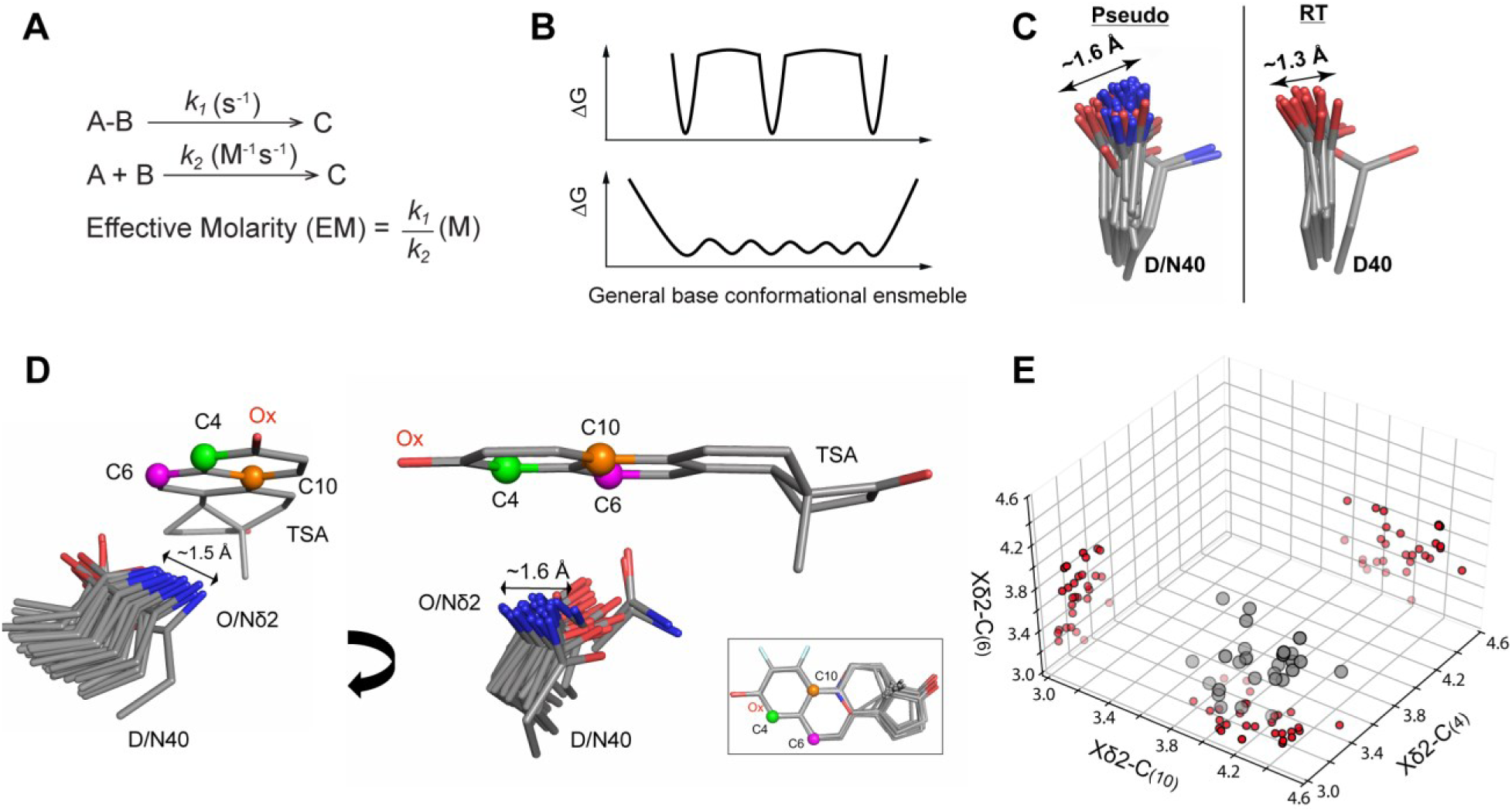
The high effective molarity of the general base in KSI is not due to a highly precise positioning. (A) When a unimolecular reaction (s^-1^) is compared to a bimolecular reaction (M^-1^s^-1^), the ratio of rate constants has units of molar and this value, referred to as the Effective Molarity (EM; in units of Molar, M) (80, 81). The EMs of the general base for the two KSI species investigated herein (KSI & KSI_homolog_) have been estimated to be 10^3^-10^5^ M (82). (B) KSI’s high EM and ability to donate and abstract protons for multiple substrates and at multiple positions could reflect narrow and distinct potential energy wells corresponding to precise positioning with respect to each of the acidic carbons on KSI substrates (top). However, EM reflects a ratio of rate constants, as illustrated in panel (A), and thus can arise without narrow positioning (bottom) with other factors responsible for the high observed catalytic efficiency. (C) The general base (D40) pseudo-ensemble and RT-ensemble. The D40 pseudo-ensemble contains both aspartate and asparagine residues, as asparagine mimics the protonated (intermediate) state of the general base and increases the affinity for TSAs (71, 83). Including only aspartate or only asparagine residues does not substantially alter the range or observed motions (Figure S21). (D) A bound TSA as “seen” by the KSI general base in a TSA-bound ensemble of cryo crystal structures (Table S2). The TSAs (equilenin and various phenols) have been aligned on the A ring with only one (PDB 1OH0) shown for clarity. The carbon positions between which protons are shuffled in KSI reactions are represented as green, magenta, and orange spheres (see Figure 1 and Figure S30 for the reaction mechanisms). The inset shows the aligned TSAs. (E) The distribution of distances between the Oδ2 (Nδ2 when asparagine at position 40) and the three carbon positions from (D) on the bound TSAs in each crystal structure. Projections onto each of the planes reveals a broad distribution rather than sub-clusters at short distances around each of the carbons.

Above we noted that the proton-abstracting oxygen of D40 (Oδ2) is not particularly well positioned, relative to other carboxylate oxygens, based on MDev comparisons (Figure 4). Nevertheless, its MDev value could represent precise positioning in narrow but spatially separate energy wells, with each corresponding to a position for proton abstraction (Figure 5B, top), or it could represent a broader, more continuous conformational ensemble (Figure 5B, bottom). The KSI pseudo-ensemble and RT-ensemble provide evidence for motions on the scale of 1-1.5 Å, with no indication of preferential sub-positions of the general base (Figure 5C and Figure S21). To further investigate this possibility, we aligned the 36 available KSI molecules with bound transition state analogs from X-ray cryo structures (Table S2), which revealed a broad range of general base positions with respect to the bound ligand (Figure 5D). We then determined the distances between the general base catalytic Oδ2 and the TSA carbon positions corresponding to the acidic carbons between which protons are shuffled in different substrates (Figure S30). A three-dimensional plot, with each dimension corresponding to one of these three distances, showed that instead of clustering around each of the three proton abstraction/donation positions, the general base catalytic Oδ2 positions are spread through this three-dimensional space (Figure 5E). We observed a similar broad general base distribution for KSI_homolog_ (Figure S22). In summary, mechanisms other than highly precise local positioning for proton abstraction appear to be needed to account for KSI’s highly efficient general base catalysis (see Discussion).

Implied above, and in numerous discussions of enzyme catalysis, is a need to balance positioning and flexibility. For example, it has been suggested that loop closure in certain enzymes can be rate limiting, so that mobile loops need sufficient flexibility to not limit the reaction rate but also sufficient rigidity to position the substrate and catalytic groups for the chemical reaction step (84– 86). Other discussions of the rigidity–flexibility continuum are often less precise and thus difficult to draw specific predictions from and assess. For KSI, we can make the prediction that increasing *or* decreasing positioning of its general base may reduce catalysis^1^: excessive rigidity would prevent access to all three proton transfer positions and excessive flexibility would result in a large fraction of unreactive conformations.

#### Ensemble-function analysis reveals a balance between general base positioning and flexibility

To evaluate the above predictions, we carried out an “ensemble-function” analysis, comparing structural ensembles for two KSIs that differ in the groups interacting with their general base in conjunction with new and prior functional data.

As expected, mutation of many residues surrounding D40 substantially reduce catalysis and also substantially increased D40 mobility, as previously evidenced by reduced electron densities in mutant cryo-X-ray data (87, 88). Mutations around a catalytic site often compromise catalysis and disrupt positioning, and these ubiquitous results provide qualitative evidence for the role of positioning in catalysis. However, for KSI there is also evidence for reduced catalysis from *restricting* motions.

The KSI focused on herein positions D40 through a hydrogen bond between its non-catalytic oxygen (Oδ1) and the side chain of W120 (Figure 6A). The hydrogen bond from W120 to Oδ1 appears to act as a pivot point that allows conformational exploration by the more distal Oδ2 (Figure 6B-D). In contrast, KSI_homolog_ replaces W120 and its hydrogen bond with a phenylalanine and an anion-aromatic interaction (Figure 6E). As expected, the less directional anion-aromatic interaction allows greater freedom of motion for D40 (Figure 6F, G).

**Figure 6.**
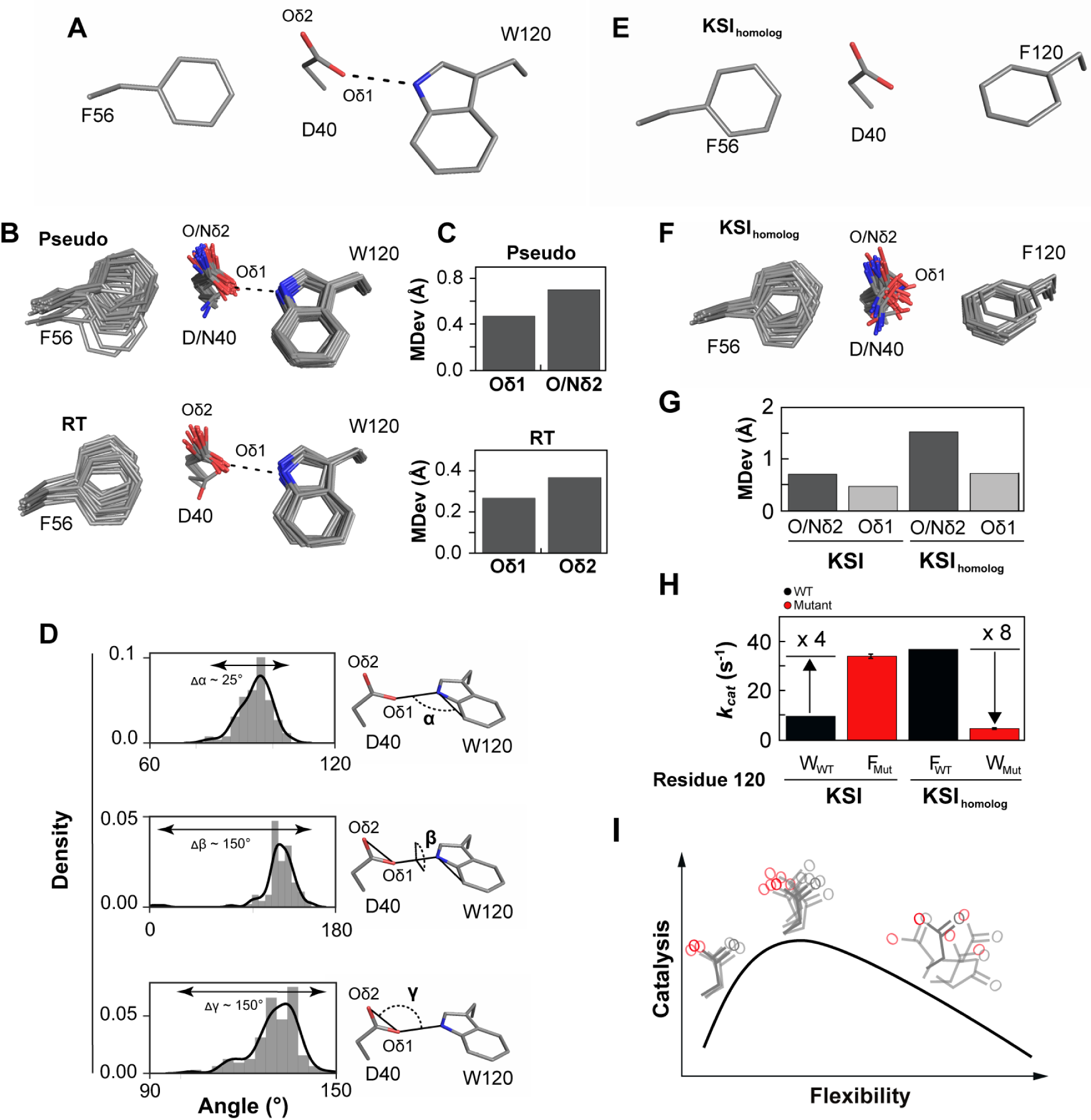
Ensemble-function analysis of general base catalysis for two KSI homologs. (A) Anion-aromatic and hydrogen bonding interactions with F56 and W120 sidechains, respectively, implicated in general base positioning in KSI (illustrated using PDB 1OH0). (B) The full pseudo-ensemble (top) and RT-ensemble (bottom) of the general base (D40), F56, and W120 sidechains. (C) MDevs of the non-catalytic (Oδ1) and catalytic Oδ2 (Nδ2 when asparagine at position 40) oxygen atoms of the general base from the full-pseudo (top) and the RT-ensemble (bottom). (D) General base D40/W120 angular sidechain orientations obtained from the pseudo-ensemble. (E) Anion-aromatic interactions with F56 and F120 sidechains implicated in general base positioning in KSI_homolog_ (illustrated using PDB 1OHP, KSI numbering used for both KSI and KSI_homolog_). (F) The KSI_homolog_ pseudo-ensemble (Table S21). This pseudo-ensemble excluded structures with mutations that alter the chemical nature of the general base (e.g., histidine or glycine) and mutations in the loop carrying the general base (residues 40-44, ‘general base loop’, KSI numbering), as these mutations alter the general base positioning (Table S21) (88). (G) MDev values for the D40 catalytic Oδ2 (dark grey bars, including asparagine Nδ2) and non-catalytic Oδ1 (light grey bars) obtained from the pseudo-ensembles in panels B and F. (H) Replacing the D40-W120 hydrogen bond in WT KSI with an anion-aromatic interaction (W120F, as in WT KSI_homolog_) results in a 4-fold rate increase (Table S58). The equivalent replacement of the D40-F120 anion aromatic interaction in WT KSI_homolog_ with a hydrogen bonding interaction (F120W, as in WT KSI) results in a 8-fold rate decrease (Table S58). (I) Conformational ensemble model for optimal general base catalysis. Optimal flexibility enables the general base to shuffle protons between different positions in different substrates (middle); reducing motion (left, e.g., when F120 in KSI_homolog_ is replaced with W120 in KSI) reduces catalysis, as does increasing motion (right, e.g., from mutations in the loop carrying the general base (87, 88)).

Intriguingly, KSI_homolog_ is 4-fold more active than KSI, and we wondered whether this increased activity might be linked to motions of D40 (89). Consistent with this possibility, mutation of F120 to tryptophan in KSI_homolog_ reduced activity 8-fold (Figure 6H). Whereas deleterious effects are common and can be difficult to trace mechanistically, favorable mutations in natural enzymes are rare. Remarkably, the W120F mutation in KSI *increases* its activity 4-fold (Figure 6H), and with this increase, its activity now matches that of KSI_homolog_.

Our ensemble–function analysis suggests that while general base flexibility appears required for function, more freedom of motion does not uniformly enhance catalysis; rather, there is presumably an optimal balance between allowing and limiting conformational motion, with too much motion reduces catalytic efficiency, as has been observed many times, but also that too-little motion hinders catalysis, presumably by lowering the occupancy of D40 in at least some of its reactive poses (Figure 6I).

#### Positioning and oxyanion hole catalysis

We expect, and observe, large deleterious effects when Y16 and D103, the hydrogen bond donors of KSI’s oxyanion hole, are replaced with large hydrophobic residues that cannot stabilize the negative charge build-up on the substrate carbonyl oxygen (Figure 1A;, Table S59) (89, 90). Such results are common but do not answer the critical question of *how* these catalytic groups provide rate advantages relative to uncatalyzed reactions in aqueous solution (91). Our ensemble data allow us to evaluate several mechanisms proposed for how replacing hydrogen bonds donated by solution water molecules with enzymatic hydrogen bond donors affords KSI its oxyanion hole catalytic advantage.

##### Assessing geometric discrimination

A widely adopted perspective on enzyme catalysis holds that the catalytic power of enzymes can be understood in terms of transition state complementarity, as opposed to the complementarity for a ground state, and this perspective is central to the interpretation of mutational effects and efforts to engineer new enzymes (13, 22, 92–95). Preferential transition state stabilization on geometrical grounds—e.g., discrimination between an sp^2^ ground state and sp^3^ transition state, has been proposed for oxyanion holes of proteases and isomerases, including that of KSI (e.g. 4, 67, 69–72, 74, 75, 96). Such geometric discrimination would require highly precise positioning of the hydrogen bond donors, so that hydrogen bonds can be suboptimal in the ground state and optimal in the transition state.

While an individual structure may show optimal or suboptimal positioning for either ground state or transition state stabilization, it is the conformational ensemble that defines the positioning and relative stabilization. Analysis of our KSI conformational ensembles indicate positioning within the oxyanion hole on the scale of ∼0.5-1 Å (Figure 7A, Figure S23), with hydrogen bonds made from a wide range of orientations (Figure 7B, Figure S24). Thus, sp^2^ *versus* sp^3^ discrimination is unlikely in KSI. This conclusion is further supported by additional analyses below.

**Figure 7.**
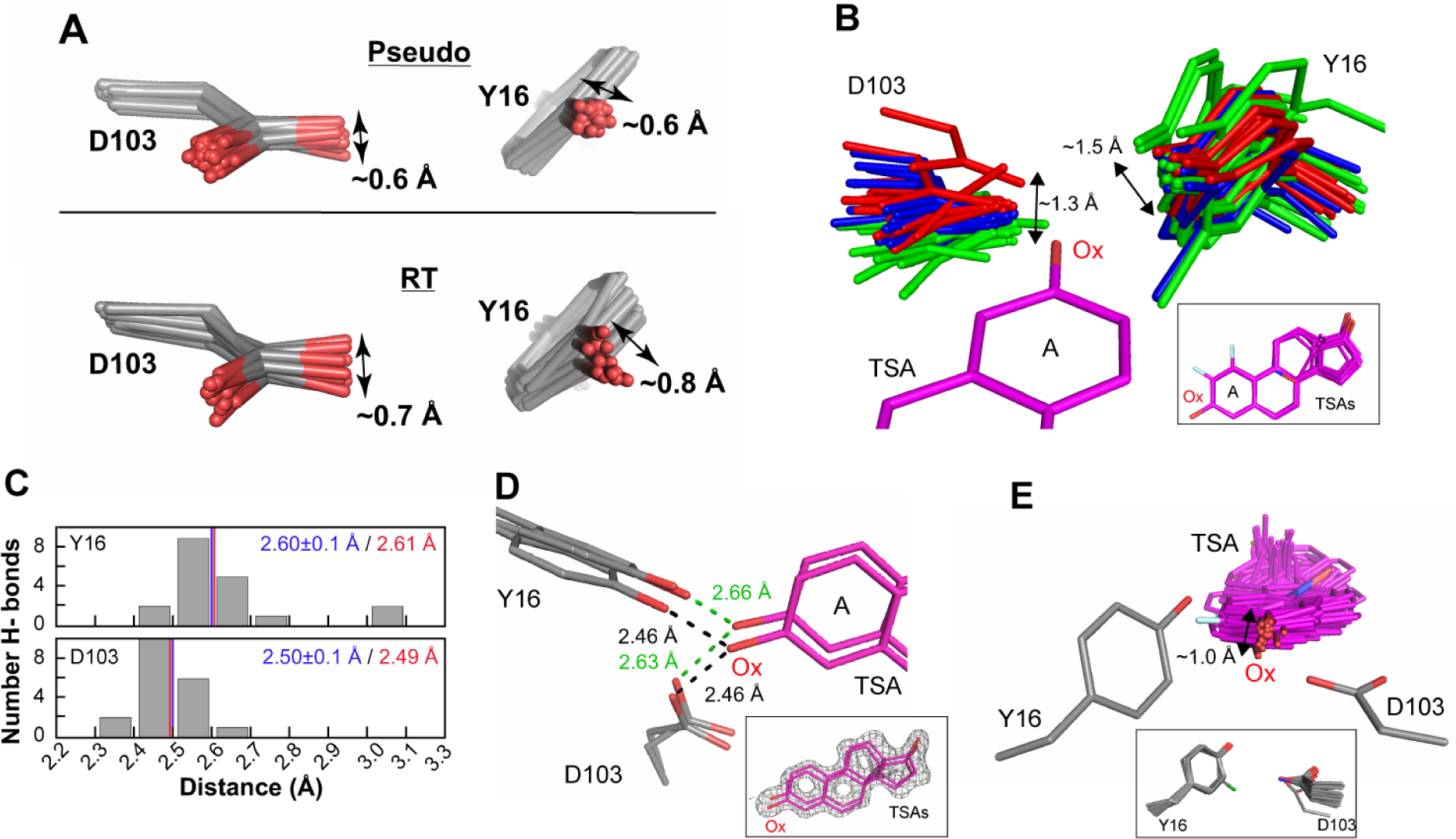
The KSI oxyanion—oxyanion hole conformational ensemble. (A) The oxyanion hole (Y16 and D103) pseudo-ensemble (top) and RT-ensembles (bottom). Phenylalanine residues at position 16 are omitted, and chlorine atoms in chemically-modified tyrosine residues have been omitted for clarity (see also Figures S16, S23). (B) The KSI oxyanion hole as “seen” by 36 bound TSAs (equilenin and various phenols) in cryo crystal structures (Table S2). Structures have been color-coded in three groups (green, blue and red) according to the D103 position in space relative to the TSA oxyanion (Ox). All TSAs have been aligned on the A ring but only one (PDB 1OH0) is shown for clarity. The inset shows the 36 aligned TSAs. (C) Distribution of Y16 (top) and D103 (bottom) hydrogen bond distances from an ensemble of KSI crystal structures of variants with WT-like activity and with the same bound TSA (equilenin, n = 19, Table S2). The mean hydrogen bond lengths and their standard deviations from the cryo crystal structure distances are shown in blue. The hydrogen bond distances obtained by solution ^1^H NMR for D40N KSI bound to equilenin are show in red (see Figure S25 for ^1^H NMR spectra from which distances are obtained). (D) The TSA-bound RT multi-conformer model shows that oxyanion hole Y16 and D103 and the bound TSA can make hydrogen bonds from different orientations. While multi-conformer models do not allow to unambiguously identify each hydrogen bonded sub-states, possible hydrogen bond lengths between Y16/D103 and the TSA are within the range obtained by cryo-structures and solution ^1^H NMR. The inset shows the TSA (purple sticks) and the experimental electron density (grey mesh, contoured at 1σ). (E) The bound TSAs from (B) as “seen” by the hydrogen bonding oxygens of Y16 and D103. Y16 and D103 have been aligned such that their hydrogen bonding oxygens overlay but only one Y16/D103 set (PDB 1OH0) is shown for clarity. The inset shows all aligned Y16 and D103 hydrogen bonding groups.

##### Assessing hydrogen bond properties

NMR studies have established that the hydrogen bonds to KSI-bound oxyanions are short (71, 97–102), and it has been suggested that short hydrogen bonds in KSI and other enzymes are conferred by properties of the enzyme and contribute substantially to catalysis (103–107). For example, these short hydrogen bonds might arise from precise geometric constraints of the oxyanion hole hydrogen bond donors. However, our ensembles described above reveal a wide range of conformational states of the oxyanion hole that donate hydrogen bonds to the bound oxyanion (Figure 7A, B, Figures S23 and S24), and the absence of strong geometrical constraints is further evidenced by lack of coordination between the two oxyanion hydrogen bond donors (Figure 7B and Figure S16). In addition, the highly similar hydrogen bond lengths across the ensemble sub-states provides no indication of hydrogen bond properties driven by conformational properties of the enzyme. Seemingly equivalent hydrogen bonds can be made from different oxyanion hole hydrogen bond donor orientations and the bound TSA can accept hydrogen bonds from a variety of bound orientations (Figure 7B-E). Indeed, the oxyanion hole hydrogen bond lengths match those observed for these donor/acceptor pairs universally, in small molecules, other proteins, and across a range of solvents (Figure 7C) (100, 108, 109). These results together suggest that the donor/acceptor electron densities determine the observed hydrogen bond lengths in KSI and that the internal protein forces are not sufficient to measurably alter these intrinsic hydrogen bond properties.

Independent experiments dissecting the energetic properties of the oxyanion hole hydrogen bonds suggest that KSI accrues a catalytic advantage by utilizing hydrogen bond donors with higher partial positive charge than the hydrogen atoms of water, without significant accentuation from the protein environment (100, 110, 111). The ensemble results provide no indication of additional catalytic advantages from geometric discrimination or from higher precision in positioning, relative to water hydrogen bond donors in solution. Indeed, the multiple oxyanion poses appear to amplify the range of conformational states of bound substrates and intermediates, perhaps acting synergistically with the motional freedom of general base (D40) to enhance KSI’s ability to carry out proton abstraction and donation at multiple positions (Figure 5C-D, Figures S21, S22 and S30). A corollary of this model is that mutations in and around the oxyanion hole that sufficiently alter its ensemble could alter the relative rates of reaction steps for proton transfer at different positions of a substrate or change substrate specificity, a prediction that remains to be tested.

### What restricts and permits motions in and around the active site?

Our KSI ensembles provide a window into molecular behaviors that are central to catalysis and allow us to begin to evaluate the interactions and forces that are responsible for positioning. This information will be needed to deeply understand enzyme function and the pathways by which enzymes have evolved, and will be needed to most effectively design new enzymes.

Traditional X-ray crystallographic models provide many insights, including identifying hydrogen bonds and hydrophobic contacts that may constrain motions. It has been noted that hydrogen bonds are more directional and thus more restricting than hydrophobic interactions in isolation (112– 117). Our analysis of what restricts and allows motion of KSI’s general base supports this view, as anion-aromatic interactions take the place of more common hydrogen bonding interactions with the carboxylate base to enhance conformational excursions, and replacement of one of the anion-aromatic interactions with a hydrogen bond restricts motion and lowers catalysis (see Figure 6).

Nevertheless, in the crowded idiosyncratic environment of a protein interior, favored conformational states and the breadth of their distributions will be determined by multiple energetic contributors, that include hydrogen bonds, van der Waals interactions, steric repulsion, and bond angle preferences, all of which need to be integrated over all the allowed states to understand the ensemble of conformations that are present and the preferences among them. Our ensemble data provide insights into the forces responsible for positioning, and, in certain instances, provide testable models for the interactions and forces that restrict and permit local motions.

To describe packing and van der Waals interactions in an accessible form, we defined a packing distance between surrounding residues to the catalytic groups, Y16, D103, and D40, yielding, for each structure from the pseudo-ensemble, a single distance of closest approach between atoms of the catalytic side chains and surrounding groups (Tables S42-48). These distances, plotted together, provide ensemble-level local packing information (Figure 8A-C). While our ensemble contains more detailed information, the simplicity of this representation facilitates interpretation, and more detailed analyses can be carried out in the future, especially to evaluate predictions from molecular dynamics simulations and other models.

**Figure 8.**
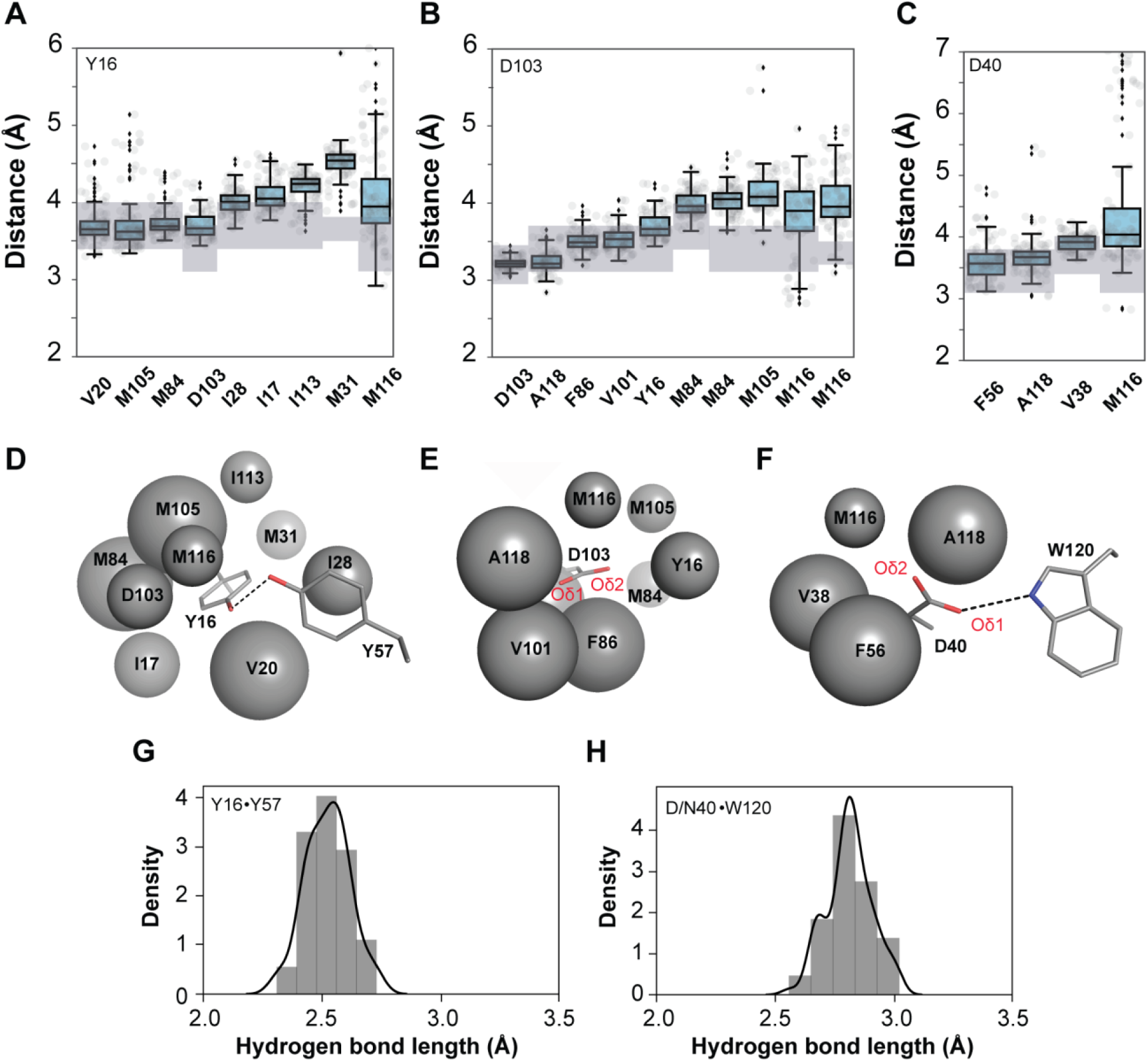
Ensemble analysis to evaluate the packing and interactions around the KSI catalytic groups. Ensemble distances between Y16 (A), D103 (B), and D40 (C) and their surrounding packing groups from all 94 independent KSI molecules from the 45 cryo crystals structures available in the PDB (Table S1). The boxes show the quartiles of the dataset and the whiskers extend to include the statistical distribution. The closest atoms making van der Waals interactions were identified and distances between specific atom pairs were measured (see Tables S42, 44, 46 for the list of atom pairs); hydrogen bonding lengths are presented separately (G, H). Two sets of distances are reported for M84 and M116, as two distinct atoms were within similar distances. Van der Waals radii (r_vdw_, shaded rectangles) are represented as a range because of uncertainty introduced by the absence of hydrogen coordinates in the X-ray structural models and because the oxygen r_vdw_ is orientation-dependent (see Table S48 for a list of r_vdw_ used here). D, E and F schematically depict the results from panels A, B and C, respectively, with the packing atoms represented as spheres; larger spheres representing more tightly packed surroundings; and Y16, D103 and D40 and the hydrogen bonding groups Y57 and W120 represented in sticks for clarity (PDB 1OH0). (G, H) Histogram of Y16-Y57 and D40-W120 hydrogen bonding distances from structures in A-C (position 40 contains both aspartate and asparagine residues).

#### Y16

Several groups, including M84, are situated within van der Waals contact distance of Y16, whereas M31 is consistently beyond van der Waals contact distance and M116 is highly variable in its positioning with respect to Y16 (Figure 8A, D, Figure S27). In addition to these packing interactions, Y16 accepts a hydrogen bond from Y57, and the Y16-Y57 distance is highly conserved (Figure 8G). The observed longer Y16/M31 distances indicate that the collection of states with tight simultaneous packing of all residues is higher in free energy than the states that are observed.

We considered two simple models to account for the absence of close Y16/M31 packing:

*Model I*: Packing is more favorable with residues other than M31 (e.g., M84), leading to the choice to not utilize the available van der Waals interaction energy from packing with M31.
*Model II*: The Y16-Y57 hydrogen bond energy and positioning dominates to constrain Y16 away from M31 and closer to M84.

*Model I* predicts that breaking the Y16-Y57 hydrogen bond will leave Y16 or its replacement residue in place, whereas in *Model II* predicts a rearrangement, most simply to make closer interactions with M31. To test these predictions, we compared KSI sub-ensembles with the Y16-Y57 hydrogen bond to those with mutations that removed this hydrogen bond (Figure 9A, B, Y16-Y57 hydrogen bond intact (grey stick) and ablated (green sticks), respectively; Table S2). Without the Y16-Y57 hydrogen bond, we observed a shift in the Y16 positions, toward M31 and away from M84 (Figure 9C-D, Y16-Y57 hydrogen bond intact (grey histograms) and ablated (green histograms), respectively). These results support *Model II* and an energetic and conformational trade-off between the Y16-Y57 hydrogen bond and what would otherwise be a more symmetrical packing of Y16 with both M31 and M84 (Figure 9A-D). Nevertheless, Y16 becomes more mobile without the hydrogen bond from Y57 (Figure 9E), and a range of Y16 distances is observed with respect to both M31 and M84, suggesting a rather flat local energy landscape in which interactions with M31 and M84 are similar in energy, whereas states rearranged to allow simultaneous packing are disfavored, presumably because the cost of losing other interactions with these side chains is too high. This flat energy landscape is then tilted by the Y16-Y57 hydrogen bond to favor Y16 positions closer to M84.

**Figure 9.**
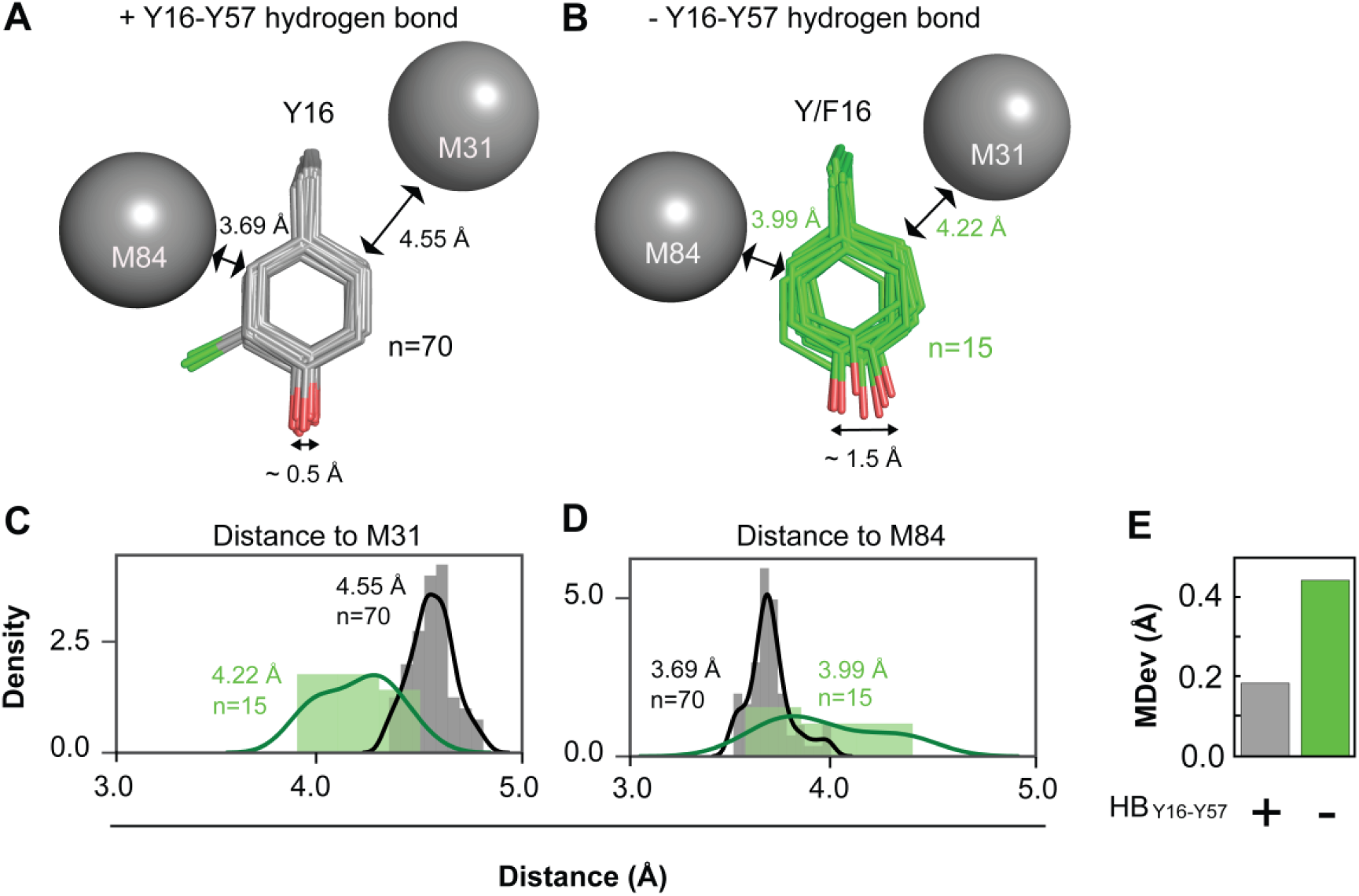
Testing models for the asymmetric packing of Y16. Y16 residues with intact Y16–Y57 hydrogen bond (A, in grey) or Y/F16 residues with the Y16–Y57 hydrogen bond ablated (B, in green) (Table S2). Residues M31 and M84 are shown as grey spheres with the mean distances between M84 and Y16 or M31 and Y16 given (n = 70 and 15 for intact and ablated Y16-Y57 hydrogen bonds, respectively). Chlorine substitutions in some of the tyrosines in A are colored in light green. (C, D) Histogram of the distribution of Y16–M31 and Y16–M84 distances for the ensembles from panels (A) and (B), respectively. (E) MDev for the Cζ atom of the phenyl ring for the ensemble from panel (A) in grey and from panel (B) in green (see Figure S19 for definition of the Cζ atom).

This simple model predicts that mutations in residues surrounding M31 and M84 might free them to allow their simultaneous interaction with Y16 and the consequences of individual mutations can be predicted via molecular dynamics and other models, underscoring the wealth of models that can be generated and tested with ensemble models and data.

Interestingly, the increased flexibility of the ring at position 16 upon removal of the Y16–Y57 hydrogen bond does not result in any rearrangements or significant structural changes in the surrounding residues, suggesting that their positioning is dominated by interactions with residues other than Y16 (Figure S28). Further, neither ablation of the Y16 hydrogen bonding group (e.g., Y16F mutation, Figure S16), nor increased flexibility or apparent mispositioning of Y16 upon ablation of the Y16–Y57 hydrogen bond (Figure S28) appear to impact D103 positioning, supporting the conclusion above that the allowed Y16 and D103 side chain orientations are not coupled (Figure 7B).

#### D103

The observed conformational constraint from the Y16–Y57 hydrogen bond leads to the question of how D103 can be well-positioned in the absence of analogous hydrogen bonding. Analysis of the D103 surroundings revealed particularly close packing of the non-catalytic Oδ1 of D103 with multiple residues: F86, V101, and A118 (Figure 8B, E). Indeed, the RT-ensemble suggested that this oxygen atom may be more restricted than the protonated catalytic oxygen (Oδ2) that sits in the oxyanion hole (Figure 4C). This atypical arrangement in which hydrophobic interactions surround a carboxylic acid oxygen atom (118) is particularly intriguing as it appears to accomplish two objectives: *i.* positioning the catalytic group, and *ii.* increasing its p*K*_a_ so that the carboxylate group remains protonated and can act as a hydrogen bond donor at physiological pH. The protonated D103 side chain provides greater oxyanion stabilization than hydrogen bond donors with higher p*K*_a_ values, such as asparagine, as those side chains have hydrogen bonding protons with lower partial positive charges (Figure 1; (110, 119) see “*Discussion*”). Thus, counterintuitively, interactions of a polar oxyanion atom with hydrophobic groups appear to provide important favorable interaction energy, from van der Waals interactions that sterically constrain the carboxylic acid group for function and disfavor conformational states where the carboxylic acid group can rearrange and be solvated to favor its anionic form. This model is testable—it predicts that mutations that reduce packing will decrease D103’s p*K*_a_ and positioning. Studies of this type will help elucidate dissect the precise relationship between positioning, energetics and function.

#### D40

Above we addressed the anion-aromatic and hydrogen bonding interactions with W120 and F120. The W120 hydrogen bond appears to be more restrictive that the anion-aromatic interaction, as expected, based on the known properties of these interactions. Thus, our ensembles do not reveal new types of forces or interactions, rather manifestations of the fundamental properties of these interactions within more complex environments and over the range of accessible states. Our packing analysis reveals interactions in addition to the W120 hydrogen bond: van der Waals interactions with V38, M84, and A118. Their roles in and effects on positioning and catalysis can be dissected in future experiments that combine functional and ensemble studies of mutants at and around these sites.

Intriguingly, the “other” oxygen of D40, the one involved in proton abstraction and donation, makes an anion-aromatic interaction, with F56 (Figure 6A). The broader potential of this interaction, relative to a hydrogen bond, may help the Oδ2 oxygen atom navigate the range of positions required for proton transfers. It may also destabilize the anionic form of D40, relative to its protonated form, to increase the driving force for proton abstract and thereby facilitate catalysis.

## Discussion

Precise positioning has often been suggested, explicitly or implicitly, as essential for enzyme catalysis and specificity (9, 11–19, 21–23). It is clear that enzymes fold and bind substrates, thereby co-localizing and restricting the motions of catalytic groups and reactants. With pseudo-ensembles from collections of cryo X-ray structures and ensembles from RT X-ray data we can now measure positioning within active sites and test specific and general catalytic models.

Our ensemble data have allowed us to evaluate whether catalytic groups are exceptionally positioned and to define the extent of their positioning in KSI’s active site. We observed that catalytic residues were more conformationally restricted than most other residues, but not extraordinarily so, with the oxyanion hole and general base residues spanning scales of ∼1–1.5 Å (Figures 4-5, 7, Figures S21, S23). Their conformational excursions were asymmetric, down to ∼0.2 Å in particular directions, reflecting the asymmetric protein environment (Figures S21, S23). Furthermore, similar extents of motion were observed for a KSI homolog, suggesting that the extent of positioning and mobility arises from a combination of certain shared structural and functional features (Figures S12, S22).

Geometric discrimination in oxyanion holes, between sp^2^ ground states and sp^3^ transition states, has been proposed for proteases and other enzymes, including KSI (e.g. 4, 67, 69–72, 74, 75). Our analyses of KSI ensembles indicate motions on the scale of 1 Å in the oxyanion hole, a large angular range of hydrogen bonds to transition state analogs, and hydrogen bonds of similar length (Figure 7, Figure S24). These observations suggest that ground state *vs.* transition state geometric discrimination for KSI is unlikely. Instead, KSI’s oxyanion hole appears to generate catalysis by providing *stronger* hydrogen bonds, relative to those from water (110, 111).

KSI exhibits highly efficient general base catalysis, as demonstrated by the effective molarity (EM) of D40, its general base (EM = 10^3^–10^5^ M; Figure 5; (82)). Nevertheless, our ensemble data revealed that D40 exhibits considerable motional freedom, providing evidence against highly precise positioning as the origin of KSI’s efficient general base catalysis and necessitating consideration of alternative models, such as general base desolvation in active site complexes driven by protein folding and ligand binding energy (80, 81, 120)

Many recent enzymatic proposals emphasize dynamics, but these proposals have been difficult to evaluate due to experimental limitations and because enzymes undergo motions for multiple reasons; indeed, physics requires that all atoms are in constant motion above 0 Kelvin. Nevertheless, one frequent catalytic function of active site motions might be successive chemical steps at multiple positions carried out by a single residue, as is the case for KSI’s general base (Figure 1A and Figure S30, D40; (4, 75, 121)). KSI’s general base flexibility is likely essential for function—to efficiently shuttle protons between different positions in substrates and to allow catalysis with different substrates (Figure 1A, 5 and Figure S30). The range of oxyanion hole conformational poses likely also contributes (Figure 7 and Figure S23). Our functional data combined with ensemble analysis suggest an optimal general base flexibility, with too much or too little negatively impacting catalysis (Figure 6I; see below).

### Testing classic proposals for enzyme function

Our KSI enzymes allowed us to test two broad catalytic models: whether enzyme groups, especially active site residues, are largely prepositioned for reaction or whether their conformational ensemble are narrowed as the reaction proceeds (the Dynamic Gradual Adaption Model, Figure 10A) and whether catalytic residues exhibit more precise positioning than other residues (the Dynamic Entatic State Model, Figure 10B).

**Figure 10.**
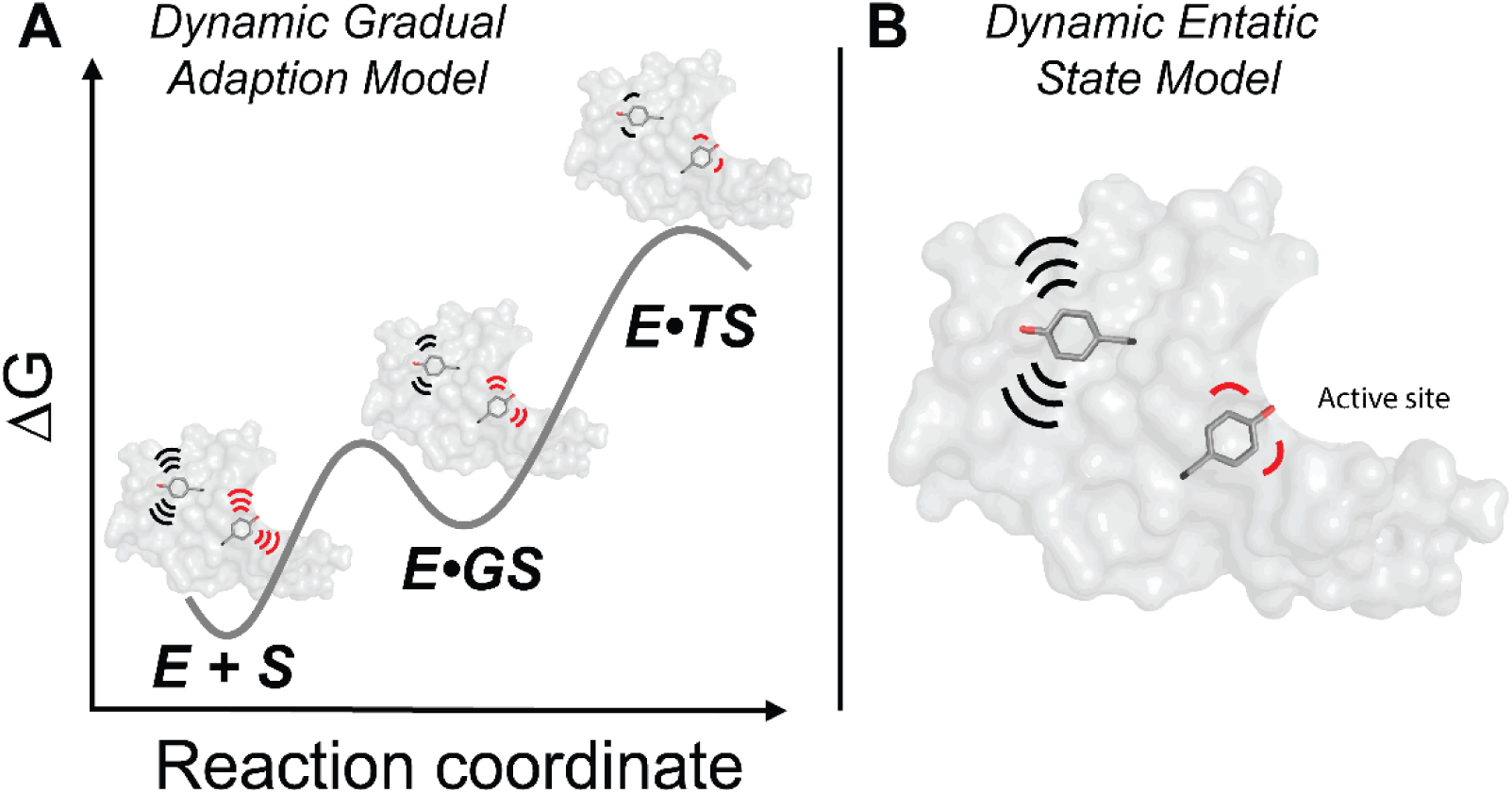
Dynamic models for enzyme catalysis. Each panel shows an enzyme with two highlighted tyrosine residues, a “non-catalytic” tyrosine (black) in the enzyme core (grey) representing the non-catalytic residues, and a “catalytic” tyrosine (red) to represent catalytic residues in the active site. (A) The Dynamic Gradual Adaption Model. Both non-catalytic and catalytic tyrosine residues become more conformationally restricted as the reaction proceeds, but with greater conformational restriction occurring in the transition state. (B) The Dynamic Entatic State Model. Folding energy and local interactions provide greater restrictions and more precise positioning of the catalytic tyrosine (red, representing active site residues), relative to a non-catalytic tyrosine (black, representing non-catalytic residues). The restriction of active site residues (reduced conformational heterogeneity) is “paid for” with folding free energy and is used to enhance catalysis according to this model. In both panels, motions are schematically depicted by the motion lines.

In 1976 Wolfenden proposed that enzymes undergo changes in shape when forming the enzyme-substrate complex and subsequent changes to give maximal complementarity to the transition state where stabilization is greatest, and we refer to this as the “Gradual Adaptation Model” (122). An updated version of this model utilizing a conformational landscape perspective recognizes that enzymes exist as ensembles, with the distribution of states determined by the relative energy, can be described as the “Dynamic Gradual Adaption Model” (Figure 10A). This model implies (re)alignment of enzyme groups upon binding of the ground state and in the transition state, such that the distribution of enzyme states is most altered or narrowed in the transition state, in agreement with the widely discussed perspective that enzymes are most complementary in charge and shape to their transition states, so that stronger transition state interactions may alter or narrow the ensemble (13, 16, 22, 120, 123).

Our results suggest that KSI does not substantially alter or narrow the distribution of enzyme states through its reaction cycle, a finding supported by both cryo pseudo-ensembles and RT X-ray data and common to both KSIs investigated here. As might be expected when new interactions are formed upon ligand binding, there is some change in conformational heterogeneity, and we estimated a modest overall decrease of ∼10–15% (Figure 3 and Figures S10, S12, S15). Nevertheless, the vast majority of KSI’s conformational restrictions, relative to groups moving freely in solution, arise from the extensive interactions that are present in the folded enzyme. Thus, at a minimum, the “Dynamic Gradual Adaptation Model” is not general to enzymes.

A second dynamic model follows from Vallee and Williams’ “entatic state” proposal that active site groups are distorted from their most stable free conformation (by folding free energy) to more closely match the conformational or electronic needs of a reaction’s transition state (124, 125) and the observation by Shoichet and colleagues that active site residues can destabilize enzymes, consistent with evolutionary selection of these residues for catalysis rather than stability (126, 127). Extending from these ideas, more precisely positioned active site relative to other residues could provide a catalytic advantage, while destabilizing the folded enzyme due to the conformational entropy loss from restriction to the most catalytically active conformers (Figure 10B).

We found that catalytic residues were generally more restricted than residues throughout the enzyme, but not extraordinarily so—they tended to match rather than exceed the positioning of chemically similar non-catalytic residues (Figure 4). Thus, our ensemble data provide evidence against the “Dynamic Entatic State Model”, at least for KSI. Other enzymes, lacking the requirement to catalyze chemical transformations at multiple positions and with greater catalytic challenges than KSI’s 10^11^-fold rate enhancement may utilize more precise positioning, a possibility to be tested in future ensemble-function studies.

### Conformational ensembles and transition states

Our ensembles have revealed a broad range of conformational states, in line with KSI’s need to abstract protons at multiple positions (Figures 1, 5-7). A subset of these conformations lie nearer to each position of proton abstraction (Figure 5). Nevertheless, our conformational ensembles represent states of high and intermediate occupancy, but are unlikely to capture rare high-energy states such as near transition states and other states of close approach in which the distance between the general base catalytic oxygen and ligand carbons is shorter than the sum of van der Waals radii. The states that we do observe are consistent with a rather smooth local conformational landscape, so that the transition states for proton abstraction are likely to lie at the “edges” of our ensemble and to retain a range of oxyanion hole and general base conformers, in line with the expectation that enzyme catalysis occurs though an ensemble of transition states rather than a single transition state (8). In addition, while no transition state analog (TSA) is a perfect mimic of the transition state, the affinities of oxyanion compounds used as KSI TSAs mirror catalytic effects of oxyanion hole mutations, suggesting that these TSA interactions faithfully represent the oxyanion interactions present in the actual transition states (89).

### The origins of KSI’s conformational heterogeneity

Given the importance of positioning and mobility, we would like to know the factors responsible for limiting and allowing conformational heterogeneity. This knowledge will lead to better understanding of how enzymes achieve catalysis and specificity and will help guide the design of new enzymes. Conformational heterogeneity data also allow computational models and conclusions to be evaluated. For example, recent conclusions that KSI efficiently discriminates between GS and TSA geometries rely on traditional single conformational cryo models and short molecular dynamics simulations, and require reevaluation in light of the KSI’s ensemble properties (96).

The general need for ensemble data is highlighted by the misleading view of positioning obtained by only considering individual structures. Different conclusions can be drawn depending on the structural snapshot considered (e.g., see Figure 8 and Figures S26). These structural snapshots are not incorrect; rather each cryo X-ray structure represents a portion of a more complex reality: there is an ensemble of states, determined by an energy landscape, which is, in turn, determined by the sum of energetic contributions from all interactions across all the available states and sub-states (8, 128–131). For example, additional Y16 positions are unleashed when a constraining hydrogen bond from Y57 is removed, revealing that the hydrogen bond was strong enough to limit the counterforce of conformational entropy (Figure 9). Without the constraints of the Y57-Y16 hydrogen bond, Y16 is now more likely to interact with M31. Nevertheless, the ensemble of Y16 states is broad and this new interaction occurs only in a subset of the observed conformational states, indicating that the Y16-M31 van der Waals interactions are not sufficiently strong to overcome the combination of Y16’s conformational entropy and other interactions that limit the ability of M31 to move to interact optimally with Y16 (Figure 9).

KSI’s general base, D40, appears to experience a “Goldilocks effect”. The catalytic oxygen of D40 explores a range of positions because the hydrogen bond of W120 to the non-catalytic oxygen provides a pivot point (Figure 6B, D). Remarkably, the W120F KSI variant is more active than WT KSI, and the W120F mutation increases the KSI activity to the KSI_homolog_’s level (Figure 6H). Mutations to other residues that more severely disrupt D40 positioning have, as expected, large deleterious effects (87, 88). Thus, our ensemble-function analysis suggests that there is an optimal amount of motion, at least for a residue that is required to carry out chemistry at more than one position (Figure 6I). It remains to be determined whether the noncovalent forces within a protein can over-dampen motions and slow individual reaction steps.

### Ensembles in enzyme design

What information and knowledge do we need to design enzymes that rival the catalysis and specificity of naturally-occurring enzymes? Unfortunately, we do not know precisely what is needed (132). In particular, we don’t know *how much* we need to restrict flexibility, or what motions we need to maintain or promote. Our KSI ensembles provide a rich dataset to investigate the origins of conformational heterogeneity. Future datasets and analyses will lead to detailed models for what allows and restricts conformational heterogeneity throughout an enzyme —e.g., what allows *vs.* limits the rotation of different side chains, what allows or limits the mobility of these surrounding groups, etc. Predictions from these models can be tested by subsequent RT X-ray crystallography of mutants. Further, ensemble-function analysis, as carried out herein, will link ensemble properties to their catalytic consequences, thereby defining necessary and optimal positioning and motion for particular catalytic tasks.

Ensemble data for natural enzymes, like that obtained herein, will provide specifications for use in the design of new enzymes, and ensemble-function studies of designed enzymes will be needed to assess successes and failures in enzyme design and suggest how to improve these enzymes using design principles (e.g., (133)).

### Ensembles in testing and improving force fields and computational approaches

A second hindrance to design is our limited understanding of the capabilities and limitations of the force fields and methods that underlie computational approaches (47, 49). Current computational approaches typically reproduce previously measured thermodynamic and kinetic constants. However, the ability of computational methods to predict unknown quantities that can be experimentally tested (“blind trials”) have provided more robust assessments (134). In addition, given the large number of parameters in a molecular force field, many different combinations of values can reproduce a small number of experimental measurements (47, 49). Conformational ensembles across all atoms of a protein provide a rich experimental source to address this limitation. Ultimately, a synergy between computational predictions and ensemble-based experiments may improve force fields and augment confidence in conclusions from computational results.

## Materials and Methods

### KSI expression and purification

The ketosteroid isomerase enzymes from *Pseudomonas putida* (pKSI, referred to herein as KSI, UniProt P07445) and *Comamonas testosteroni* (tKSI, referred to herein as KSI_homolog_, UniProt P00947) were expressed and purified as previously described with minor modifications (82, 135). KSI W120F and KSI_homolog_ F120W (KSI numbering) variants were obtained using standard mutagenesis protocols and the presence of the desired mutations was confirmed via DNA sequencing (see Table S60). Briefly, BL21 cells transformed with plasmid carrying the desired KSI construct were grown at 37 °C to OD 0.5-0.6 in LB media (EMD Millipore Corp, Billerica, MA, USA) containing 50 μg/mL carbenicillin (Goldbio, St Lousi, MO, USA), and protein expression was induced with 1 mM isopropyl-β-D-1-thiogalactopyranoside (Goldbio, St Lousi, MO, USA). After induction, cultures were grown for 10-12 h at 37 °C. Cells were harvested by centrifugation at 5000 g for 30 min at 4 °C and lysed using sonication. Lysed cells were centrifuged at 48000 g for 30 min at 4 °C. Enzymes were purified from the soluble fraction, first using an affinity column (deoxycholate resin) followed by a size exclusion chromatography column (SEC) Superdex 200. Prior to the purification of each enzyme, the affinity column, FPLC loops, and SEC column were washed with 40 mM potassium phosphate (JT Baker, Omaha, NE, USA), 6 M guanidine (JT Baker, Omaha, NE, USA), pH 7.2 buffer, and then equilibrated with 40 mM potassium phosphate, 1 mM sodium EDTA, 2 mM DTT (Goldbio, St Lousi, MO, USA), pH 7.2 buffer.

### KSI solution kinetics

KSI Michaelis–Menten parameters were obtained by monitoring the 5(10)-estrene-3,17-dione ((5(10)-EST), Steraloids, Newport, RI, USA) reaction at 248 nm (extinction coefficient 14,800 M^−1^ cm^−1^) in a PerkinElmer Lambda 25 spectrophotometer. Reactions were measured at 25 °C in 4 mM sodium phosphate, pH 7.2 buffer with 2% DMSO (JT Baker, Omaha, NE, USA) added for substrate solubility. Low buffer concentrations were used to minimize the background reaction rate. Values of *k*_*cat*_ and *K*_M_ were determined by fitting the initial rates as a function of substrate concentration to the Michaelis–Menten equation. Typically, seven to eight substrate concentrations, varying from 2 to 600 μM, were used for each mutant. The *k*_*cat*_ and *K*_M_ values were averaged from two independent measurements using different enzyme concentrations varied over 2–3 fold. Averaged values and errors representing the standard deviations are given in Table S58.

### KSI ^1^H solution Nuclear Magnetic Resonance

The ^1^H NMR spectrum of KSI D40N bound to equilenin was acquired at the Stanford Magnetic Resonance Laboratory using an 800 MHz Varian ^UNITY^INOVA spectrometer running VNMRJ 3.1A and equipped with a Varian 5 mm triple resonance, pulsed field gradient ^1^H[^13^C,^15^N] cold probe, as previously described (71). The sample contained 1.0 mM KSI and 2.0 mM equilenin (Steraloids, Newport, RI, USA) in 40 mM potassium phosphate (pH 7.2), 1 mM sodium·EDTA, 2 mM DTT, and 10% DMSO-*d*_*6*_ (v/v) (Cambridge Isotope Laboratories, Tewksbury, MA, USA). DMSO-*d*_*6*_ served as the deuterium lock solvent and prevented freezing at low temperatures. The spectrum was obtained in a 5 mm Shigemi symmetrical microtube at –3.5 °C, following temperature calibration with a 100% methanol standard. The 1331 binomial pulse sequence was used to suppress the water signal with a spectral width of 35 ppm (carrier frequency set on the water resonance) and an excitation maximum between 14-18 ppm (136). The data was processed using 10 Hz line broadening and baseline correction applied over the peaks of interest. Chemical shifts were referenced internally to the water resonance.

### Protein crystallization and X-ray data collection

All enzymes were crystallized as previously described (100). Briefly, enzyme were crystallized by mixing 1 μL of enzyme at 1 mM and 1 μL of crystallization solution (17-23% PEG 3350 (Hampton Research, Aliso Viejo, CA, USA) and 0.2 M MgCl_2_ (JT Baker, Omaha, NE, USA)) in a vapor diffusion hanging drop setup at room temperature. For crystallization of KSI bound to the transition state analog (equilenin) or the ground-state analog (4-androstenedione (Steraloids, Newport, RI, USA)), equilenin or 4-androstenedione were first dissolved in methanol (JT Baker, Omaha, NE, USA) at 20 mM and 40 mM concentration, respectively. Each ligand was then mixed with enzyme to achieve final concentrations of 1 mM enzyme and 2 mM equilenin or 4 mM 4-androstenedione (10% methanol in the final enzyme-ligand solution). As a ground-state analog, 4-androstenedione binds more weakly than the transition-state analog equilenin, and thus higher concentration was used to achieve higher occupancy. Crystals typically appeared after 24-72 h. Prior to data collection, crystals with minimum dimensions 0.2 x 0.2 x 0.2 mm were transferred from the crystallization solution to paratone N oil (Hampton Research, Aliso Viejo, CA, USA) where excess crystallization solution was stripped and crystals where then either frozen in liquid nitrogen for 100 K data collection and then mounted on the goniometer or directly mounted on the goniometer for 250 K or 280 K data collection. Data collection temperature was controlled using a N_2_ cooler/heater. Single-crystal diffraction data were collected at SSRL, beamline BL9-2, using wavelengths of either 0.787 Å or 0.886 Å. See Supplementary file Table S3 for diffraction data statistics.

### Crystallographic data processing and model building

Data processing was carried out with in-house scripts: http://smb.slac.stanford.edu/facilities/software/xds/#autoxds_script. Briefly, data reduction was done using the XDS package (137), scaling and merging was done using *Aimless* (138, 139) and structure factor amplitudes were obtained using *Truncate* (138, 140). Initial phases were obtained via molecular replacement using *PHASER* (141) and the PDB entry 3VSY as a search model. Model building was carried out with the program *ARP/wARP* (142) and manually in *Coot* (143). Traditional, single conformation models, in which major alternative side chain and backbone conformations were modeled, were refined manually after visual inspection with *Coot* and using *phenix.refine* (144). Torsion-angle simulated annealing (as implemented in *phenix.refine*) was used during the initial stages of refinement. Riding hydrogens were added in the late stages of refinement and their scattering contribution was accounted for in the refinement. Ligand restraints were generated using the *GRADE* server (http://grade.globalphasing.org/cgi-bin/grade/server.cgi). Model quality was assessed using *Molprobity* (145) as implemented in *phenix.refine* and via the PDB Validation server (https://validate-rcsb-2.wwpdb.org/). See Supplementary file Table S3 for refinement statistics.

Multi-conformer models were obtained from the 250 K diffraction datasets, using previously described methods (53, 54, 56, 64, 146). As a large body of work identified the 180–220 K temperature range as an inflection point above which various protein motions are activated, providing strong evidence that at and above 250 K both harmonic and anharmonic protein motions are enabled (56, 57, 59, 147, 148), and as the 250 K diffraction data was of higher resolution than our 280 K data (Table S3), we used the higher-resolution 250 K to obtain multi-conformer models of Apo, GSA-bound, and TSA-bound KSI. Briefly, the program *qFit* was used to obtain multi-conformation models (53, 54) using as input the traditional single-conformation models obtained above after removing the riding hydrogen atoms. Subsequent to the automated multi-conformer model building, ill-defined water molecules were deleted and alternative protein side and main chain conformations and orientations were edited manually after visual inspection in *Coot* and based on the fit to the electron density (149). Models were subsequently refined with *phenix.refine* (144). Riding hydrogen atoms were added in the late stages of refinement and their scattering contribution was accounted for in the refinement. Final multi-conformer model quality was checked by *MolProbity* (145) and via the PDB Validation server (https://validate-rcsb-2.wwpdb.org/) and deposited on the PDB (Table S3). See Supplementary file Table S3 for refinement statistics.

### Crystallographic order parameters calculation

Crystallographic order parameters, S^2^, were obtained from the 250 K multi-conformer models as previously described (64). These order parameters include both harmonic and anharmonic contributions as captured by the crystallographic atomic displacement parameters (B-factors) and by the occupancies of alternative rotameric states, respectively; these values correlate well with solution NMR-derived S^2^ (64). The analysis was applied to the bond most closely associated with the first side-chain dihedral angle (χ_1_), using Cβ—H for all amino acids other than Gly and Cα—H for Gly. Because S^2^ varies from 0 to 1 as a measure of order, we used 1-S^2^ as a measure of disorder.

The KSI crystals obtained in this study contained two molecules in the asymmetric unit and the average of the two molecules was used for analysis. Because the total ground-state analog (4-Androstenedione) occupancy in the final refined model of the two crystallographically-independent KSI molecules was 1.4 instead of 2.0 (corresponding to the maximum occupancy of 1.0 for each of the KSI molecules), the (1-S^2^) values for each residue were corrected using the equation: 

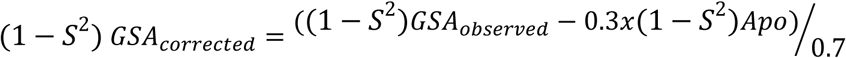

### Ensemble building

To obtain KSI pseudo-ensembles, all KSI cryo crystal structures were downloaded from the PDB (62) and parsed into individual KSI monomers (Table S1). While KSI is a dimer, we focused the analysis on the individual KSI molecule as: i) all individual KSI molecules were highly similar, with RMSDs below 0.5 Å after excluding two flexible loops (see Results); ii) KSI is known to also crystallize with one molecule in the crystallographic asymmetric unit (see Table S1), indicating that the two molecules of the dimer are identical; and iii) each monomer has the full catalytic machinery required for catalysis. All KSI molecules were aligned using PyMOL and standard commands (The PyMOL Molecular Graphics System, Version 2.0 Schrödinger, LLC.) on the protein backbone (N, Cα, C, O atoms) of residues 5-125 on either the highest resolution crystal structure of KSI bound to the transitions state analog equilenin (PDB 1OH0, (60)) or as otherwise indicated. Residues 1-4 (N-terminal) and 126-131(C-terminal) were excluded from the analyses because these residues appeared highly flexible, as is common for N/C-terminal residues, and were also not modeled in some of the KSI structures. No alignment gaps were allowed during the alignment, and allowing for gaps (default PyMOL alignment procedure) did not appreciably change the results (Figure S9). The same approach was used to obtain KSI_homolog_ pseudo-ensembles, with the backbone of residues 3-122 aligned on the highest resolution crystal structure of KSI_homolog_ bound to the ground state analog 4-androstene-3,17-dione (PDB 3NHX). Different sets of crystal structures were used to obtain different types of pseudo-ensembles (sub-ensembles) and the structures included in the different types of pseudo-ensembles are listed in Tables S1, 2, 20-22 and explicitly indicated in the legends of figures from the main text and supplemental information. The KSI RT-ensemble was obtained from the Apo, GSA-bound, and TSA-bound RT multi-conformer models using the same alignment procedure and the Apo state as alignment template.

### Calculating mean deviations

From the ensemble of aligned crystal structures or multi-conformer models, a list of *xyz* coordinates was defined for each atom using PyMOL and standard code. For *n* copies of each particular atom in the ensemble, a coordinate list of size 3*n* was obtained for this atom. The mean deviation (MDev) was then calculated by computing the spread of these points about the center point *j*: 

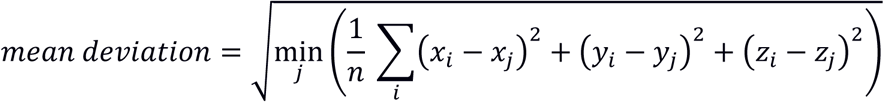

Here *i* is an index over each of the *n* atoms in the ensemble and the average distance of all points to a center point from atom *j* was calculated. The center point *j* was chosen to give the minimum average distance to all other points in the given ensemble of atoms and was determined by calculating the average distance between each atom in a given ensemble and all other atoms in the same ensemble of atoms.

### Bootstrap analysis

A bootstrap analysis was used to estimate the errors associated with the sum of Cα MDev values of the Apo and the transition-state bound KSI pseudo-ensembles (ΣMDev Apo and TSA-bound, Figure 3E from the main text). Briefly, from the number of distances, *n* (e.g. for the full pseudo-ensemble of 94 structures (*q* = 94), *n* = *q* – 1), used to calculate the Cα MDev for a given KSI residue *x* (where *x* denotes residues 5 to 125), a random number of distances, *m*_*1*_, was randomly selected and excluded from *n*. A second small number of distances, *m*_*2*_, equal to *m*_*1*_, was randomly selected from *n* to replace the excluded *m*_*1*_ and then the sum of the MDevs for residues 5-125 was calculated (ΣMDev). The procedure was repeated 300 times and the standard deviations associated with an increasing number of bootstrapped ΣMDev (*n* = 2, 5, 10, 20, 30, 40, 50, 100, 200 and 300 cycles, Figure S8, Table S27) was used to estimate the standard deviation of the ΣMDev in Figure 3E from the main text.

## Acknowledgments

This work was funded by a National Science Foundation (NSF) Grant (MCB-1714723) to DH. FY was supported in part by a long-term Human Frontiers Science Program postdoctoral fellowship and in part by the NSF grant MCB-1714723. MMP was supported in part by an NSF Graduate Research Fellowship and in part by a Lieberman Fellowship from Stanford University. ASP was supported in part by the National Science Foundation Graduate Research Fellowship and in part by the Stanford ChEM-H Chemistry/Biology Interface Predoctoral Training Program and the National Institute of General Medical Sciences of the NIH under Award Number T32GM120007. We thank Dr. Steve Bonilla for help with the bootstrap analysis, Stanford Synchrotron Radiation Lighthouse (SSRL) and Lisa Dunn for beam time allocation and access, Dr. Corey W. Liu, Stanford Magnetic Resonance Laboratory, and Dr. Mark Kelly, UCSF Nuclear Magnetic Resonance Laboratory, for assistance with NMR spectroscopy, and members of the Herschlag lab for helpful suggestions and feedback on the manuscript. The SMRL [MMP1] 800 MHz NMR was supported in part by NIH Shared Instrumentation Grant 1 S10 RR025612-01A1.

## Footnotes

1 A trivial example of decreased catalysis via reduced positioning is when a catalytic group is restricted to positions incompatible with catalysis. In such a case, increasing its mobility can increase catalysis, even if only one reaction step is involved. The KSI example in the text represents a distinct case that is common in which mobility is needed to carry out multiple reaction steps.

