## Supplementary file for "Assessment of enzyme active site positioning and tests of catalytic mechanisms through X-ray-derived conformational ensembles"

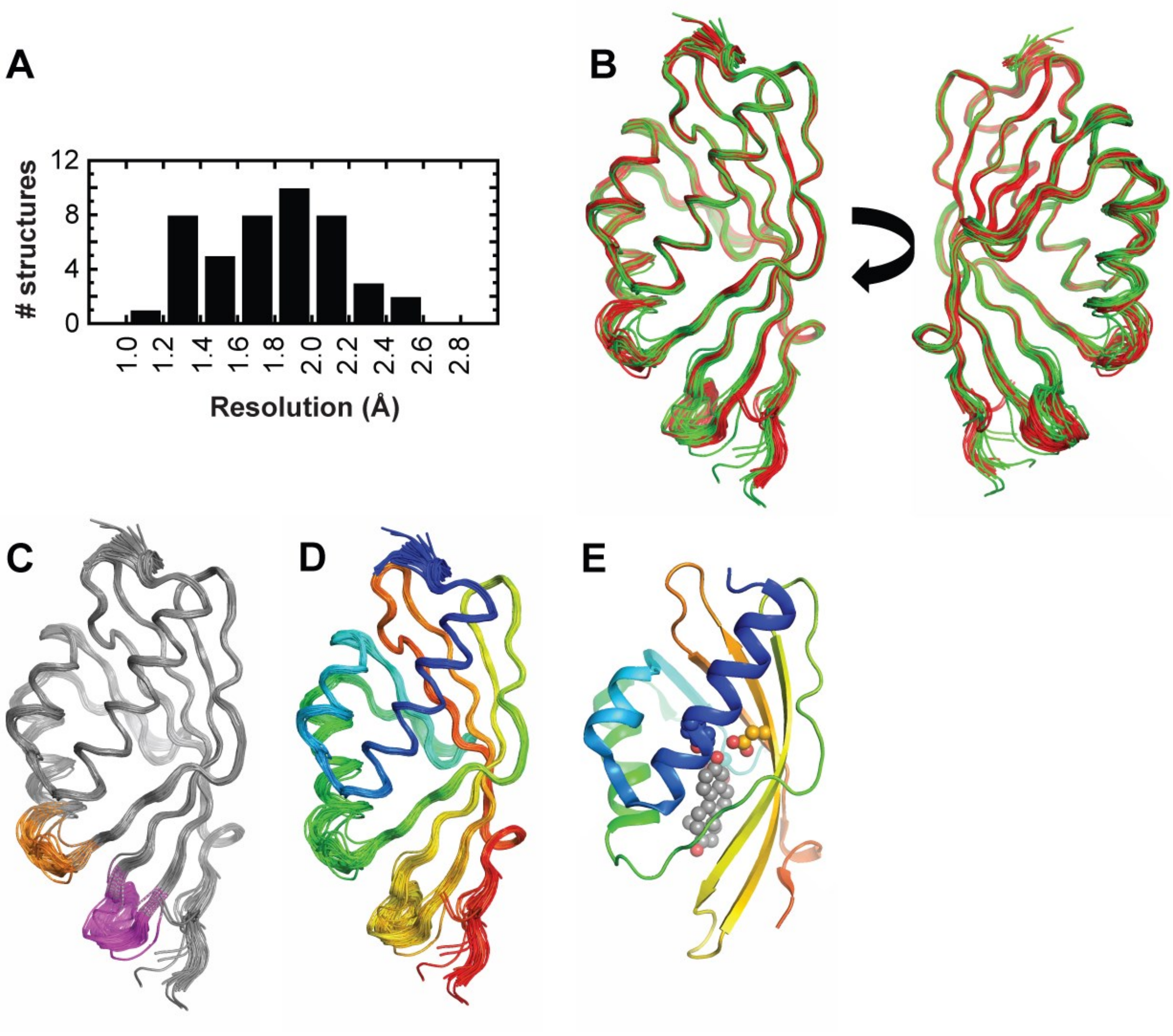

**Figure S1. The KSI pseudo-ensemble**. (A) Resolution of the KSI cryo crystal structures used to obtain pseudo-ensembles (Table S1). (B-D) Individual KSI molecules are colored, as follows: (B) according to the catalytic state: Apo (green), GSA-bound (blue), and TSA-bound (red); (C) in grey, with the exception of the 62-65 loop (in orange) and 91-96 loop (in magenta); and (D) from N– (blue) to C–terminal (red). (E) KSI bound to a TSA (PDB 1OH0) using the same color code as (D), with Y16 (blue), D103 (orange), D40 (teal), colored according to their associated secondary structure element, and bound TSA (grey) shown in spheres.

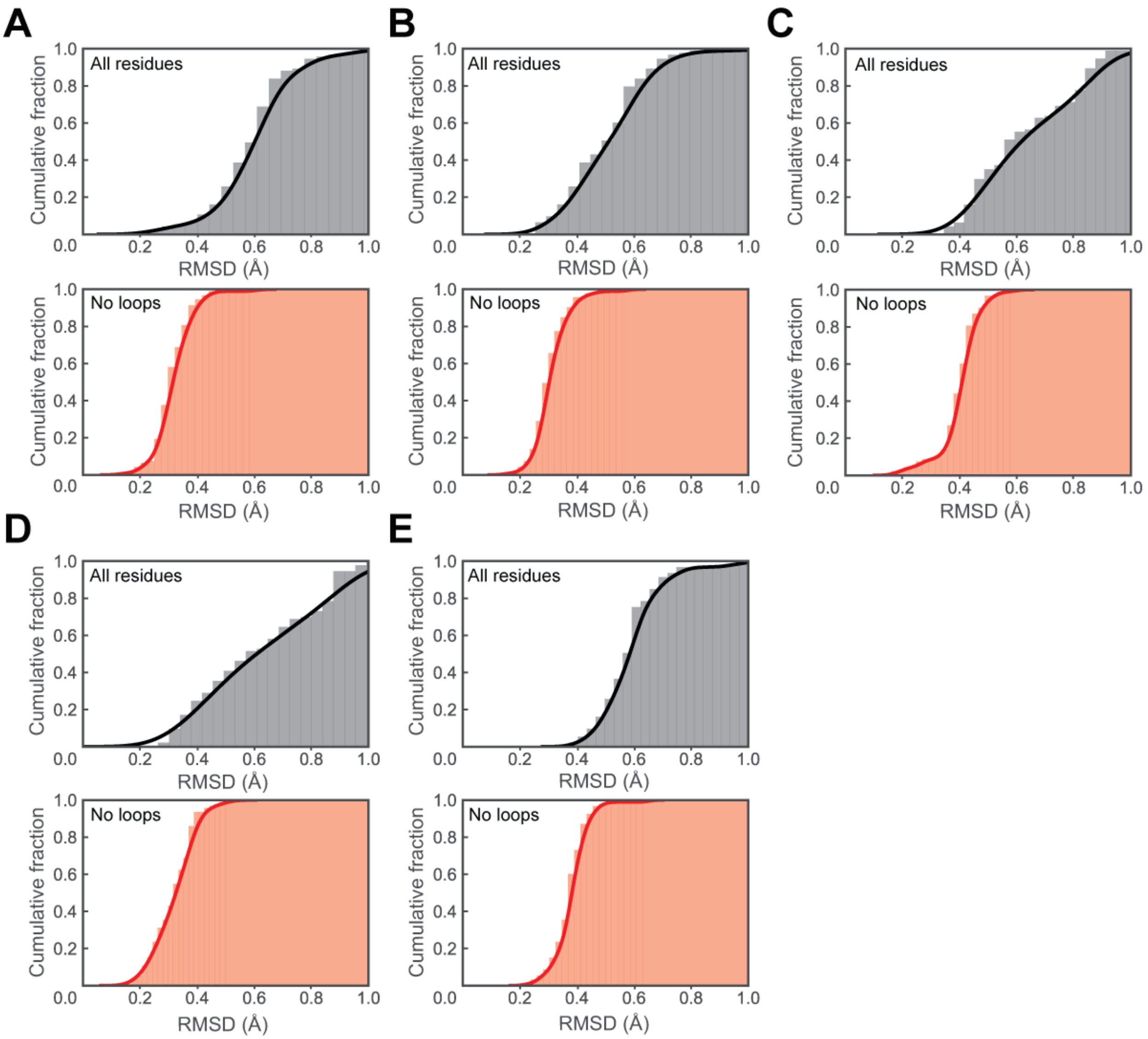

**Figure S2. RMSDs for all available KSI cryo structures**. Structures were aligned on the backbone of residues 5–125. Backbone RMSDs for WT Apo (PDB 3VSY, A and B), WT GSA-bound (PDB 5KP4, C), and WT TSA-bound (PDB 1OH0, D and E) relative to each of the remaining crystallographically-independent KSI molecules from the PDB (N = 95). A and B represent each of the two crystallographically-independent KSI molecules of WT Apo (PDB 3VSY) and similarly D and E represent each of the two crystallographically-independent KSI molecules of WT TSA-bound (PDB 1OH0), whereas for PDB 5KP4 (C), there is only the crystallographically-independent KSI molecule with bound GSA, molecule B. The larger RMSD values and variability in the "All residues" panels reflect the greater structural variability of the 62–65 and 91–96 loops.

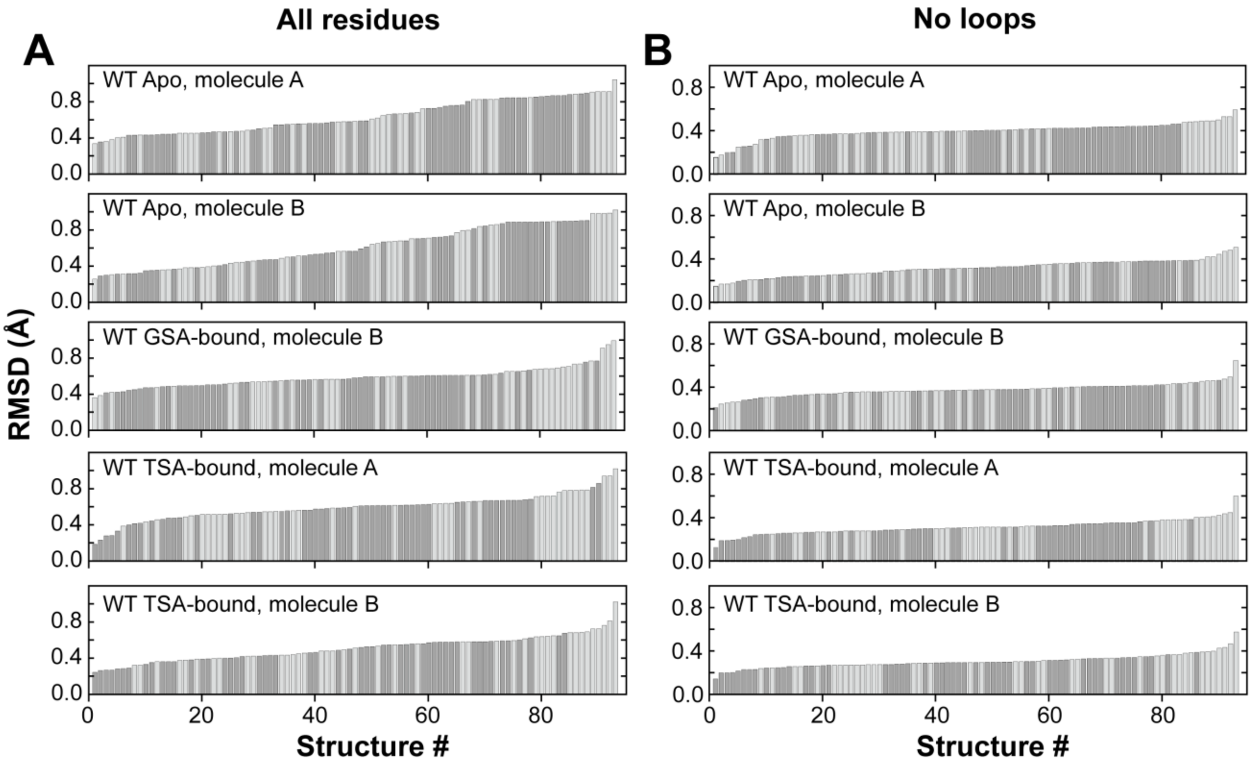

**Figure S3.** Ranked RMSDs for all available KSI cryo structures. Backbone RMSDs from Figure supplement 2 have been ranked according to increasing RMSD values for (A) the entire sequence and (B) excluding loops 62–65 and 91–96. RMSDs with Apo and Ligand-bound KSI molecules are colored in light and dark grey, respectively.

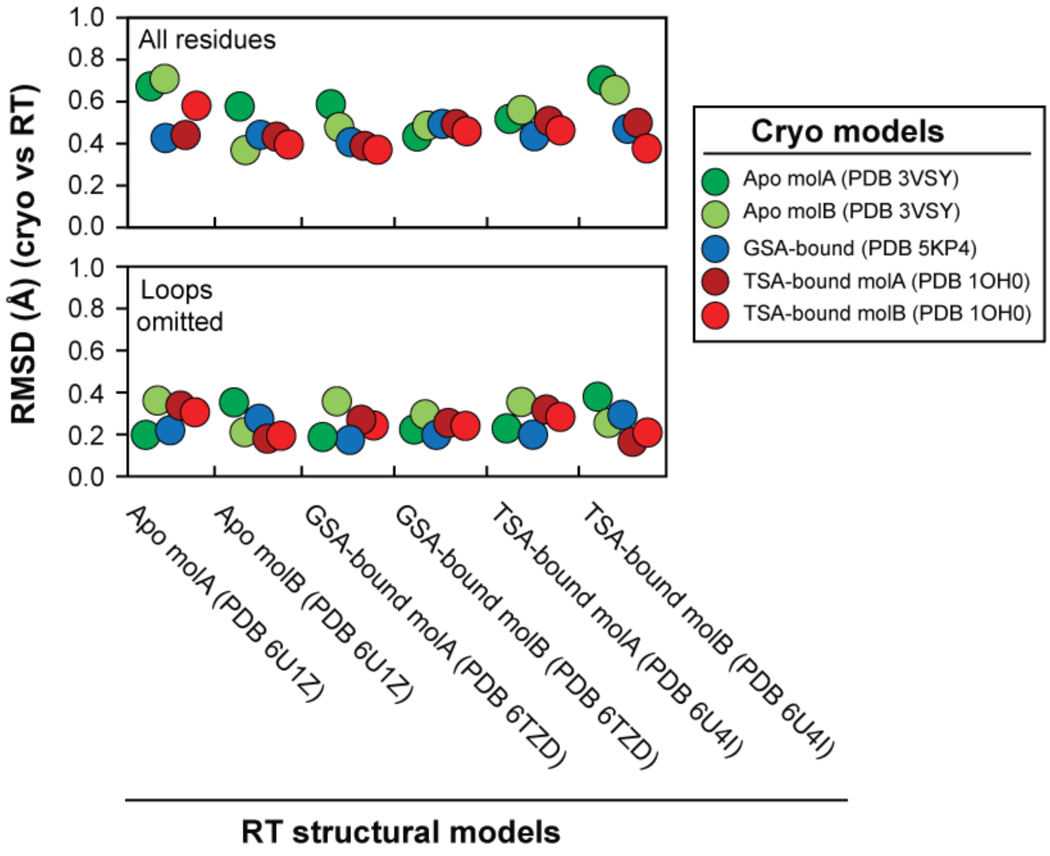

**Figure S4.** Backbone RMSDs of Apo, GSA-bound and TSA-bound KSI single-conformation structures obtained at RT (280 K, this study; x-axis) *vs.* the highest-resolution cryo (100 K, from the PDB; y-axis). There are two crystallographically-independent molecules for each enzymes state, for both RT and cryo structures, except for the GSA-bound cryo structure, for which there was only one GSA-bound molecule. Each circle represents the RMSD between each of the two independent KSI molecules from the RT structures (x-axis) and each of the independent molecules from the cryo structures (color coded in legend). Figure 2 from the main text represents the average RMSD between each of the two independent monomers from the KSI dimer from the RT structures and each of the two independent monomers from the KSI dimer the cryo structures.

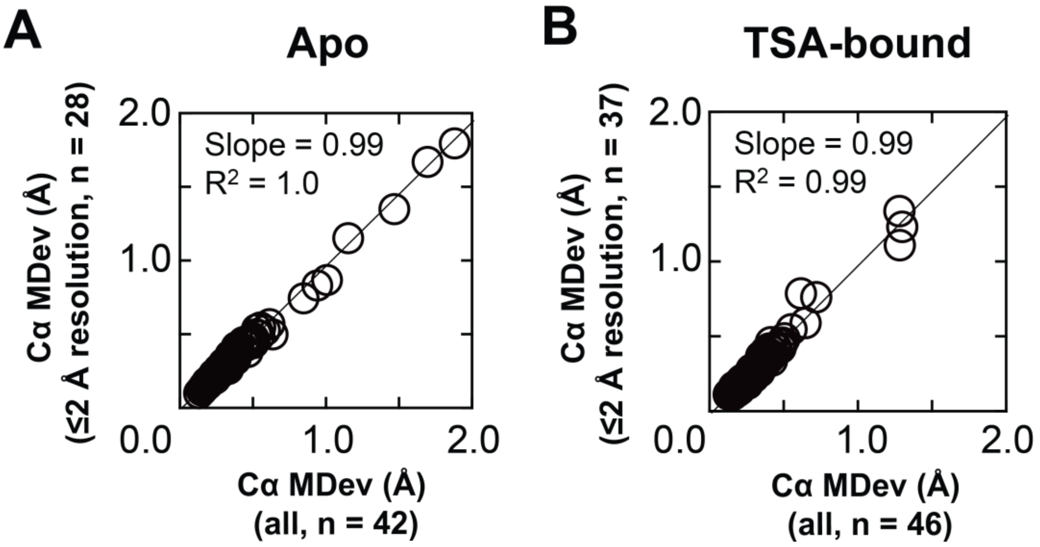

**Figure S5. Exclusion of lower resolution structures does not alter the KSI pseudo-ensemble properties.** Correlation between Cα MDevs for the full pseudo-ensemble *vs.* a pseudo-ensemble including only higher-resolution Apo structures for (A) Apo pseudo-ensembles (42 and 28 structures, respectively) and (B) TSA-bound pseudo-ensembles (46 and 37 structures, respectively). High-resolution structures are defined as structures with resolutions ≤2 Å. For a given atom in a structure, the MDev describes the average displacement of equivalent atoms within the ensemble of structures, with lower and higher values representing smaller and larger positional fluctuations, respectively, corresponding to less or more conformational heterogeneity (also see Materials and Methods for a more complete definition of MDev).

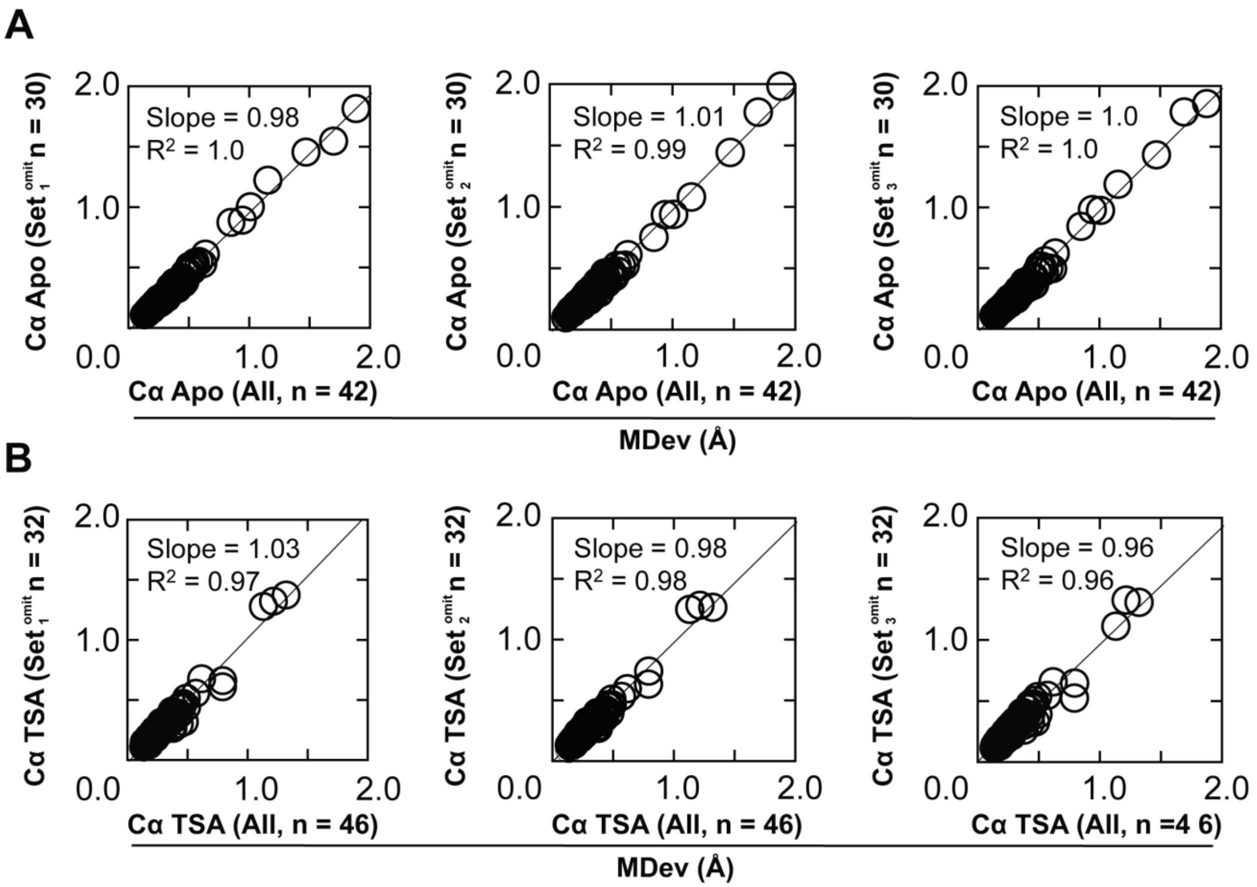

**Figure S6. Random omission of KSI molecules does not alter the pseudo-ensemble properties.** Comparison of Cα MDevs for (A) Apo and (B) TSA-bound from the full pseudo-ensembles (obtained from 42 (Apo) and 46 (TSA-bound) KSI molecules, respectively) and pseudo-ensembles from which 30% of the structures were randomly omitted (12 out of 42 and 14 out of 46 KSI molecules, respectively); random selection and exclusion of molecules was repeated three times to generate three independent sets (Set1-3); Table S8).

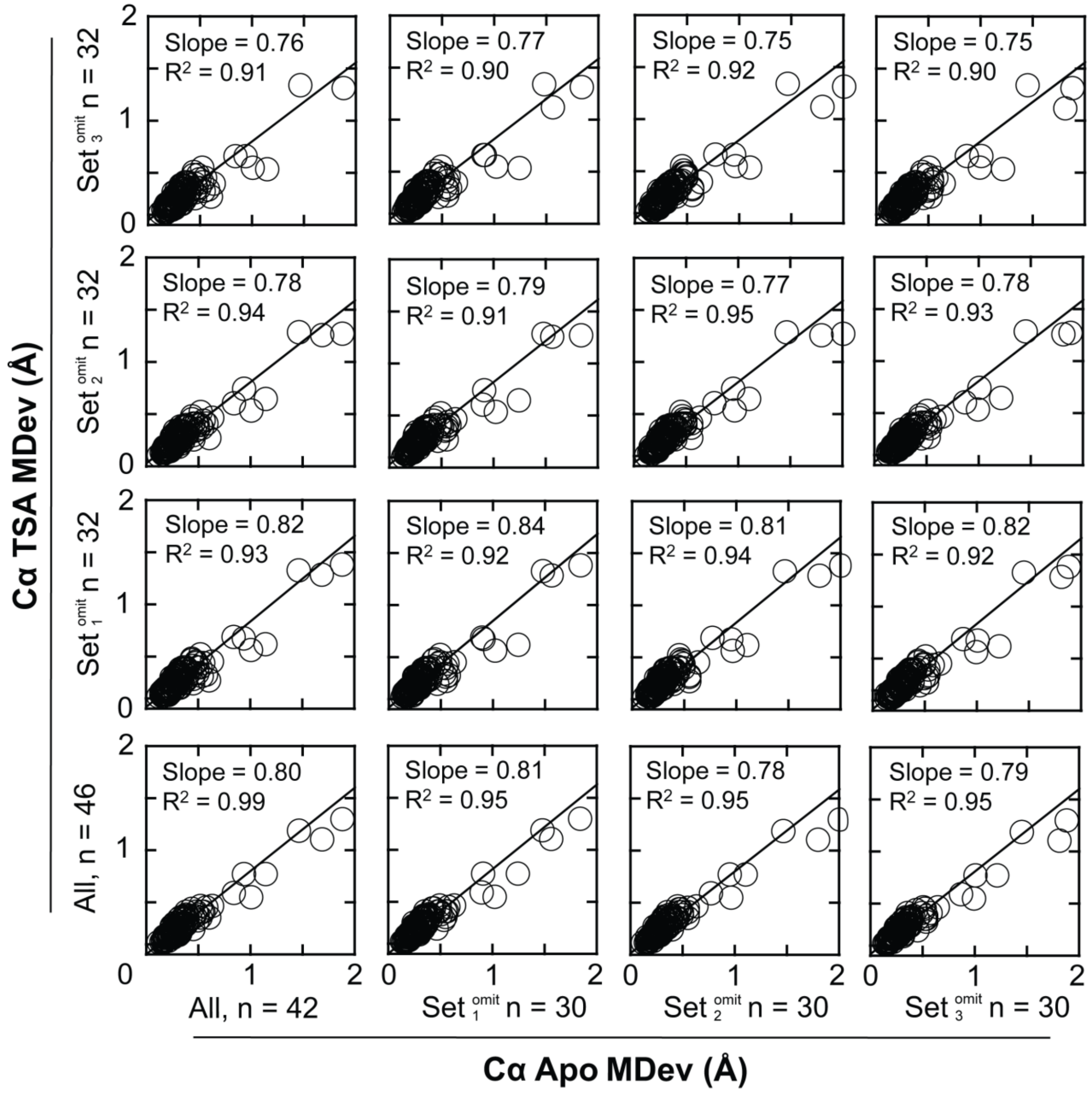

**Figure S7. Random omission of KSI molecules from pseudo-ensembles has no significant impact on the conformational heterogeneity dampening in Apo KSI upon TSA binding.** Comparison of correlation plots between Cα MDevs for Apo and TSA-bound pseudo-ensembles composed of all structures (42 and 46 KSI molecules, respectively) and pseudo-ensembles from which 30% of the structures have been randomly omitted (12 out of 42 and 14 out of 46 KSI molecules, respectively; data from Figure S5, Tables S8 and S9). The average slope is 0.79 ± 0.02 and average $R^2$ is 0.93 ± 0.02.

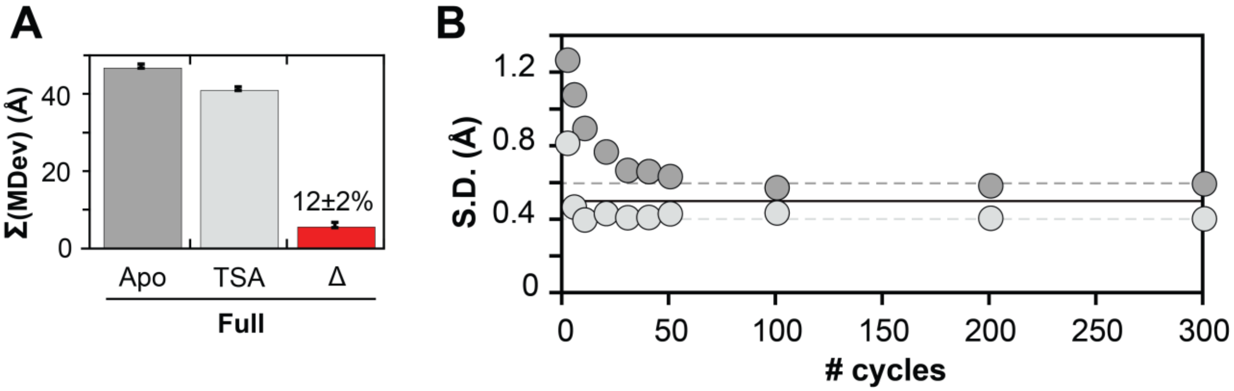

**Figure S8. Estimating the error in the sum of Cα MDevs from the Apo and TSA-bound pseudo-ensembles.** (A) Sum of Cα MDevs for Apo (dark grey bars), TSA-bound (light grey bars) and the sum of their difference (Δ, red bars) for the entire enzyme (reproduced from Figure 3E from main text). The error bars in (A) were estimated using a bootstrap analysis. Briefly, ΣMDev Apo and TSA-bound were obtained by summing the Cα MDevs for residues 5-125 in Apo and TSA-bound pseudo-ensembles, respectively. The Cα MDev for each residue is the average distance of all Cα atoms within the Cα ensemble for a given residues, to the center of the same Cα ensemble. For a number of distances $n$, (in either the Apo or the TSA-bound pseudo-ensemble), a random number of distances, $m_1$, was randomly selected and replaced by second number of distances, $m_2$, equal in number to $m_1$, and randomly selected from the same ensemble. A new Cα ΣMDev for residues 5–125 was then obtained and the procedure was repeated 300 times (see Materials and Methods for a more complete description). (B) The standard deviation (S.D.) over an increasing number of bootstrapped Cα ΣMDevs. The S.D. levels off as the number of bootstrapped Cα ΣMDevs increases, as expected (dark and light grey dashed lines for Apo and TSA-bound pseudo-ensembles, respectively). The solid black line indicates the average, 0.5 Å, which was used as a measure of the error in (A) and in the Figure 3E of the main text.

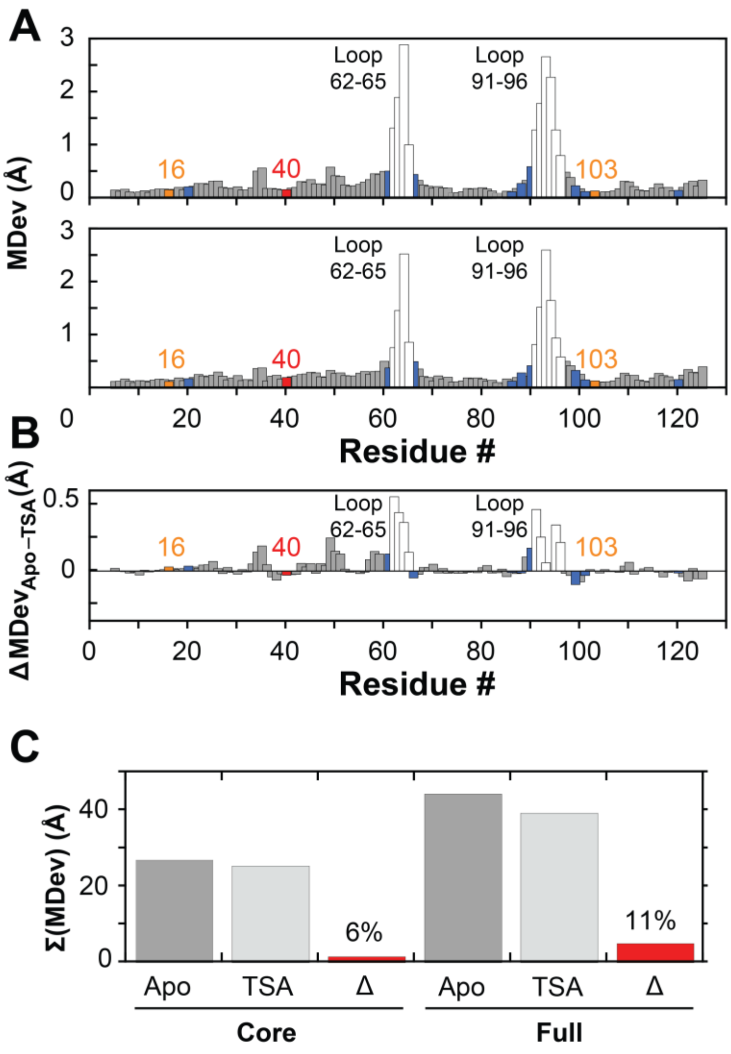

**Figure S9. The alignment procedure does not impact the change in conformational heterogeneity for Apo *vs.* TSA-bound KSI obtained from pseudo-ensembles.** Apo and TSA-bound KSI pseudo-ensembles were obtained using an alignment procedure different from the alignment procedure used in the main text (see Materials and Methods). (A) Cα MDevs for KSI Apo (top) and TSA–bound (bottom) states. (B) Difference Cα MDev values between the Apo and TSA-bound states ($MDevs_{Apo\text{-}TSA}$), such that positive values indicate lower MDevs for the TSA-bound state. The differences are similar throughout the enzyme, except for larger changes in the 62-65 and 91-96 loops (white bars). (C) Sum of Cα MDevs for Apo, TSA-bound and their difference. The color code is the same as in Figure 3 of the main text.

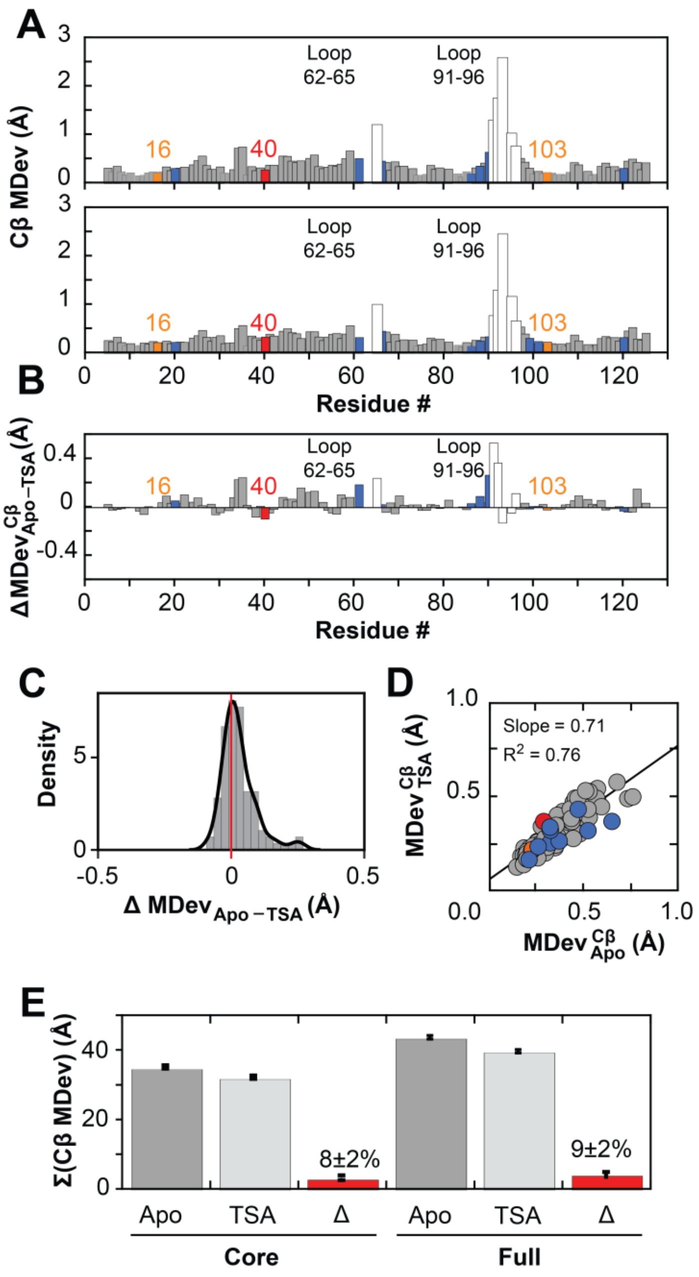

**Figure S10. Quantifying changes in conformational heterogeneity through the KSI catalytic cycle via pseudo-ensemble Cβ MDevs.** (A) Cβ MDevs for KSI Apo (top) and TSA–bound (bottom) states. (B) Difference Cβ MDev values between the Apo and TSA-bound states ($MDevs_{Apo\text{-}TSA}$), such that positive values indicate lower MDevs for the TSA-bound state. (C) Histogram of MDev differences from part (B; $MDevs_{Apo\text{-}TSA}$) for the enzyme core (i.e., loops excluded). (D) Correlation plot of Apo and TSA-bound Cβ MDevs (excluding loops 62–65 and 91–96). (E) Sum of Cα MDevs for Apo, TSA-bound and their difference; colors as in Figure 4 main text. The slightly lower dampening of conformational heterogeneity for the full enzyme estimated using Cβ MDevs *vs.* Cα MDevs (12±2% and 9±2% *vs.* 8±2% and 9±2%, respectively) arises at least in part because both 62–65 and 91–96 loops contain Gly residues (3 and 1 Gly residues, respectively) which do not have Cβ atoms. Color code as in Figure 3 of the main text.

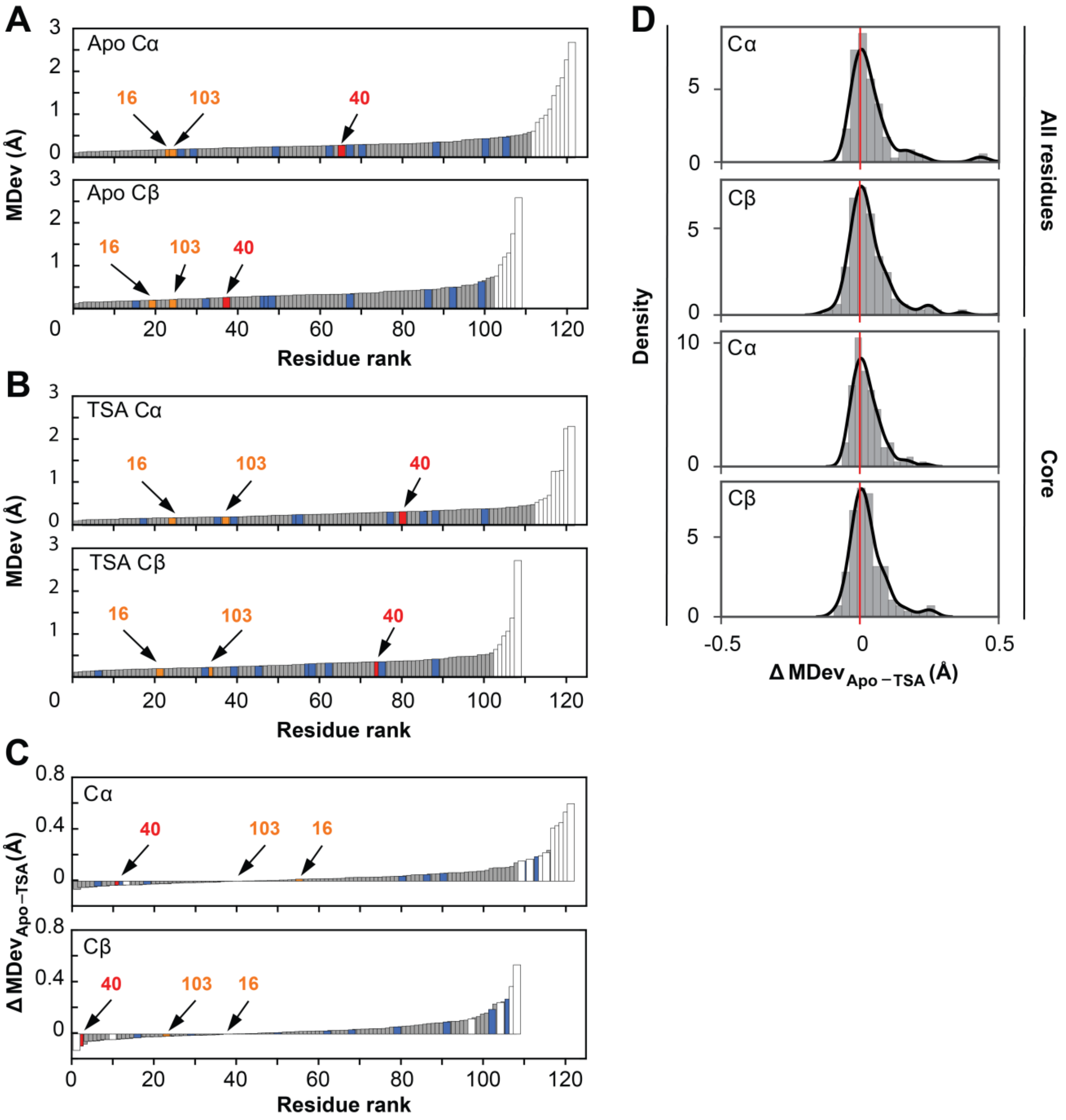

**Figure S11. Conformational heterogeneity through the KSI catalytic cycle obtained from pseudo-ensembles.** MDevs for KSI Apo (A) and TSA–bound state (B) Cα (top) and Cβ (bottom) in rank order. (C) Difference Cα (top) and Cβ (bottom) MDev values between the Apo and TSA-bound states ($MDevs_{Apo\text{-}TSA}$) ordered according to increasing $MDevs_{Apo\text{-}TSA}$ values. Positive values indicate lower MDevs for the TSA-bound state. The largest changes occur in the 62–65 and 91–96 loops (white bars). (D) Histogram of $\Delta MDev_{Apo\text{-}TSA\text{–}bound}$ values for all residues (two top panels) and for the enzyme core (two bottom panels). Color code as in Figure 4 of the main text.

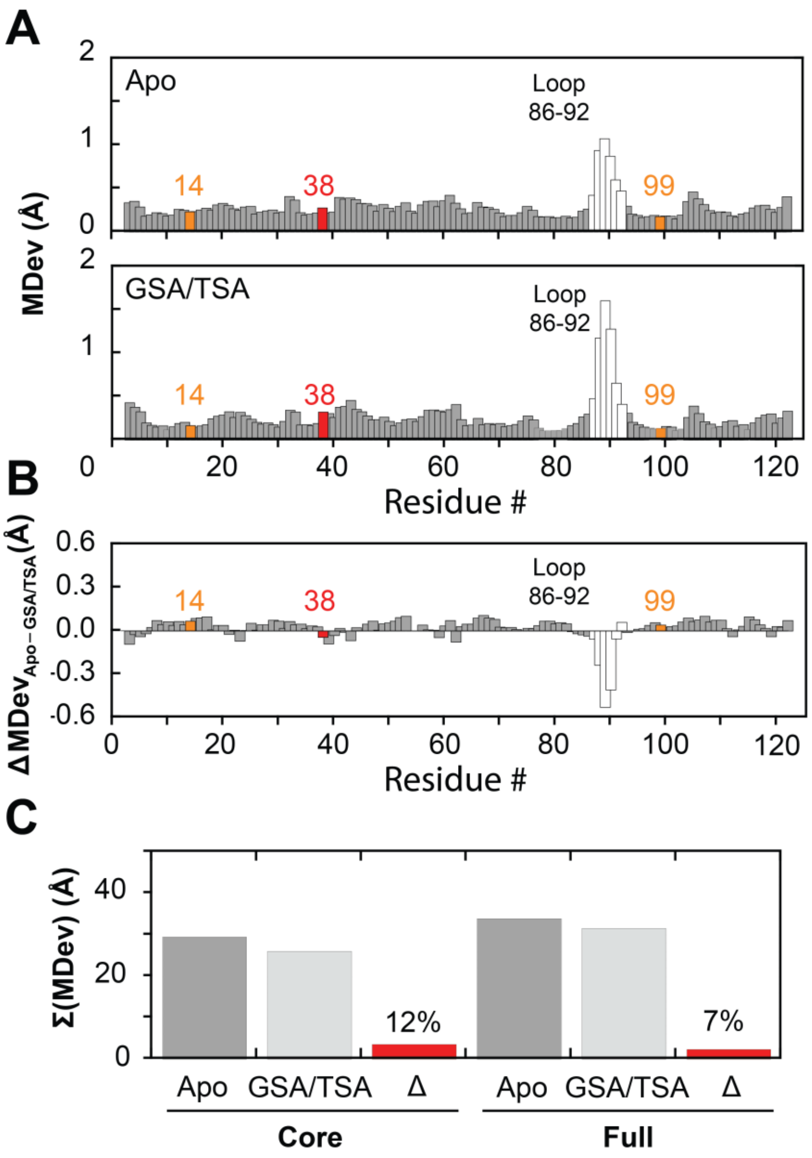

**Figure S12. Quantifying changes in conformational heterogeneity for a homologous KSI ($KSI_{homolog}$) catalytic cycle via pseudo-ensembles.** (A) Cα MDevs for KSI Apo (n = 24, top) and GSA/TSA–bound (n = 18, bottom) states. GSA-bound and TSA-bound structures were pooled due to the low number of GSA-bound and TSA-bound molecules (9 and 9, respectively) from PDB crystal structures (Table S21-23). The flexible 86–92 loop is presented as white bars; positions 14 and 99 (oxyanion hole, corresponding to positons 16 and 103, respectively in the main text) in orange and position 38 (general base; D40 in main text) in red. (B) Difference Cα MDev values between the Apo and GSA/TSA-bound states ($\Delta MDevs_{Apo\text{-}TSA}$), such that positive values indicate lower MDevs for the GSA/TSA-bound state. (C) Sum of Cα MDevs for Apo, GSA/TSA-bound and their difference (Δ); colors as in Figure 4 of the main text. From the above comparisons it appears that the 86–92 loop becomes more flexible upon GSA/TSA binding; nevertheless not that the ΔMDev for the loop represents a small difference between large values (note the different scale of the y-axis in A *vs.* B).

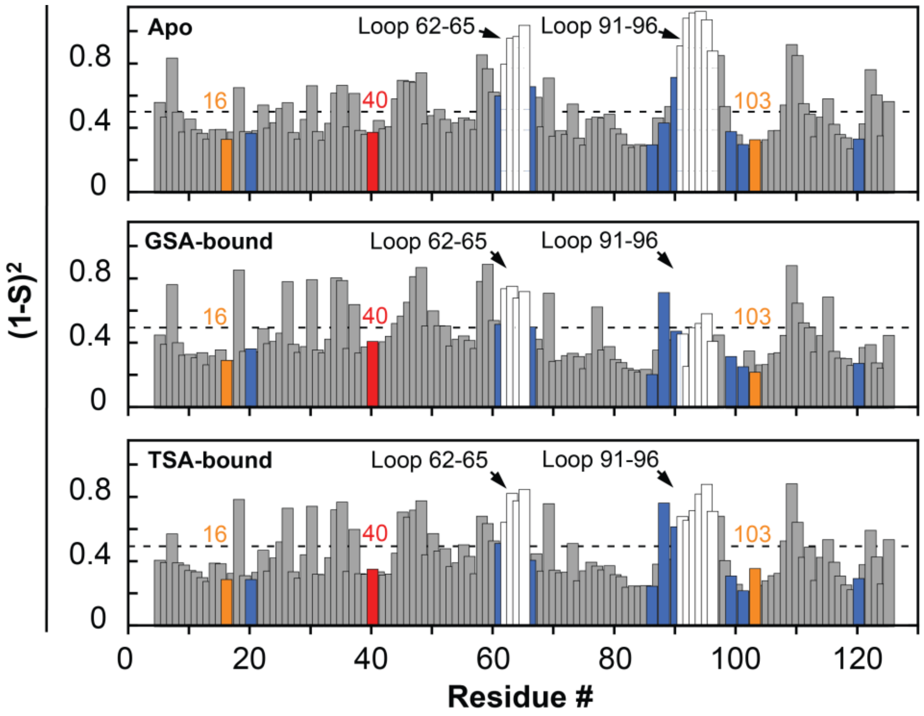

**Figure S13. Conformational heterogeneity in KSI catalytic states from RT X-ray data.** Disorder parameters, $(1\text{-}S^2)$, obtained from multi-conformer models for KSI Apo (top), GSA-bound (middle) and TSA-bound (bottom). $(1\text{-}S^2)$ for the GSA-bound state corrected for 70% GSA occupancy (see Figure S14 for uncorrected values). Dashed lines represent average $(1\text{-}S^2)$ values: 0.52, 0.45, and 0.46 for Apo, GSA- and TSA-bound, respectively. Y16 and D103 are in orange; D40 in red; binding residues in blue; and loop residues in white, as in Figure 3A from main text.

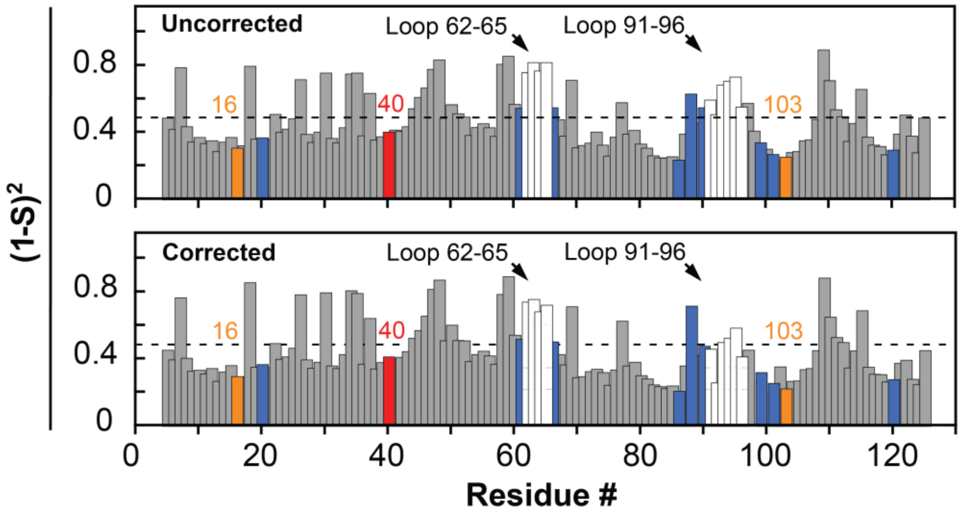

**Figure S14. Correcting for incomplete occupancy does not appreciably alter the conformational heterogeneity in the KSI GSA-bound multi-conformer model.** The observed (uncorrected, top) and GSA occupancy–corrected (70% occupancy, bottom) GSA-bound (1-$S^2$) obtained from the 250 K multi-conformer model are highly similar and give analogous results and conclusions (see Table S32). The bottom panel is repeated from the Figure S13 (middle). Color code as in Figure 3A from main text.

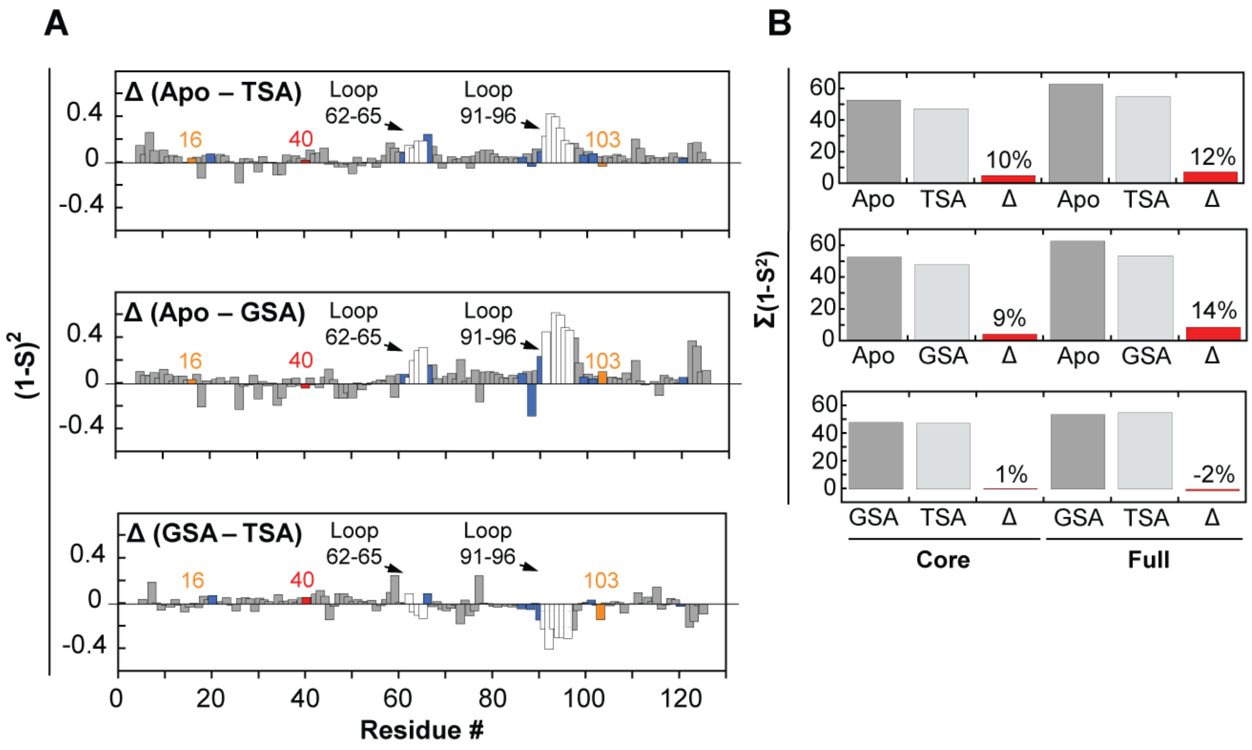

**Figure S15. Changes in KSI conformational heterogeneity during its catalytic cycle from RT X-ray data.** (A) Difference (1-$S^2$) between Apo and TSA-bound (top), Apo and GSA-bound (middle) and GSA and TSA-bound (bottom). Y16 and D103 in orange, D40 in red, and binding residues in blue. (B) Sum of (1-$S^2$) values for the different catalytic states (grey bars) and their difference (Δ, red bars). Each panel in B is the summed (1-$S^2$) value or difference (Δ) from the respective panel in A. Color code as in Figure 3A from main text.

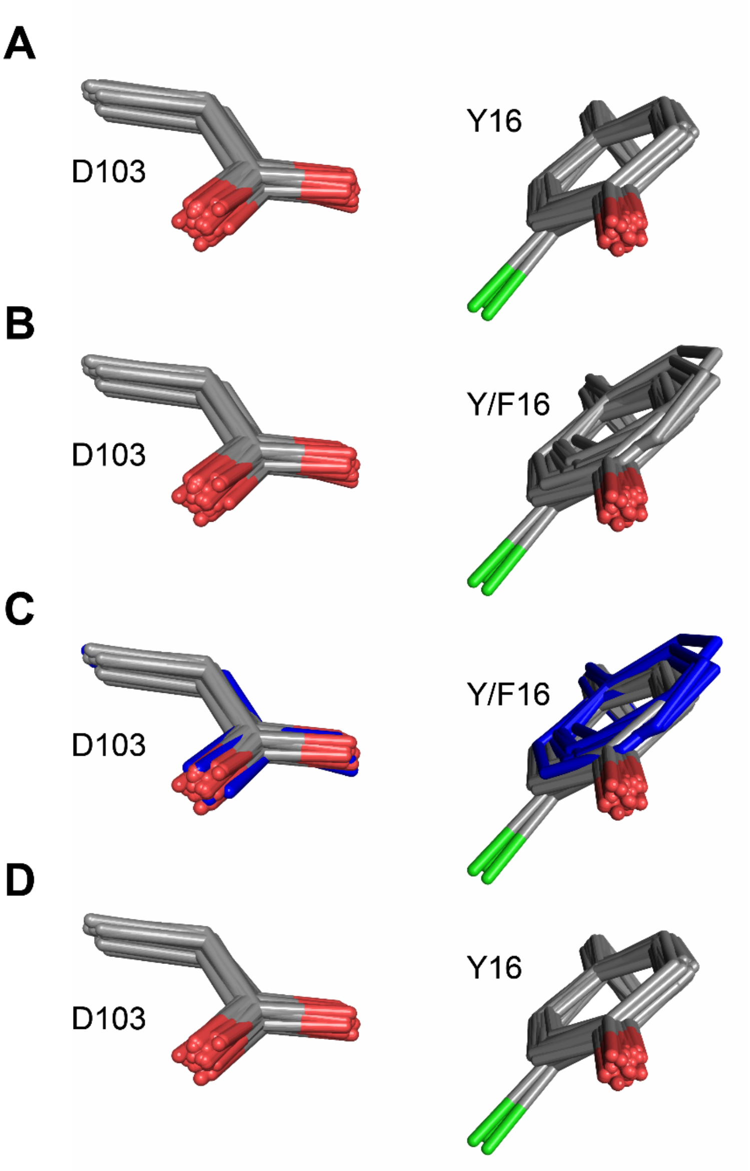

**Figure S16. Ablation of the Y16 hydrogen bonding group does not alter D103 positioning.** (**A**) The oxyanion hole D103 and Y16 reduced pseudo-ensemble, which does not include structures with i) mutations in the oxyanion hole that alter the chemical nature of the hydrogen bonding groups or ii) mutations in the Y16 hydrogen bond network (e.g. Y57F) (Table S2). Phenylalanine residues at position 16 are omitted in this panel. Chlorine atoms in chemically modified tyrosine residues are shown in green. (B) The reduced pseudo-ensemble with phenylalanine residues at position 16 included (Table S2). (C) The reduced pseudo-ensemble from B in which structures with phenylalanine residues at position 16 are colored in blue. The aspartate residues from structures with phenylalanine at position 16 (blue) are within the ensemble of aspartate residues from structures with tyrosine at position 16. (D) The reduced pseudo-ensemble from which all structures with phenylalanine residues at position 16 have been excluded.

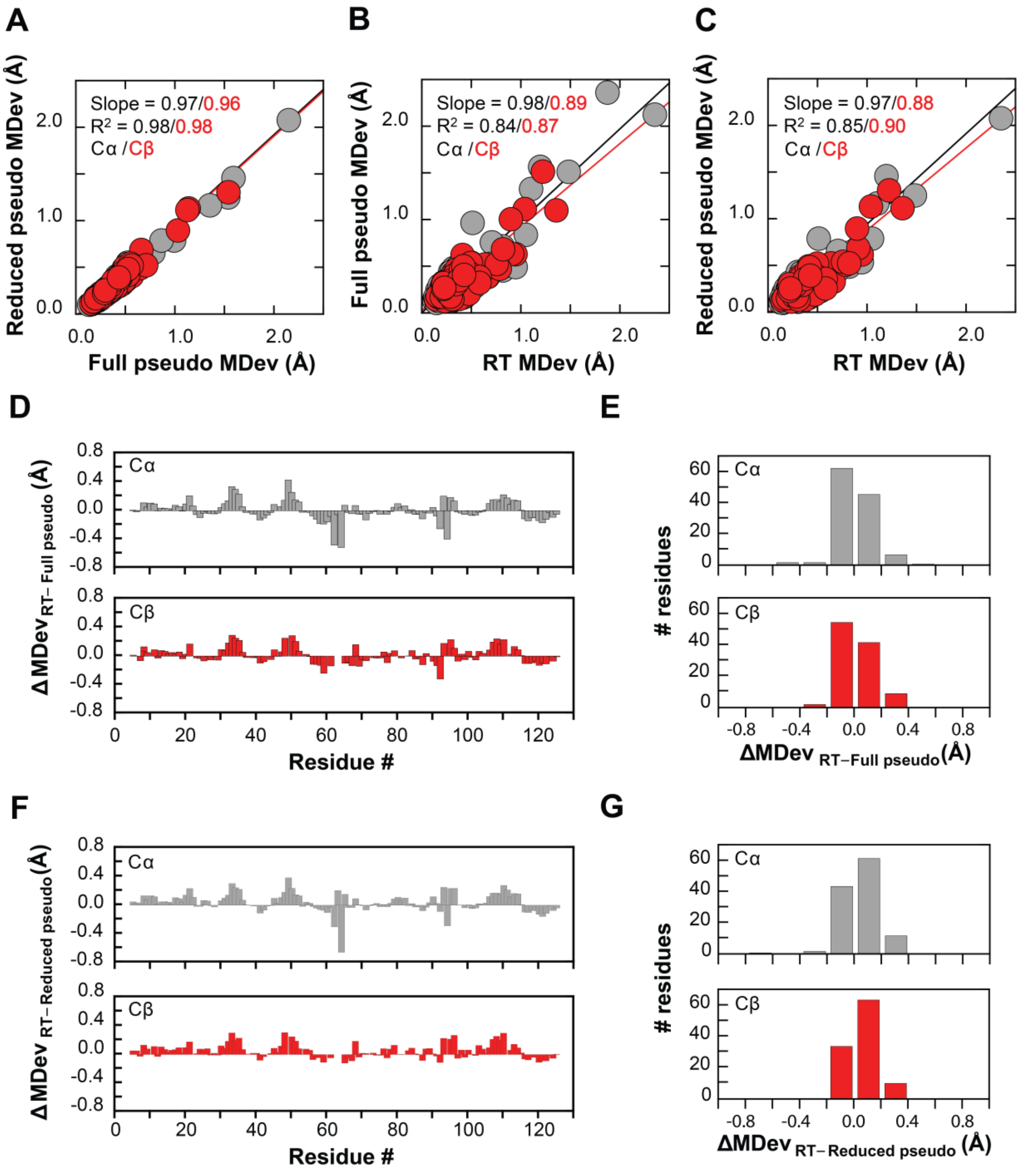

**Figure S17. Similar overall KSI conformational heterogeneity is obtained from the full pseudo-ensemble, the reduced pseudo-ensemble, and the RT-ensemble** (see Table S2 and Materials and Methods). Correlation plots of MDev values for the full pseudo-ensemble and reduced pseudo-ensembles (A), for RT-ensemble and full pseudo-ensembles, (B) and for RT-ensemble and reduced pseudo-ensembles (C); backbone (Cα, grey symbols) and side-chain (Cβ, red symbols) MDevs. ΔMDevs between (D) RT-ensemble and full pseudo-ensemble and (F) RT-ensemble and reduced pseudo-ensemble backbone (Cα, grey, top panels) and side-chain (Cβ, red bottom panels). All ΔMDevs for the full pseudo-ensemble *vs.* reduced pseudo-ensemble are

≤0.2 Å and are not shown. The histograms in (E) and (G) represent the ΔMDevs from D and F, respectively.

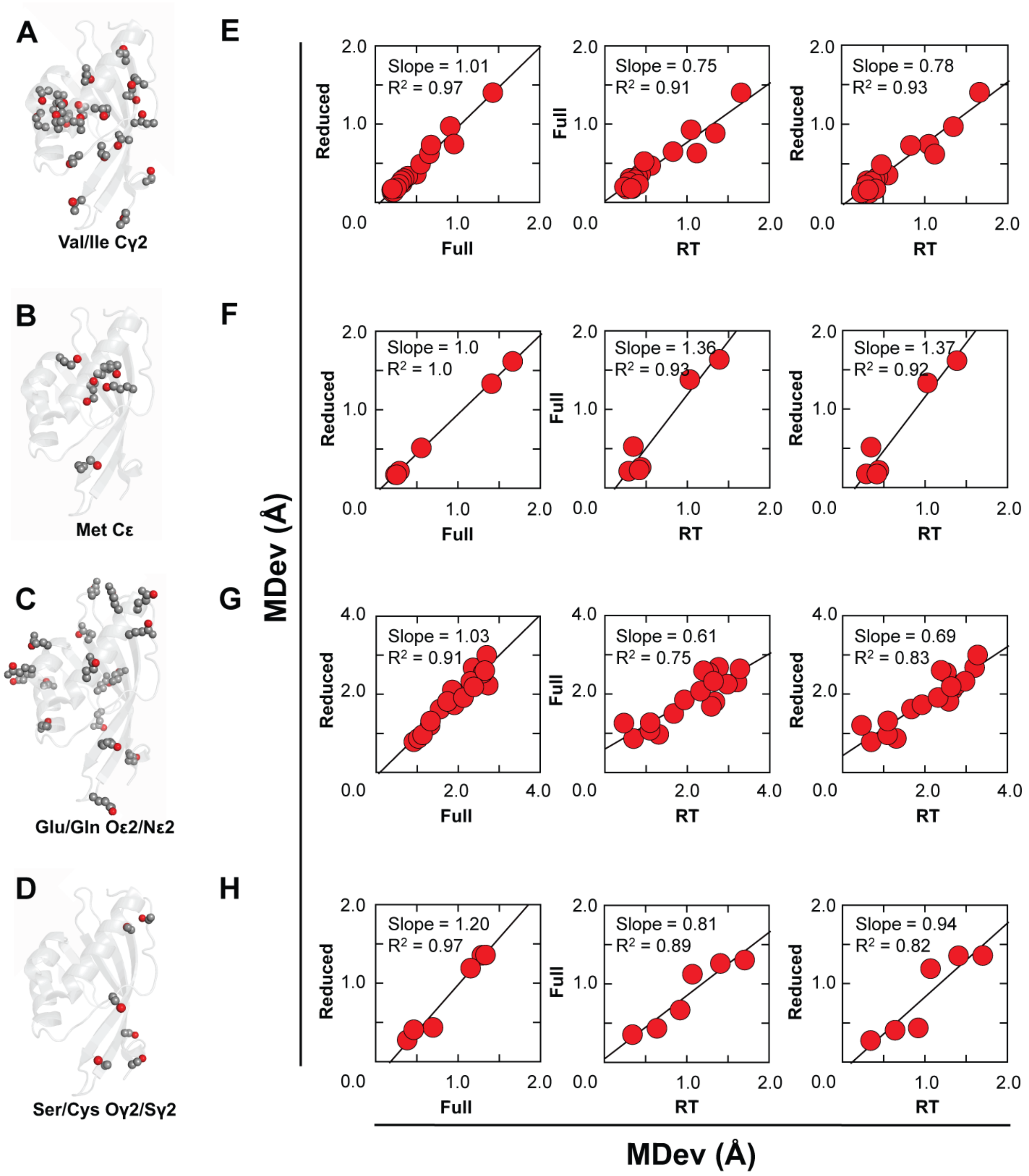

**Figure S18. High correlation coefficients indicate similar KSI side chain conformational heterogeneity obtained from the full pseudo-ensemble, the reduced pseudo-ensemble, and the RT-ensemble.** Cartoon representation of the KSI structure (PDB 1OH0) in which different classes of side-chains are depicted as spheres: Ile/Val and Met (hydrophobic, (A) and (B), respectively) and Glu/Gln and Ser/Cys (polar/charged, (C) and (D), respectively). Side chains are colored in grey and the red color indicates specific atoms for which conformational heterogeneity from the different types of KSI ensembles has been quantified and compared in E-H. Correlation plots of full pseudo-ensemble and reduced pseudo-ensemble (left row), RT-ensemble and full pseudo-ensemble (middle row) and RT-ensemble and reduced pseudo-ensemble (right row) MDevs for representative side chain atoms: valine/isoleucine Cγ2 (E), glutamate/glutamine Oε2/Nε2 (F), serine/cysteine Oγ2/Sγ2 (G), and methionine Cε (H).

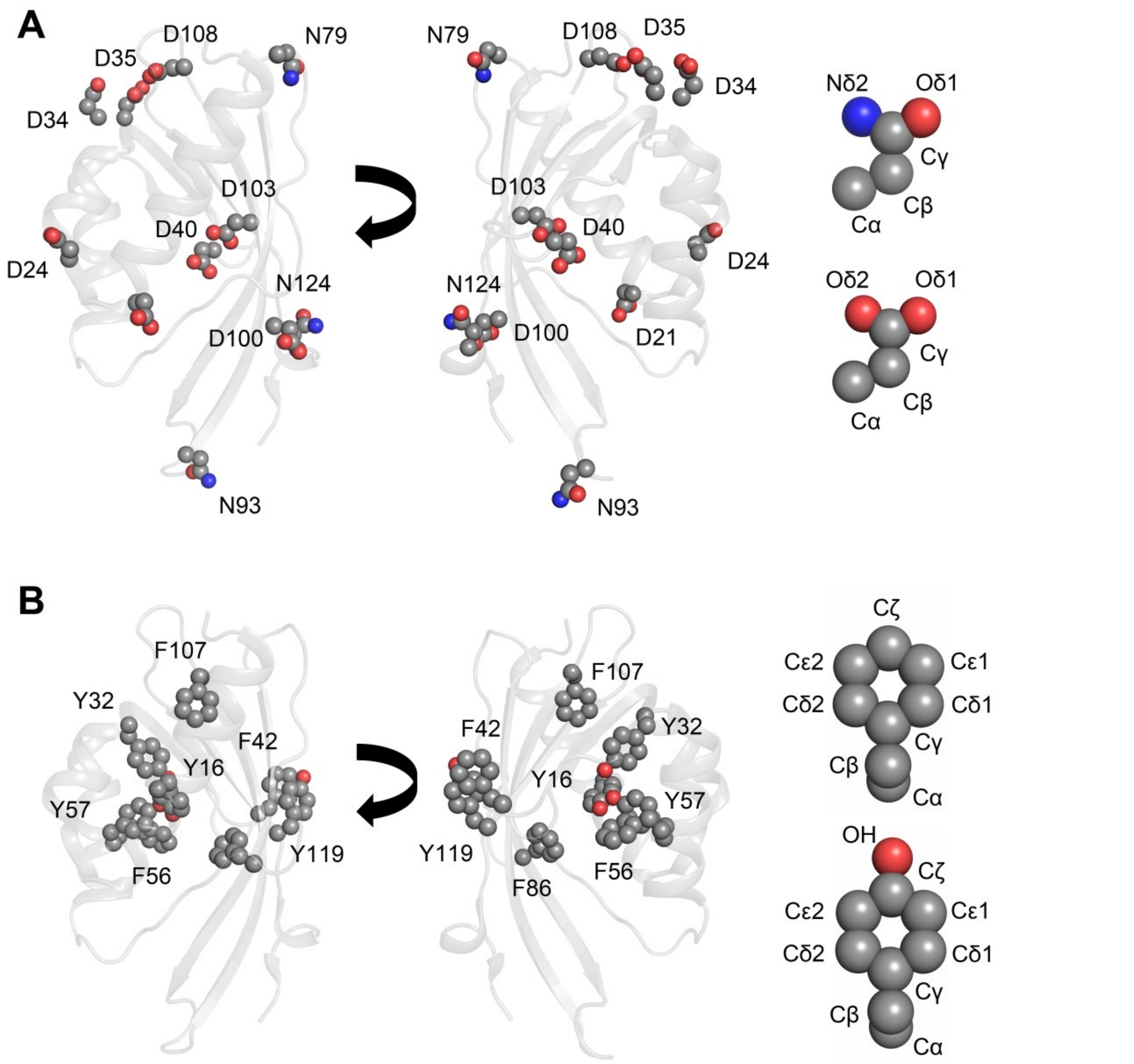

**Figure S19.** Cartoon depiction of KSI (PDB 1OH0) with all aspartate and asparagine residues (A) and all tyrosine and phenylalanine residues (B) represented as spheres. Right panels show aspartate, asparagine, tyrosine and phenylalanine side-chains and atom nomenclature. The assignment of Oδ1 *vs.* Oδ2 in aspartate residues is arbitrary, but consistent across all KSI crystal structures. N93 was not included in the analyses in this work, as it is situated within the flexible 91–96 loop.

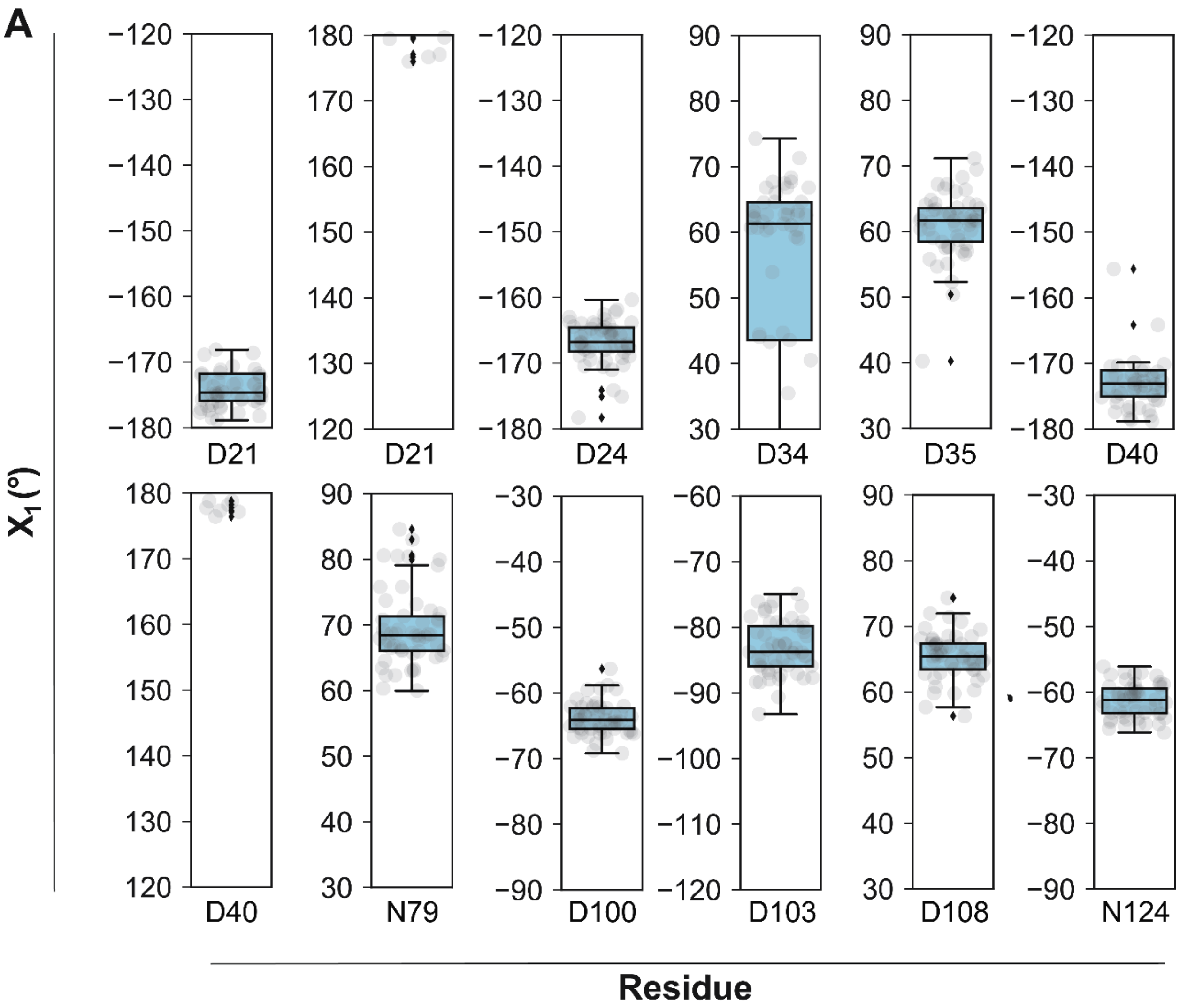
A
X1 (°)
Residue
D21
D21
D24
D34
D35
D40
D40
N79
D100
D103
D108
N124

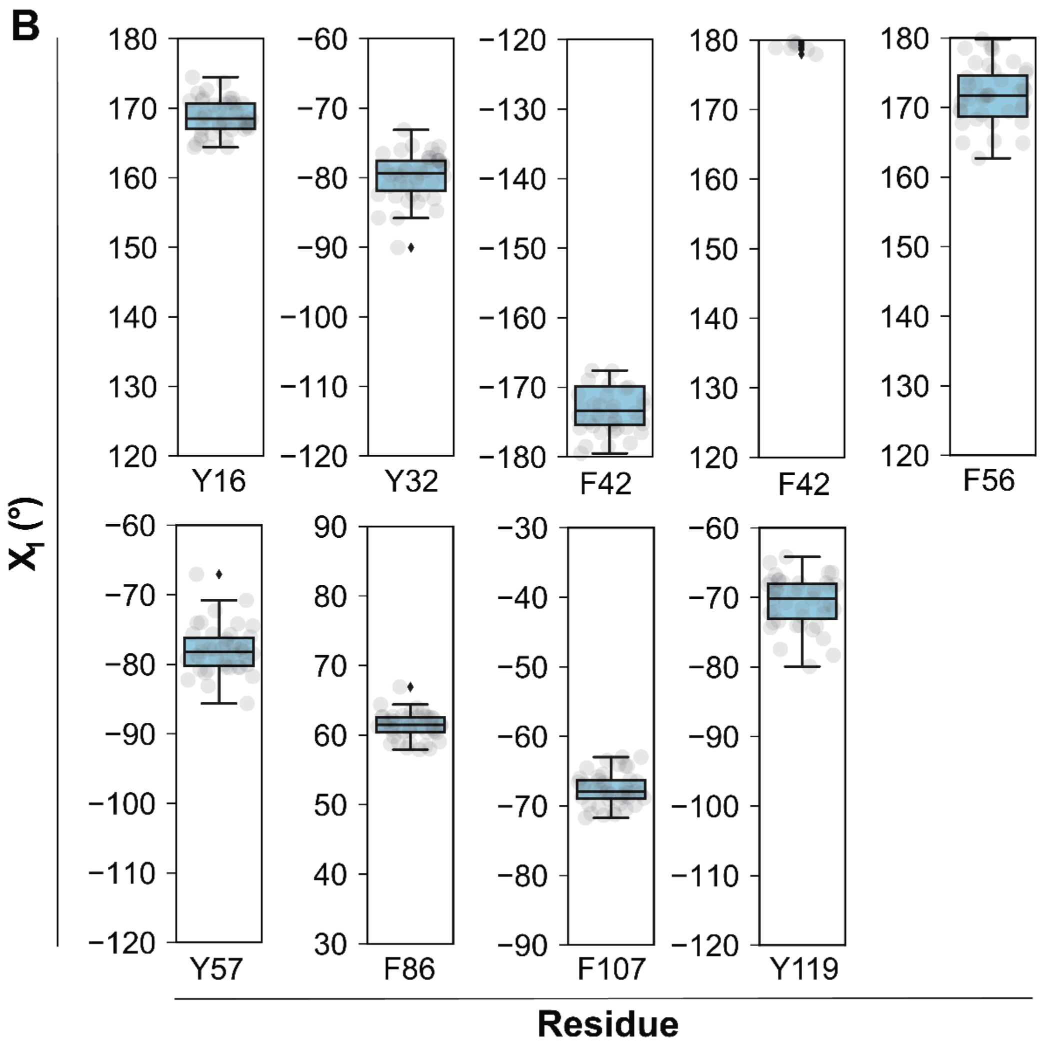

**Figure S20. Sidechain dihedral angles analysis provides additional evidence against exceptional positioning of oxyanion hole Y16 and D103 and general base D40**. Side chain χ1 dihedral angles for (A) all aspartate and asparagine residues and (B) tyrosine and phenylalanine residues from the reduced pseudo-ensemble. Asparagine residues at position 2 and position 93 have been excluded from the analysis because these residues are situated in the highly flexible N-terminus and 91-96 loop, respectively. χ1 angles for position 16 include tyrosine residues only (phenylalanine substitutions have been omitted).

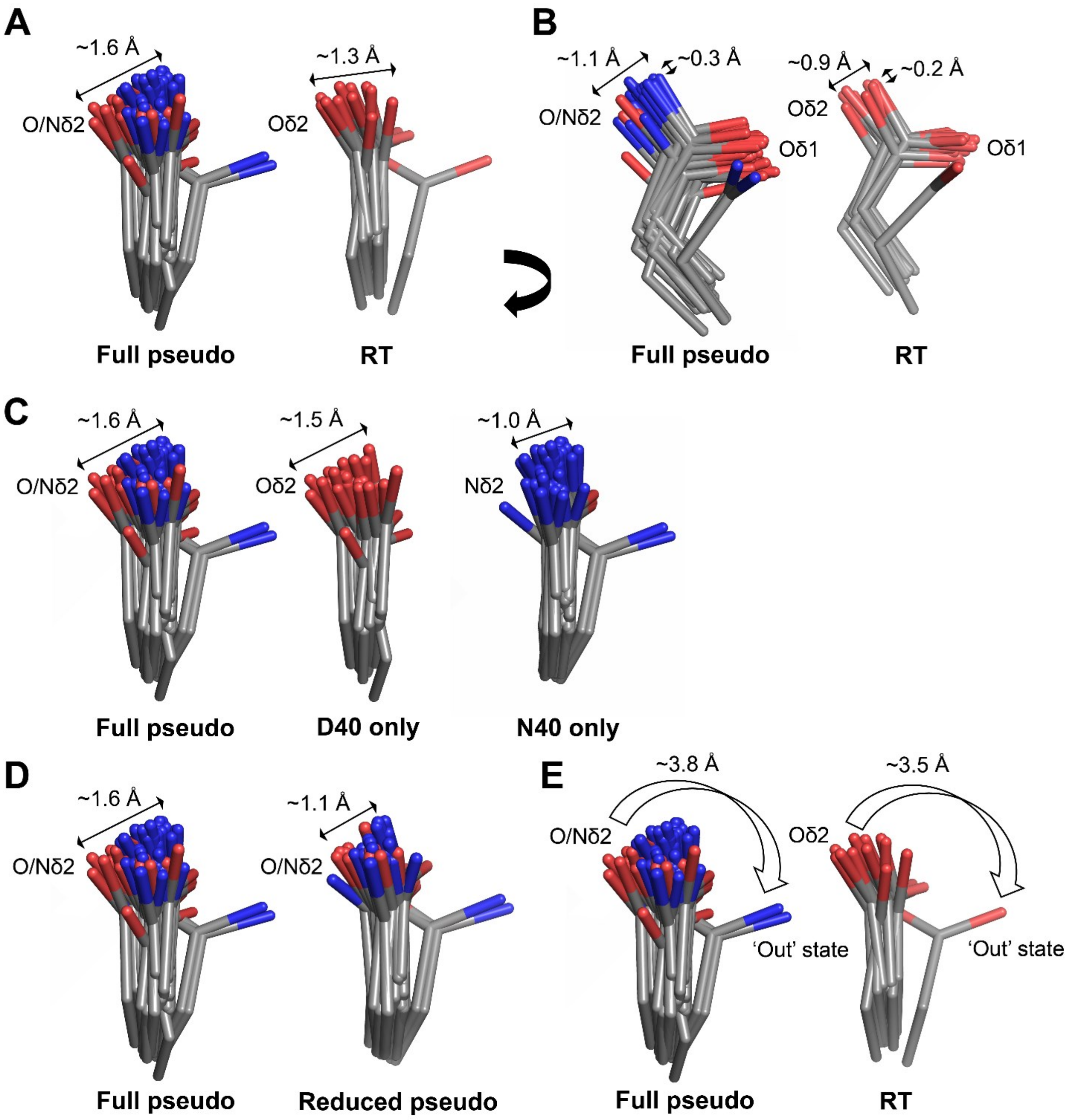

**Figure S21. Estimating the precision in positioning of the KSI general base.** Comparison of the general base D40 full pseudo-ensemble (left) and RT-ensemble (right) (A and B show orthogonal orientations); asparagine at position 40 mimics the protonated (intermediate) state of the KSI general base (1, 2). (C) Comparison of the full general base pseudo-ensemble (left) with sub-ensembles composed of aspartate only (middle) or asparagine only (right) at position 40. These ensembles exhibit similar extents of motion. (D) The full pseudo-ensemble (left) and the reduced pseudo-ensemble (right) exhibit similar extents of motion (Table S2). (E) The pseudo-ensemble and the RT-ensemble both provide evidence for general base 'out' state, suggesting that the KSI general base can undergo motion of up to ~4 Å.

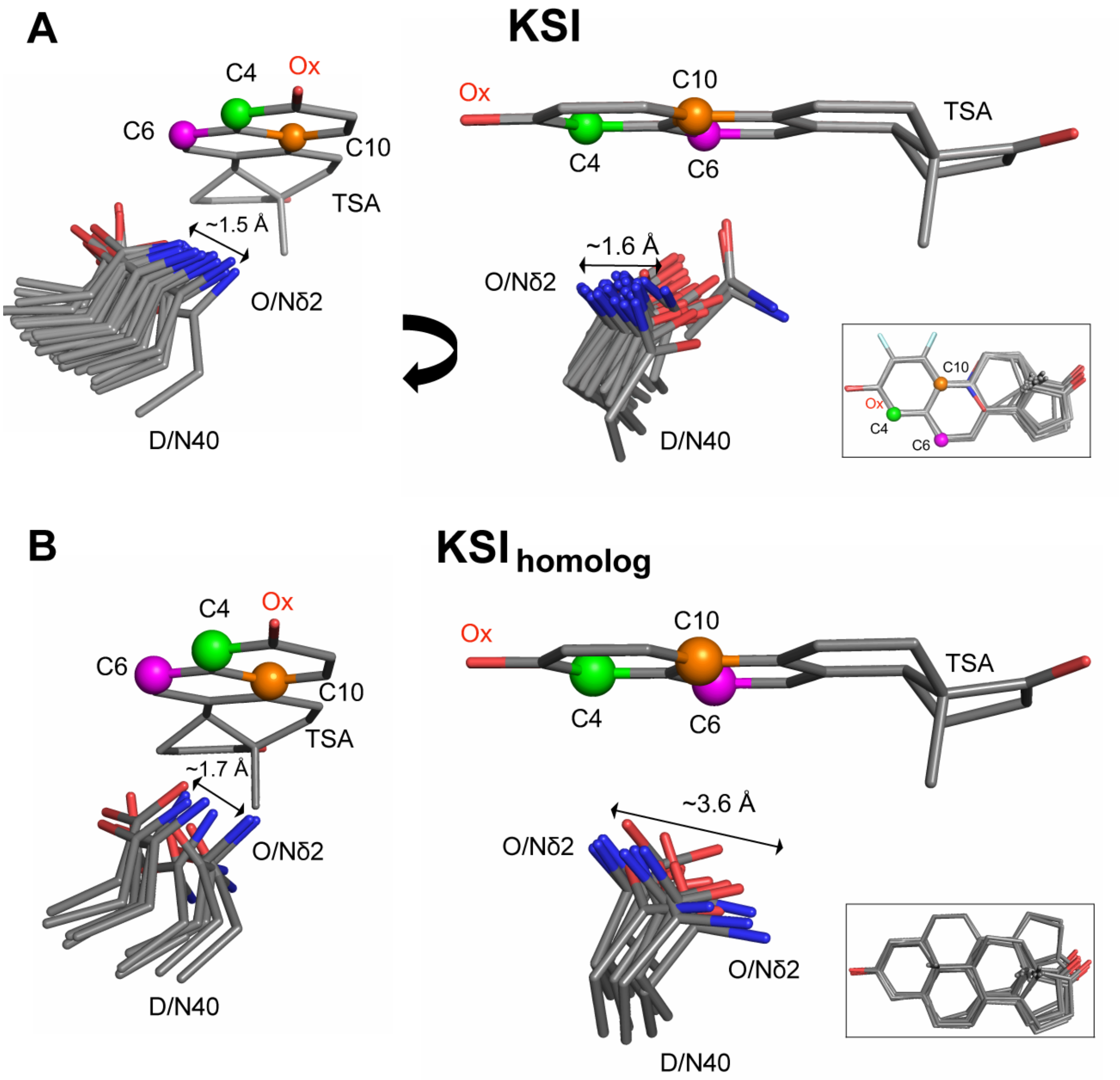

**Figure S22.** A bound TSA as "seen" by the KSI (A) and $KSI_{homolog}$ (B) general base in a TSA-bound ensemble of cryo crystal structures (n = 36 and 11, respectively) (see Table S2 for structures in (A) and Table S22 for structures in (B) (for (B) only $KSI_{homolog}$-TSA bound complexes for which the steroid is bound with its ring A facing the oxyanion hole are included (PDB codes 1OHP, 1QJG, 3NHX, 3NUV)). Panel A is reproduced from Figure 5D from the main text. The TSAs (equilenin and various phenols) have been aligned on the A ring with only one (PDB 1OH0) shown for clarity. The carbon positions between which protons are shuffled in KSI reactions are represented as green, magenta, and orange spheres (see Figure 1 and Figure S30 for the reaction mechanisms). The insets show the aligned TSAs.

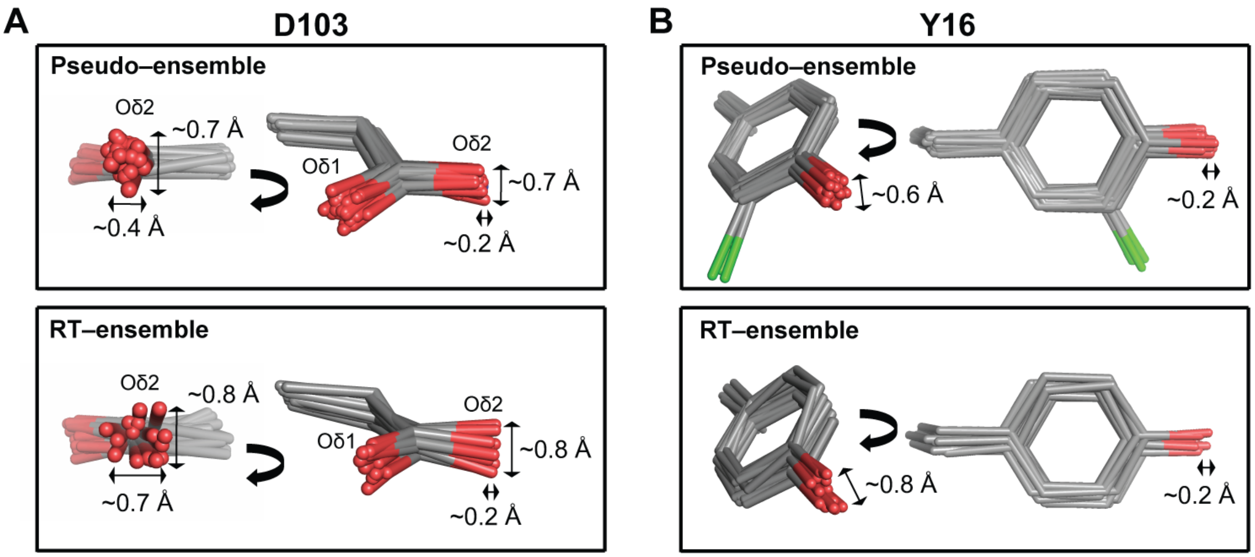

**Figure S23. Estimating the precision in positioning within the KSI oxyanion hole.** The oxyanion hole Y16 (A) and D103 (B) reduced pseudo-ensemble (top panels) and RT-ensembles (bottom panels). Chlorine atoms of Cl-tyrosine residues are colored in green.

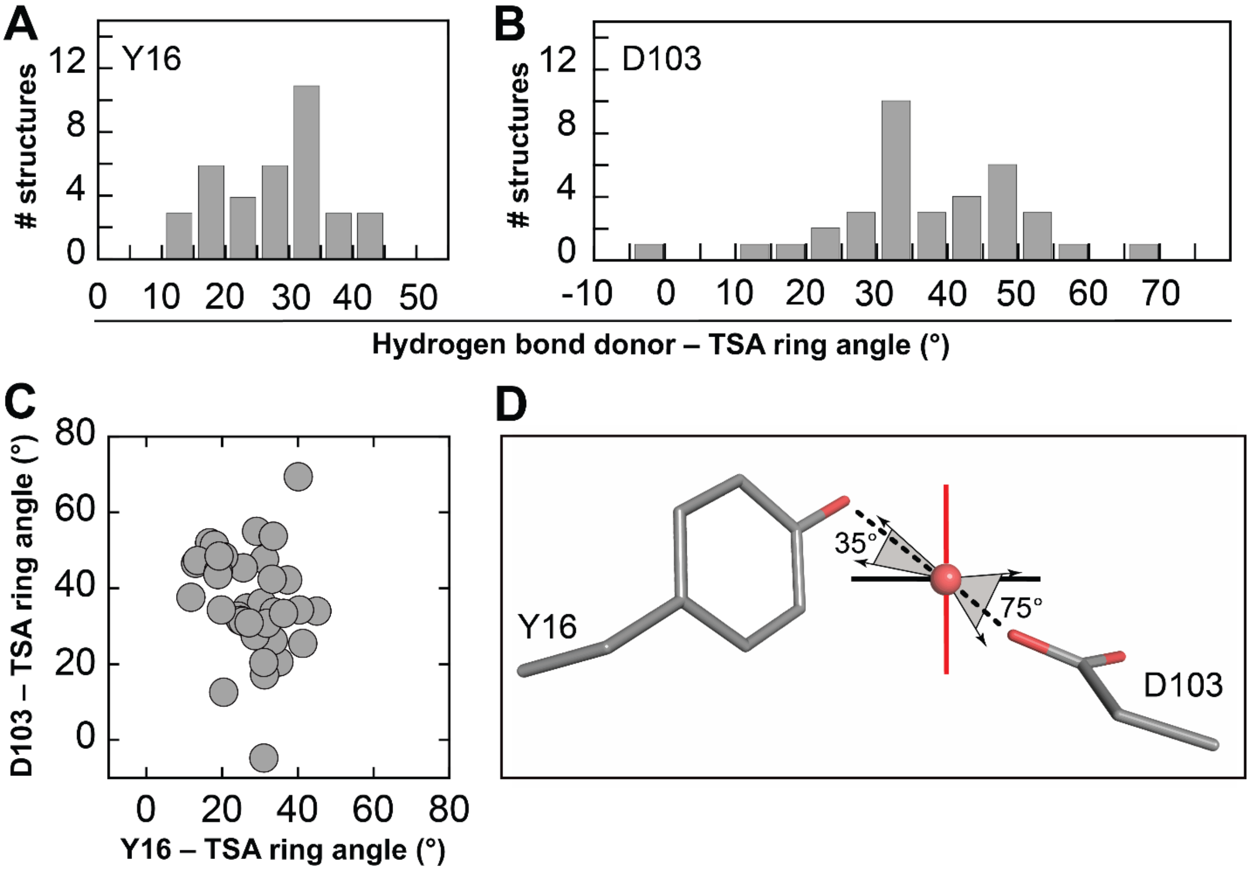

**Figure S24. The KSI oxyanion hole does not appear to be precisely positioned for ground state (sp$^2$) *vs.* transition state (sp$^3$) geometric discrimination.** Distribution of the angles between the hydrogen bond donors Y16 (A) and D103 (B) and the plane of steroid ligands from KSI crystal structures of variants with WT-like activity bound to TSAs (Table S2); angles range from 10° to 45° and from -5° to 75°, respectively. (C) Plot of the Y16 and D103 angles from A and B showing lack of correlation between Y16 and D103 hydrogen bond angles. (D) Cartoon representing the range of angles from (A) and (B) and the plane of a steroid ligand (black solid line). The black and red solid lines represent planes that are parallel and orthogonal to the steroid ring, respectively; the red line indicates the extreme plane from which sp$^3$ ligands could be more stabilized than sp$^2$ ligands. We note that rotation around C–O(H) bond in Y16 and D103 will allow the hydrogen bonding H (H is only expected to be observed in crystal structure of resolution better than 0.6-0.8 Å and therefore not observed in the crystal structures used in this work) to approach the ligand oxyanion from an additional range of angles, which could potentially contribute to the range of angles made with the plane of the steroid.

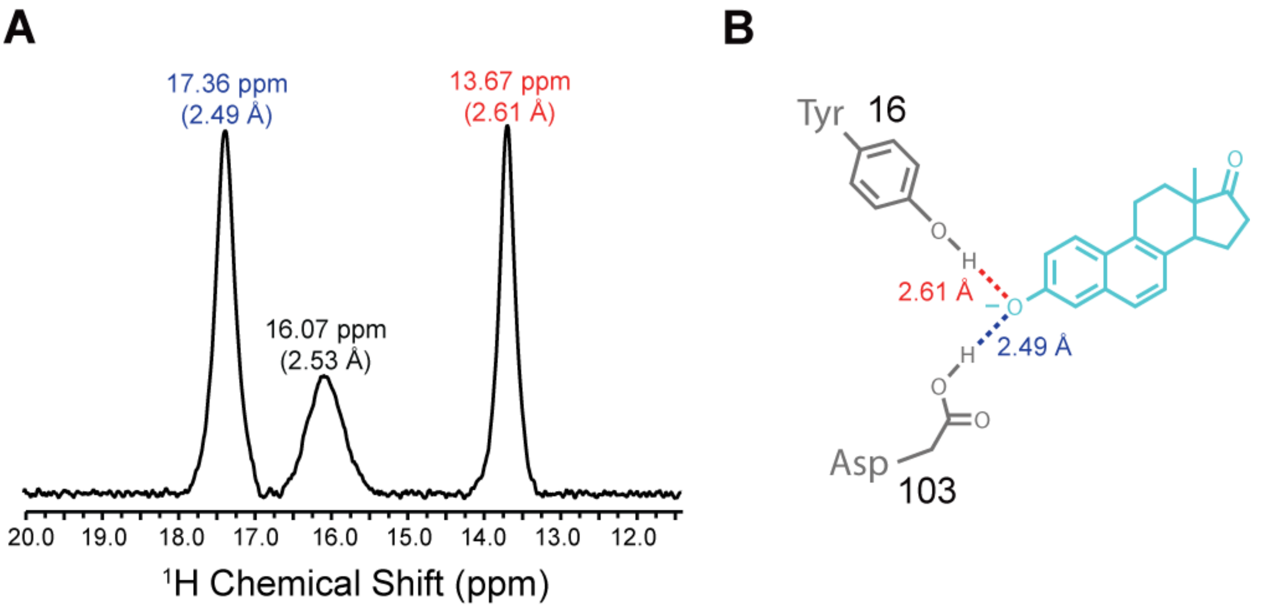

**Figure S25. Measuring Y16 and D103 hydrogen bond lengths in a KSI–TSA complex in solution by $^{1}$H NMR.** (A) $^{1}$H NMR spectra of KSI D40N variant bound to the TSA equilenin (see Materials and Methods for data collection). The D40N substitution mimics the protonated (intermediate) general base state and has been used to increase TSA affinity (1, 2). Shown are the chemical shifts for the $^{1}$H peaks (in ppm) in the downfield region of the spectrum. The respective hydrogen bonds lengths were calculated from the chemical shifts and using standard procedures (Harris & Mildvan, 1999; Pinney et al., 2018) (Materials and Methods). Three peaks are observed in the downfield region of the $^{1}$H spectrum. Comparison with previously published $^{1}$H NMR spectra of KSI bound to a variety of TSAs and consideration of the coupling between the hydrogen bond lengths suggests that the 13.67 ppm and the 17.36 ppm peaks (2.61 Å and 2.49 Å lengths, respectively) correspond to the Y16–TSA and D103–TSA hydrogen bonds, respectively (1, 3) Alternative assignments are possible but do not alter any of the conclusions in the main text. We note that the peak at 16.07 ppm could originate from alternative hydrogen bonding conformations or from a non-active site hydrogen bond, possibilities that will be investigated in future work. (B) Schematic depiction of the oxyanion hole residues Y16 and D103 (in grey) and the bound TSA (in cyan) and the assigned hydrogen bond lengths (Y16–TSA in red and D103–TSA in blue).

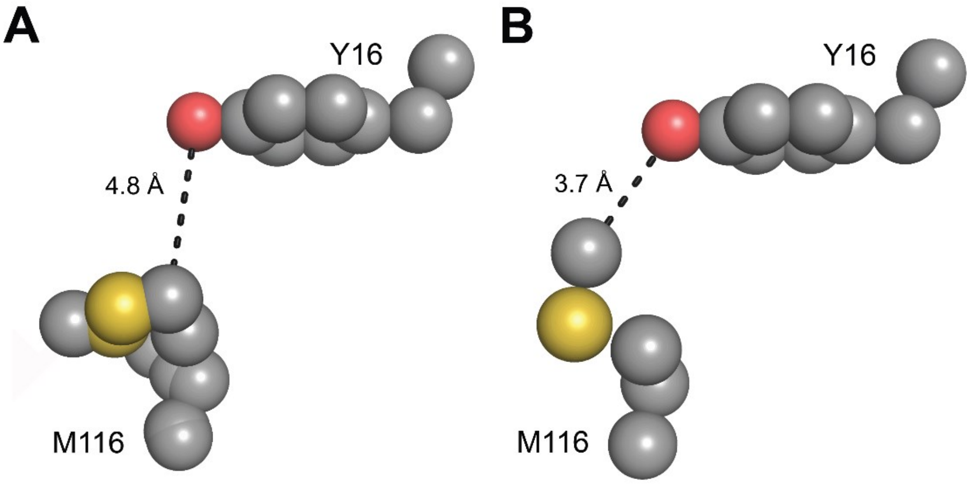

**Figure S26. Packing interactions cannot be uniquely evaluated from single X-ray crystallography models.** Distance between M116 and Y16 could be interpreted as beyond van der Waals contact radius (‘loose’ packing, (A), PDB 5D81) or within this contact radius (‘tight’ packing, (B), PDB 5D83), depending on the analyzed crystal structure. The $r_{vdw}$ methyl is ~2.0 Å, $r_{vdw}$ oxygen is 1.4–1.7 Å (see Table S48).

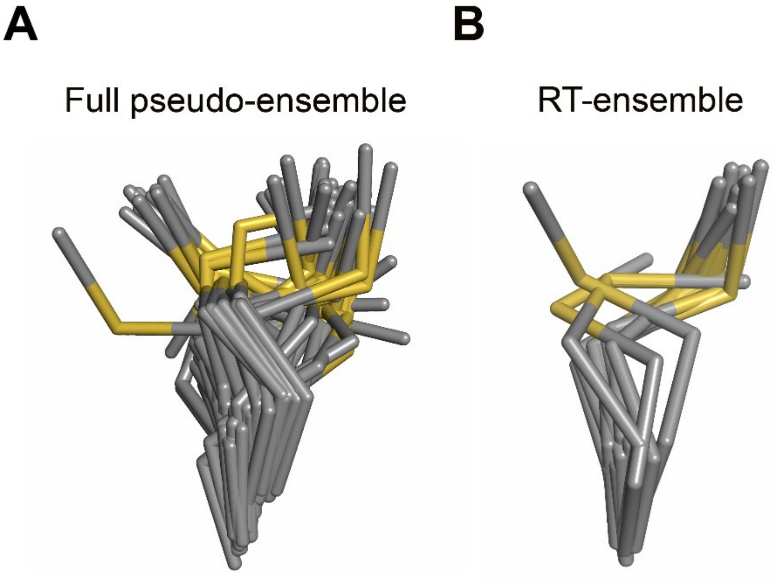

**Figure S27. M116 has a broad conformational ensemble.** (A) The M116 full pseudo-ensemble and (B) and RT-ensemble.

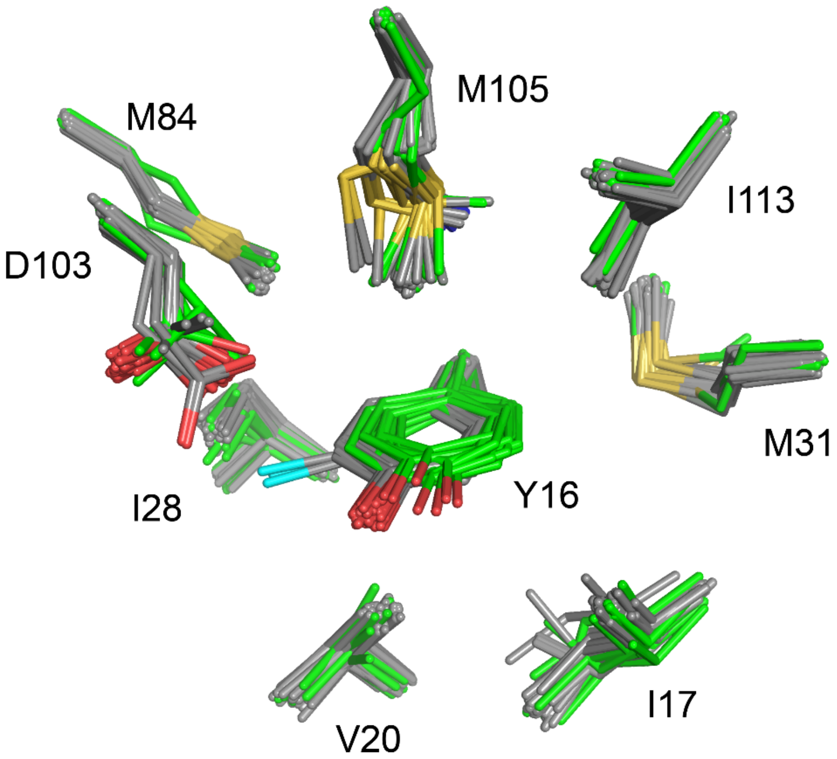

**Figure S28. Ablation of the Y16–Y57 hydrogen bond and the increased flexibility of the residue at position 16 does not lead to significant rearrangements or enhanced mobility in the surrounding residues**. Shown are all structures with intact (grey, n = 70) or ablated (green, n = 15) Y16–Y57 hydrogen bond; position 103 includes various substitutions (see also Figures S16 and 29).

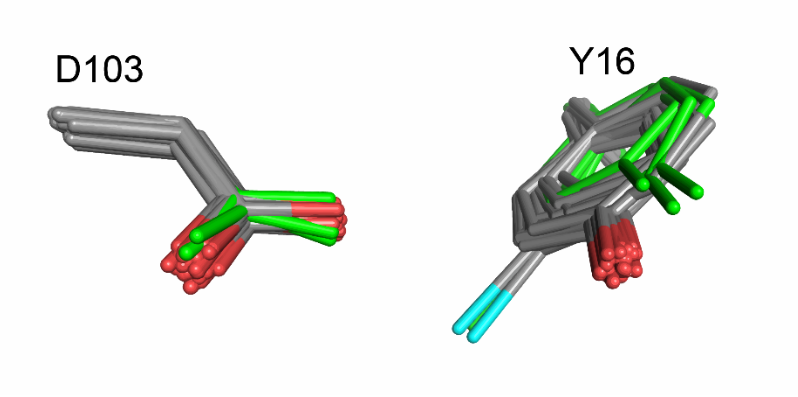

**Figure S29. Increased flexibility or mispositioning of Y16 does not impact D103 positioning.** Shown are the reduced pseudo-ensemble (grey, Table S2) and a subset of KSI structures with intact Y16 and D103, in which the Y16-Y57 hydrogen bond is ablated via Y57 substitutions (green; PDB 1DMM, Y57F and PDB 1K41, Y57S).

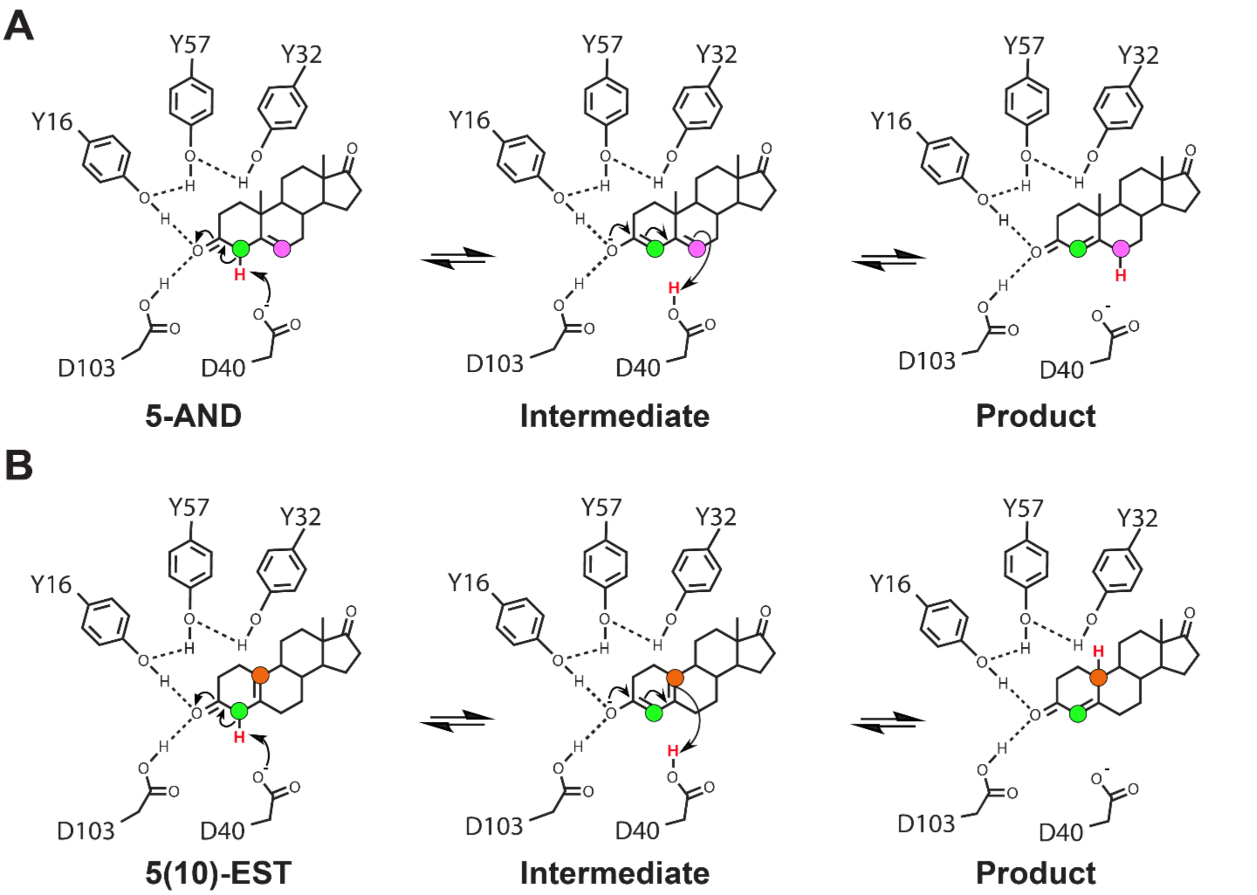

**Figure S30.** KSI reaction mechanism with the steroid substrates 5-androstenedione (5-AND, (A)) and 5(10)-estrene-3,17-dione (5(10)-EST, (B)). The shuffled proton is colored in red. The donating and accepting protons (forward reaction) are colored in green and magenta for 5-AND, respectively, and green and orange for 5(10)-EST, respectively; the same colors are used in Figure 5 in the main text to map donating and accepting carbon positions onto KSI-bound TSAs.

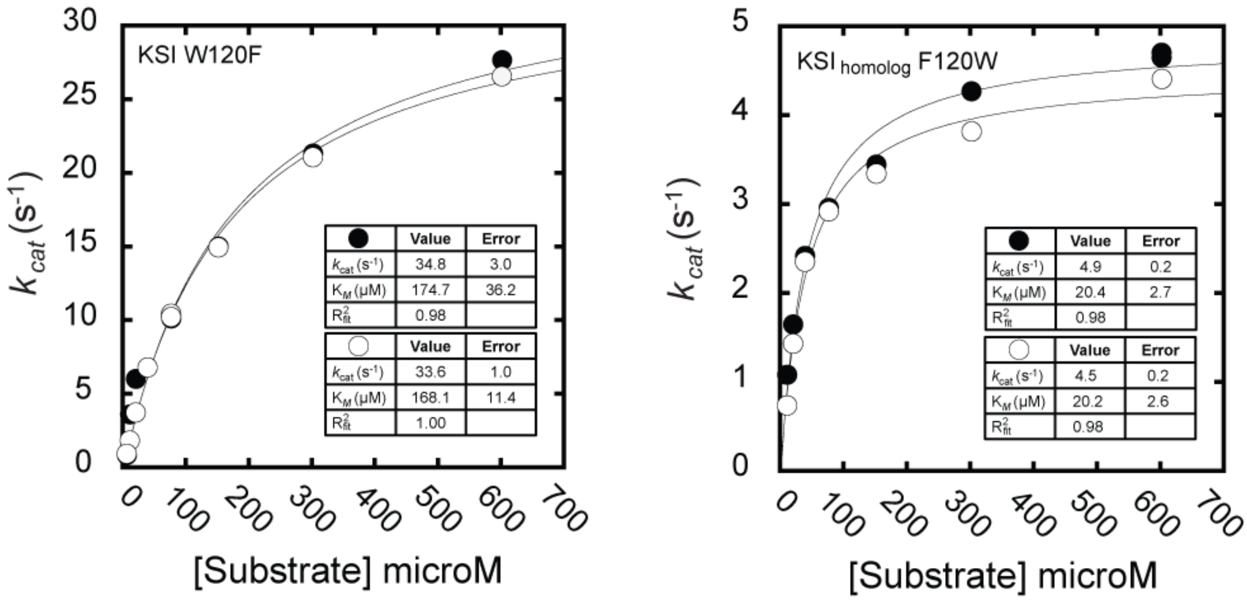

**Figure S31. Michaelis-Menten kinetics of KSI W120F and KSI$_{homolog}$ F120W.** Kinetics were measured with the substrate 5(10)-Estrene-3,17-dione as the chemical step for this substrate is rate limiting (4). Kinetics were measured in duplicate (both shown) with the enzyme concentration varied ~3 fold (Table S58).

| Structure (PDB code) | Mutations | Ligand | Resolution (Å) | Space group | # molecules asymmetric unit |
|---|---|---|---|---|---|
| 1DMM | Y57F | - | 1.9 | $C222_1$ | 1 |
| 1DMN | Y32F/Y57F | - | 2.05 | $C222_1$ | 1 |
| 1DMQ | Y32F | - | 2.15 | $C222_1$ | 1 |
| 1E97 | Y16F/Y32F/Y57F | - | 2.0 | $C222_1$ | 1 |
| 1EA2 | Y16F | - | 1.8 | $C222_1$ | 1 |
| 1GS3 | Y32F/Y57F/Y119F/D40N | Equilenin | 2.1 | $C222_1$ | 1 |
| 1OHO | Y16F/D40N | Equilenin | 1.9 | $C222_1$ | 1 |
| 1OPY | WT | - | 1.9 | $C222_1$ | 1 |
| 1W02 | Y16F/D103L | - | 2.3 | $C222_1$ | 1 |
| 1W6Y | W92A | Equilenin | 2.1 | $C222_1$ | 1 |
| 2INX | D40N | 2,6-difluorophenol | 1.5 | $C222_1$ | 1 |
| 3CPO | D40N | 2-fluorophenol | 1.24 | $C222_1$ | 1 |
| 3RGR | M116A | - | 1.59 | $C222_1$ | 1 |
| 3SED | M105A | - | 1.3 | $C222_1$ | 1 |
| 4K1V | Y16F/Y57F | - | 1.8 | $C222_1$ | 1 |
| 5AI1 | Y32F/Y57F/Y119F/D40N | Equilenin | 2.1 | $C222_1$ | 1 |
| 1CQS | D103E/D40N | Equilenin | 1.9 | $C2_1$ | 2 |
| 1E3R | D40N | Androsten-3β-ol-17-one | 2.5 | $C2_1$ | 2 |
| 1E3V | WT | Deoxycholate | 2.0 | $P2_12_12_1$ | 2 |
| 1K41 | Y57S | - | 2.2 | $C2_1$ | 2 |
| 1OGX | D40N | Equilenin | 2.0 | $C2_1$ | 2 |
| 1OH0 | WT | Equilenin | 1.1 | $C2_1$ | 2 |
| 1VZZ | Y32F/D103L | - | 2.3 | $P2_12_12_1$ | 2 |
| 1W00 | D103L | - | 2.2 | $P2_12_12_1$ | 2 |
| 1W01 | Y57F/D103L | - | 2.2 | $P2_12_12_1$ | 2 |
| 3FZW | D40N/D103N | Equilenin | 1.32 | C21 | 2 |
| 3OWS | D40N/C69S/C81S/C97S/M116C-CN | Equilenin | 1.71 | $C2_1$ | 4 |
| 3OWU | D40N/C69S/C81S/C97S/F86C-CN | Equilenin | 1.7 | $P2_1$ | 4 |
| 3VGN | D40N | 3-fluoro-4-nitrophenol | 1.3 | $P2_12_12_1$ | 2 |
| 3VSY | WT | - | 1.5 | $P2_12_12_1$ | 2 |
| 4K1U | Y16F/Y32F | - | 2.0 | $P2_12_12_1$ | 2 |
| 5D83 | D40N, Y32 (Cl-Y) | - | 1.7 | $P2_12_12_1$ | 2 |
| 5D82 | D40N, Y16 (Cl-Y) | - | 1.37 | $P2_12_12_1$ | 2 |
| 5D81 | D40N, Y57 (Cl-Y) | - | 1.39 | $C222_1$ | 1 |
| 5KP4 | WT | 19-nortestosterone | 1.71 | $P2_12_12_1$ | 2 |
| 2PZV | D40N | Phenol | 1.25 | P1 | 4 |
| 3IPT | Y16S/D40N | Equilenin | 1.63 | $P2_1$ | 4 |
| 3OWY | D40N/C69S/C81S/C97S/M105C-CN | Equilenin | 2.3 | $P2_1$ | 8 |
| 3OX9 | D40N/C69S/C81S/C97S/F86C-CN | - | 2.0 | $P2_1$ | 4 |
| 3OXA | D40N/C69S/C81S/C97S/M116C-CN | - | 1.89 | $P2_1$ | 4 |
| 3T8N | Y16AD103A | - | 1.47 | $C2_1$ | 4 |
| 5KP1 | D40N, Y16(Cl-Y) | Equilenin | 1.22 | P1 | 4 |
| 5KP3 | D40N, Y57(Cl-Y) | Equilenin | 1.7 | $P2_12_12_1$ | 2 |
| 5G2G | M116K | Equilenin | 1.6 | $P2_12_12_1$ | 2 |
| 1C7H | R75A | - | 2.5 | $C2_12_12_1$ | 1 |

**Table S1.** KSI cryo crystal structures available from the Protein Data Bank (PDB).

| PDB | Pseudo-ensemble | | | | | | | |
|---|---|---|---|---|---|---|---|---|
| | Full | Apo | TSA-bound | Reduced | TSA-bound* | Intact Y16–Y57 Hbond | Ablated Y16–Y57 Hbond[#] | Equilenin-bound[%] |
| 1DMM | X | X | | | | | X | |
| 1DMN | X | X | | | | | X | |
| 1DMQ | X | X | | | | X | | |
| 1E97 | X | X | | X | | | X | |
| 1EA2 | X | X | | X | | | X | |
| 1GS3 | X | | X | | X | | X | |
| 1OHO | X | | X | X | | | X | |
| 1OPY | X | X | | X | | X | | |
| 1W02 | X | X | | | | | X | |
| 1W6Y | X | | X | | X | X | | X |
| 2INX | X | | X | X | | X | | |
| 3CPO | X | | X | X | X | X | | |
| 3RGR | X | X | | X | | X | | |
| 3SED | X | X | | X | | X | | |
| 4K1V | X | X | | X | | | X | |
| 5AI1 | X | | X | | X | | X | |
| 1CQS | X | | X | | | X | | |
| 1E3R | X | | | | | X | | |
| 1E3V | X | | | X | | X | | |
| 1K41 | X | X | | | | | X | |
| 1OGX | X | | X | X | X | X | | X |
| 1OH0 | X | | X | X | X | X | | X |
| 1VZZ | X | X | | | | X | | |
| 1W00 | X | X | | | | X | | |
| 1W01 | X | X | | | | | X | |
| 3FZW | X | | X | X | X | X | | |
| 3OWS | X | | X | X | X | X | | X |
| 3OWU | X | | X | | X | X | | X |
| 3VGN | X | | X | X | X | X | | |
| 3VSY | X | X | | X | | X | | |
| 4K1U | X | X | | X | | | X | |
| 5D83 | X | X | | X | | X | | |
| 5D82 | X | X | | X | | X | | |
| 5D81 | X | X | | X | | X | | |
| 5KP4 | X | | | X[2] | | X[2] | | |
| 2PZV | X | | X | X | X | X | | |
| 3IPT | X | | X | X | | | | |
| 3OWY | X | X[1] | X | | X | X | | X |
| 3OX9 | X | X | | | | X | | |
| 3OXA | X | X | | X | | X | | |
| 3T8N | X | X | | | | | | |
| 5KP1 | X | | X | X | X | X | | |
| 5KP3 | X | | X | X | X | X | | |
| 5G2G | X | | X | X | | X | | |
| 1C7H | X | X | | | | X | | |
| # KSI monomers | 94 | 42 | 46 | 54 | 36 | 70 | 15 | 19 |

*TSA-bound structures with activity within ~10–fold from WT and containing both Asp and Asn residues at position 40. TSAs include equilenin and various phenols. Even though D40N substitution leads to a rate decrease, the Asn at position 40 mimics the protonated (intermediate) state of the general base and increases the KSI affinity for TSAs (1, 2); # KSI crystals structures in which Y16 – Y57 hydrogen bond (Hbond) is ablated but the phenyl ring at position 16 is preserved (Y16F or Y57X substitutions, X being any residue). % Equilenin-bound structures with intact oxyanion hole residues, activity within ~10–fold from WT and containing both Asp and Asn residues at position 40. [1] Although the structure has been reported as TSA-bound, two out of the 8 KSI molecules in the asymmetric unit were Apo and were thus included in the Apo pseudo-ensemble. [2] The asymmetric unit of this crystal structure contains two molecules and the molecule B was not bound to a GSA. Further, in molecule B, Y16 orientation appears misaligned when compared to all known KSI crystal structures to date and was not included in the pseudo-ensembles in this work.

**Table S2**. KSI crystal structures used to obtain the various pseudo-ensembles in this work. To most accurately estimate the precision in positioning in the KSI oxyanion hole and remove potential artifacts, the KSI reduced pseudo-ensemble does not include structures with i) mutations in the oxyanion hole that alter the chemical nature of the hydrogen bonding groups (e.g. D103L mutations); ii) mutations in the Y16 hydrogen bond network (e.g. Y57F) as these mutations have been suggested to alter Y16 positioning (3); iii) resolution worse than 2.0 Å. We did not exclude structures with Y16F mutations, as this mutation did not appear to alter D103 positioning (Figure S16), further suggesting that Y16 and D103 hydrogen bonding orientations are not coupled.

| | Apo | | GSA-bound | | | TSA-bound | |
|---|---|---|---|---|---|---|---|
| | **250 K** | **280 K** | **100 K** | **250 K** | **280 K** | **250 K** | **280 K** |
| **PDB code** | **6UCW** | **6U1Z** | **6UBQ** | **6UCY** | **6TZD** | **6UCN** | **6U4I** |
| **Data collection*** | | | | | | | |
| **Wavelength (Å)** | 0.88557 | 0.88557 | 0.78719 | 0.88557 | 0.88557 | 0.88557 | 0.88557 |
| **Resolution range** | 36.72-1.25 (1.27-1.25) | 36.97-1.5 (1.53-1.50) | 37.16-1.30 (1.32-1.30) | 36.06-1.15 (1.17-1.15) | 37.19-1.45 (1.48-1.45) | 35.91-1.32 (1.34-1.32) | 36.83-1.55 (1.58-1.55) |
| **Space group** | $P2_12_12_1$ | $P2_12_12_1$ | $P2_12_12_1$ | $P2_12_12_1$ | $P2_12_12_1$ | $P2_12_12_1$ | $P2_12_12_1$ |
| **Unit cell** | 35.85 73.44 95.99 90 90 90 | 36.03 73.94 95.69 90 90 90 | 36.21 74.32 95.56 90 90 90 | 36.06 73.85 95.62 90 90 90 | 36.23 74.39 95.35 90 90 90 | 35.91 73.72 95.73 90 90 90 | 35.54 73.66 95.68 90 90 90 |
| **Total reflections** | 336888 (16253) | 227920 (10482) | 851076 (37730) | 594265 (27460) | 256187 (12435) | 286924 (13737) | 162032 (7526) |
| **Unique reflections** | 70573 (3438) | 41654 (1902) | 64423 (3045) | 89597 (4212) | 46456 (2224) | 58753 (2829) | 37064 (1762) |
| **Multiplicity** | 4.8 (4.7) | 5.5 (5.5) | 13.2 (12.4) | 6.6 (6.5) | 5.5 (5.6) | 4.9 (4.9) | 4.4 (4.3) |
| **Completeness (%)** | 99.5 (98.6) | 98.6 (95.3) | 99.8 (97.4) | 98.1 (94.4) | 99.8 (99.1) | 97.3 (95.2) | 99.4 (98.0) |
| **Mean I/sigma(I)** | 9.9 (1.1) | 10.9 (1.1) | 19.4 (1.4) | 11.8 (1.2) | 18.0 (1.1) | 12.4 (1.0) | 11.8 (1.1) |
| **R-merge** | 0.076 (1.459) | 0.074 (1.737) | 0.062 (1.972) | 0.069 (1.504) | 0.048 (1.685) | 0.066 (1.446) | 0.075 (1.457) |
| **R-meas** | 0.085 (1.644) | 0.082 (1.916) | 0.064 (2.058) | 0.075 (1.632) | 0.054 (1.863) | 0.074 (1.621) | 0.085 (1.658) |
| **R-pim** | 0.038 (0.746) | 0.035 (0.796) | 0.018 (0.577) | 0.029 (0.626) | 0.023 (0.783) | 0.033 (0.720) | 0.040 (0.776) |
| **$CC_{1/2}$** | 0.997 (0.410) | 0.997 (0.466) | 0.995 (0.586) | 0.999 (0.583) | 0.999 (0.557) | 0.999 (0.430) | 0.999 (0.505) |
| **Refinement*** | | | | | | | |
| **Model type** | Multi-conformer | Traditional | Traditional | Multi-conformer | Traditional | Multi-conformer | Traditional |
| **Resolution range** | 36.72-1.25 (1.27-1.25) | 36.97-1.50 (1.53-1.50) | 34.64-1.30 (1.32- 1.30) | 32.40-1.15 (1.16-1.15) | 34.65-1.45 (1.48-1.45) | 32.29-1.32 (1.34-1.32) | 36.84-1.55 (1.59-1.55) |
| **Unique reflections used in refinement**** | 70423 (2569) | 41570 (2498) | 64304 (2532) | 88966 (2689) | 46342 (2652) | 58644 (2549) | 36978 (2629) |
| **Unique reflections used for R-free** | 3491 (145) | 2035 (136) | 3176 (135) | 4435 (145) | 2273 (135) | 2894 (137) | 1847 (138) |
| **R-work** | 0.1488 (0.2943) | 0.1360 (0.2528) | 0.1520 (0.2681) | 0.1456 (0.2672) | 0.1456 (0.2915) | 0.1444 (0.3078) | 0.1367 (0.2465) |
| **R-free** | 0.1731 (0.2771) | 0.1668 (0.2965) | 0.1699 (0.3067) | 0.1636 (0.2595) | 0.1741 (0.3415) | 0.1738 (0.3323) | 0.1752 (0.3190) |
| **non-hydrogen atoms** | 5018 | 2480 | 2565 | 5351 | 2436 | 5294 | 2420 |
| **macromolecules** | 4719 | 2317 | 2228 | 4970 | 2265 | 4949 | 2223 |
| **ligands** | 4 | 3 | 45 | 67 | 45 | 84 | 43 |
| **solvent** | 295 | 160 | 292 | 314 | 126 | 261 | 154 |
| **Protein residues** | 255 | 254 | 254 | 259 | 252 | 255 | 254 |
| **RMS(bonds)** | 0.008 | 0.009 | 0.008 | 0.009 | 0.008 | 0.009 | 0.010 |
| **RMS(angles)** | 1.25 | 0.91 | 0.97 | 1.11 | 0.94 | 1.06 | 1.00 |
| **Ramachandran favored (%)** | 97.5 | 98.8 | 97.6 | 95.7 | 98.8 | 97.6 | 98.0 |
| **Ramachandran allowed (%)** | 2.5 | 1.2 | 2.4 | 4.2 | 1.2 | 2.4 | 2.0 |
| **Ramachandran outliers (%)** | 0.0 | 0.0 | 0.0 | 0.1[#] | 0.0 | 0.0 | 0.0 |
| **Average B-factor** | 17.9 | 28.7 | 27.0 | 15.4 | 30.6 | 17.2 | 24.9 |
| **macromolecules** | 17.1 | 28.0 | 25.9 | 14.6 | 29.9 | 16.5 | 24.0 |
| **ligands** | 20.0 | 29.1 | 31.5 | 20.4 | 39.2 | 19.5 | 29.2 |
| **solvent** | 29.5 | 39.1 | 34.5 | 27.6 | 39.4 | 30.2 | 35.9 |

* values in parenthesis are for the highest resolution shell; ** values in parenthesis indicate the number of reflections (working set) in the highest resolution shell; [#] 1 out of 629 peptide bonds represents an outlier with clear electron density.

**Table S3.** X-ray diffraction data collection and model refinement statistics.

| 280 K<br>100K | 6U1Z A (Apo) | 6U1Z B (Apo) | 6TZD A (GSA) | 6TZD B (GSA) | 6U4I A (TSA) | 6U4I B (TSA) |
|---|---|---|---|---|---|---|
| **3VSY A (Apo)** | 0.67 | 0.58 | 0.59 | 0.43 | 0.52 | 0.70 |
| **3VSY B (Apo)** | 0.71 | 0.37 | 0.48 | 0.49 | 0.56 | 0.66 |
| **5KP4 B (GSA)** | 0.43 | 0.44 | 0.41 | 0.49 | 0.44 | 0.47 |
| **1OH0 A (TSA)** | 0.44 | 0.43 | 0.39 | 0.49 | 0.51 | 0.50 |
| **1OH0 B (TSA)** | 0.58 | 0.40 | 0.37 | 0.46 | 0.46 | 0.38 |

**Table S4.** RMSDs between crystal structures for different KSI catalytic states obtained at cryo (100 K) and room temperature (280 K). Cryo KSI Apo, GSA-bound and TSA-bound structures have been obtained from the PDB (PDB 3VSY, 5KP4 and 1OH0 for Apo, GSA-bound and TSA-bound, respectively), while the corresponding RT (280 K) structures were obtained in this study (PDB 6U1Z, 6TZD and 6U4I for Apo, GSA-bound and TSA-bound, respectively). KSI models were aligned on the backbone of residues 5–125 and RMSDs obtained for all residues.

| **280 K** **100K** | **6U1Z A (Apo)** | **6U1Z B (Apo)** | **6TZD A (GSA)** | **6TZD B (GSA)** | **6U4I A (TSA)** | **6U4I B (TSA)** |
|---|---|---|---|---|---|---|
| **3VSY A (Apo)** | 0.20 | 0.35 | 0.36 | 0.22 | 0.23 | 0.38 |
| **3VSY B (Apo)** | 0.36 | 0.21 | 0.24 | 0.30 | 0.36 | 0.26 |
| **5KP4 B (GSA)** | 0.22 | 0.28 | 0.27 | 0.20 | 0.20 | 0.29 |
| **1OH0 A (TSA)** | 0.34 | 0.18 | 0.17 | 0.26 | 0.32 | 0.17 |
| **1OH0 B (TSA)** | 0.31 | 0.19 | 0.19 | 0.24 | 0.28 | 0.21 |

**Table S5.** RMSDs between crystal structures for different KSI catalytic states obtained at cryo (100 K) and room temperature (280 K). Cryo KSI Apo, GSA-bound and TSA-bound structures have been obtained from the PDB (PDB 3VSY, 5KP4 and 1OH0 for Apo, GSA-bound and TSA-bound, respectively), while the corresponding RT (280 K) structures were obtained in this study (PDB 6U1Z, 6TZD and 6U4I for Apo, GSA-bound and TSA-bound, respectively). KSI models were aligned on the backbone of residues 5–125 and RMSDs obtained for all residues excluding loops 62–65 and 91–96.

| PDB | Mutations | Ligand | Resolution | Space group | # molecules AU |
|---|---|---|---|---|---|
| 1DMM | Y57F | - | 1.9 | $C222_1$ | 1 |
| 1E97 | Y16F/Y32F/Y57F | - | 2.0 | $C222_1$ | 1 |
| 1EA2 | Y16F | - | 1.8 | $C222_1$ | 1 |
| 1OPY | WT | - | 1.9 | $C222_1$ | 1 |
| 3RGR | M116A | - | 1.59 | $C222_1$ | 1 |
| 3SED | M105A | - | 1.3 | $C222_1$ | 1 |
| 4K1V | Y16F/Y57F | - | 1.8 | $C222_1$ | 1 |
| 3VSY | WT | - | 1.5 | $P2_12_12_1$ | 2 |
| 4K1U | Y16F/Y32F | - | 2.0 | $P2_12_12_1$ | 2 |
| 5D83 | D40N, Y32 (Cl-Y) | - | 1.7 | $P2_12_12_1$ | 2 |
| 5D82 | D40N, Y16 (Cl-Y) | - | 1.37 | $P2_12_12_1$ | 2 |
| 5D81 | D40N, Y57 (Cl-Y) | - | 1.39 | $C222_1$ | 1 |
| 3OX9 | D40N/C69S/C81S/C97S/ F86C-CN | - | 2.0 | $P2_1$ | 4 |
| 3OXA | D40N/C69S/C81S/C97S/ M116C-CN | - | 1.89 | $P2_1$ | 4 |
| 3T8N | Y16AD103A | - | 1.47 | $C2_1$ | 4 |

**Table S6.** Cryo KSI Apo crystal structures of high-resolution (≤ 2 Å) available from the PDB.

| PDB | Mutations | Ligand | Resolution | Space group | # molecules AU |
|---|---|---|---|---|---|
| 1OHO | Y16F/D40N | Equilenin | 1.9 | $C222_1$ | 1 |
| 2INX | D40N | 2,6-difluorophenol | 1.5 | $C222_1$ | 1 |
| 3CPO | D40N | 2-fluorophenol | 1.24 | $C222_1$ | 1 |
| 1CQS | D103E/D40N | Equilenin | 1.9 | $C2_1$ | 2 |
| 1OGX | D40N | Equilenin | 2.0 | $C2_1$ | 2 |
| 1OH0 | WT | Equilenin | 1.1 | $C2_1$ | 2 |
| 3FZW | D40N/D103N | Equilenin | 1.32 | $C2_1$ | 2 |
| 3OWS | D40N/C69S/C81S/C97S/ M116C-CN | Equilenin | 1.71 | $P2_1$ | 4 |
| 3OWU | D40N/C69S/C81S/C97S/ F86C-CN | Equilenin | 1.7 | $P2_1$ | 4 |
| 3VGN | D40N | 3-fluoro-4-nitrophenol | 1.3 | $P2_12_12_1$ | 2 |
| 2PZV | D40N | Phenol | 1.25 | P1 | 4 |
| 3IPT | Y16S/D40N | Equilenin | 1.63 | $P2_1$ | 4 |
| 5KP1 | D40N, Y16(Cl-Y) | Equilenin | 1.22 | P1 | 4 |
| 5KP3 | D40N, Y57(Cl-Y) | Equilenin | 1.7 | $P2_12_12_1$ | 2 |
| 5G2G | M116K | Equilenin | 1.6 | $P2_12_12_1$ | 2 |

**Table S7.** Cryo KSI TSA-bound crystal structures of high-resolution (≤ 2 Å) available from the PDB.

| Apo | | | TSA-bound | | |
|---|---|---|---|---|---|
| $Set_1^{omit}$ | $Set_2^{omit}$ | $Set_3^{omit}$ | $Set_1^{omit}$ | $Set_2^{omit}$ | $Set_3^{omit}$ |
| 1C7H | 1C7H | 1C7H | 1GS3 | 1GS3 | 1GS3 |
| 1DMM | 1DMM | 1DMM | 1OHO | 1OHO | 1OHO |
| 1DMN | 1DMN | 1DMN | 1W6Y | 1W6Y | 1W6Y |
| 1DMQ | 1DMQ | 1DMQ | 2INX | 2INX | 2INX |
| 1E97 | 1E97 | 1E97 | 3CPO | 3CPO | 3CPO |
| 1EA2 | 1EA2 | 1EA2 | 5AI1 | 5AI1 | 5AI1 |
| 1OPY | 1OPY | 1OPY | 1OH0_A | 1OH0_A | 1OH0_A |
| 1W02 | 1W02 | 1W02 | 1OH0_B | 1OH0_B | 1OH0_B |
| 3RGR | 3RGR | 3RGR | 1CQS_A | 1CQS_A | 1CQS_A |
| 3SED | 3SED | 3SED | 1OGX_A | 1OGX_A | 1OGX_A |
| 4K1V | 4K1V | 4K1V | 3FZW_A | 3FZW_A | 3FZW_A |
| 5D81 | 5D81 | 5D81 | 3VGN_A | 3VGN_A | 3VGN_A |
| 3VSY_A | 3VSY_A | 3VSY_A | 5KP3_A | 5KP3_A | 5KP3_A |
| 3VSY_B | 3VSY_B | 3VSY_B | 5G2G_A | 5G2G_A | 5G2G_A |
| 1K41_A | 1K41_A | 1K41_A | 1CQS_B | 1CQS_B | 1CQS_B |
| 1VZZ_A | 1VZZ_A | 1VZZ_A | 1OGX_B | 1OGX_B | 1OGX_B |
| 1W00_A | 1W00_A | 1W00_A | 3FZW_B | 3FZW_B | 3FZW_B |
| 1W01_A | 1W01_A | 1W01_A | 3VGN_B | 3VGN_B | 3VGN_B |
| 4K1U_A | 4K1U_A | 4K1U_A | 5KP3_B | 5KP3_B | 5KP3_B |
| 5D82_A | 5D82_A | 5D82_A | 5G2G_B | 5G2G_B | 5G2G_B |
| 5D83_A | 5D83_A | 5D83_A | 3OWS_A | 3OWS_A | 3OWS_A |
| 1K41_B | 1K41_B | 1K41_B | 3OWS_B | 3OWS_B | 3OWS_B |
| 1VZZ_B | 1VZZ_B | 1VZZ_B | 3OWS_C | 3OWS_C | 3OWS_C |
| 1W00_B | 1W00_B | 1W00_B | 3OWS_D | 3OWS_D | 3OWS_D |
| 1W01_B | 1W01_B | 1W01_B | 3OWU_A | 3OWU_A | 3OWU_A |
| 4K1U_B | 4K1U_B | 4K1U_B | 3OWU_B | 3OWU_B | 3OWU_B |
| 5D82_B | 5D82_B | 5D82_B | 3OWU_C | 3OWU_C | 3OWU_C |
| 5D83_B | 5D83_B | 5D83_B | 3OWU_D | 3OWU_D | 3OWU_D |
| 3OWY_F | 3OWY_F | 3OWY_F | 2PZV_A | 2PZV_A | 2PZV_A |
| 3OWY_G | 3OWY_G | 3OWY_G | 2PZV_B | 2PZV_B | 2PZV_B |
| 3OX9_A | 3OX9_A | 3OX9_A | 2PZV_C | 2PZV_C | 2PZV_C |
| 3OX9_B | 3OX9_B | 3OX9_B | 2PZV_D | 2PZV_D | 2PZV_D |
| 3OX9_C | 3OX9_C | 3OX9_C | 3IPT_A | 3IPT_A | 3IPT_A |
| 3OX9_D | 3OX9_D | 3OX9_D | 3IPT_B | 3IPT_B | 3IPT_B |
| 3OXA_A | 3OXA_A | 3OXA_A | 3IPT_C | 3IPT_C | 3IPT_C |
| 3OXA_B | 3OXA_B | 3OXA_B | 3IPT_D | 3IPT_D | 3IPT_D |
| 3OXA_C | 3OXA_C | 3OXA_C | 3OWY_A | 3OWY_A | 3OWY_A |
| 3OXA_D | 3OXA_D | 3OXA_D | 3OWY_B | 3OWY_B | 3OWY_B |
| 3T8N_A | 3T8N_A | 3T8N_A | 3OWY_C | 3OWY_C | 3OWY_C |
| 3T8N_B | 3T8N_B | 3T8N_B | 3OWY_D | 3OWY_D | 3OWY_D |
| 3T8N_D | 3T8N_D | 3T8N_D | 3OWY_E | 3OWY_E | 3OWY_E |
| 3T8N_F | 3T8N_F | 3T8N_F | 3OWY_H | 3OWY_H | 3OWY_H |
| | | | 5KP1_A | 5KP1_A | 5KP1_A |
| | | | 5KP1_B | 5KP1_B | 5KP1_B |
| | | | 5KP1_C | 5KP1_C | 5KP1_C |
| | | | 5KP1_D | 5KP1_D | 5KP1_D |

**Table S8.** KSI molecules used to obtain Apo and TSA-bound pseudo-ensembles in which ~30% of all molecules have been randomly omitted (light grey, 12 out of 42 and 14 out of 46 KSI molecules omitted, respectively); random selection and exclusion of molecules was repeated three times to generate three independent sets ($\mathrm{Set}_{1-3}^{\mathrm{omit}}$) for both Apo and TSA-bound.

| | Cα Apo MDev (Å) | | | | CαTSA MDev (Å) | | | |
|---|---|---|---|---|---|---|---|---|
| **Residues** | **Full** | $Set_1^{omit}$ | $Set_2^{omit}$ | $Set_3^{omit}$ | **Full** | $Set_1^{omit}$ | $Set_2^{omit}$ | $Set_3^{omit}$ |
| **5** | 0.284 | 0.267 | 0.292 | 0.287 | 0.262 | 0.285 | 0.263 | 0.261 |
| **6** | 0.242 | 0.231 | 0.248 | 0.237 | 0.263 | 0.278 | 0.266 | 0.272 |
| **7** | 0.273 | 0.264 | 0.288 | 0.272 | 0.294 | 0.318 | 0.307 | 0.315 |
| **8** | 0.220 | 0.216 | 0.225 | 0.223 | 0.228 | 0.253 | 0.238 | 0.248 |
| **9** | 0.165 | 0.165 | 0.174 | 0.168 | 0.166 | 0.172 | 0.171 | 0.164 |
| **10** | 0.190 | 0.191 | 0.189 | 0.196 | 0.202 | 0.203 | 0.207 | 0.208 |
| **11** | 0.222 | 0.218 | 0.224 | 0.226 | 0.193 | 0.202 | 0.199 | 0.196 |
| **12** | 0.168 | 0.170 | 0.177 | 0.172 | 0.170 | 0.191 | 0.175 | 0.168 |
| **13** | 0.178 | 0.185 | 0.183 | 0.184 | 0.170 | 0.174 | 0.174 | 0.171 |
| **14** | 0.193 | 0.202 | 0.186 | 0.192 | 0.184 | 0.181 | 0.190 | 0.200 |
| **15** | 0.212 | 0.226 | 0.210 | 0.219 | 0.193 | 0.195 | 0.214 | 0.193 |
| **16** | 0.200 | 0.205 | 0.204 | 0.205 | 0.185 | 0.204 | 0.192 | 0.193 |
| **17** | 0.216 | 0.221 | 0.201 | 0.219 | 0.182 | 0.186 | 0.190 | 0.187 |
| **18** | 0.248 | 0.222 | 0.209 | 0.224 | 0.175 | 0.180 | 0.183 | 0.167 |
| **19** | 0.259 | 0.232 | 0.237 | 0.235 | 0.225 | 0.239 | 0.230 | 0.228 |
| **20** | 0.306 | 0.273 | 0.284 | 0.271 | 0.256 | 0.261 | 0.256 | 0.263 |
| **21** | 0.304 | 0.264 | 0.267 | 0.259 | 0.238 | 0.248 | 0.237 | 0.239 |
| **22** | 0.309 | 0.274 | 0.255 | 0.274 | 0.206 | 0.205 | 0.198 | 0.215 |
| **23** | 0.346 | 0.313 | 0.290 | 0.317 | 0.248 | 0.246 | 0.245 | 0.254 |
| **24** | 0.377 | 0.351 | 0.359 | 0.369 | 0.305 | 0.296 | 0.324 | 0.305 |
| **25** | 0.422 | 0.372 | 0.369 | 0.385 | 0.366 | 0.365 | 0.372 | 0.366 |
| **26** | 0.448 | 0.420 | 0.436 | 0.446 | 0.409 | 0.422 | 0.413 | 0.410 |
| **27** | 0.318 | 0.291 | 0.324 | 0.318 | 0.344 | 0.349 | 0.347 | 0.344 |
| **28** | 0.241 | 0.219 | 0.224 | 0.225 | 0.264 | 0.255 | 0.264 | 0.283 |
| **29** | 0.242 | 0.231 | 0.230 | 0.238 | 0.291 | 0.301 | 0.278 | 0.314 |
| **30** | 0.258 | 0.240 | 0.265 | 0.261 | 0.322 | 0.323 | 0.320 | 0.338 |
| **31** | 0.194 | 0.193 | 0.195 | 0.194 | 0.204 | 0.208 | 0.204 | 0.220 |
| **32** | 0.183 | 0.182 | 0.187 | 0.184 | 0.186 | 0.184 | 0.185 | 0.192 |
| **33** | 0.216 | 0.220 | 0.208 | 0.210 | 0.153 | 0.147 | 0.159 | 0.153 |
| **34** | 0.533 | 0.541 | 0.536 | 0.487 | 0.391 | 0.314 | 0.372 | 0.338 |
| **35** | 0.572 | 0.565 | 0.536 | 0.516 | 0.401 | 0.333 | 0.411 | 0.344 |
| **36** | 0.210 | 0.217 | 0.199 | 0.196 | 0.155 | 0.142 | 0.153 | 0.147 |
| **37** | 0.240 | 0.248 | 0.232 | 0.241 | 0.273 | 0.276 | 0.297 | 0.231 |
| **38** | 0.258 | 0.254 | 0.246 | 0.251 | 0.307 | 0.313 | 0.342 | 0.301 |
| **39** | 0.283 | 0.275 | 0.277 | 0.282 | 0.304 | 0.313 | 0.308 | 0.316 |
| **40** | 0.296 | 0.290 | 0.300 | 0.298 | 0.327 | 0.340 | 0.318 | 0.348 |
| **41** | 0.358 | 0.357 | 0.376 | 0.375 | 0.387 | 0.398 | 0.390 | 0.421 |
| **42** | 0.376 | 0.376 | 0.403 | 0.376 | 0.386 | 0.394 | 0.387 | 0.421 |
| **43** | 0.492 | 0.506 | 0.487 | 0.524 | 0.414 | 0.408 | 0.402 | 0.450 |
| **44** | 0.471 | 0.473 | 0.484 | 0.489 | 0.440 | 0.450 | 0.420 | 0.489 |
| **45** | 0.475 | 0.481 | 0.465 | 0.488 | 0.431 | 0.457 | 0.412 | 0.481 |
| **46** | 0.358 | 0.337 | 0.316 | 0.338 | 0.353 | 0.355 | 0.334 | 0.378 |
| **47** | 0.337 | 0.332 | 0.303 | 0.333 | 0.288 | 0.308 | 0.287 | 0.305 |
| **48** | 0.317 | 0.294 | 0.297 | 0.302 | 0.212 | 0.204 | 0.218 | 0.203 |
| **49** | 0.602 | 0.546 | 0.540 | 0.511 | 0.363 | 0.278 | 0.284 | 0.277 |
| **50** | 0.442 | 0.384 | 0.386 | 0.372 | 0.334 | 0.293 | 0.299 | 0.317 |
| **51** | 0.452 | 0.399 | 0.414 | 0.378 | 0.367 | 0.353 | 0.344 | 0.360 |
| **52** | 0.328 | 0.310 | 0.320 | 0.293 | 0.312 | 0.314 | 0.301 | 0.346 |
| **53** | 0.287 | 0.263 | 0.249 | 0.263 | 0.270 | 0.268 | 0.280 | 0.314 |
| **54** | 0.318 | 0.304 | 0.299 | 0.307 | 0.332 | 0.340 | 0.338 | 0.367 |
| **55** | 0.352 | 0.352 | 0.355 | 0.362 | 0.390 | 0.406 | 0.395 | 0.411 |
| **56** | 0.342 | 0.325 | 0.308 | 0.340 | 0.375 | 0.384 | 0.385 | 0.380 |
| **57** | 0.392 | 0.339 | 0.317 | 0.349 | 0.318 | 0.323 | 0.323 | 0.333 |
| **58** | 0.514 | 0.536 | 0.446 | 0.530 | 0.399 | 0.408 | 0.397 | 0.419 |
| **59** | 0.539 | 0.557 | 0.490 | 0.576 | 0.436 | 0.452 | 0.439 | 0.466 |
| **60** | 0.513 | 0.491 | 0.448 | 0.504 | 0.493 | 0.529 | 0.525 | 0.551 |
| **61** | 0.492 | 0.524 | 0.432 | 0.535 | 0.335 | 0.352 | 0.363 | 0.361 |

| | | | | | | | | |
|---|---|---|---|---|---|---|---|---|
| **62** | 1.141 | 1.237 | 1.095 | 1.205 | 0.605 | 0.624 | 0.649 | 0.540 |
| **63** | 1.684 | 1.561 | 1.786 | 1.802 | 1.271 | 1.293 | 1.264 | 1.128 |
| **64** | 2.697 | 2.591 | 2.559 | 2.641 | 2.264 | 2.209 | 2.124 | 1.993 |
| **65** | 0.835 | 0.888 | 0.767 | 0.860 | 0.636 | 0.691 | 0.610 | 0.672 |
| **66** | 0.369 | 0.394 | 0.365 | 0.400 | 0.392 | 0.404 | 0.404 | 0.408 |
| **67** | 0.319 | 0.314 | 0.318 | 0.337 | 0.367 | 0.393 | 0.391 | 0.392 |
| **68** | 0.244 | 0.255 | 0.232 | 0.258 | 0.203 | 0.197 | 0.202 | 0.202 |
| **69** | 0.279 | 0.282 | 0.268 | 0.273 | 0.286 | 0.290 | 0.291 | 0.296 |
| **70** | 0.269 | 0.261 | 0.268 | 0.269 | 0.232 | 0.229 | 0.234 | 0.232 |
| **71** | 0.333 | 0.304 | 0.348 | 0.321 | 0.304 | 0.329 | 0.301 | 0.319 |
| **72** | 0.273 | 0.259 | 0.273 | 0.269 | 0.207 | 0.208 | 0.223 | 0.215 |
| **73** | 0.239 | 0.255 | 0.231 | 0.250 | 0.221 | 0.222 | 0.221 | 0.225 |
| **74** | 0.139 | 0.145 | 0.134 | 0.138 | 0.142 | 0.136 | 0.142 | 0.149 |
| **75** | 0.153 | 0.152 | 0.152 | 0.154 | 0.175 | 0.182 | 0.174 | 0.176 |
| **76** | 0.172 | 0.166 | 0.154 | 0.168 | 0.184 | 0.188 | 0.188 | 0.203 |
| **77** | 0.280 | 0.281 | 0.273 | 0.284 | 0.248 | 0.233 | 0.230 | 0.264 |
| **78** | 0.265 | 0.243 | 0.250 | 0.256 | 0.247 | 0.252 | 0.261 | 0.275 |
| **79** | 0.265 | 0.234 | 0.270 | 0.247 | 0.261 | 0.270 | 0.275 | 0.290 |
| **80** | 0.248 | 0.243 | 0.253 | 0.265 | 0.198 | 0.204 | 0.196 | 0.194 |
| **81** | 0.218 | 0.216 | 0.219 | 0.223 | 0.169 | 0.168 | 0.160 | 0.154 |
| **82** | 0.179 | 0.168 | 0.174 | 0.170 | 0.190 | 0.190 | 0.182 | 0.184 |
| **83** | 0.136 | 0.133 | 0.134 | 0.130 | 0.151 | 0.164 | 0.145 | 0.151 |
| **84** | 0.119 | 0.123 | 0.113 | 0.127 | 0.117 | 0.123 | 0.112 | 0.130 |
| **85** | 0.182 | 0.177 | 0.190 | 0.186 | 0.191 | 0.205 | 0.194 | 0.204 |
| **86** | 0.213 | 0.201 | 0.214 | 0.209 | 0.174 | 0.178 | 0.171 | 0.176 |
| **87** | 0.268 | 0.259 | 0.276 | 0.270 | 0.249 | 0.264 | 0.255 | 0.261 |
| **88** | 0.263 | 0.265 | 0.259 | 0.268 | 0.203 | 0.198 | 0.206 | 0.194 |
| **89** | 0.312 | 0.299 | 0.279 | 0.307 | 0.245 | 0.246 | 0.247 | 0.232 |
| **90** | 0.452 | 0.465 | 0.393 | 0.444 | 0.260 | 0.252 | 0.262 | 0.260 |
| **91** | 0.996 | 1.018 | 0.954 | 0.990 | 0.541 | 0.571 | 0.547 | 0.561 |
| **92** | 1.457 | 1.470 | 1.458 | 1.448 | 1.287 | 1.338 | 1.297 | 1.342 |
| **93** | 2.288 | 2.237 | 2.434 | 2.205 | 2.318 | 2.483 | 2.300 | 2.367 |
| **94** | 1.870 | 1.831 | 1.999 | 1.869 | 1.268 | 1.389 | 1.281 | 1.324 |
| **95** | 0.929 | 0.907 | 0.951 | 0.996 | 0.707 | 0.676 | 0.756 | 0.662 |
| **96** | 0.623 | 0.626 | 0.624 | 0.639 | 0.467 | 0.458 | 0.478 | 0.407 |
| **97** | 0.374 | 0.343 | 0.350 | 0.345 | 0.403 | 0.397 | 0.397 | 0.401 |
| **98** | 0.392 | 0.368 | 0.392 | 0.384 | 0.387 | 0.388 | 0.387 | 0.394 |
| **99** | 0.288 | 0.289 | 0.299 | 0.284 | 0.319 | 0.322 | 0.316 | 0.342 |
| **100** | 0.216 | 0.211 | 0.227 | 0.219 | 0.214 | 0.222 | 0.216 | 0.220 |
| **101** | 0.210 | 0.200 | 0.209 | 0.207 | 0.207 | 0.214 | 0.209 | 0.217 |
| **102** | 0.189 | 0.188 | 0.185 | 0.193 | 0.172 | 0.174 | 0.171 | 0.176 |
| **103** | 0.204 | 0.198 | 0.202 | 0.197 | 0.205 | 0.214 | 0.205 | 0.222 |
| **104** | 0.172 | 0.171 | 0.183 | 0.177 | 0.181 | 0.177 | 0.177 | 0.187 |
| **105** | 0.150 | 0.159 | 0.155 | 0.160 | 0.139 | 0.142 | 0.140 | 0.142 |
| **106** | 0.155 | 0.154 | 0.160 | 0.158 | 0.150 | 0.156 | 0.159 | 0.145 |
| **107** | 0.163 | 0.167 | 0.167 | 0.164 | 0.147 | 0.149 | 0.147 | 0.151 |
| **108** | 0.206 | 0.216 | 0.218 | 0.213 | 0.194 | 0.187 | 0.197 | 0.207 |
| **109** | 0.333 | 0.355 | 0.350 | 0.349 | 0.275 | 0.275 | 0.270 | 0.304 |
| **110** | 0.329 | 0.327 | 0.320 | 0.319 | 0.262 | 0.279 | 0.260 | 0.284 |
| **111** | 0.254 | 0.245 | 0.248 | 0.249 | 0.227 | 0.240 | 0.235 | 0.246 |
| **112** | 0.175 | 0.188 | 0.163 | 0.182 | 0.143 | 0.141 | 0.144 | 0.144 |
| **113** | 0.157 | 0.164 | 0.168 | 0.169 | 0.130 | 0.126 | 0.131 | 0.129 |
| **114** | 0.165 | 0.174 | 0.165 | 0.170 | 0.135 | 0.136 | 0.144 | 0.133 |
| **115** | 0.230 | 0.243 | 0.226 | 0.217 | 0.200 | 0.194 | 0.221 | 0.205 |
| **116** | 0.329 | 0.333 | 0.343 | 0.322 | 0.325 | 0.324 | 0.340 | 0.327 |
| **117** | 0.295 | 0.294 | 0.316 | 0.304 | 0.332 | 0.342 | 0.345 | 0.351 |
| **118** | 0.331 | 0.315 | 0.335 | 0.322 | 0.328 | 0.341 | 0.340 | 0.351 |
| **119** | 0.281 | 0.273 | 0.289 | 0.281 | 0.282 | 0.289 | 0.279 | 0.304 |
| **120** | 0.313 | 0.304 | 0.320 | 0.312 | 0.353 | 0.360 | 0.351 | 0.384 |

| | | | | | | | | |
|---|---|---|---|---|---|---|---|---|
| **121** | 0.434 | 0.430 | 0.455 | 0.436 | 0.480 | 0.487 | 0.481 | 0.519 |
| **122** | 0.434 | 0.423 | 0.476 | 0.436 | 0.463 | 0.466 | 0.465 | 0.491 |
| **123** | 0.377 | 0.368 | 0.380 | 0.378 | 0.299 | 0.280 | 0.304 | 0.287 |
| **124** | 0.292 | 0.277 | 0.289 | 0.289 | 0.273 | 0.269 | 0.277 | 0.291 |
| **125** | 0.329 | 0.317 | 0.324 | 0.311 | 0.345 | 0.333 | 0.342 | 0.371 |

**Table S9.** Cα MDevs for Apo and TSA-bound pseudo-ensembles composed of all structures (42 and 46 KSI molecules, respectively, indicated as 'Full') and pseudo-ensembles from which 30% of the structures have been randomly omitted (12 out of 42 and 14 out of 46 KSI molecules omitted, respectively; random selection and exclusion of molecules was repeated three times to generate three independent sets ($Set_{1-3}^{\mathrm{omit}}$) for both Apo and TSA-bound).

| | | **Cα ΣMDev entire enzyme (Å)** | **Cα ΣMDev enzyme core (Å)** |
|---|---|---|---|
| **Apo** | **Full** | 47.068 | 32.547 |
| | $\boldsymbol{Set_1^{omit}}$ | 46.087 | 31.722 |
| | $\boldsymbol{Set_2^{omit}}$ | 46.179 | 31.555 |
| | $\boldsymbol{Set_3^{omit}}$ | 46.647 | 31.992 |
| **TSA-bound** | **Full** | 41.225 | 29.860 |
| | $\boldsymbol{Set_1^{omit}}$ | 41.856 | 30.124 |
| | $\boldsymbol{Set_2^{omit}}$ | 41.346 | 30.040 |
| | $\boldsymbol{Set_3^{omit}}$ | 41.847 | 30.850 |

**Table S10.** Sum of Cα MDev values (ΣMDev, from Table S9) for Apo and TSA-bound pseudo-ensembles composed of all structures (42 and 46 KSI molecules, respectively, indicated as 'Full') and pseudo-ensembles from which 30% of the structures have been randomly omitted (12 out of 42 and 14 out of 46 KSI molecules omitted, respectively; indicated as '$\mathrm{Set}_{1-3}^{\mathrm{omit}}$').

| | | Cα TSA-bound ΣMDev (Å) | | | |
|---|---|---|---|---|---|
| | | **Full** | $Set_1^{omit}$ | $Set_2^{omit}$ | $Set_3^{omit}$ |
| **Cα Apo ΣMDev (Å)** | **Full** | 5.843 | 5.213 | 5.723 | 5.221 |
| | $Set_1^{omit}$ | 4.862 | 4.231 | 4.741 | 4.240 |
| | $Set_2^{omit}$ | 4.954 | 4.324 | 4.834 | 4.333 |
| | $Set_3^{omit}$ | 5.422 | 4.791 | 5.301 | 4.800 |

**Table S11.** The values represent the differences (ΔMDev) between the sums of Cα MDev values for Apo (Cα Apo ΣMDev from Table S10) and the sums of Cα MDev values for TSA-bound pseudo-ensembles (Cα TSA-bound ΣMDev from Table S10) for the entire enzyme. The 16 values represent all combinations between the Cα Apo ΣMDev from Apo Full and Sets1-3 and TSA-bound Full and Sets1-3.

| | | Cα TSA-bound ΣMDev (Å) | | | |
|---|---|---|---|---|---|
| | | **Full** | $Set_1^{omit}$ | $Set_2^{omit}$ | $Set_3^{omit}$ |
| **Cα Apo ΣMDev (Å)** | **Full** | 2.687 | 2.423 | 2.507 | 1.697 |
| | $Set_1^{omit}$ | 1.862 | 1.598 | 1.682 | 0.872 |
| | $Set_2^{omit}$ | 1.695 | 1.431 | 1.515 | 0.704 |
| | $Set_3^{omit}$ | 2.132 | 1.868 | 1.952 | 1.142 |

**Table S11.** The values represent the differences (ΔMDev) between the sums of Cα MDev values for Apo (Cα Apo ΣMDev from Table S10) and the sums of Cα MDev values for TSA-bound pseudo-ensembles (Cα TSA-bound ΣMDev from Table S10) for the enzyme core. The 16 values represent all combinations between the Cα Apo ΣMDev from Apo Full and Sets1-3 and TSA-bound Full and Sets1-3.

| Cα Apo ΣMDev (Å) | | Cα TSA-bound ΣMDev (Å) | | $\Delta MDev_{Apo\text{-}TSA}$ | $\Delta MDev_{Apo\text{-}TSA}$ / $\Sigma MDev_{Apo}$ |
|---|---|---|---|---|---|
| **Full** | 47.068 | **Full** | 41.225 | 5.843 | 0.124 |
| | | $Set_1^{omit}$ | 41.856 | 5.213 | 0.111 |
| | | $Set_2^{omit}$ | 41.346 | 5.723 | 0.122 |
| | | $Set_3^{omit}$ | 41.847 | 5.221 | 0.111 |
| $Set_1^{omit}$ | 46.087 | **Full** | 41.225 | 4.862 | 0.105 |
| | | $Set_1^{omit}$ | 41.856 | 4.231 | 0.092 |
| | | $Set_2^{omit}$ | 41.346 | 4.741 | 0.103 |
| | | $Set_3^{omit}$ | 41.847 | 4.240 | 0.092 |
| $Set_2^{omit}$ | 46.179 | **Full** | 41.225 | 4.954 | 0.107 |
| | | $Set_1^{omit}$ | 41.856 | 4.324 | 0.094 |
| | | $Set_2^{omit}$ | 41.346 | 4.834 | 0.105 |
| | | $Set_3^{omit}$ | 41.847 | 4.333 | 0.094 |
| $Set_3^{omit}$ | 46.647 | **Full** | 41.225 | 5.422 | 0.116 |
| | | $Set_1^{omit}$ | 41.856 | 4.791 | 0.103 |
| | | $Set_2^{omit}$ | 41.346 | 5.301 | 0.114 |
| | | $Set_3^{omit}$ | 41.847 | 4.800 | 0.103 |

**Table S13.** The values in the far right column represent the conformational heterogeneity dampening in entire Apo enzyme upon TSA binding as obtained by dividing the difference between Cα Apo and TSA-bound MDevs ($\Delta MDev_{Apo\text{-}TSA}$) by the sum of Cα Apo MDevs ($\Sigma MDev_{Apo}$). The values in column 2, 4 and 5 are taken from Tables S10 and S11.

| Cα Apo ΣMDev (Å) | | Cα TSA-bound ΣMDev (Å) | | $\Delta MDev_{Apo\text{-}TSA}$ | $\Delta MDev_{Apo\text{-}TSA}$ / $\Sigma MDev_{Apo}$ |
|---|---|---|---|---|---|
| **Full** | 32.547 | **Full** | 29.860 | 2.687 | 0.083 |
| | | $\boldsymbol{Set_1^{omit}}$ | 30.124 | 2.423 | 0.074 |
| | | $\boldsymbol{Set_2^{omit}}$ | 30.040 | 2.507 | 0.077 |
| | | $\boldsymbol{Set_3^{omit}}$ | 30.850 | 1.697 | 0.052 |
| $\boldsymbol{Set_1^{omit}}$ | 31.722 | **Full** | 29.860 | 1.862 | 0.059 |
| | | $\boldsymbol{Set_1^{omit}}$ | 30.124 | 1.598 | 0.050 |
| | | $\boldsymbol{Set_2^{omit}}$ | 30.040 | 1.682 | 0.053 |
| | | $\boldsymbol{Set_3^{omit}}$ | 30.850 | 0.872 | 0.027 |
| $\boldsymbol{Set_2^{omit}}$ | 31.555 | **Full** | 29.860 | 1.695 | 0.054 |
| | | $\boldsymbol{Set_1^{omit}}$ | 30.124 | 1.431 | 0.045 |
| | | $\boldsymbol{Set_2^{omit}}$ | 30.040 | 1.515 | 0.048 |
| | | $\boldsymbol{Set_3^{omit}}$ | 30.850 | 0.704 | 0.022 |
| $\boldsymbol{Set_3^{omit}}$ | 31.992 | **Full** | 29.860 | 2.132 | 0.067 |
| | | $\boldsymbol{Set_1^{omit}}$ | 30.124 | 1.868 | 0.058 |
| | | $\boldsymbol{Set_2^{omit}}$ | 30.040 | 1.952 | 0.061 |
| | | $\boldsymbol{Set_3^{omit}}$ | 30.850 | 1.142 | 0.036 |

**Table S14.** The values in the far right column represent the conformational heterogeneity dampening in Apo enzyme core (excluding 62-65 and 91-96 loops) upon TSA binding as obtained by dividing the difference between Cα Apo and TSA-bound MDevs ($\Delta MDev_{Apo\text{-}TSA}$) by the sum of Cα Apo MDevs ($\Sigma MDev_{Apo}$). The values in column 2, 4 and 5 are taken from Tables S10 and S12.

| Residue | Apo Cα MDev (Å) | TSA Cα MDev (Å) |
|---|---|---|
| **5** | 0.159 | 0.138 |
| **6** | 0.140 | 0.143 |
| **7** | 0.179 | 0.183 |
| **8** | 0.126 | 0.141 |
| **9** | 0.106 | 0.115 |
| **10** | 0.124 | 0.152 |
| **11** | 0.154 | 0.147 |
| **12** | 0.121 | 0.126 |
| **13** | 0.160 | 0.163 |
| **14** | 0.184 | 0.178 |
| **15** | 0.186 | 0.175 |
| **16** | 0.167 | 0.135 |
| **17** | 0.179 | 0.160 |
| **18** | 0.215 | 0.202 |
| **19** | 0.173 | 0.154 |
| **20** | 0.224 | 0.186 |
| **21** | 0.238 | 0.207 |
| **22** | 0.286 | 0.265 |
| **23** | 0.288 | 0.241 |
| **24** | 0.272 | 0.224 |
| **25** | 0.329 | 0.254 |
| **26** | 0.332 | 0.313 |
| **27** | 0.216 | 0.234 |
| **28** | 0.197 | 0.169 |
| **29** | 0.193 | 0.203 |
| **30** | 0.222 | 0.245 |
| **31** | 0.160 | 0.151 |
| **32** | 0.170 | 0.175 |
| **33** | 0.208 | 0.153 |
| **34** | 0.524 | 0.381 |
| **35** | 0.581 | 0.392 |
| **36** | 0.192 | 0.145 |
| **37** | 0.193 | 0.236 |
| **38** | 0.185 | 0.252 |
| **39** | 0.164 | 0.184 |
| **40** | 0.172 | 0.207 |
| **41** | 0.199 | 0.227 |
| **42** | 0.216 | 0.241 |
| **43** | 0.325 | 0.267 |
| **44** | 0.310 | 0.279 |
| **45** | 0.363 | 0.307 |
| **46** | 0.282 | 0.298 |
| **47** | 0.282 | 0.226 |
| **48** | 0.260 | 0.216 |
| **49** | 0.596 | 0.345 |
| **50** | 0.435 | 0.284 |
| **51** | 0.425 | 0.296 |

| | | |
|---|---|---|
| **52** | 0.260 | 0.244 |
| **53** | 0.235 | 0.242 |
| **54** | 0.243 | 0.239 |
| **55** | 0.272 | 0.291 |
| **56** | 0.316 | 0.324 |
| **57** | 0.358 | 0.273 |
| **58** | 0.471 | 0.323 |
| **59** | 0.459 | 0.328 |
| **60** | 0.518 | 0.518 |
| **61** | 0.522 | 0.392 |
| **62** | 1.334 | 0.774 |
| **63** | 1.914 | 1.472 |
| **64** | 2.908 | 2.542 |
| **65** | 1.019 | 0.877 |
| **66** | 0.460 | 0.511 |
| **67** | 0.346 | 0.371 |
| **68** | 0.260 | 0.232 |
| **69** | 0.233 | 0.209 |
| **70** | 0.198 | 0.145 |
| **71** | 0.228 | 0.242 |
| **72** | 0.202 | 0.184 |
| **73** | 0.198 | 0.215 |
| **74** | 0.113 | 0.124 |
| **75** | 0.114 | 0.131 |
| **76** | 0.131 | 0.148 |
| **77** | 0.195 | 0.202 |
| **78** | 0.182 | 0.186 |
| **79** | 0.193 | 0.204 |
| **80** | 0.207 | 0.183 |
| **81** | 0.183 | 0.142 |
| **82** | 0.145 | 0.159 |
| **83** | 0.104 | 0.116 |
| **84** | 0.105 | 0.090 |
| **85** | 0.139 | 0.182 |
| **86** | 0.134 | 0.144 |
| **87** | 0.192 | 0.211 |
| **88** | 0.286 | 0.301 |
| **89** | 0.345 | 0.238 |
| **90** | 0.607 | 0.434 |
| **91** | 1.299 | 0.835 |
| **92** | 1.796 | 1.540 |
| **93** | 2.679 | 2.615 |
| **94** | 2.293 | 1.660 |
| **95** | 1.301 | 0.953 |
| **96** | 0.816 | 0.599 |
| **97** | 0.502 | 0.519 |
| **98** | 0.402 | 0.434 |
| **99** | 0.243 | 0.345 |
| **100** | 0.190 | 0.269 |

| | | |
|---|---|---|
| **101** | 0.136 | 0.168 |
| **102** | 0.127 | 0.116 |
| **103** | 0.146 | 0.147 |
| **104** | 0.110 | 0.124 |
| **105** | 0.134 | 0.123 |
| **106** | 0.129 | 0.132 |
| **107** | 0.145 | 0.148 |
| **108** | 0.190 | 0.201 |
| **109** | 0.331 | 0.293 |
| **110** | 0.320 | 0.252 |
| **111** | 0.227 | 0.250 |
| **112** | 0.162 | 0.163 |
| **113** | 0.155 | 0.139 |
| **114** | 0.145 | 0.118 |
| **115** | 0.205 | 0.203 |
| **116** | 0.281 | 0.289 |
| **117** | 0.202 | 0.246 |
| **118** | 0.216 | 0.227 |
| **119** | 0.152 | 0.150 |
| **120** | 0.157 | 0.172 |
| **121** | 0.217 | 0.257 |
| **122** | 0.287 | 0.362 |
| **123** | 0.326 | 0.306 |
| **124** | 0.239 | 0.297 |
| **125** | 0.352 | 0.410 |

**Table S15.** Cα MDevs for Apo and TSA-bound pseudo-ensembles composed of all structures (42 and 46 KSI molecules, respectively) and aligned using and alternative alignment procedure (see Materials and Methods).

| | Cα ΣMDev entire enzyme (Å) | Cα ΣMDev enzyme core (Å) |
|---|---|---|
| **Apo** | 44.36 | 27.00 |
| **TSA** | 39.32 | 25.45 |
| **$\Delta MDev_{Apo-TSA}$** | 5.04 | 1.55 |

**Table S16.** Sum of Cα MDev values (ΣMDev; from Table S15) for Apo and TSA-bound pseudo-ensembles composed of all structures (42 and 46 KSI molecules, respectively) aligned using an alternative alignment procedure (see Materials and Methods). $\Delta MDev_{Apo-TSA}$ indicates the difference between ΣMDev for Apo and ΣMDev for TSA-bound for either the entire enzyme or for the enzyme core (excluding loops 62-65 and 91-96).

| | $\Delta MDev_{Apo\text{-}TSA}$ / $\Sigma MDev_{Apo}$ |
|---|---|
| **Entire enzyme** | 0.11 |
| **Enzyme core** | 0.06 |

**Table S17.** The values represent the conformational heterogeneity dampening in Apo enzyme core upon TSA binding as obtained by dividing the difference between Cα Apo and TSA-bound MDevs ($\Delta MDev_{Apo\text{-}TSA}$, from Table S16) by the sum of Cα Apo MDevs ($\Sigma MDev_{Apo}$, from Table S16).

| Residue | Apo Cβ MDev (Å) | TSA Cβ MDev (Å) |
|---|---|---|
| **5** | 0.318 | 0.290 |
| **6** | 0.247 | 0.267 |
| **7** | 0.355 | 0.370 |
| **8** | 0.244 | 0.252 |
| **9** | 0.182 | 0.178 |
| **10** | 0.222 | 0.220 |
| **11** | - | - |
| **12** | 0.160 | 0.189 |
| **13** | 0.176 | 0.232 |
| **14** | 0.226 | 0.217 |
| **15** | 0.248 | 0.248 |
| **16** | 0.216 | 0.217 |
| **17** | 0.246 | 0.204 |
| **18** | 0.343 | 0.247 |
| **19** | 0.266 | 0.230 |
| **20** | 0.315 | 0.260 |
| **21** | 0.319 | 0.283 |
| **22** | 0.342 | 0.235 |
| **23** | - | - |
| **24** | 0.386 | 0.339 |
| **25** | 0.472 | 0.390 |
| **26** | 0.567 | 0.550 |
| **27** | 0.371 | 0.372 |
| **28** | 0.294 | 0.264 |
| **29** | 0.293 | 0.350 |
| **30** | 0.346 | 0.387 |
| **31** | 0.220 | 0.217 |
| **32** | 0.207 | 0.232 |
| **33** | 0.290 | 0.206 |
| **34** | 0.729 | 0.497 |
| **35** | 0.755 | 0.506 |
| **36** | 0.181 | 0.162 |
| **37** | 0.348 | 0.357 |
| **38** | 0.269 | 0.347 |
| **39** | 0.325 | 0.313 |
| **40** | 0.281 | 0.374 |
| **41** | 0.382 | 0.426 |
| **42** | 0.383 | 0.397 |
| **43** | - | - |
| **44** | 0.569 | 0.482 |
| **45** | 0.563 | 0.498 |
| **46** | 0.468 | 0.396 |
| **47** | 0.434 | 0.380 |
| **48** | 0.471 | 0.308 |
| **49** | - | - |
| **50** | 0.524 | 0.397 |
| **51** | 0.589 | 0.453 |

| | | |
|---|---|---|
| **52** | 0.347 | 0.346 |
| **53** | 0.305 | 0.262 |
| **54** | 0.341 | 0.363 |
| **55** | 0.383 | 0.427 |
| **56** | 0.353 | 0.407 |
| **57** | 0.402 | 0.296 |
| **58** | 0.516 | 0.440 |
| **59** | 0.671 | 0.584 |
| **60** | - | - |
| **61** | 0.517 | 0.326 |
| **62** | - | - |
| **63** | - | - |
| **64** | - | - |
| **65** | 1.219 | 0.975 |
| **66** | 0.464 | 0.439 |
| **67** | 0.438 | 0.476 |
| **68** | 0.272 | 0.231 |
| **69** | 0.450 | 0.499 |
| **70** | 0.347 | 0.248 |
| **71** | 0.359 | 0.339 |
| **72** | - | - |
| **73** | 0.307 | 0.285 |
| **74** | 0.174 | 0.150 |
| **75** | 0.172 | 0.194 |
| **76** | 0.192 | 0.215 |
| **77** | 0.377 | 0.353 |
| **78** | 0.291 | 0.269 |
| **79** | 0.323 | 0.328 |
| **80** | - | - |
| **81** | 0.300 | 0.237 |
| **82** | - | - |
| **83** | 0.206 | 0.204 |
| **84** | 0.138 | 0.131 |
| **85** | 0.246 | 0.264 |
| **86** | 0.203 | 0.170 |
| **87** | 0.323 | 0.307 |
| **88** | 0.363 | 0.270 |
| **89** | 0.421 | 0.379 |
| **90** | 0.644 | 0.374 |
| **91** | 1.311 | 0.776 |
| **92** | 1.767 | 1.398 |
| **93** | 2.608 | 2.735 |
| **94** | - | - |
| **95** | 1.040 | 1.083 |
| **96** | 0.778 | 0.661 |
| **97** | 0.522 | 0.534 |
| **98** | 0.442 | 0.404 |
| **99** | 0.317 | 0.327 |
| **100** | 0.200 | 0.190 |

| | | |
|---|---|---|
| **101** | 0.251 | 0.240 |
| **102** | 0.209 | 0.182 |
| **103** | 0.228 | 0.246 |
| **104** | 0.221 | 0.225 |
| **105** | 0.186 | 0.195 |
| **106** | 0.191 | 0.203 |
| **107** | 0.175 | 0.152 |
| **108** | 0.237 | 0.211 |
| **109** | 0.431 | 0.373 |
| **110** | 0.386 | 0.287 |
| **111** | - | - |
| **112** | 0.249 | 0.180 |
| **113** | 0.174 | 0.144 |
| **114** | 0.192 | 0.174 |
| **115** | 0.355 | 0.294 |
| **116** | 0.422 | 0.441 |
| **117** | 0.338 | 0.346 |
| **118** | 0.347 | 0.354 |
| **119** | 0.267 | 0.258 |
| **120** | 0.314 | 0.343 |
| **121** | 0.463 | 0.499 |
| **122** | 0.504 | 0.539 |
| **123** | 0.439 | 0.284 |
| **124** | 0.288 | 0.284 |
| **125** | 0.428 | 0.390 |

**Table S18.** Cβ MDevs of residues 5-125 for Apo and TSA-bound pseudo-ensembles composed of all structures (42 and 46 KSI molecules, respectively). Glycine residues lack Cβ and therefore no MDevs could be obtained (indicated with “-“).

| | Cβ ΣMDev entire enzyme (Å) | Cβ ΣMDev enzyme core (Å) |
|---|---|---|
| **Apo** | 43.481 | 34.759 |
| **TSA** | 39.478 | 31.851 |
| **$\Delta MDev_{Apo-TSA}$** | 4.003 | 2.908 |

**Table S19.** Sum of Cβ MDev values (ΣMDev; from Table S18) for Apo and TSA-bound pseudo-ensembles composed of all structures (42 and 46 KSI molecules, respectively). $\Delta MDev_{Apo-TSA}$ indicates the difference between ΣMDev for Apo and ΣMDev for TSA-bound for either the entire enzyme or for the enzyme core (excluding loops 62-65 and 91-96).

| | $\Delta MDev_{Apo\text{-}TSA}$ / $\Sigma MDev_{Apo}$ |
|---|---|
| **Entire enzyme** | 0.092 |
| **Enzyme core** | 0.084 |

**Table S20.** The values represent the conformational heterogeneity dampening in Apo enzyme core upon TSA binding as obtained by dividing the difference between Cβ Apo and TSA-bound MDevs ($\Delta MDev_{Apo\text{-}TSA}$, from Table S16) by the sum of Cβ Apo MDevs ($\Sigma MDev_{Apo}$, from Table S16).

| PDB ID | Mutations | Ligand | Resolution (Å) | Space group | # molecules in AU |
|---|---|---|---|---|---|
| **1OCV** | F116W | - | 2.0 | $P3_1$ | 4 |
| **1OGZ** | P39A | Equilenin | 2.3 | $P6_522$ | 1 |
| **1OHP** | D38N | *5α-Estran-3,17-Dione* | 1.53 | $P2_1$ | 4 |
| **1OHS** | Y14F/D38N | *5α -androstane-3,17-dione* | 1.7 | $P2_1$ | 4 |
| **1QJG** | WT | Equilenin | 2.3 | $P2_1$ | 6 |
| **3M8C** | D99N | Equilenin | 2.1 | $P6_122$ | 4 |
| **3MHE** | P39A | - | 1.72 | $P2_12_12_1$ | 2 |
| **3MKI** | D38E/D99N | - | 2.0 | $P6_122$ | 4 |
| **3MYT** | D38H/D99N | Equilenin | 1.96 | $P6_122$ | 4 |
| **3NBR** | D38N/P39G/D99N | *4-Androstene-3,17-dione* | 1.73 | $P6_522$ | 1 |
| **3NHX** | D99N | *4-Androstene-3,17-dione* | 1.59 | $P6_522$ | 1 |
| **3NM2** | D38E/P39G/V40G/S42G | - | 1.89 | $P6_522$ | 1 |
| **3NUV** | D38N/D99N | *4-Androstene-3,17-dione* | 1.76 | $P3_112$ | 2 |
| **3NXJ** | D99N | - | 1.97 | $C222_1$ | 2 |
| **3OV4** | P39G/V40G/S42G | Equilenin | 1.83 | $P6_122$ | 4 |
| **3T8U** | Y14A/Y55F/D99A | - | 2.5 | $P6_122$ | 4 |
| **3UNL** | F54G | - | 2.52 | $P6_122$ | 4 |
| **4L7K** | D38E | - | 2.1 | $P6_122$ | 12 |
| **5DRE** | D38G/P39G/D99N | - | 2.15 | $P6_522$ | 1 |
| **5UGI** | D38G/F54A | Equilenin | 1.8 | $P6_522$ | 1 |
| **8CHO** | WT | - | 2.3 | $P6_522$ | 1 |

**Table S21.** All $KSI_{homolog}$ crystal structures available from the PDB (here we use '$KSI_{homolog}$' to refer to the KSI from the organism *C. testosterone,* see Materials and Methods).

| PDB ID# | Apo | GSA/TSA-bound | Reduced |
|---|---|---|---|
| **1OCV** | X | | |
| **1OGZ** | | | |
| **1OHP** | | X | X |
| **1OHS** | X* | X | X |
| **1QJG** | | X | X |
| **3M8C** | X* | X | X |
| **3MHE** | | | |
| **3MKI** | X | | |
| **3MYT** | | | |
| **3NBR** | | | |
| **3NHX** | | X | X |
| **3NM2** | | | |
| **3NUV** | | X | X |
| **3NXJ** | X | | X |
| **3OV4** | | | |
| **3T8U** | X | | X |
| **3UNL** | X | | |
| **4L7K** | X | | |
| **5DRE** | | | |
| **5UGI** | | | |
| **8CHO** | | | X |

* $KSI_{homolog}$ molecules from these structures contained both ligand-bound and Apo molecules which were included in either the GSA/TSA-bound or in the Apo pseudo-ensembles, respectively (see **Table S23**).

**Table S22.** Different $KSI_{homolog}$ crystal structures included in the various pseudo-ensembles used in this work. Due to the relatively low total number of bound $KSI_{homolog}$ GSA- and TSA-bound molecules (9 and 9 molecules, respectively) available from the different bound structures, we did not attempt building separate GSA-bound and TSA-bound pseudo-ensembles. $KSI_{homolog}$ Apo structures with molecules in which active site bound ligands were not catalytic cycle analogs (i.e. sulfate or glycerol) were excluded from the pseudo-ensembles (PDB 3MKI, molecules B-D; PDB 4L7K, molecules A, D, F, H, J; PDB 8CHO). $KSI_{homolog}$ GSA/TSA-bound structures with molecules lacking any bound ligand (PDB 1OHS, molecule D; PDB 3M8C, molecule A) were included in the Apo pseudo-ensemble. PDB 1OHP, molecule B was excluded from both ensembles, as the GSA was not fully bound in the active site and the catalytic state represented by this KSI structure in not defined. $KSI_{homolog}$ crystal structures with mutations that alter the chemical nature of the general base or with mutations in the general base loop (residues 38-43) which are known to substantially increase the local flexibility were also excluded to eliminate artifacts (PDBs 1OGZ, 3MHE, 3MYT, 3NBR, 3NM2, 3OV4, 5DRE, and 5UGI).

| Apo | GSA/TSA–bound |
|---|---|
| 1OCV_A | 1OHS_A |
| 1OCV_B | 1OHS_B |
| 1OCV_C | 1OHS_C |
| 1OCV_D | 1OHP_A |
| 1OHS_D | 1OHP_C |
| 3NXJ_A | 1OHP_D |
| 3NXJ_B | 1QJG_E |
| 3M8C_A | 1GJG_F |
| 3MKI_A | 3NHX_A |
| 3T8U_A | 3NUV_A |
| 3T8U_B | 3NUV_B |
| 3T8U_C | 1QJG_A |
| 3T8U_D | 1QJG_B |
| 3UNL_A | 1QJG_C |
| 3UNL_B | 1QJG_D |
| 3UNL_C | 3M8C_B |
| 3UNL_D | 3M8C_C |
| 4L7K_B | 3M8C_D |
| 4L7K_C | |
| 4L7K_E | |
| 4L7K_G | |
| 4L7K_I | |
| 4L7K_K | |
| 4L7K_O | |

**Table S23.** $KSI_{homolog}$ molecules from the PDB crystal structures used to obtain Apo and GSA/TSA–bound pseudo-ensembles.

| Residue | Apo Cα MDev (Å) | GSA/TSA Cα MDev (Å) |
|---|---|---|
| **3** | 0.338 | 0.430 |
| **4** | 0.356 | 0.380 |
| **5** | 0.287 | 0.326 |
| **6** | 0.183 | 0.201 |
| **7** | 0.197 | 0.172 |
| **8** | 0.225 | 0.151 |
| **9** | 0.212 | 0.165 |
| **10** | 0.185 | 0.120 |
| **11** | 0.194 | 0.154 |
| **12** | 0.259 | 0.193 |
| **13** | 0.252 | 0.207 |
| **14** | 0.235 | 0.166 |
| **15** | 0.237 | 0.149 |
| **16** | 0.256 | 0.164 |
| **17** | 0.275 | 0.176 |
| **18** | 0.255 | 0.246 |
| **19** | 0.314 | 0.273 |
| **20** | 0.329 | 0.289 |
| **21** | 0.305 | 0.327 |
| **22** | 0.267 | 0.261 |
| **23** | 0.251 | 0.322 |
| **24** | 0.288 | 0.283 |
| **25** | 0.295 | 0.241 |
| **26** | 0.211 | 0.204 |
| **27** | 0.231 | 0.186 |
| **28** | 0.243 | 0.199 |
| **29** | 0.253 | 0.178 |
| **30** | 0.220 | 0.151 |
| **31** | 0.234 | 0.215 |
| **32** | 0.411 | 0.343 |
| **33** | 0.368 | 0.324 |
| **34** | 0.221 | 0.173 |
| **35** | 0.195 | 0.178 |
| **36** | 0.205 | 0.168 |
| **37** | 0.220 | 0.191 |
| **38** | 0.280 | 0.325 |
| **39** | 0.218 | 0.307 |
| **40** | 0.230 | 0.261 |
| **41** | 0.401 | 0.381 |
| **42** | 0.380 | 0.389 |
| **43** | 0.393 | 0.459 |
| **44** | 0.377 | 0.354 |
| **45** | 0.329 | 0.326 |
| **46** | 0.327 | 0.280 |
| **47** | 0.302 | 0.233 |
| **48** | 0.241 | 0.235 |
| **49** | 0.317 | 0.265 |

| | | |
|---|---|---|
| **50** | 0.274 | 0.217 |
| **51** | 0.236 | 0.167 |
| **52** | 0.295 | 0.199 |
| **53** | 0.338 | 0.236 |
| **54** | 0.242 | 0.243 |
| **55** | 0.216 | 0.215 |
| **56** | 0.314 | 0.271 |
| **57** | 0.354 | 0.344 |
| **58** | 0.357 | 0.349 |
| **59** | 0.294 | 0.339 |
| **60** | 0.361 | 0.324 |
| **61** | 0.424 | 0.351 |
| **62** | 0.334 | 0.411 |
| **63** | 0.270 | 0.239 |
| **64** | 0.214 | 0.167 |
| **65** | 0.222 | 0.174 |
| **66** | 0.340 | 0.251 |
| **67** | 0.287 | 0.178 |
| **68** | 0.281 | 0.191 |
| **69** | 0.261 | 0.179 |
| **70** | 0.219 | 0.191 |
| **71** | 0.179 | 0.155 |
| **72** | 0.193 | 0.168 |
| **73** | 0.192 | 0.181 |
| **74** | 0.224 | 0.221 |
| **75** | 0.298 | 0.279 |
| **76** | 0.252 | 0.234 |
| **77** | 0.176 | 0.138 |
| **78** | 0.175 | 0.106 |
| **79** | 0.178 | 0.109 |
| **80** | 0.163 | 0.113 |
| **81** | 0.167 | 0.108 |
| **82** | 0.183 | 0.133 |
| **83** | 0.164 | 0.167 |
| **84** | 0.170 | 0.190 |
| **85** | 0.241 | 0.268 |
| **86** | 0.264 | 0.299 |
| **87** | 0.423 | 0.480 |
| **88** | 0.940 | 1.177 |
| **89** | 1.078 | 1.607 |
| **90** | 0.874 | 1.282 |
| **91** | 0.603 | 0.657 |
| **92** | 0.472 | 0.412 |
| **93** | 0.300 | 0.311 |
| **94** | 0.207 | 0.207 |
| **95** | 0.192 | 0.179 |
| **96** | 0.167 | 0.151 |
| **97** | 0.178 | 0.139 |
| **98** | 0.189 | 0.131 |

| | | |
|---|---|---|
| **99** | 0.178 | 0.136 |
| **100** | 0.183 | 0.159 |
| **101** | 0.180 | 0.149 |
| **102** | 0.149 | 0.113 |
| **103** | 0.194 | 0.131 |
| **104** | 0.350 | 0.260 |
| **105** | 0.462 | 0.394 |
| **106** | 0.382 | 0.305 |
| **107** | 0.317 | 0.214 |
| **108** | 0.246 | 0.183 |
| **109** | 0.215 | 0.141 |
| **110** | 0.233 | 0.158 |
| **111** | 0.228 | 0.212 |
| **112** | 0.187 | 0.227 |
| **113** | 0.169 | 0.161 |
| **114** | 0.209 | 0.176 |
| **115** | 0.221 | 0.123 |
| **116** | 0.277 | 0.198 |
| **117** | 0.302 | 0.258 |
| **118** | 0.311 | 0.290 |
| **119** | 0.262 | 0.301 |
| **120** | 0.228 | 0.196 |
| **121** | 0.282 | 0.254 |
| **122** | 0.407 | 0.333 |

**Table S24.** Cα MDev values for Apo and GSA/TSA-bound pseudo-ensembles composed of all Apo or GSA/TSA-bound $KSI_{homolog}$ molecules (from Table S23).

| | Cα ΣMDev entire enzyme (Å) | Cα ΣMDev enzyme core (Å) |
|---|---|---|
| **Apo** | 33.945 | 29.555 |
| **TSA** | 31.626 | 26.012 |
| **$\Delta$MDev$_{Apo-GSA/TSA}$** | 2.319 | 3.543 |

**Table S25.** Sum of Cα MDev values (ΣMDev; from Table S24) for Apo and GSA/TSA-bound pseudo-ensembles composed of the $KSI_{homolog}$ molecules (24 and 18 KSI molecules, respectively, Table S23). $\Delta MDev_{Apo-GSA/TSA}$ indicates the difference between ΣMDev for Apo and ΣMDev for TSA-bound for either the entire enzyme or for the enzyme core (excluding the 86-92 loop).

| | $\Delta MDev_{Apo\text{-}GSA/TSA}$ / $\Sigma MDev_{Apo}$ |
|---|---|
| **Entire enzyme** | 0.068 |
| **Enzyme core** | 0.112 |

**Table S26.** The values represent the conformational heterogeneity dampening in Apo enzyme core upon GSA/TSA binding as obtained by dividing the difference between Cα Apo and TSA-bound MDevs ($\Delta MDev_{Apo\text{-}TSA}$, from Table S25) by the sum of Cα Apo MDevs ($\Sigma MDev_{Apo}$, from Table S25).

| Bootstrap cycles | Standard deviation (Å) | |
|---|---|---|
| | Apo pseudo-ensemble | TSA-bound pseudo-ensemble |
| 2 | 1.2746 | 0.82326 |
| 5 | 1.0870 | 0.47490 |
| 10 | 0.90315 | 0.40654 |
| 20 | 0.77570 | 0.44091 |
| 30 | 0.67611 | 0.41783 |
| 40 | 0.66826 | 0.41948 |
| 50 | 0.64100 | 0.43871 |
| 100 | 0.58084 | 0.44315 |
| 200 | 0.59049 | 0.41500 |
| 300 | 0.60243 | 0.41161 |

**Table S27.** Standard deviation values from increasing number of bootstrap cycles used to estimate the error associated with Apo and TSA-bound Cα ΣMDev (Figure 4E from main text and Figure supplement 7, see also Materials and Methods).

| (1-S$^2$) | | | | |
|---|---|---|---|---|
| **Residue #** | **Apo** | **GSA-bound observed** | **GSA-bound corrected** | **TSA-bound** |
| **5** | 0.566 | 0.489 | 0.456 | 0.414 |
| **6** | 0.473 | 0.421 | 0.398 | 0.399 |
| **7** | 0.842 | 0.792 | 0.770 | 0.577 |
| **8** | 0.506 | 0.437 | 0.407 | 0.398 |
| **9** | 0.382 | 0.346 | 0.330 | 0.384 |
| **10** | 0.461 | 0.373 | 0.335 | 0.349 |
| **11** | 0.395 | 0.338 | 0.314 | 0.342 |
| **12** | 0.374 | 0.354 | 0.346 | 0.306 |
| **13** | 0.337 | 0.291 | 0.271 | 0.283 |
| **14** | 0.396 | 0.347 | 0.326 | 0.396 |
| **15** | 0.397 | 0.374 | 0.364 | 0.392 |
| **16** | 0.336 | 0.310 | 0.299 | 0.294 |
| **17** | 0.383 | 0.324 | 0.299 | 0.332 |
| **18** | 0.658 | 0.798 | 0.858 | 0.790 |
| **19** | 0.384 | 0.364 | 0.355 | 0.318 |
| **20** | 0.374 | 0.372 | 0.371 | 0.295 |
| **21** | 0.390 | 0.363 | 0.351 | 0.338 |
| **22** | 0.550 | 0.513 | 0.496 | 0.476 |
| **23** | 0.405 | 0.404 | 0.403 | 0.346 |
| **24** | 0.440 | 0.424 | 0.417 | 0.428 |
| **25** | 0.527 | 0.485 | 0.467 | 0.529 |
| **26** | 0.564 | 0.720 | 0.787 | 0.738 |
| **27** | 0.379 | 0.392 | 0.398 | 0.342 |
| **28** | 0.337 | 0.345 | 0.348 | 0.301 |
| **29** | 0.460 | 0.406 | 0.383 | 0.448 |
| **30** | 0.669 | 0.760 | 0.799 | 0.748 |
| **31** | 0.383 | 0.369 | 0.362 | 0.326 |
| **32** | 0.331 | 0.341 | 0.346 | 0.324 |
| **33** | 0.436 | 0.431 | 0.429 | 0.430 |
| **34** | 0.623 | 0.754 | 0.810 | 0.728 |
| **35** | 0.672 | 0.758 | 0.795 | 0.775 |
| **36** | 0.414 | 0.385 | 0.372 | 0.345 |
| **37** | 0.621 | 0.638 | 0.645 | 0.606 |
| **38** | 0.393 | 0.358 | 0.343 | 0.331 |
| **39** | 0.369 | 0.377 | 0.381 | 0.326 |
| **40** | 0.380 | 0.405 | 0.416 | 0.357 |
| **41** | 0.454 | 0.418 | 0.402 | 0.343 |
| **42** | 0.401 | 0.409 | 0.412 | 0.323 |
| **43** | 0.417 | 0.439 | 0.448 | 0.329 |
| **44** | 0.588 | 0.546 | 0.528 | 0.458 |
| **45** | 0.704 | 0.613 | 0.574 | 0.714 |
| **46** | 0.696 | 0.670 | 0.659 | 0.681 |
| **47** | 0.691 | 0.781 | 0.819 | 0.726 |
| **48** | 0.749 | 0.837 | 0.875 | 0.783 |
| **49** | 0.433 | 0.488 | 0.512 | 0.447 |
| **50** | 0.484 | 0.569 | 0.605 | 0.580 |

| | | | | |
|---|---|---|---|---|
| **51** | 0.522 | 0.516 | 0.514 | 0.473 |
| **52** | 0.460 | 0.496 | 0.511 | 0.486 |
| **53** | 0.380 | 0.385 | 0.387 | 0.373 |
| **54** | 0.440 | 0.433 | 0.430 | 0.397 |
| **55** | 0.470 | 0.455 | 0.449 | 0.509 |
| **56** | 0.457 | 0.432 | 0.421 | 0.437 |
| **57** | 0.399 | 0.380 | 0.371 | 0.331 |
| **58** | 0.862 | 0.811 | 0.789 | 0.687 |
| **59** | 0.774 | 0.859 | 0.895 | 0.642 |
| **60** | 0.630 | 0.571 | 0.546 | 0.533 |
| **61** | 0.606 | 0.548 | 0.523 | 0.517 |
| **62** | 0.803 | 0.761 | 0.742 | 0.650 |
| **63** | 0.964 | 0.821 | 0.759 | 0.830 |
| **64** | 0.975 | 0.772 | 0.684 | 0.784 |
| **65** | 1.045 | 0.822 | 0.726 | 0.852 |
| **66** | 0.663 | 0.552 | 0.505 | 0.413 |
| **67** | 0.597 | 0.480 | 0.429 | 0.453 |
| **68** | 0.414 | 0.381 | 0.367 | 0.346 |
| **69** | 0.717 | 0.716 | 0.715 | 0.764 |
| **70** | 0.356 | 0.312 | 0.292 | 0.352 |
| **71** | 0.391 | 0.326 | 0.298 | 0.336 |
| **72** | 0.369 | 0.340 | 0.327 | 0.340 |
| **73** | 0.556 | 0.406 | 0.342 | 0.517 |
| **74** | 0.343 | 0.328 | 0.322 | 0.286 |
| **75** | 0.300 | 0.263 | 0.246 | 0.348 |
| **76** | 0.461 | 0.376 | 0.339 | 0.401 |
| **77** | 0.473 | 0.582 | 0.629 | 0.378 |
| **78** | 0.468 | 0.394 | 0.362 | 0.355 |
| **79** | 0.491 | 0.416 | 0.384 | 0.380 |
| **80** | 0.402 | 0.334 | 0.305 | 0.324 |
| **81** | 0.370 | 0.310 | 0.284 | 0.313 |
| **82** | 0.303 | 0.264 | 0.248 | 0.246 |
| **83** | 0.287 | 0.252 | 0.237 | 0.254 |
| **84** | 0.304 | 0.250 | 0.227 | 0.252 |
| **85** | 0.296 | 0.259 | 0.243 | 0.256 |
| **86** | 0.302 | 0.238 | 0.210 | 0.252 |
| **87** | 0.469 | 0.394 | 0.362 | 0.390 |
| **88** | 0.437 | 0.634 | 0.718 | 0.769 |
| **89** | 0.540 | 0.460 | 0.426 | 0.419 |
| **90** | 0.721 | 0.552 | 0.480 | 0.621 |
| **91** | 0.916 | 0.598 | 0.462 | 0.684 |
| **92** | 1.090 | 0.509 | 0.260 | 0.660 |
| **93** | 1.123 | 0.688 | 0.502 | 0.721 |
| **94** | 1.131 | 0.709 | 0.528 | 0.825 |
| **95** | 1.078 | 0.735 | 0.588 | 0.884 |
| **96** | 0.884 | 0.556 | 0.415 | 0.718 |
| **97** | 0.857 | 0.577 | 0.456 | 0.687 |
| **98** | 0.539 | 0.411 | 0.356 | 0.412 |
| **99** | 0.384 | 0.341 | 0.323 | 0.316 |

| | | | | |
|---|---|---|---|---|
| **100** | 0.364 | 0.304 | 0.278 | 0.263 |
| **101** | 0.304 | 0.272 | 0.258 | 0.224 |
| **102** | 0.276 | 0.229 | 0.209 | 0.219 |
| **103** | 0.333 | 0.257 | 0.224 | 0.363 |
| **104** | 0.319 | 0.285 | 0.270 | 0.259 |
| **105** | 0.333 | 0.290 | 0.272 | 0.287 |
| **106** | 0.390 | 0.355 | 0.339 | 0.318 |
| **107** | 0.392 | 0.360 | 0.346 | 0.332 |
| **108** | 0.550 | 0.482 | 0.453 | 0.533 |
| **109** | 0.924 | 0.898 | 0.887 | 0.889 |
| **110** | 0.857 | 0.714 | 0.652 | 0.649 |
| **111** | 0.558 | 0.539 | 0.531 | 0.431 |
| **112** | 0.492 | 0.498 | 0.501 | 0.496 |
| **113** | 0.353 | 0.351 | 0.350 | 0.289 |
| **114** | 0.474 | 0.459 | 0.452 | 0.431 |
| **115** | 0.588 | 0.660 | 0.691 | 0.542 |
| **116** | 0.429 | 0.377 | 0.354 | 0.402 |
| **117** | 0.366 | 0.328 | 0.311 | 0.317 |
| **118** | 0.346 | 0.320 | 0.309 | 0.260 |
| **119** | 0.279 | 0.271 | 0.267 | 0.229 |
| **120** | 0.337 | 0.298 | 0.281 | 0.300 |
| **121** | 0.432 | 0.394 | 0.378 | 0.385 |
| **122** | 0.770 | 0.508 | 0.395 | 0.600 |
| **123** | 0.611 | 0.381 | 0.282 | 0.434 |
| **124** | 0.357 | 0.283 | 0.251 | 0.267 |
| **125** | 0.572 | 0.490 | 0.454 | 0.541 |

**Table S28.** Crystallographic disorder parameters obtained from the 250 K multi-conformer models of KSI Apo, GSA-bound and TSA-bound (Table S3) .The values are the average of the two molecules from the crystallographic asymmetric unit. The total occupancy of the GSA in the two GSA-bound molecules was 1.4 instead of 2.0 (1.0 occupancy for each KSI molecule in the asymmetric unit). The corrected (1-$S^2$) values were obtained using the relationship $(1-S^2)\ GSA_{corrected} = ((1-S^2)GSA_{observed} - 0.3x(1-S^2)Apo) \big/ 0.7$

| Δ(1-$S^2$) | | | |
|---|---|---|---|
| **Residue #** | **Apo – GSA $_{corrected}$** | **GSA$_{corrected}$ – TSA** | **Apo – TSA** |
| **5** | 0.11 | 0.042 | 0.152 |
| **6** | 0.075 | -0.001 | 0.074 |
| **7** | 0.072 | 0.193 | 0.265 |
| **8** | 0.099 | 0.009 | 0.108 |
| **9** | 0.052 | -0.054 | -0.002 |
| **10** | 0.126 | -0.014 | 0.112 |
| **11** | 0.081 | -0.028 | 0.053 |
| **12** | 0.028 | 0.04 | 0.068 |
| **13** | 0.066 | -0.012 | 0.054 |
| **14** | 0.07 | -0.07 | 0 |
| **15** | 0.033 | -0.028 | 0.005 |
| **16** | 0.037 | 0.005 | 0.042 |
| **17** | 0.084 | -0.033 | 0.051 |
| **18** | -0.2 | 0.068 | -0.132 |
| **19** | 0.029 | 0.037 | 0.066 |
| **20** | 0.003 | 0.076 | 0.079 |
| **21** | 0.039 | 0.013 | 0.052 |
| **22** | 0.054 | 0.02 | 0.074 |
| **23** | 0.002 | 0.057 | 0.059 |
| **24** | 0.023 | -0.011 | 0.012 |
| **25** | 0.06 | -0.062 | -0.002 |
| **26** | -0.223 | 0.049 | -0.174 |
| **27** | -0.019 | 0.056 | 0.037 |
| **28** | -0.011 | 0.047 | 0.036 |
| **29** | 0.077 | -0.065 | 0.012 |
| **30** | -0.13 | 0.051 | -0.079 |
| **31** | 0.021 | 0.036 | 0.057 |
| **32** | -0.015 | 0.022 | 0.007 |
| **33** | 0.007 | -0.001 | 0.006 |
| **34** | -0.187 | 0.082 | -0.105 |
| **35** | -0.123 | 0.02 | -0.103 |
| **36** | 0.042 | 0.027 | 0.069 |
| **37** | -0.024 | 0.039 | 0.015 |
| **38** | 0.05 | 0.012 | 0.062 |
| **39** | -0.012 | 0.055 | 0.043 |
| **40** | -0.036 | 0.059 | 0.023 |
| **41** | 0.052 | 0.059 | 0.111 |
| **42** | -0.011 | 0.089 | 0.078 |
| **43** | -0.031 | 0.119 | 0.088 |
| **44** | 0.06 | 0.07 | 0.13 |
| **45** | 0.13 | -0.14 | -0.01 |
| **46** | 0.037 | -0.022 | 0.015 |
| **47** | -0.128 | 0.093 | -0.035 |
| **48** | -0.126 | 0.092 | -0.034 |
| **49** | -0.079 | 0.065 | -0.014 |
| **50** | -0.121 | 0.025 | -0.096 |

| | | | |
|---|---|---|---|
| **51** | 0.008 | 0.041 | 0.049 |
| **52** | -0.051 | 0.025 | -0.026 |
| **53** | -0.007 | 0.014 | 0.007 |
| **54** | 0.01 | 0.033 | 0.043 |
| **55** | 0.021 | -0.06 | -0.039 |
| **56** | 0.036 | -0.016 | 0.02 |
| **57** | 0.028 | 0.04 | 0.068 |
| **58** | 0.073 | 0.102 | 0.175 |
| **59** | -0.121 | 0.253 | 0.132 |
| **60** | 0.084 | 0.013 | 0.097 |
| **61** | 0.083 | 0.006 | 0.089 |
| **62** | 0.061 | 0.092 | 0.153 |
| **63** | 0.205 | -0.071 | 0.134 |
| **64** | 0.291 | -0.1 | 0.191 |
| **65** | 0.319 | -0.126 | 0.193 |
| **66** | 0.158 | 0.092 | 0.25 |
| **67** | 0.168 | -0.024 | 0.144 |
| **68** | 0.047 | 0.021 | 0.068 |
| **69** | 0.002 | -0.049 | -0.047 |
| **70** | 0.064 | -0.06 | 0.004 |
| **71** | 0.093 | -0.038 | 0.055 |
| **72** | 0.042 | -0.013 | 0.029 |
| **73** | 0.214 | -0.175 | 0.039 |
| **74** | 0.021 | 0.036 | 0.057 |
| **75** | 0.054 | -0.102 | -0.048 |
| **76** | 0.122 | -0.062 | 0.06 |
| **77** | -0.156 | 0.251 | 0.095 |
| **78** | 0.106 | 0.007 | 0.113 |
| **79** | 0.107 | 0.004 | 0.111 |
| **80** | 0.097 | -0.019 | 0.078 |
| **81** | 0.086 | -0.029 | 0.057 |
| **82** | 0.055 | 0.002 | 0.057 |
| **83** | 0.05 | -0.017 | 0.033 |
| **84** | 0.077 | -0.025 | 0.052 |
| **85** | 0.053 | -0.013 | 0.04 |
| **86** | 0.092 | -0.042 | 0.05 |
| **87** | 0.107 | -0.028 | 0.079 |
| **88** | -0.281 | -0.051 | -0.332 |
| **89** | 0.114 | 0.007 | 0.121 |
| **90** | 0.241 | -0.141 | 0.1 |
| **91** | 0.454 | -0.222 | 0.232 |
| **92** | 0.83 | -0.4 | 0.43 |
| **93** | 0.621 | -0.219 | 0.402 |
| **94** | 0.603 | -0.297 | 0.306 |
| **95** | 0.49 | -0.296 | 0.194 |
| **96** | 0.469 | -0.303 | 0.166 |
| **97** | 0.401 | -0.231 | 0.17 |
| **98** | 0.183 | -0.056 | 0.127 |
| **99** | 0.061 | 0.007 | 0.068 |

| | | | |
|---|---|---|---|
| **100** | 0.086 | 0.015 | 0.101 |
| **101** | 0.046 | 0.034 | 0.08 |
| **102** | 0.067 | -0.01 | 0.057 |
| **103** | 0.109 | -0.139 | -0.03 |
| **104** | 0.049 | 0.011 | 0.06 |
| **105** | 0.061 | -0.015 | 0.046 |
| **106** | 0.051 | 0.021 | 0.072 |
| **107** | 0.046 | 0.014 | 0.06 |
| **108** | 0.097 | -0.08 | 0.017 |
| **109** | 0.037 | -0.002 | 0.035 |
| **110** | 0.205 | 0.003 | 0.208 |
| **111** | 0.027 | 0.1 | 0.127 |
| **112** | -0.009 | 0.005 | -0.004 |
| **113** | 0.003 | 0.061 | 0.064 |
| **114** | 0.022 | 0.021 | 0.043 |
| **115** | -0.103 | 0.149 | 0.046 |
| **116** | 0.075 | -0.048 | 0.027 |
| **117** | 0.055 | -0.006 | 0.049 |
| **118** | 0.037 | 0.049 | 0.086 |
| **119** | 0.012 | 0.038 | 0.05 |
| **120** | 0.056 | -0.019 | 0.037 |
| **121** | 0.054 | -0.007 | 0.047 |
| **122** | 0.375 | -0.205 | 0.17 |
| **123** | 0.329 | -0.152 | 0.177 |
| **124** | 0.106 | -0.016 | 0.09 |
| **125** | 0.118 | -0.087 | 0.031 |

**Table S29.** Difference crystallographic disorder parameters ($\Delta(1\text{-}S^2)$ values from Table S28).

| | $\Sigma(1\text{-}S^2)$ entire enzyme | $\Sigma(1\text{-}S^2)$ `enzyme core (Å) |
|---|---|---|
| **Apo** | 63.02 | 53.01 |
| **GSA-bound observed** | 56.71 | 49.74 |
| **GSA-bound corrected** | 54.00 | 48.34 |
| **TSA-bound** | 55.30 | 47.69 |

**Table S30**. Sum of crystallographic disorder parameters ($1\text{-}S^2$) (from Table S29) obtained for Apo, GSA-bound (observed and corrected) and TSA-bound 250 K KSI multi-conformer models.

| | **Δ(1-$S^2$)**<br>**entire enzyme** | **Δ(1-$S^2$)**<br>**`enzyme core (Å)** |
|---|---|---|
| **Apo – $GSA_{observed}$** | 6.31 | 3.27 |
| **Apo – $GSA_{corrected}$** | 9.04 | 4.67 |
| **$GSA_{observed}$ – TSA** | 1.41 | 2.05 |
| **$GSA_{corrected}$ – TSA** | -1.30 | 0.65 |
| **Apo – TSA** | 7.72 | 5.32 |

**Table S31**. Difference crystallographic disorder parameters (Δ(Σ1-$S^2$) values from Table S30)

| | **Entire enzyme** | | | **Enzyme core** | | |
|---|---|---|---|---|---|---|
| | **$\Delta\Sigma(1-S^2)$** | **$\Sigma(1-S^2)_{Apo}$** | **$\Delta\Sigma(1-S^2)$ / $\Sigma(1-S^2)_{Apo}$** | **$\Delta\Sigma(1-S^2)$** | **$\Sigma(1-S^2)_{Apo}$** | **$\Delta\Sigma(1-S^2)$ / $\Sigma(1-S^2)_{Apo}$** |
| **Apo → GSA (observed)** | 6.31 | 63.02 | 0.10 | 3.27 | 53.01 | 0.06 |
| **Apo → GSA (corrected)** | 9.02 | | 0.14 | 4.67 | | 0.09 |
| | **$\Delta\Sigma(1-S^2)$** | **$\Sigma(1-S^2)_{GSA}$** | **$\Delta\Sigma(1-S^2)$ / $\Sigma(1-S^2)_{GSA}$** | **$\Delta\Sigma(1-S^2)$** | **$\Sigma(1-S^2)_{GSA}$** | **$\Delta\Sigma(1-S^2)$ / $\Sigma(1-S^2)_{GSA}$** |
| **GSA (observed) → TSA** | 1.41 | 56.71 | 0.02 | 2.05 | 49.74 | 0.04 |
| **GSA (corrected) → TSA** | -1.30 | 54.00 | -0.02 | 0.65 | 48.34 | 0.01 |
| | **$\Delta\Sigma(1-S^2)$** | **$\Sigma(1-S^2)_{Apo}$** | **$\Delta\Sigma(1-S^2)$ / $\Sigma(1-S^2)_{Apo}$** | **$\Delta\Sigma(1-S^2)$** | **$\Sigma(1-S^2)_{Apo}$** | **$\Delta\Sigma(1-S^2)$ / $\Sigma(1-S^2)_{Apo}$** |
| **Apo → TSA** | 7.72 | 63.02 | 0.12 | 5.32 | 53.01 | 0.1 |

**Table S32**. The values in the $\Delta\Sigma(1-S^2)$ / $\Delta(1-S^2)$ columns represent the conformational heterogeneity dampening in a given KSI catalytic state relative to the preceding catalytic state – i.e. in the GSA-bound state relative to the Apo state and in the TSA-bound state relative to the GSA-bound state. The last row indicates the conformational heterogeneity dampening in the TSA-bound state with respect to the Apo state. The conformational heterogeneity dampening for both the entire enzyme or the enzyme core (excluding 62-65 and 91-96 loops) was obtained by dividing the difference of the $(1-S^2)$ sums of two states *x* and *y* ($\Delta\Sigma(1-S^2)$), by the sum of $(1-S^2)$ for *x*. The conformational heterogeneity dampening was calculated using both the observed and corrected $(1-S^2)$ for the GSA-bound state. Using either the observed or corrected values led to analogous results and conclusions.

| Residue # | Reduced pseudo-ensemble | | RT-ensemble | |
|---|---|---|---|---|
| | **Oδ1** | **O/Nδ1** | **Oδ1** | **O/Nδ2** |
| **D21** | 0.270 | 0.271 | 0.395 | 0.524 |
| **D24** | 0.390 | 0.544 | 0.679 | 0.759 |
| **D34** | 1.608 | 1.564 | 2.068 | 1.768 |
| **D35** | 1.005 | 1.057 | 2.800 | 1.405 |
| **D40** | 0.528 | 0.809 | 0.337 | 0.954 |
| **N79** | 1.322 | 1.294 | 0.979 | 1.081 |
| **D100** | 0.226 | 0.278 | 0.456 | 0.461 |
| **D103** | 0.191 | 0.196 | 0.271 | 0.370 |
| **D108** | 0.688 | 0.688 | 0.585 | 0.891 |
| **N124** | 0.256 | 0.233 | 0.296 | 0.355 |

**Table S33**. Reduced pseudo-ensemble and RT-ensemble MDev values for all aspartate and asparagine residues Oδ1 and O/Nδ2 atoms (Oδ2 if the residue is aspartate and Nδ2 if the residue is asparagine). Asparagine 93 is situated in the middle of the 91-96 flexible loop and was therefore not included in the comparison.

| Residue # | Reduced pseudo-ensemble | | RT-ensemble | |
|---|---|---|---|---|
| | **OH** | **Cζ** | **OH** | **Cζ** |
| **Y16** | 0.214 | | 0.335 | |
| **Y32** | 0.305 | | 0.367 | |
| **F42** | | 0.436 | | 0.484 |
| **F56** | | 0.687 | | 0.668 |
| **Y57** | 0.318 | | 0.321 | |
| **F86** | | 0.177 | | 0.234 |
| **F107** | | 0.184 | | 0.279 |
| **Y119** | 0.416 | | 0.414 | |

**Table S34**. Reduced pseudo-ensemble and RT-ensemble MDev values for all tyrosine and phenylalanine residues OH and Cζ atoms, respectively.

| Residue # | Cα MDev (Å) | | | Cα ΔMDev (Å) | |
|---|---|---|---|---|---|
| | Full | Reduced | RT | RT–Full | RT–reduced |
| **5** | 0.269 | 0.231 | 0.278 | 0.009 | 0.047 |
| **6** | 0.250 | 0.203 | 0.234 | -0.016 | 0.031 |
| **7** | 0.281 | 0.252 | 0.286 | 0.005 | 0.033 |
| **8** | 0.223 | 0.210 | 0.331 | 0.108 | 0.121 |
| **8** | 0.164 | 0.141 | 0.273 | 0.109 | 0.132 |
| **9** | 0.191 | 0.165 | 0.243 | 0.051 | 0.078 |
| **10** | 0.206 | 0.179 | 0.313 | 0.107 | 0.134 |
| **11** | 0.171 | 0.146 | 0.266 | 0.095 | 0.120 |
| **12** | 0.173 | 0.152 | 0.170 | -0.004 | 0.017 |
| **13** | 0.187 | 0.174 | 0.223 | 0.036 | 0.049 |
| **14** | 0.201 | 0.181 | 0.232 | 0.030 | 0.051 |
| **15** | 0.194 | 0.180 | 0.222 | 0.028 | 0.042 |
| **16** | 0.201 | 0.193 | 0.212 | 0.010 | 0.019 |
| **17** | 0.212 | 0.182 | 0.289 | 0.076 | 0.107 |
| **18** | 0.239 | 0.202 | 0.296 | 0.058 | 0.094 |
| **19** | 0.279 | 0.226 | 0.275 | -0.005 | 0.049 |
| **20** | 0.270 | 0.222 | 0.365 | 0.096 | 0.143 |
| **21** | 0.265 | 0.239 | 0.472 | 0.207 | 0.233 |
| **22** | 0.299 | 0.256 | 0.366 | 0.067 | 0.111 |
| **23** | 0.339 | 0.300 | 0.283 | -0.056 | -0.017 |
| **24** | 0.390 | 0.309 | 0.293 | -0.096 | -0.016 |
| **25** | 0.424 | 0.320 | 0.388 | -0.035 | 0.068 |
| **26** | 0.329 | 0.252 | 0.313 | -0.016 | 0.060 |
| **27** | 0.251 | 0.215 | 0.210 | -0.041 | -0.005 |
| **28** | 0.273 | 0.252 | 0.228 | -0.044 | -0.023 |
| **29** | 0.285 | 0.251 | 0.252 | -0.033 | 0.001 |
| **30** | 0.204 | 0.183 | 0.248 | 0.043 | 0.065 |
| **31** | 0.191 | 0.188 | 0.290 | 0.099 | 0.101 |
| **32** | 0.194 | 0.189 | 0.332 | 0.139 | 0.143 |
| **33** | 0.470 | 0.502 | 0.806 | 0.336 | 0.304 |
| **34** | 0.511 | 0.564 | 0.810 | 0.298 | 0.245 |
| **35** | 0.188 | 0.197 | 0.421 | 0.233 | 0.223 |
| **36** | 0.253 | 0.239 | 0.311 | 0.058 | 0.072 |
| **37** | 0.279 | 0.249 | 0.255 | -0.024 | 0.006 |
| **39** | 0.286 | 0.251 | 0.240 | -0.046 | -0.012 |
| **40** | 0.321 | 0.283 | 0.276 | -0.045 | -0.007 |
| **41** | 0.367 | 0.349 | 0.238 | -0.129 | -0.111 |
| **42** | 0.370 | 0.334 | 0.266 | -0.104 | -0.068 |
| **43** | 0.442 | 0.392 | 0.382 | -0.060 | -0.010 |
| **44** | 0.448 | 0.424 | 0.423 | -0.025 | -0.001 |
| **45** | 0.441 | 0.430 | 0.529 | 0.087 | 0.099 |
| **46** | 0.356 | 0.326 | 0.358 | 0.002 | 0.032 |
| **47** | 0.313 | 0.263 | 0.372 | 0.059 | 0.109 |
| **48** | 0.259 | 0.220 | 0.403 | 0.144 | 0.183 |
| **49** | 0.506 | 0.550 | 0.935 | 0.429 | 0.385 |
| **50** | 0.399 | 0.419 | 0.655 | 0.256 | 0.236 |
| **51** | 0.416 | 0.420 | 0.567 | 0.151 | 0.147 |

| | | | | | |
|---|---|---|---|---|---|
| **52** | 0.316 | 0.306 | 0.439 | 0.122 | 0.132 |
| **53** | 0.274 | 0.268 | 0.252 | -0.022 | -0.016 |
| **54** | 0.325 | 0.281 | 0.323 | -0.002 | 0.042 |
| **55** | 0.381 | 0.329 | 0.356 | -0.024 | 0.027 |
| **56** | 0.379 | 0.325 | 0.306 | -0.073 | -0.019 |
| **57** | 0.376 | 0.296 | 0.256 | -0.120 | -0.040 |
| **58** | 0.455 | 0.335 | 0.274 | -0.181 | -0.061 |
| **59** | 0.494 | 0.377 | 0.307 | -0.187 | -0.070 |
| **60** | 0.504 | 0.448 | 0.346 | -0.158 | -0.102 |
| **61** | 0.412 | 0.359 | 0.331 | -0.080 | -0.028 |
| **62** | 0.980 | 0.803 | 0.500 | -0.480 | -0.303 |
| **63** | 1.526 | 1.264 | 1.470 | -0.056 | 0.206 |
| **64** | 2.375 | 2.519 | 1.861 | -0.514 | -0.658 |
| **65** | 0.728 | 0.650 | 0.804 | 0.076 | 0.154 |
| **66** | 0.369 | 0.376 | 0.344 | -0.025 | -0.032 |
| **67** | 0.341 | 0.333 | 0.300 | -0.041 | -0.033 |
| **68** | 0.233 | 0.212 | 0.305 | 0.072 | 0.093 |
| **69** | 0.291 | 0.234 | 0.240 | -0.051 | 0.006 |
| **70** | 0.239 | 0.199 | 0.194 | -0.045 | -0.005 |
| **71** | 0.315 | 0.253 | 0.188 | -0.127 | -0.065 |
| **72** | 0.243 | 0.212 | 0.207 | -0.037 | -0.005 |
| **73** | 0.233 | 0.209 | 0.217 | -0.015 | 0.009 |
| **74** | 0.142 | 0.123 | 0.147 | 0.005 | 0.024 |
| **75** | 0.167 | 0.137 | 0.116 | -0.050 | -0.021 |
| **76** | 0.180 | 0.168 | 0.179 | -0.001 | 0.011 |
| **77** | 0.265 | 0.238 | 0.215 | -0.050 | -0.022 |
| **78** | 0.268 | 0.229 | 0.257 | -0.011 | 0.028 |
| **79** | 0.267 | 0.226 | 0.316 | 0.049 | 0.090 |
| **80** | 0.224 | 0.213 | 0.323 | 0.099 | 0.111 |
| **81** | 0.195 | 0.163 | 0.236 | 0.041 | 0.073 |
| **82** | 0.181 | 0.146 | 0.248 | 0.067 | 0.102 |
| **83** | 0.145 | 0.119 | 0.114 | -0.032 | -0.006 |
| **84** | 0.122 | 0.114 | 0.137 | 0.015 | 0.023 |
| **85** | 0.185 | 0.157 | 0.134 | -0.051 | -0.023 |
| **86** | 0.189 | 0.162 | 0.144 | -0.045 | -0.018 |
| **87** | 0.257 | 0.220 | 0.144 | -0.113 | -0.076 |
| **88** | 0.234 | 0.237 | 0.210 | -0.023 | -0.027 |
| **89** | 0.281 | 0.214 | 0.250 | -0.032 | 0.035 |
| **90** | 0.371 | 0.293 | 0.414 | 0.043 | 0.121 |
| **91** | 0.770 | 0.670 | 0.695 | -0.075 | 0.025 |
| **92** | 1.342 | 1.178 | 1.090 | -0.252 | -0.089 |
| **93** | 2.139 | 2.091 | 2.341 | 0.202 | 0.250 |
| **94** | 1.579 | 1.470 | 1.182 | -0.397 | -0.288 |
| **95** | 0.852 | 0.802 | 1.037 | 0.185 | 0.234 |
| **96** | 0.539 | 0.437 | 0.675 | 0.136 | 0.238 |
| **97** | 0.424 | 0.406 | 0.405 | -0.018 | 0.000 |
| **98** | 0.393 | 0.355 | 0.360 | -0.033 | 0.005 |
| **99** | 0.300 | 0.281 | 0.247 | -0.053 | -0.034 |
| **100** | 0.210 | 0.201 | 0.208 | -0.002 | 0.007 |

| | | | | | |
|---|---|---|---|---|---|
| **101** | 0.209 | 0.190 | 0.142 | -0.067 | -0.048 |
| **102** | 0.182 | 0.160 | 0.122 | -0.059 | -0.038 |
| **103** | 0.202 | 0.157 | 0.179 | -0.023 | 0.022 |
| **104** | 0.172 | 0.150 | 0.160 | -0.012 | 0.010 |
| **105** | 0.143 | 0.124 | 0.156 | 0.013 | 0.032 |
| **106** | 0.154 | 0.132 | 0.255 | 0.102 | 0.123 |
| **107** | 0.153 | 0.137 | 0.307 | 0.154 | 0.170 |
| **108** | 0.200 | 0.187 | 0.356 | 0.156 | 0.169 |
| **109** | 0.305 | 0.277 | 0.437 | 0.132 | 0.160 |
| **110** | 0.296 | 0.241 | 0.518 | 0.223 | 0.278 |
| **111** | 0.243 | 0.208 | 0.422 | 0.179 | 0.214 |
| **112** | 0.162 | 0.135 | 0.254 | 0.092 | 0.119 |
| **113** | 0.144 | 0.127 | 0.291 | 0.148 | 0.164 |
| **114** | 0.154 | 0.133 | 0.293 | 0.139 | 0.160 |
| **115** | 0.210 | 0.207 | 0.217 | 0.007 | 0.009 |
| **116** | 0.320 | 0.303 | 0.207 | -0.113 | -0.096 |
| **117** | 0.308 | 0.271 | 0.167 | -0.141 | -0.103 |
| **118** | 0.318 | 0.282 | 0.219 | -0.099 | -0.064 |
| **119** | 0.283 | 0.262 | 0.185 | -0.098 | -0.077 |
| **120** | 0.344 | 0.333 | 0.206 | -0.138 | -0.127 |
| **121** | 0.457 | 0.445 | 0.289 | -0.169 | -0.157 |
| **122** | 0.445 | 0.438 | 0.325 | -0.120 | -0.113 |
| **123** | 0.332 | 0.315 | 0.305 | -0.027 | -0.010 |
| **124** | 0.279 | 0.262 | 0.187 | -0.092 | -0.075 |
| **125** | 0.345 | 0.333 | 0.294 | -0.051 | -0.038 |

**Table S35**. Full pseudo-ensemble, reduced pseudo-ensemble and RT-ensemble MDev values for Cα atoms of residues 5-125.

| Residue # | Cβ MDev (Å) | | | Cβ ΔMDev (Å) | |
|---|---|---|---|---|---|
| | Full | Reduced | RT | RT–Full | RT–reduced |
| **5** | 0.299 | 0.260 | 0.313 | 0.014 | 0.053 |
| **6** | 0.263 | 0.207 | 0.262 | -0.001 | 0.055 |
| **7** | 0.359 | 0.321 | 0.297 | -0.062 | -0.025 |
| **8** | 0.248 | 0.242 | 0.378 | 0.130 | 0.136 |
| **8** | 0.177 | 0.147 | 0.248 | 0.071 | 0.101 |
| **9** | 0.216 | 0.188 | 0.267 | 0.051 | 0.080 |
| **10** | - | - | - | - | - |
| **11** | 0.173 | 0.149 | 0.263 | 0.090 | 0.114 |
| **12** | 0.202 | 0.201 | 0.175 | -0.027 | -0.026 |
| **13** | 0.223 | 0.193 | 0.302 | 0.079 | 0.109 |
| **14** | 0.243 | 0.236 | 0.310 | 0.067 | 0.074 |
| **15** | 0.221 | 0.220 | 0.254 | 0.033 | 0.034 |
| **16** | 0.226 | 0.216 | 0.277 | 0.050 | 0.061 |
| **17** | 0.295 | 0.261 | 0.330 | 0.034 | 0.069 |
| **18** | 0.248 | 0.210 | 0.308 | 0.061 | 0.099 |
| **19** | 0.292 | 0.233 | 0.257 | -0.035 | 0.024 |
| **20** | 0.301 | 0.230 | 0.330 | 0.029 | 0.100 |
| **21** | 0.298 | 0.291 | 0.469 | 0.170 | 0.178 |
| **22** | - | - | - | - | - |
| **23** | 0.360 | 0.321 | 0.336 | -0.024 | 0.015 |
| **24** | 0.424 | 0.315 | 0.393 | -0.031 | 0.077 |
| **25** | 0.552 | 0.421 | 0.486 | -0.066 | 0.066 |
| **26** | 0.372 | 0.286 | 0.368 | -0.004 | 0.082 |
| **27** | 0.278 | 0.235 | 0.226 | -0.051 | -0.009 |
| **28** | 0.320 | 0.294 | 0.313 | -0.008 | 0.019 |
| **29** | 0.366 | 0.319 | 0.312 | -0.054 | -0.008 |
| **30** | 0.222 | 0.193 | 0.315 | 0.092 | 0.122 |
| **31** | 0.222 | 0.211 | 0.302 | 0.080 | 0.091 |
| **32** | 0.248 | 0.245 | 0.433 | 0.185 | 0.188 |
| **33** | 0.643 | 0.632 | 0.934 | 0.291 | 0.302 |
| **34** | 0.648 | 0.701 | 0.900 | 0.252 | 0.199 |
| **35** | 0.177 | 0.167 | 0.398 | 0.221 | 0.231 |
| **36** | 0.350 | 0.330 | 0.404 | 0.054 | 0.074 |
| **37** | 0.308 | 0.281 | 0.291 | -0.018 | 0.010 |
| **39** | 0.310 | 0.268 | 0.289 | -0.022 | 0.021 |
| **40** | 0.337 | 0.304 | 0.325 | -0.012 | 0.021 |
| **41** | 0.395 | 0.378 | 0.298 | -0.097 | -0.080 |
| **42** | 0.379 | 0.333 | 0.303 | -0.076 | -0.030 |
| **43** | - | - | - | - | - |
| **44** | 0.514 | 0.450 | 0.466 | -0.048 | 0.016 |
| **45** | 0.534 | 0.538 | 0.625 | 0.091 | 0.088 |
| **46** | 0.419 | 0.385 | 0.416 | -0.003 | 0.031 |
| **47** | 0.427 | 0.368 | 0.489 | 0.063 | 0.121 |
| **48** | 0.399 | 0.341 | 0.649 | 0.250 | 0.308 |
| **49** | - | - | - | - | - |
| **50** | 0.487 | 0.523 | 0.773 | 0.287 | 0.251 |
| **51** | 0.539 | 0.554 | 0.744 | 0.205 | 0.190 |

| | | | | | |
|---|---|---|---|---|---|
| **52** | 0.348 | 0.335 | 0.419 | 0.071 | 0.084 |
| **53** | 0.284 | 0.279 | 0.284 | -0.001 | 0.005 |
| **54** | 0.352 | 0.300 | 0.378 | 0.025 | 0.078 |
| **55** | 0.417 | 0.352 | 0.413 | -0.005 | 0.061 |
| **56** | 0.411 | 0.367 | 0.296 | -0.114 | -0.070 |
| **57** | 0.380 | 0.309 | 0.264 | -0.116 | -0.044 |
| **58** | 0.468 | 0.381 | 0.341 | -0.126 | -0.040 |
| **59** | 0.636 | 0.506 | 0.398 | -0.238 | -0.108 |
| **60** | - | - | - | - | - |
| **61** | 0.440 | 0.350 | 0.303 | -0.137 | -0.046 |
| **62** | - | - | - | - | - |
| **63** | - | - | - | - | - |
| **64** | - | - | - | - | - |
| **65** | 1.127 | 1.146 | 1.027 | -0.100 | -0.119 |
| **66** | 0.445 | 0.409 | 0.364 | -0.081 | -0.045 |
| **67** | 0.441 | 0.396 | 0.307 | -0.134 | -0.089 |
| **68** | 0.260 | 0.234 | 0.420 | 0.161 | 0.187 |
| **69** | 0.478 | 0.370 | 0.336 | -0.142 | -0.034 |
| **70** | 0.292 | 0.223 | 0.244 | -0.048 | 0.021 |
| **71** | 0.341 | 0.260 | 0.277 | -0.064 | 0.017 |
| **72** | - | - | - | - | - |
| **73** | 0.296 | 0.251 | 0.275 | -0.020 | 0.024 |
| **74** | 0.159 | 0.143 | 0.132 | -0.027 | -0.011 |
| **75** | 0.184 | 0.171 | 0.160 | -0.025 | -0.011 |
| **76** | 0.201 | 0.179 | 0.261 | 0.060 | 0.083 |
| **77** | 0.364 | 0.320 | 0.274 | -0.090 | -0.046 |
| **78** | 0.292 | 0.254 | 0.331 | 0.039 | 0.077 |
| **79** | 0.326 | 0.272 | 0.401 | 0.075 | 0.129 |
| **80** | - | - | - | - | - |
| **81** | 0.265 | 0.235 | 0.241 | -0.024 | 0.007 |
| **82** | - | - | - | - | - |
| **83** | 0.210 | 0.158 | 0.214 | 0.004 | 0.057 |
| **84** | 0.142 | 0.128 | 0.221 | 0.079 | 0.094 |
| **85** | 0.260 | 0.207 | 0.220 | -0.040 | 0.013 |
| **86** | 0.184 | 0.168 | 0.162 | -0.022 | -0.007 |
| **87** | 0.312 | 0.276 | 0.195 | -0.118 | -0.081 |
| **88** | 0.320 | 0.273 | 0.355 | 0.035 | 0.081 |
| **89** | 0.393 | 0.304 | 0.391 | -0.002 | 0.087 |
| **90** | 0.516 | 0.434 | 0.446 | -0.070 | 0.012 |
| **91** | 1.018 | 0.910 | 0.888 | -0.131 | -0.022 |
| **92** | 1.529 | 1.319 | 1.210 | -0.318 | -0.109 |
| **93** | 2.529 | 2.479 | 2.709 | 0.179 | 0.229 |
| **94** | - | - | - | - | - |
| **95** | 1.114 | 1.128 | 1.346 | 0.232 | 0.218 |
| **96** | 0.697 | 0.537 | 0.807 | 0.111 | 0.270 |
| **97** | 0.550 | 0.535 | 0.483 | -0.068 | -0.052 |
| **98** | 0.440 | 0.406 | 0.469 | 0.029 | 0.064 |
| **99** | 0.322 | 0.309 | 0.271 | -0.050 | -0.038 |
| **100** | 0.195 | 0.197 | 0.262 | 0.067 | 0.066 |

| | | | | | |
|---|---|---|---|---|---|
| **101** | 0.246 | 0.224 | 0.230 | -0.015 | 0.006 |
| **102** | 0.198 | 0.161 | 0.147 | -0.051 | -0.014 |
| **103** | 0.236 | 0.175 | 0.303 | 0.067 | 0.129 |
| **104** | 0.216 | 0.178 | 0.255 | 0.039 | 0.078 |
| **105** | 0.187 | 0.168 | 0.189 | 0.002 | 0.021 |
| **106** | 0.194 | 0.184 | 0.281 | 0.087 | 0.098 |
| **107** | 0.163 | 0.146 | 0.360 | 0.197 | 0.214 |
| **108** | 0.220 | 0.206 | 0.462 | 0.242 | 0.256 |
| **109** | 0.400 | 0.370 | 0.569 | 0.170 | 0.199 |
| **110** | 0.336 | 0.268 | 0.569 | 0.233 | 0.301 |
| **111** | | | | | |
| **112** | 0.221 | 0.185 | 0.248 | 0.027 | 0.063 |
| **113** | 0.160 | 0.149 | 0.295 | 0.134 | 0.146 |
| **114** | 0.190 | 0.184 | 0.278 | 0.088 | 0.095 |
| **115** | 0.322 | 0.306 | 0.332 | 0.010 | 0.026 |
| **116** | 0.418 | 0.380 | 0.317 | -0.101 | -0.063 |
| **117** | 0.333 | 0.306 | 0.231 | -0.102 | -0.075 |
| **118** | 0.356 | 0.296 | 0.261 | -0.094 | -0.035 |
| **119** | 0.273 | 0.243 | 0.204 | -0.069 | -0.039 |
| **120** | 0.334 | 0.313 | 0.207 | -0.127 | -0.107 |
| **121** | 0.477 | 0.468 | 0.381 | -0.097 | -0.087 |
| **122** | 0.514 | 0.497 | 0.408 | -0.107 | -0.089 |
| **123** | 0.364 | 0.371 | 0.361 | -0.003 | -0.010 |
| **124** | 0.284 | 0.268 | 0.214 | -0.070 | -0.054 |
| **125** | 0.422 | 0.405 | 0.413 | -0.009 | 0.008 |

**Table S36**. Full pseudo-ensemble, reduced pseudo-ensemble and RT-ensemble MDev values for Cβ atoms of residues 5-125.

| Residue # | Cγ2 MDev (Å) | | |
|---|---|---|---|
| | **Full** | **Reduced** | **RT** |
| **9** | 0.191 | 0.146 | 0.321 |
| **17** | 0.275 | 0.243 | 0.323 |
| **20** | 0.313 | 0.259 | 0.309 |
| **22** | 0.898 | 0.985 | 1.331 |
| **25** | 0.485 | 0.375 | 0.546 |
| **28** | 0.323 | 0.300 | 0.292 |
| **29** | 0.382 | 0.351 | 0.423 |
| **38** | 0.336 | 0.319 | 0.388 |
| **47** | 0.943 | 0.760 | 1.036 |
| **53** | 0.306 | 0.291 | 0.354 |
| **66** | 0.537 | 0.505 | 0.465 |
| **74** | 0.187 | 0.175 | 0.251 |
| **88** | 0.645 | 0.638 | 1.107 |
| **91** | 1.416 | 1.417 | 1.646 |
| **101** | 0.277 | 0.245 | 0.282 |
| **102** | 0.213 | 0.151 | 0.225 |
| **104** | 0.247 | 0.198 | 0.398 |
| **113** | 0.194 | 0.186 | 0.308 |
| **123** | 0.665 | 0.749 | 0.816 |

**Table S37**. Full pseudo-ensemble, reduced pseudo-ensemble and RT-ensemble MDev values for Cγ2 atoms of all KSI isoleucine and valine residues.

| Residue # | Cε MDev (Å) | | |
|---|---|---|---|
| | **Full** | **Reduced** | **RT** |
| **13** | 0.228 | 0.189 | 0.275 |
| **31** | 0.278 | 0.233 | 0.427 |
| **84** | 0.243 | 0.187 | 0.402 |
| **90** | 1.651 | 1.631 | 1.375 |
| **105** | 0.543 | 0.530 | 0.331 |
| **116** | 1.395 | 1.348 | 1.017 |

**Table S38**. Full pseudo-ensemble, reduced pseudo-ensemble and RT-ensemble MDev values for Cε atoms of all KSI methionine residues.

| Residue # | Oγ2/Sγ2 MDev (Å) | | |
|---|---|---|---|
| | **Full** | **Reduced** | **RT** |
| **69** | 1.276 | 1.370 | 1.395 |
| **77** | 0.682 | 0.450 | 0.907 |
| **81** | 0.370 | 0.290 | 0.329 |
| **97** | 1.319 | 1.373 | 1.690 |
| **121** | 0.449 | 0.421 | 0.626 |
| **126** | 1.138 | 1.206 | 1.056 |

**Table S39**. Full pseudo-ensemble, reduced pseudo-ensemble and RT-ensemble MDev values for Oγ2 and Sγ2 atoms of all KSI serine and cysteine residues, respectively.

| Residue # | Oε2/Nε2 MDev (Å) | | |
|---|---|---|---|
| | **Full** | **Reduced** | **RT** |
| **7** | 1.829 | 2.140 | 2.646 |
| **8** | 0.893 | 0.815 | 0.670 |
| **10** | 1.289 | 1.233 | 0.435 |
| **18** | 2.336 | 2.705 | 3.180 |
| **26** | 2.708 | 2.261 | 2.733 |
| **30** | 2.670 | 3.026 | 3.252 |
| **39** | 1.002 | 0.900 | 1.281 |
| **44** | 1.531 | 1.649 | 1.645 |
| **51** | 1.880 | 1.758 | 1.903 |
| **52** | 1.712 | 1.846 | 2.556 |
| **59** | 2.578 | 2.572 | 2.493 |
| **89** | 1.104 | 0.991 | 1.066 |
| **95** | 2.280 | 2.359 | 2.945 |
| **109** | 2.621 | 2.635 | 2.367 |
| **114** | 2.103 | 1.944 | 2.295 |
| **117** | 1.300 | 1.346 | 1.072 |
| **122** | 2.355 | 2.218 | 2.610 |

**Table S40**. Full pseudo-ensemble, reduced pseudo-ensemble and RT-ensemble MDev values for Oε2 and Nε2 atoms of all KSI glutamate and glutamine residues, respectively.

| Structure | Distance (Å) | |
|---|---|---|
| | Y16 OH – Equ Ox | D103 Oδ2 – Equ Ox |
| **1W6Y** | 2.56 | 2.58 |
| **1OH0_A** | 2.54 | 2.48 |
| **1OH0_B** | 2.55 | 2.56 |
| **3OWU_D** | 2.76 | 2.56 |
| **3OWU_A** | 2.7 | 2.5 |
| **3OWU_B** | 2.56 | 2.47 |
| **3OWU_C** | 2.69 | 2.47 |
| **1OGX_A** | 2.68 | 2.54 |
| **1OGX_B** | 2.66 | 2.55 |
| **3OWS_A** | 2.58 | 2.46 |
| **3OWS_B** | 2.53 | 2.5 |
| **3OWS_C** | 2.53 | 2.45 |
| **3OWS_D** | 2.57 | 2.49 |
| **3OWY_A** | 2.48 | 2.52 |
| **3OWY_B** | 3.07 | 2.38 |
| **3OWY_C** | 2.6 | 2.63 |
| **3OWY_D** | 2.66 | 2.38 |
| **3OWY_E** | 2.47 | 2.48 |
| **3OWY_H** | 3.09 | 2.42 |
| **Mean** | 2.60 | 2.50 |

**Table S41**. Lengths of Y16 and D103 hydrogen bonds made to the transition state analog equilenin in the ensemble of KSI crystal structures of variants with WT-like activity (Table S1-S2). The Y16 OH – Equ Ox distance from PDB 3OWY molecules B and H were not included in the calculation of the mean, because the measured distances of 3.07 and 3.09 Å appear too long for equilenin to be making a hydrogen bond with Y16. However the corresponding D103 Oδ2 – Equ Ox hydrogen bond lengths were included as the distances of 2.38 and 2.42 Å are within the expected range. This observation is onsistent with previous observations that Y16 and D103 hydrogen bonds can be formed independent from one another.

|  | **Packing residue** | **Packing atom** | **Y16 atom** | **Van der Waals sum** |
|---|---|---|---|---|
| **1** | V20 | Cγ2 | Cζ | 3.4-4.0 |
| **2** | M105 | Cε | Ring | 3.4-4.0 |
| **3** | M84 | Cε | Cδ2 | 3.4-4.0 |
| **4** | D103 | Oδ2 | Cε2 | 3.1-3.7 |
| **5** | I28 | Cγ2 | Cε1 | 3.4-4.0 |
| **6** | I17 | Cδ1 | Cδ2 | 3.4-4.0 |
| **7** | I113 | Cδ1 | Cδ1 | 3.4-4.0 |
| **8** | M31 | Sδ | Cδ1 | 3.5-3.8 |
| **9** | M116 | Cε | OH | 3.1-3.8 |

**Table S42**. Y16 packing residues and closest atoms making van der Waals interactions with Y16 identified from KSI crystal structures. Van der Waals sum indicates the sum of the van der Waals radii and is represented as a range because of uncertainty introduced by the absence of hydrogen coordinates in the X-ray structural models and because the oxygen $r_{vdw}$ is orientation-dependent (see **Table S48** for van der Waals radii values).

| # distance | 1 | 2 | 3 | 4 | 5 | 6 | 7 | 8 | 9 |
|---|---|---|---|---|---|---|---|---|---|
| **1** | 4.03 | 3.71 | 3.84 | 4.00 | 4.05 | 4.03 | 4.29 | 6.20 | 3.83 |
| **2** | 4.14 | 3.56 | 3.69 | 4.02 | 4.03 | 4.39 | 3.96 | 3.99 | 3.94 |
| **3** | 3.95 | 3.66 | 3.73 | 3.80 | 4.32 | 4.24 | 4.03 | 4.23 | 3.10 |
| **4** | 3.85 | 3.64 | 3.55 | 3.56 | 4.26 | 4.08 | 4.21 | 4.41 | 3.49 |
| **5** | 4.12 | 3.60 | 3.80 | 3.89 | 4.09 | 3.98 | 4.10 | 4.24 | 2.92 |
| **6** | 4.24 | 3.60 | 4.07 | 4.19 | 3.91 | 4.63 | 3.81 | 3.98 | 4.02 |
| **7** | 3.85 | 3.56 | 3.57 | 3.85 | 4.34 | 4.04 | 4.05 | 4.34 | 4.05 |
| **8** | 4.02 | 3.59 | 3.86 | 4.03 | 4.04 | 4.29 | 3.94 | 4.15 | 3.83 |
| **9** | 3.73 | 3.55 | 3.59 | 3.75 | 4.11 | 3.96 | 4.28 | 4.43 | 5.80 |
| **10** | 4.08 | 3.82 | 4.38 | 3.88 | 3.66 | 4.54 | 4.06 | 3.89 | 3.89 |
| **11** | 3.62 | 3.57 | 3.85 | 3.52 | 3.91 | 4.10 | 4.24 | 4.49 | 3.42 |
| **12** | 3.75 | 3.56 | 3.58 | 3.54 | 3.85 | 3.86 | 4.16 | 4.54 | 4.45 |
| **13** | 3.63 | 3.65 | 3.68 | 3.53 | 4.09 | 3.91 | 4.22 | 4.54 | 3.29 |
| **14** | 3.68 | 3.76 | 3.74 | 3.45 | 4.00 | 3.91 | 4.43 | 4.46 | 4.77 |
| **15** | 3.58 | 3.72 | 3.71 | 3.44 | 4.08 | 3.97 | 4.33 | 4.62 | 6.00 |
| **16** | 3.88 | 3.58 | 3.77 | 3.92 | 3.90 | 4.02 | 4.23 | 4.36 | 4.27 |
| **17** | 4.01 | 3.39 | 3.74 | 3.62 | 4.16 | 4.50 | 4.30 | 4.40 | 3.86 |
| **18** | 3.68 | 3.61 | 3.71 | 3.77 | 4.01 | 3.94 | 4.15 | 4.63 | 4.07 |
| **19** | 3.63 | 3.62 | 4.13 | 3.91 | 3.81 | 3.99 | 4.42 | 4.44 | 3.88 |
| **20** | 3.81 | 3.56 | 3.68 | 4.25 | 4.07 | 3.78 | 4.12 | 4.58 | 3.84 |
| **21** | 3.62 | 3.63 | 3.77 | 3.79 | 3.94 | 3.96 | 4.36 | 4.51 | 3.94 |
| **22** | 4.19 | 3.62 | 4.35 | 3.68 | 3.76 | 4.48 | 3.72 | 4.59 | 3.45 |
| **23** | 3.75 | 3.82 | 3.80 | 3.55 | 4.02 | 3.96 | 4.22 | 4.45 | 3.91 |
| **24** | 3.65 | 3.67 | 3.61 | 3.66 | 3.93 | 3.87 | 4.44 | 4.54 | 4.32 |
| **25** | 3.68 | 3.69 | 3.97 | 3.50 | 4.05 | 4.13 | 4.20 | 6.08 | 3.55 |
| **26** | 3.75 | 3.76 | 3.90 | 3.75 | 3.93 | 4.06 | 4.31 | 6.06 | 4.95 |
| **27** | 4.31 | 3.66 | 4.27 | 3.92 | 4.06 | 4.05 | 3.95 | 3.98 | 3.93 |
| **28** | 3.58 | 5.14 | 3.79 | 3.61 | 3.96 | 3.88 | 4.23 | 4.47 | 3.37 |
| **29** | 3.64 | 3.43 | 3.66 | 3.62 | 3.84 | 3.83 | 4.42 | 4.61 | 5.01 |
| **30** | 3.58 | 3.59 | 3.73 | 3.88 | 3.83 | 4.00 | 4.34 | 4.35 | 5.53 |
| **31** | 3.84 | 4.68 | 3.98 | 3.52 | 3.68 | 4.33 | 3.77 | 4.29 | 5.09 |
| **32** | 3.64 | 3.69 | 3.68 | 3.66 | 4.06 | 4.31 | 4.18 | 4.63 | 4.08 |
| **33** | 3.75 | 3.67 | 3.64 | 3.77 | 4.11 | 3.97 | 4.00 | 4.44 | 4.80 |
| **34** | 3.64 | 3.38 | 3.65 | 4.25 | 3.91 | 4.11 | 4.27 | 6.19 | 3.73 |
| **35** | 4.53 | 3.64 | 4.04 | 3.74 | 3.70 | 4.57 | 3.73 | 5.94 | 3.36 |
| **36** | 3.57 | 3.53 | 3.66 | 3.62 | 3.88 | 3.94 | 4.23 | 4.60 | 4.36 |
| **37** | 3.70 | 3.61 | 3.91 | 3.50 | 4.15 | 3.98 | 4.29 | 4.57 | 3.70 |
| **38** | 3.77 | 3.44 | 3.52 | 3.63 | 4.06 | 4.20 | 4.00 | 4.51 | 3.99 |
| **39** | 3.70 | 3.64 | 3.75 | 3.49 | 3.96 | 4.07 | 4.27 | 4.44 | 3.95 |
| **40** | 4.73 | 3.82 | 4.31 | 3.64 | 4.56 | 4.27 | 3.63 | 4.50 | 4.07 |
| **41** | 3.72 | 3.51 | 3.68 | 3.83 | 4.13 | 3.92 | 4.21 | 4.61 | 3.80 |
| **42** | 3.58 | 3.50 | 3.67 | 3.59 | 3.99 | 3.91 | 4.29 | 4.52 | 3.95 |
| **43** | 3.54 | 3.79 | 3.99 | 3.59 | 3.97 | 4.12 | 4.48 | 4.62 | 3.98 |
| **44** | 3.65 | 3.65 | 3.95 | 3.62 | 3.98 | 4.08 | 4.49 | 4.61 | 4.17 |
| **45** | 4.51 | 3.76 | 4.39 | 3.53 | 3.79 | 4.32 | 3.84 | 4.07 | 3.95 |
| **46** | 3.61 | 3.87 | 3.76 | 3.63 | 4.02 | 3.93 | 4.18 | 4.56 | 5.31 |
| **47** | 3.55 | 3.44 | 3.63 | 3.75 | 3.90 | 3.83 | 4.24 | 4.56 | 4.01 |

| | | | | | | | | | |
|---|---|---|---|---|---|---|---|---|---|
| **48** | 3.59 | 3.64 | 3.63 | 3.82 | 3.88 | 4.06 | 4.35 | 4.61 | 3.87 |
| **49** | 3.71 | 3.71 | 3.91 | 3.84 | 3.76 | 4.41 | 3.89 | 4.25 | 3.62 |
| **50** | 3.64 | 3.62 | 3.84 | 3.63 | 4.04 | 4.43 | 4.34 | 6.27 | 4.93 |
| **51** | 3.73 | 4.89 | 3.73 | 3.66 | 4.05 | 4.05 | 4.10 | 4.74 | 3.65 |
| **52** | 3.55 | 4.78 | 3.71 | 3.72 | 4.00 | 3.77 | 4.03 | 4.35 | 5.48 |
| **53** | 3.67 | 3.60 | 3.68 | 3.65 | 3.90 | 4.01 | 4.31 | 4.35 | 4.08 |
| **54** | 3.59 | 3.54 | 3.69 | 3.57 | 3.93 | 4.05 | 4.11 | 4.38 | 3.30 |
| **55** | 3.63 | 3.50 | 3.63 | 3.58 | 4.22 | 4.05 | 4.43 | 4.77 | 3.87 |
| **56** | 3.44 | 3.55 | 3.67 | 3.58 | 3.86 | 3.78 | 4.23 | 4.50 | 3.86 |
| **57** | 3.61 | 3.52 | 3.70 | 3.55 | 4.13 | 4.02 | 4.26 | 4.56 | 3.95 |
| **58** | 3.62 | 3.51 | 3.69 | 3.80 | 3.96 | 4.04 | 4.20 | 4.62 | 3.85 |
| **59** | 3.60 | 3.60 | 3.77 | 3.60 | 4.19 | 4.58 | 4.23 | 4.77 | 5.18 |
| **60** | 3.42 | 3.42 | 3.65 | 3.63 | 4.04 | 4.00 | 4.24 | 4.47 | 4.85 |
| **61** | 3.54 | 3.45 | 3.56 | 3.64 | 4.18 | 4.23 | 4.27 | 4.55 | 4.86 |
| **62** | 3.48 | 3.34 | 3.72 | 3.72 | 4.49 | 4.02 | 4.36 | 4.52 | 5.15 |
| **63** | 3.63 | 3.44 | 3.60 | 3.63 | 4.05 | 3.89 | 4.34 | 4.55 | 3.94 |
| **64** | 3.65 | 3.44 | 3.64 | 3.75 | 4.08 | 3.93 | 4.24 | 4.54 | 3.94 |
| **65** | 3.67 | 3.51 | 3.63 | 3.69 | 4.05 | 3.95 | 4.30 | 4.52 | 3.83 |
| **66** | 3.66 | 3.90 | 3.63 | 3.52 | 4.09 | 4.02 | 4.32 | 4.72 | 3.95 |
| **67** | 3.43 | 3.54 | 3.70 | 3.51 | 4.26 | 4.08 | 4.21 | 4.52 | 4.02 |
| **68** | 3.51 | 3.55 | 3.57 | 3.50 | 4.36 | 4.07 | 4.28 | 4.81 | 5.00 |
| **69** | 3.33 | 3.95 | 3.51 | 3.52 | 3.97 | 4.14 | 4.27 | 4.51 | 4.25 |
| **70** | 3.55 | 3.55 | 3.51 | 3.69 | 3.95 | 4.24 | 4.19 | 4.68 | 4.24 |
| **71** | 3.51 | 3.98 | 3.52 | 3.68 | 4.17 | 4.00 | 4.27 | 4.70 | 3.95 |
| **72** | 3.49 | 3.52 | 3.51 | 3.69 | 4.18 | 4.20 | 4.26 | 4.64 | 3.65 |
| **73** | 3.30 | 3.89 | 3.53 | 3.69 | 4.19 | 3.94 | 4.39 | 4.62 | 3.73 |
| **74** | 3.37 | 4.78 | 3.61 | 3.96 | 3.92 | 4.16 | 4.25 | 4.57 | 3.62 |
| **75** | 3.62 | 4.41 | 3.68 | 3.92 | 3.98 | 4.19 | 4.35 | 4.63 | 3.88 |
| **76** | 3.64 | 4.74 | 3.66 | 3.92 | 4.00 | 4.20 | 4.40 | 4.67 | 3.53 |
| **77** | 3.68 | 4.43 | 3.66 | 3.97 | 3.98 | 4.15 | 4.32 | 4.63 | 3.71 |
| **78** | 3.62 | 4.32 | 3.73 | | 3.89 | 4.16 | 4.37 | 4.61 | 3.63 |
| **79** | 3.78 | 4.30 | 3.68 | | 3.90 | 3.95 | 4.27 | 4.59 | 3.56 |
| **80** | 3.75 | 4.79 | 3.69 | | 3.83 | 4.00 | 4.26 | 4.63 | 3.53 |
| **81** | 3.76 | 4.48 | 3.66 | | 3.84 | 4.02 | 4.27 | 4.64 | 3.48 |
| **82** | 3.75 | 3.46 | 3.69 | | 3.98 | 4.00 | 4.29 | 4.51 | 3.50 |
| **83** | 3.67 | 3.43 | 3.69 | | 4.10 | 4.42 | 4.24 | 4.51 | 4.57 |
| **84** | 3.53 | 3.40 | 3.72 | | 4.08 | 4.46 | 4.14 | 4.52 | 4.55 |
| **85** | 3.53 | 3.48 | 3.73 | | 4.01 | 4.46 | 4.13 | 4.53 | 4.60 |
| **86** | 3.65 | 3.40 | 3.65 | | | 4.40 | 4.26 | | 4.63 |
| **87** | | 3.40 | | | | | | | 4.10 |
| **88** | | 3.42 | | | | | | | 4.16 |
| **89** | | 3.42 | | | | | | | 4.14 |
| **90** | | 3.65 | | | | | | | 4.07 |
| **91** | | 3.65 | | | | | | | |
| **92** | | 3.63 | | | | | | | |
| **93** | | 3.66 | | | | | | | |
| **94** | | | | | | | | | |

**Table S43**. Full pseudo-ensemble distances between Y16 atoms and packing residues closest atoms from **Table S42**. The number of distances varies depending on the packing group as a result of the presence of various mutations and, in some cases, alternative conformations for some residues.

| | Packing residue | Packing atom | D103 atom | Van der Waals sum |
|---|---|---|---|---|
| **1** | D103 | NH | Oδ1 | 2.95-3.25* |
| **2** | A118 | Cβ | Oδ1 | 3.1-3.7 |
| **3** | F86 | Cε1 | Oδ1 | 3.1-3.7 |
| **4** | V101 | Cγ1 | Oδ1 | 3.1-3.7 |
| **5** | Y16 | Cε2 | Oδ2 | 3.1-3.7 |
| **6** | M84 | Cε | Cβ | 3.4-4.0 |
| **7** | M84 | Cε | Oδ2 | 3.1-3.7 |
| **8** | M105 | Cε | Oδ2 | 3.1-3.7 |
| **9** | M116 | Cε | Oδ2 | 3.1-3.7 |
| **10** | M116 | Sδ | Oδ2 | 3.2-3.5 |

* The $r_{vdw}$ for N was used instead of NH.

**Table S44**. D103 packing residues and closest atoms making van der Waals interactions with D103 identified from KSI crystal structures. Van der Waals sum indicates the sum of the van der Waals radii and is represented as a range because of uncertainty introduced by the absence of hydrogen coordinates in the X-ray structural models and because the oxygen $r_{vdw}$ is orientation-dependent (see **Table S48** for van der Waals radii values).

| # distance | 1 | 2 | 3 | 4 | 5 | 6 | 7 | 8 | 9 | 10 |
|---|---|---|---|---|---|---|---|---|---|---|
| **1** | 3.13 | 3.52 | 3.40 | 3.25 | 4.00 | 4.08 | 4.46 | 4.51 | 3.59 | 3.63 |
| **2** | 3.25 | 3.34 | 3.64 | 3.62 | 4.02 | 4.14 | 4.17 | 4.02 | 4.02 | 3.86 |
| **3** | 3.44 | 3.19 | 3.78 | 3.85 | 3.80 | 3.93 | 3.99 | 3.90 | 3.73 | 3.86 |
| **4** | 3.32 | 3.26 | 3.59 | 3.65 | 3.56 | 3.82 | 3.98 | 4.29 | 4.03 | 3.86 |
| **5** | 3.32 | 3.24 | 3.67 | 3.65 | 3.89 | 3.99 | 4.12 | 4.41 | 3.91 | 3.88 |
| **6** | 3.26 | 3.46 | 3.46 | 3.62 | 4.19 | 3.99 | 3.97 | 3.93 | 4.06 | 4.06 |
| **7** | 3.26 | 3.56 | 3.37 | 3.55 | 3.85 | 3.93 | 3.98 | 4.28 | 4.00 | 4.00 |
| **8** | 3.28 | 3.36 | 3.58 | 3.62 | 4.03 | 4.16 | 4.05 | 4.16 | 4.21 | 3.76 |
| **9** | 3.23 | 3.26 | 3.53 | 3.58 | 3.75 | 4.19 | 4.30 | 4.08 | 4.00 | 3.92 |
| **10** | 3.34 | 3.01 | 3.84 | 3.83 | 3.88 | 4.46 | 4.21 | 4.35 | 4.14 | 4.23 |
| **11** | 3.18 | 3.25 | 3.53 | 3.45 | 3.52 | 4.13 | 4.08 | 4.08 | 3.90 | 4.09 |
| **12** | 3.22 | 3.21 | 3.56 | 3.46 | 3.54 | 4.03 | 4.14 | 4.29 | 3.64 | 3.43 |
| **13** | 3.20 | 3.15 | 3.51 | 3.45 | 3.53 | 4.08 | 4.05 | 3.74 | 3.65 | 3.68 |
| **14** | 3.33 | 3.21 | 3.59 | 3.68 | 3.45 | 3.97 | 3.87 | 4.08 | 4.02 | 3.94 |
| **15** | 3.24 | 3.17 | 3.46 | 3.61 | 3.44 | 3.84 | 4.08 | 3.72 | 3.80 | 3.09 |
| **16** | 3.28 | 3.30 | 3.31 | 3.70 | 3.92 | 3.84 | 4.64 | 3.99 | 3.17 | 4.71 |
| **17** | 3.20 | 3.21 | 3.46 | 3.42 | 3.62 | 4.10 | 4.05 | 4.07 | 3.57 | 3.83 |
| **18** | 3.20 | 2.98 | 3.65 | 3.91 | 3.77 | 3.94 | 4.12 | 4.30 | 3.99 | 3.90 |
| **19** | 3.24 | 3.12 | 3.69 | 3.48 | 3.91 | 4.11 | 4.34 | 3.74 | 3.77 | 4.58 |
| **20** | 3.35 | 3.26 | 3.97 | 4.03 | 4.25 | 3.88 | 3.93 | 4.13 | 3.86 | 4.39 |
| **21** | 3.28 | 3.37 | 3.50 | 3.73 | 3.79 | 4.10 | 3.78 | 4.04 | 4.04 | 4.01 |
| **22** | 3.22 | 3.21 | 3.41 | 3.56 | 3.68 | 3.65 | 4.02 | 5.46 | 3.93 | 3.94 |
| **23** | 3.18 | 3.19 | 3.39 | 3.54 | 3.55 | 3.84 | 3.93 | 4.09 | 3.26 | 3.26 |
| **24** | 3.21 | 3.26 | 3.49 | 3.41 | 3.66 | 3.92 | 4.06 | 3.94 | 3.69 | 3.80 |
| **25** | 3.21 | 3.16 | 3.48 | 3.49 | 3.50 | 4.16 | 4.04 | 4.35 | 4.04 | 3.19 |
| **26** | 3.26 | 3.04 | 3.29 | 3.40 | 3.75 | 4.14 | 4.19 | 4.22 | 4.51 | 4.30 |
| **27** | 3.23 | 3.16 | 3.48 | 3.45 | 3.92 | 4.40 | 4.08 | 3.87 | 4.61 | 3.88 |
| **28** | 3.21 | 3.19 | 3.54 | 3.53 | 3.61 | 3.96 | 4.01 | 4.01 | 3.72 | 3.95 |
| **29** | 3.15 | 3.20 | 3.46 | 3.58 | 3.62 | 4.04 | 4.01 | 4.06 | 4.19 | 4.19 |
| **30** | 3.13 | 3.19 | 3.55 | 3.51 | 3.88 | 4.04 | 4.05 | 4.04 | 4.02 | 3.73 |
| **31** | 3.37 | 3.16 | 3.68 | 3.80 | 3.52 | 3.93 | 3.96 | 4.13 | 4.24 | 3.96 |
| **32** | 3.23 | 3.65 | 3.36 | 3.68 | 3.66 | 4.11 | 3.87 | 3.64 | 4.34 | 4.28 |
| **33** | 3.23 | 3.13 | 3.55 | 3.54 | 3.77 | 3.91 | 4.26 | 4.03 | 4.10 | 3.89 |
| **34** | 3.24 | 2.84 | 3.88 | 4.04 | 4.25 | 3.94 | 4.57 | 4.07 | 3.98 | 3.90 |
| **35** | 3.36 | 3.31 | 3.58 | 3.62 | 3.74 | 3.99 | 4.00 | 4.00 | 3.86 | 4.02 |
| **36** | 3.24 | 3.18 | 3.43 | 3.51 | 3.62 | 3.86 | 4.13 | 4.16 | 4.01 | 3.95 |
| **37** | 3.20 | 3.15 | 3.38 | 3.43 | 3.50 | 4.25 | 4.12 | 4.25 | 3.80 | 3.89 |
| **38** | 3.19 | 3.39 | 3.46 | 3.42 | 3.63 | 3.90 | 4.09 | 3.94 | 2.89 | 3.54 |
| **39** | 3.16 | 3.18 | 3.48 | 3.40 | 3.49 | 4.04 | 4.20 | 4.12 | 3.31 | 3.30 |
| **40** | 3.32 | 3.02 | 3.26 | 3.59 | 3.64 | 4.03 | 3.81 | 6.21 | 3.83 | 4.40 |
| **41** | 3.20 | 3.18 | 3.45 | 3.50 | 3.83 | 3.94 | 4.06 | 5.76 | 4.97 | 3.58 |
| **42** | 3.22 | 3.23 | 3.51 | 3.52 | 3.59 | 3.96 | 4.02 | 4.09 | 4.27 | 4.09 |
| **43** | 3.23 | 3.22 | 3.45 | 3.60 | 3.59 | 4.21 | 3.99 | 3.97 | 4.42 | 4.42 |
| **44** | 3.20 | 3.22 | 3.53 | 3.53 | 3.62 | 4.26 | 4.05 | 4.12 | 3.03 | 4.75 |
| **45** | 3.34 | 3.32 | 3.72 | 3.83 | 3.53 | 4.24 | 3.91 | 4.06 | 3.22 | 4.91 |
| **46** | 3.23 | 3.25 | 3.55 | 3.60 | 3.63 | 4.04 | 4.20 | 3.97 | 3.25 | 4.98 |
| **47** | 3.15 | 3.38 | 3.44 | 3.53 | 3.75 | 4.08 | 3.99 | 3.93 | 3.18 | 4.84 |

| | | | | | | | | | | |
|---|---|---|---|---|---|---|---|---|---|---|
| **48** | 3.22 | 3.18 | 3.43 | 3.57 | 3.82 | 4.02 | 4.33 | 4.05 | 4.54 | 4.54 |
| **49** | 3.23 | 3.24 | 3.51 | 3.59 | 3.84 | 3.93 | 4.07 | 4.14 | 4.38 | 4.38 |
| **50** | 3.14 | 3.17 | 3.39 | 3.33 | 3.63 | 4.01 | 4.10 | 4.46 | 4.27 | 4.27 |
| **51** | 3.16 | 3.24 | 3.53 | 3.38 | 3.66 | 3.90 | 3.87 | 4.51 | 4.49 | 4.49 |
| **52** | 3.10 | 3.11 | 3.55 | 3.37 | 3.72 | 3.80 | 3.85 | 4.50 | 3.84 | 4.11 |
| **53** | 3.06 | 3.23 | 3.19 | 3.33 | 3.65 | 3.87 | 4.00 | 4.40 | 3.97 | 3.97 |
| **54** | 3.23 | 3.33 | 3.53 | 3.42 | 3.57 | 3.82 | 4.21 | 3.48 | 3.88 | 4.07 |
| **55** | 3.20 | 3.31 | 3.58 | 3.50 | 3.58 | 3.96 | 4.28 | 3.94 | 3.98 | 4.09 |
| **56** | 3.21 | 3.28 | 3.58 | 3.40 | 3.58 | 3.91 | 4.18 | 3.95 | 3.61 | 3.80 |
| **57** | 3.21 | 3.27 | 3.48 | 3.42 | 3.55 | 4.07 | 4.24 | 3.96 | 3.87 | 3.84 |
| **58** | 3.32 | 3.30 | 3.45 | 3.55 | 3.80 | 3.90 | 4.01 | 4.03 | 3.78 | 3.89 |
| **59** | 3.36 | 3.36 | 3.27 | 3.42 | 3.60 | 3.91 | 3.82 | 4.08 | 3.65 | 3.70 |
| **60** | 3.38 | 3.40 | 3.32 | 3.45 | 3.63 | 3.72 | 3.80 | 4.07 | 2.70 | 3.60 |
| **61** | 3.34 | 3.34 | 3.32 | 3.53 | 3.64 | 4.00 | 3.78 | 3.92 | 2.78 | 3.83 |
| **62** | 3.22 | 2.99 | 3.56 | 3.44 | 3.72 | 3.84 | 4.19 | 4.42 | 2.85 | 4.02 |
| **63** | 3.14 | 3.06 | 3.36 | 3.50 | 3.63 | 4.23 | 3.78 | 4.41 | 2.69 | 3.82 |
| **64** | 3.19 | 3.05 | 3.57 | 3.58 | 3.75 | 4.24 | 3.65 | 4.45 | 3.03 | 4.12 |
| **65** | 3.11 | 3.15 | 3.39 | 3.63 | 3.69 | 4.28 | 3.89 | 4.46 | 2.76 | 3.82 |
| **66** | 3.21 | 3.26 | 3.47 | 3.60 | 3.52 | 4.28 | 3.62 | 4.11 | 2.91 | 3.93 |
| **67** | 3.19 | 3.19 | 3.55 | 3.65 | 3.51 | 4.01 | 3.97 | 4.11 | 2.94 | 3.96 |
| **68** | 3.13 | 3.01 | 3.53 | 3.45 | 3.50 | 3.84 | 4.06 | 4.10 | 4.19 | 4.62 |
| **69** | 3.14 | 3.16 | 3.42 | 3.48 | 3.52 | 3.81 | 3.71 | 4.12 | 4.14 | 4.54 |
| **70** | 3.04 | 3.56 | 3.39 | 3.34 | 3.69 | 3.71 | 3.89 | 3.95 | 4.19 | 4.59 |
| **71** | 3.06 | 3.55 | 3.44 | 3.29 | 3.68 | 3.90 | 3.90 | 4.04 | 4.16 | 4.61 |
| **72** | 3.09 | 3.55 | 3.52 | 3.34 | 3.69 | 3.64 | 3.90 | 4.06 | 4.16 | 4.16 |
| **73** | 3.16 | 3.56 | 3.44 | 3.31 | 3.69 | 3.92 | 3.85 | 3.96 | 4.17 | 4.17 |
| **74** | 3.22 | 3.05 | 3.67 | 3.67 | 3.96 | 3.89 | 4.12 | | 4.21 | 4.21 |
| **75** | 3.18 | 3.09 | 3.60 | 3.67 | 3.92 | 3.70 | 4.13 | | 4.16 | 4.16 |
| **76** | 3.16 | 3.10 | 3.62 | 3.65 | 3.92 | 3.88 | 4.08 | | 3.83 | 3.78 |
| **77** | 3.18 | 3.06 | 3.65 | 3.71 | 3.97 | 3.88 | 4.11 | | 3.64 | 3.73 |
| **78** | 3.18 | 3.26 | 3.43 | 3.42 | | 3.69 | 4.01 | | 3.65 | 3.73 |
| **79** | 3.17 | 3.19 | 3.42 | 3.44 | | 3.91 | 4.22 | | 3.81 | 3.77 |
| **80** | 3.21 | 3.18 | 3.44 | 3.42 | | 3.88 | 4.21 | | | |
| **81** | 3.17 | 3.50 | 3.44 | 3.43 | | 3.93 | 4.01 | | | |
| **82** | | | | | | 3.86 | | | | |
| **83** | | | | | | 3.85 | | | | |
| **84** | | | | | | 3.89 | | | | |
| **85** | | | | | | 3.86 | | | | |
| **86** | | | | | | 3.86 | | | | |
| **87** | | | | | | 4.15 | | | | |
| **88** | | | | | | 4.18 | | | | |
| **89** | | | | | | 4.08 | | | | |
| **90** | | | | | | 4.16 | | | | |
| **91** | | | | | | 4.06 | | | | |
| **92** | | | | | | 4.17 | | | | |
| **93** | | | | | | 4.18 | | | | |
| **94** | | | | | | 4.07 | | | | |

**Table S45**. Full pseudo-ensemble distances between D103 atoms and packing residues closest atoms from **Table S44**. The number of distances varies depending on the packing group as a result of the presence of various mutations and, in some cases, alternative conformations for some residues.

| | Packing residue | Packing atom | General base atom | Van der Waals range |
|---|---|---|---|---|
| **1** | F56 | Cγ | Xδ2 | 3.1-3.7* |
| **2** | A118 | Cβ | Xδ2 | 3.1-3.7* |
| **3** | V38 | Cγ1 | Cβ | 3.4-4.0 |
| **4** | M116 | Cε | Xδ2 | 3.1-3.7* |

* A $r_{vdw}$ range of 1.4-1.7 Å was used for both O and $NH_2$ groups; The average N $r_{vdw}$ is ~ 1.55 Å, close to the average O $r_{vdw}$ of 1.5 Å. Xδ2 indicates either Nδ2 (asparagine) or Oδ2 (aspartate).

**Table S46**. General base packing residues and closest atoms making van der Waals interactions with general base identified from KSI crystal structures. Van der Waals sum indicates the sum of the van der Waals radii and is represented as a range because of uncertainty introduced by the absence of hydrogen coordinates in the X-ray structural models and because the oxygen $r_{vdw}$ is orientation-dependent (see **Table S48** for van der Waals radii values).

| # distances | 1 | 2 | 3 | 4 |
|---|---|---|---|---|
| **1** | 3.93 | 3.48 | 4.06 | 3.92 |
| **2** | 3.67 | 3.67 | 3.95 | 4.01 |
| **3** | 4.01 | 3.30 | 3.90 | 4.48 |
| **4** | 3.81 | 3.37 | 3.91 | 4.20 |
| **5** | 3.80 | 3.53 | 4.03 | 5.03 |
| **6** | 3.75 | 3.39 | 3.96 | 4.04 |
| **7** | 3.54 | 3.68 | 4.05 | 5.14 |
| **8** | 4.31 | 3.06 | 4.12 | 3.65 |
| **9** | 3.71 | 3.51 | 3.98 | 3.84 |
| **10** | 4.08 | 3.04 | 4.00 | 3.42 |
| **11** | 3.46 | 3.94 | 4.00 | 4.22 |
| **12** | 3.19 | 3.64 | 3.75 | 3.87 |
| **13** | 3.44 | 3.62 | 3.77 | 6.85 |
| **14** | 3.38 | 3.52 | 3.75 | 7.80 |
| **15** | 3.49 | 3.55 | 3.70 | 3.89 |
| **16** | 3.72 | 3.45 | 3.87 | 4.59 |
| **17** | 3.71 | 3.60 | 4.24 | 5.65 |
| **18** | 3.49 | 3.67 | 3.94 | 3.96 |
| **19** | 3.77 | 4.05 | 4.03 | 4.29 |
| **20** | 3.24 | 4.24 | 3.90 | 3.73 |
| **21** | 3.21 | 3.48 | 3.90 | 3.94 |
| **22** | 3.57 | 3.41 | 4.19 | 3.89 |
| **23** | 4.08 | 3.82 | 3.81 | 3.85 |
| **24** | 3.12 | 3.88 | 3.82 | 4.01 |
| **25** | 3.55 | 3.24 | 4.13 | 3.74 |
| **26** | 4.04 | 3.66 | 3.98 | 3.76 |
| **27** | 3.57 | 3.90 | 3.95 | 6.49 |
| **28** | 4.17 | 3.69 | 3.74 | 6.17 |
| **29** | 3.55 | 5.35 | 3.96 | 6.28 |
| **30** | 3.60 | 3.46 | 3.90 | 3.96 |
| **31** | 3.53 | 3.58 | 3.90 | 4.46 |
| **32** | 3.41 | 3.57 | 3.77 | 4.04 |
| **33** | 4.31 | 3.71 | 3.66 | 3.97 |
| **34** | 3.74 | 3.69 | 4.14 | 4.39 |
| **35** | 3.63 | 4.28 | 3.81 | 2.83 |
| **36** | 3.50 | 3.76 | 4.14 | 4.00 |
| **37** | 4.14 | 4.33 | 3.87 | 4.33 |
| **38** | 4.80 | 3.73 | 4.02 | 4.05 |
| **39** | 3.58 | 3.70 | 4.08 | 3.75 |
| **40** | 3.27 | 3.71 | 3.69 | 3.95 |
| **41** | 3.63 | 3.99 | 3.89 | 3.69 |
| **42** | 3.35 | 3.75 | 3.80 | 3.70 |
| **43** | 3.84 | 3.56 | 3.93 | 3.66 |
| **44** | 3.18 | 3.62 | 3.88 | 4.07 |
| **45** | 3.60 | 3.90 | 4.24 | 3.90 |
| **46** | 3.74 | 3.77 | 3.70 | 6.52 |
| **47** | 3.80 | 5.46 | 4.04 | 3.66 |
| **48** | 4.32 | 3.42 | 3.83 | 5.92 |

| | | | | |
|---|---|---|---|---|
| **49** | 3.45 | 3.65 | 3.86 | 6.53 |
| **50** | 3.52 | 3.56 | 3.90 | 3.82 |
| **51** | 3.56 | 3.82 | 3.68 | 4.39 |
| **52** | 3.63 | 3.70 | 4.05 | 3.67 |
| **53** | 4.14 | 3.44 | 3.81 | 2.85 |
| **54** | 3.53 | 3.67 | 4.14 | 3.66 |
| **55** | 3.53 | 3.61 | 4.01 | 3.82 |
| **56** | 3.67 | 3.66 | 3.95 | 3.73 |
| **57** | 3.95 | 3.55 | 4.03 | 3.65 |
| **58** | 3.64 | 3.56 | 3.93 | 3.59 |
| **59** | 3.41 | 3.81 | 4.00 | 4.17 |
| **60** | 3.21 | 3.76 | 3.92 | 4.09 |
| **61** | 3.15 | 3.65 | 3.96 | 3.92 |
| **62** | 3.15 | 3.58 | 3.80 | 3.94 |
| **63** | 3.17 | 3.81 | 3.81 | 4.08 |
| **64** | 3.34 | 3.77 | 3.68 | 4.14 |
| **65** | 3.23 | 3.88 | 3.84 | 4.09 |
| **66** | 3.35 | 3.81 | 3.77 | 7.02 |
| **67** | 3.34 | 3.70 | 3.70 | 6.81 |
| **68** | 3.48 | 3.72 | 3.91 | 6.71 |
| **69** | 3.64 | 3.59 | 3.88 | 6.55 |
| **70** | 3.19 | 3.61 | 3.89 | 6.94 |
| **71** | 3.58 | 3.79 | 4.03 | 6.45 |
| **72** | 3.27 | 3.75 | 4.11 | 6.41 |
| **73** | 3.57 | 3.86 | 4.04 | 6.95 |
| **74** | 3.67 | 3.71 | 3.93 | 4.14 |
| **75** | 3.21 | 3.85 | 4.07 | 4.04 |
| **76** | 3.53 | 3.62 | 4.00 | 4.13 |
| **77** | 3.61 | 3.45 | 4.01 | 4.07 |
| **78** | 3.61 | 3.89 | 4.03 | 3.98 |
| **79** | 3.57 | 3.74 | 4.00 | 3.81 |
| **80** | 3.67 | 3.69 | 3.91 | 4.09 |
| **81** | 3.66 | 3.70 | 3.98 | 3.95 |
| **82** | 3.73 | 3.72 | 3.99 | 4.10 |
| **83** | 3.67 | 3.41 | 3.96 | 3.95 |
| **84** | 3.45 | 3.42 | 3.95 | 4.00 |
| **85** | 3.33 | 3.34 | 3.98 | 4.06 |
| **86** | 4.69 | 3.42 | 3.98 | |
| **87** | 3.56 | 3.63 | 3.77 | |
| **88** | 3.49 | 3.40 | 3.86 | |
| **89** | 3.17 | 3.87 | 3.76 | |
| **90** | 3.16 | 3.59 | 3.80 | |
| **91** | 3.32 | 3.76 | 3.65 | |
| **92** | | 3.74 | 3.73 | |
| **93** | | 3.74 | 3.79 | |
| **94** | | 3.73 | 3.63 | |

**Table S47**. Full pseudo-ensemble distances between general base D40 atoms and packing residues closest atoms from **Table S46**. The number of distances varies depending on the packing group as a result of the presence of various mutations and, in some cases, alternative conformations for some residues.

| Atom/group | $r_{vdw}$ (Å) | Reference |
|---|---|---|
| C | 1.70 | (4) |
| N | 1.55 | (4) |
| O | 1.40-1.70 | (4) |
| S | 1.8 | (4) |
| $CH_3$ | 2.0 | (5) |

**Table S48.** Van der Waals radii of elements and groups.

| Structure | Y16 – Y57 hydrogen bond length (Å) |
|---|---|
| **1C7H** | 2.47 |
| **1DMQ** | 2.73 |
| **1OPY** | 2.66 |
| **1W6Y** | 2.33 |
| **2INX** | 2.43 |
| **3CPO** | 2.54 |
| **3RGR** | 2.57 |
| **3SED** | 2.59 |
| **5D81** | 2.42 |
| **1OH0_A** | 2.51 |
| **1OH0_B** | 2.53 |
| **3VSY_A** | 2.59 |
| **3VSY_A** | 2.52 |
| **1CQS_A** | 2.7 |
| **1E3R_A** | 2.52 |
| **1E3V_A** | 2.67 |
| **1OGX_A** | 2.59 |
| **1VZZ_A** | 2.58 |
| **1W00_A** | 2.6 |
| **3FZW** | 2.43 |
| **3VGN_A** | 2.55 |
| **5D82_A** | 2.41 |
| **5D83_A** | 2.61 |
| **5KP3_A** | 2.43 |
| **5G2G_A** | 2.33 |
| **1CQS_B** | 2.42 |
| **1E3R_B** | 2.51 |
| **1E3V_B** | 2.71 |
| **1OGX_B** | 2.42 |
| **1VZZ_B** | 2.52 |
| **1W00_B** | 2.61 |
| **3FZW_B** | 2.4 |
| **3VGN_B** | 2.55 |
| **5D82_B** | 2.49 |
| **5D83_B** | 2.45 |
| **5KP3_B** | 2.31 |
| **5KP4_B** | 2.76 |
| **5G2G_B** | 2.43 |
| **3OWS_A** | 2.5 |
| **3OWS_B** | 2.55 |
| **3OWS_C** | 2.43 |
| **3OWS_D** | 2.45 |
| **3OWU_A** | 2.44 |
| **3OWU_B** | 2.49 |

| | |
|---|---|
| **3OWU_C** | 2.46 |
| **3OWU_D** | 2.53 |
| **2PZV_A** | 2.51 |
| **2PZV_B** | 2.53 |
| **2PZV_C** | 2.49 |
| **2PZV_D** | 2.5 |
| **3OWY_A** | 2.59 |
| **3OWY_B** | 2.54 |
| **3OWY_C** | 2.69 |
| **3OWY_D** | 2.65 |
| **3OWY_E** | 2.56 |
| **3OWY_F** | 2.62 |
| **3OWY_G** | 2.73 |
| **3OWY_H** | 2.6 |
| **3OX9_A** | 2.55 |
| **3OX9_B** | 2.57 |
| **3OX9_C** | 2.55 |
| **3OX9_D** | 2.56 |
| **3OXA_A** | 2.63 |
| **3OXA_B** | 2.61 |
| **3OXA_C** | 2.57 |
| **3OXA_D** | 2.59 |
| **5KP1_A** | 2.48 |
| **5KP1_B** | 2.47 |
| **5KP1_C** | 2.46 |
| **5KP1_D** | 2.46 |

**Table S49**. Lengths of the Y16 – Y57 hydrogen bond in all KSI crystal structures that contain intact Y16 and Y57 (n=70, **Table S1**). 5KP4 molecule B was not included, as in this molecule, Y16 and Y32 instead Y57 make a hydrogen bond.

| Structure | D/N 40 Xδ1 – W120 Nε1 Hydrogen bond length (Å) |
|---|---|
| **1C7H** | 2.56 |
| **1DMM** | 2.81 |
| **1DMN** | 2.92 |
| **1DMQ** | 2.94 |
| **1E97** | 2.68 |
| **1EA2** | 2.71 |
| **1GS3** | 2.68 |
| **1OHO** | 2.74 |
| **1OPY** | 2.89 |
| **1W02** | 2.83 |
| **1W6Y** | 2.64 |
| **2INX** | 2.86 |
| **3CPO** | 2.81 |
| **3RGR** | 2.81 |
| **3SED** | 2.84 |
| **4K1V** | 2.75 |
| **5AI1** | 2.84 |
| **5D81** | 2.84 |
| **1CQS_A** | 2.79 |
| **1E3R_A** | 2.75 |
| **1E3V_A** | 2.91 |
| **1K41_A** | 2.85 |
| **1OGX_A** | 2.82 |
| **1OH0_A** | 2.83 |
| **1VZZ_A** | 2.77 |
| **1W00_A** | 2.68 |
| **1W01_A** | 2.79 |
| **3FZW_A** | 2.80 |
| **3VGN_A** | 2.77 |
| **3VSY_A** | 2.84 |
| **4K1U_A** | 2.72 |
| **5D82_A** | 2.81 |
| **5D83_A** | 2.82 |
| **5KP3_A** | 2.76 |
| **5KP4_A** | 2.77 |
| **5G2G_A** | 2.93 |
| **1CQS_B** | 2.89 |
| **1E3R_B** | 2.82 |
| **1E3V_B** | 2.78 |
| **1K41_B** | 2.98 |
| **1OGX_B** | 2.77 |
| **1OH0_B** | 2.86 |
| **1VZZ_B** | 2.77 |
| **1W00_B** | 2.94 |
| **1W01_B** | 2.68 |
| **3FZW_B** | 2.86 |

| | |
|---|---|
| **3VGN_B** | 2.79 |
| **3VSY_B** | 2.90 |
| **4K1U_B** | 2.67 |
| **5D82_B** | 2.87 |
| **5D83_B** | 2.85 |
| **5KP3_B** | 2.85 |
| **5KP4_B** | 2.81 |
| **5G2G_B** | 2.85 |
| **3OWU_A** | 2.74 |
| **3OWU_B** | 2.72 |
| **3OWU_C** | 2.69 |
| **3OWU_D** | 2.78 |
| **3OWS_A** | 2.90 |
| **3OWS_B** | 2.80 |
| **3OWS_C** | 2.68 |
| **3OWS_D** | 2.89 |
| **2PZV_A** | 2.88 |
| **2PZV_B** | 2.82 |
| **2PZV_C** | 2.80 |
| **2PZV_D** | 2.84 |
| **3IPT_A** | 2.80 |
| **3IPT_B** | 2.80 |
| **3IPT_C** | 2.79 |
| **3IPT_D** | 2.68 |
| **3OWY_A** | 2.78 |
| **3OWY_B** | 2.64 |
| **3OWY_C** | 2.98 |
| **3OWY_D** | 2.67 |
| **3OWY_E** | 2.69 |
| **3OWY_F** | 2.64 |
| **3OWY_G** | 2.82 |
| **3OWY_H** | 2.68 |
| **3OX9_A** | 2.94 |
| **3OX9_B** | 2.97 |
| **3OX9_C** | 2.91 |
| **3OX9_D** | 2.94 |
| **3OXA_A** | 2.99 |
| **3OXA_B** | 3.00 |
| **3OXA_C** | 3.02 |
| **3OXA_D** | 3.02 |
| **3T8N_A** | 2.81 |
| **3T8N_B** | 2.84 |
| **3T8N_D** | 2.82 |
| **3T8N_F** | 2.81 |
| **5KP1_A** | 2.82 |
| **5KP1_A** | 2.91 |
| **5KP1_C** | 2.89 |
| **5KP1_D** | 2.81 |

**Table S50.** Lengths of the D/N40 – W120 hydrogen bond in all KSI crystal structures (n=94, **Table S1**).

| Structure | Distance (Å) | |
|---|---|---|
| | M84 – Y16 | M31 – Y16 |
| 1C7H | 3.84 | 4.43 |
| 1DMQ | 3.55 | 4.41 |
| 1OPY | 3.58 | 4.43 |
| 1W6Y | 3.85 | 4.49 |
| 2INX | 3.58 | 4.54 |
| 3CPO | 3.67 | 4.54 |
| 3RGR | 3.74 | 4.46 |
| 3SED | 3.71 | 4.62 |
| 5D81 | 3.71 | 4.63 |
| 1OH0_A | 3.61 | 4.54 |
| 1OH0_B | 3.67 | 4.52 |
| 3VSY_A | 3.73 | 4.35 |
| 3VSY_B | 3.63 | 4.61 |
| 1CQS_A | 4.13 | 4.44 |
| 1E3R_A | 3.68 | 4.58 |
| 1E3V_A | 3.77 | 4.51 |
| 1OGX_A | 3.80 | 4.45 |
| 1VZZ_A | 3.97 | 4.41 |
| 1W00_A | 3.90 | 4.34 |
| 3FZW_A | 3.79 | 4.47 |
| 3VGN_A | 3.66 | 4.61 |
| 5D82_A | 3.68 | 4.63 |
| 5D83_A | 3.64 | 4.44 |
| 5KP3_A | 3.65 | 4.71 |
| 5G2G_A | 3.66 | 4.60 |
| 1CQS_B | 3.91 | 4.57 |
| 1E3R_B | 3.52 | 4.51 |
| 1E3V_B | 3.75 | 4.44 |
| 1OGX_B | 3.68 | 4.61 |
| 1VZZ_B | 3.99 | 4.62 |
| 1W00_B | 3.95 | 4.61 |
| 3FZW_B | 3.76 | 4.56 |
| 3VGN_B | 3.63 | 4.56 |
| 5D82_B | 3.84 | 4.24 |
| 5D83_B | 3.73 | 4.69 |
| 5KP3_B | 3.71 | 4.74 |
| 5KP4_B | 3.68 | 4.35 |
| 5G2G_B | 3.69 | 4.35 |
| 3OWS_A | 3.63 | 4.38 |
| 3OWS_B | 3.67 | 4.77 |
| 3OWS_C | 3.70 | 4.50 |
| 3OWS_D | 3.69 | 4.56 |
| 3OWU_A | 3.77 | 4.62 |
| 3OWU_B | 3.65 | 4.77 |
| 3OWU_C | 3.55 | 4.47 |
| 3OWU_D | 3.72 | 4.55 |

| | | |
|---|---|---|
| **2PZV_A** | 3.60 | 4.51 |
| **2PZV_B** | 3.64 | 4.55 |
| **2PZV_C** | 3.63 | 4.54 |
| **2PZV_D** | 3.63 | 4.52 |
| **3OWY_A** | 3.70 | 4.72 |
| **3OWY_B** | 3.57 | 4.52 |
| **3OWY_C** | 3.51 | 4.81 |
| **3OWY_D** | 3.51 | 4.50 |
| **3OWY_E** | 3.52 | 4.68 |
| **3OWY_F** | 3.51 | 4.70 |
| **3OWY_G** | 3.53 | 4.64 |
| **3OWY_H** | 3.61 | 4.62 |
| **3OX9_A** | 3.68 | 4.57 |
| **3OX9_B** | 3.66 | 4.63 |
| **3OX9_C** | 3.66 | 4.67 |
| **3OX9_D** | 3.73 | 4.63 |
| **3OXA_A** | 3.68 | 4.61 |
| **3OXA_B** | 3.69 | 4.59 |
| **3OXA_C** | 3.66 | 4.63 |
| **3OXA_D** | 3.69 | 4.64 |
| **5KP1_A** | 3.68 | 4.51 |
| **5KP1_B** | 3.72 | 4.51 |
| **5KP1_C** | 3.73 | 4.52 |
| **5KP1_D** | 3.65 | 4.53 |
| **Mean** | 3.69 | 4.55 |

**Table S51.** Y16 Cδ2 – M84 Cε and Y16 Cδ1 – M31 Sδ distances in all KSI crystal structures with intact Y16-Y57 hydrogen bond (n=70, see **Table S1 and S2**).

| Structure | Distance (Å) | |
|---|---|---|
| | M84 – Y16 | M31 – Y16 |
| **1DMM** | 3.69 | 3.99 |
| **1DMN** | 3.73 | 4.23 |
| **1E97** | 3.80 | 4.24 |
| **1EA2** | 4.07 | 3.98 |
| **1GS3** | 3.57 | 4.34 |
| **1OHO** | 3.86 | 4.15 |
| **1W02** | 4.38 | 3.89 |
| **4KIV** | 3.77 | 4.36 |
| **5AI1** | 3.74 | 4.40 |
| **1K41_A** | 4.35 | 4.59 |
| **1W01_A** | 4.27 | 3.98 |
| **4K1U_A** | 3.98 | 4.29 |
| **1K41_B** | 4.31 | 4.50 |
| **1W01_B** | 4.39 | 4.07 |
| **4K1U_B** | 3.91 | 4.25 |
| **Mean** | 3.99 | 4.22 |

**Table S52**. Y16 Cδ2 – M84 Cε and Y16 Cδ1 – M31 Sδ distances in all KSI crystal structures in which the Y16-Y57 hydrogen bond has been ablated (i.e. crystal structures with Y16F and Y57X mutations, in which X is any residue (n=15, see **Table S1 and S2**).

| Structure | Side chain χ1 angle (°) | | | | | | | | | |
|---|---|---|---|---|---|---|---|---|---|---|
| | D21 | D24 | D34 | D35 | D40 | N79 | D100 | D103 | D108 | N124 |
| 1E97 | -171.07 | -160.35 | 74.26 | 58.44 | -169.88 | 67.72 | -68.74 | -93.21 | 68.12 | -56.11 |
| 1EA2 | -175.87 | -163.34 | 71.30 | 60.17 | -164.16 | 66.61 | -64.25 | -87.88 | 63.33 | -58.86 |
| 1OHO | -175.17 | -164.57 | 63.22 | 58.08 | -155.61 | 63.44 | -62.30 | -84.39 | 66.07 | -59.84 |
| 1OPY | -174.66 | -164.75 | 67.48 | 62.03 | -172.13 | 66.06 | -64.05 | -87.95 | 64.79 | -59.48 |
| 2INX | -174.86 | -164.45 | 62.74 | 64.23 | -175.99 | 70.89 | -62.19 | -78.58 | 63.46 | -64.24 |
| 3CPO | -176.99 | -165.41 | 64.60 | 60.38 | -175.15 | 66.23 | -65.98 | -79.28 | 64.37 | -65.17 |
| 3RGR | -173.70 | -165.41 | 64.60 | 60.38 | -173.51 | 70.80 | -63.05 | -82.54 | 68.09 | -64.01 |
| 3SED | -177.11 | -167.98 | 63.13 | 63.48 | -178.03 | 73.17 | -66.08 | -86.14 | 67.90 | -63.85 |
| 4K1V | -176.03 | -165.01 | 62.59 | 58.71 | -173.04 | 67.58 | -69.17 | -87.67 | 63.46 | -57.32 |
| 5D81 | -175.46 | -165.47 | 62.71 | 66.37 | -175.61 | 69.32 | -63.48 | -81.35 | 65.77 | -62.99 |
| 1OH0_A | -175.11 | -163.84 | 67.70 | 67.21 | 178.81 | 68.69 | -63.13 | -83.82 | 64.58 | -60.38 |
| 1OH0_B | -175.11 | -163.84 | 67.70 | 67.21 | 178.81 | 68.69 | -63.13 | -83.82 | 64.58 | -60.38 |
| 3VSY_A | -176.65 | -165.77 | -69.90 | 55.08 | -173.80 | 72.19 | -65.03 | -79.25 | 68.53 | -60.87 |
| 3VSY_B | -176.16 | -168.88 | 59.33 | 62.11 | -174.21 | 70.43 | -64.03 | -77.23 | 69.72 | -62.06 |
| 1E3V_A | -174.48 | -167.10 | 66.02 | 61.67 | -170.46 | 71.34 | -61.53 | -85.93 | 62.51 | -57.05 |
| 1OGX_A | -168.62 | -167.92 | 62.35 | 62.31 | -176.81 | 66.49 | -65.66 | -82.48 | 61.73 | -58.48 |
| 3FZW_A | -174.86 | -170.30 | 63.69 | 56.97 | -177.17 | 67.03 | -62.27 | -79.56 | 68.41 | -60.26 |
| 3VGN_A | -178.84 | -162.20 | 62.67 | 57.77 | 176.43 | 68.73 | -65.47 | -77.56 | 65.07 | -64.82 |
| 4K1U_A | -174.57 | -167.14 | 116.05 | 57.60 | -177.16 | 64.54 | -66.19 | -86.95 | 62.93 | -57.52 |
| 5D82_A | -175.45 | -168.29 | 62.09 | 64.09 | -175.57 | 68.55 | -64.75 | -80.74 | 64.79 | -62.91 |
| 5D83_A | -168.86 | -165.75 | 53.93 | 62.78 | -171.10 | 63.03 | -64.35 | -83.76 | 62.15 | -63.95 |
| 5KP3_A | 176.68 | -163.74 | 60.41 | 61.78 | -173.24 | 66.83 | -64.69 | -80.01 | 66.37 | -61.16 |
| 5G2G_A | 179.67 | -167.29 | 62.68 | 61.01 | 177.35 | 65.70 | -64.33 | -88.33 | 64.15 | -61.55 |
| 1E3V_B | -175.67 | -164.13 | -83.58 | 61.60 | -171.84 | 75.79 | -65.42 | -84.79 | 65.40 | -60.27 |
| 1OGX_B | -175.67 | -164.13 | -83.58 | 61.60 | -171.84 | 75.79 | -65.42 | -84.79 | 65.40 | -60.27 |
| 3FZW_B | -172.99 | -165.80 | -83.58 | 71.16 | -174.90 | 68.34 | -66.99 | -81.14 | 66.86 | -63.12 |
| 3VGN_B | -178.22 | -167.63 | 60.72 | 64.78 | 177.78 | 71.74 | -65.27 | -76.77 | 65.28 | -65.56 |
| 4K1U_B | -174.79 | -162.99 | -76.62 | 50.39 | -174.54 | 66.89 | -58.83 | -79.15 | 56.34 | -60.55 |
| 5D82_B | -172.94 | -161.87 | -66.63 | 66.57 | -172.66 | 68.24 | -63.43 | -83.99 | 65.17 | -60.64 |
| 5D83_B | -176.75 | -170.43 | 64.55 | 56.53 | -171.97 | 68.59 | -59.37 | -83.38 | 62.82 | -61.27 |
| 5KP3_B | 176.00 | -167.54 | 43.30 | 66.11 | -172.65 | 62.35 | -65.92 | -87.78 | 59.79 | -63.21 |
| 5KP4_B | -171.60 | -165.72 | -56.37 | 57.47 | -174.17 | 60.32 | -66.13 | -82.65 | 57.69 | -61.44 |
| 5G2G_B | -174.20 | -165.43 | -73.41 | 40.27 | 177.17 | 65.10 | -63.36 | -86.54 | 67.33 | -64.87 |
| 3OWS_A | -168.11 | -163.97 | 22.95 | 54.68 | -177.42 | 84.59 | -60.86 | -85.77 | 69.53 | -57.00 |
| 3OWS_B | 179.44 | -162.58 | 35.48 | 62.51 | -176.69 | 83.02 | -59.34 | -79.84 | 67.14 | -58.09 |
| 3OWS_C | -173.40 | -164.92 | 40.53 | 52.35 | -175.04 | 73.73 | -63.00 | -85.76 | 65.43 | -64.83 |
| 3OWS_D | -173.36 | -166.94 | 25.51 | 55.79 | -178.82 | 79.97 | -59.50 | -87.27 | 67.42 | -64.52 |
| 2PZV_A | -177.68 | -168.20 | 61.60 | 64.27 | -174.67 | 67.98 | -66.63 | -82.50 | 74.32 | -62.02 |
| 2PZV_B | -174.82 | -169.22 | 63.26 | 61.58 | -173.80 | 70.58 | -66.10 | -82.83 | 66.98 | -63.12 |
| 2PZV_C | -177.37 | -167.37 | 59.08 | 63.20 | -178.41 | 63.26 | -66.67 | -78.40 | 70.69 | -64.18 |
| 2PZV_D | -178.52 | -169.64 | 60.73 | 62.26 | -172.54 | 66.15 | -61.97 | -80.71 | 67.37 | -66.15 |
| 3IPT_A | 177.06 | -178.32 | 60.71 | 60.91 | -178.63 | 62.34 | -65.40 | -83.23 | 60.25 | -62.99 |
| 3IPT_B | -175.62 | -174.14 | 61.03 | 59.06 | 178.25 | 63.02 | -67.70 | -88.35 | 59.76 | -57.27 |
| 3IPT_C | -177.74 | -171.01 | 61.28 | 57.06 | -170.16 | 65.37 | -62.06 | -84.40 | 64.42 | -61.61 |
| 3IPT_D | -176.46 | -175.11 | 60.35 | 59.35 | -172.47 | 60.00 | -56.31 | -90.61 | 60.77 | -58.71 |
| 3OXA_A | -171.92 | -167.32 | 44.46 | 64.09 | -170.32 | 80.60 | -64.49 | -87.07 | 66.56 | -60.02 |
| 3OXA_B | -171.17 | -167.64 | 44.69 | 63.60 | -171.30 | 80.50 | -61.97 | -84.84 | 67.33 | -59.17 |
| 3OXA_C | -172.20 | -167.51 | 44.07 | 62.32 | -172.12 | 80.43 | -62.07 | -84.78 | 66.79 | -60.51 |
| 3OXA_D | -170.61 | -168.96 | 43.57 | 63.37 | -171.15 | 79.11 | -63.58 | -85.88 | 68.47 | -59.25 |
| 5KP1_A | -173.20 | -170.56 | 66.80 | 61.42 | -173.89 | 69.24 | -64.10 | -76.09 | 71.97 | -58.82 |
| 5KP1_B | -171.72 | -166.80 | 66.80 | 68.26 | -172.69 | 68.44 | -63.57 | -76.01 | 66.42 | -63.31 |
| 5KP1_C | -171.73 | -165.94 | 66.80 | 69.49 | -173.62 | 66.00 | -65.05 | -74.93 | 65.57 | -62.16 |
| 5KP1_D | -173.35 | -169.47 | 68.31 | 62.38 | -173.79 | 68.82 | -61.58 | -76.83 | 71.38 | -59.43 |

**Table S53**. Side chain χ1 dihedral angles for all aspartate and asparagine residues in KSI from the reduced pseudo-ensemble. Asparagine residues at position 2 and position 93 have been excluded from the analysis, because these residues are situated in the highly flexible N-terminus of the protein and the 91-96 loop, respectively.

| Structure | Side chain χ1 angle (°) | | | | | | | |
|---|---|---|---|---|---|---|---|---|
| | **Y16** | **Y32** | **F42** | **F56** | **Y57** | **F86** | **F107** | **Y119** |
| **1OPY** | 170.46 | -76.03 | 177.96 | 164.93 | -78.58 | 66.91 | -65.80 | -67.48 |
| **2INX** | 170.59 | -77.61 | -172.79 | 172.35 | -78.19 | 63.32 | -68.42 | -70.77 |
| **3CPO** | 169.66 | -79.43 | -175.94 | 174.96 | -78.21 | 61.84 | -68.12 | -73.40 |
| **3RGR** | 168.38 | -77.54 | -175.33 | 171.22 | -73.99 | 57.90 | -67.40 | -71.06 |
| **3SED** | 169.33 | -76.54 | -178.03 | 176.45 | -77.83 | 63.90 | -70.43 | -74.11 |
| **5D81** | 168.81 | -80.65 | -171.25 | 179.82 | -77.83 | 60.51 | -68.86 | -68.90 |
| **1OH0_A** | 165.56 | -78.96 | -178.52 | -179.44 | -80.43 | 62.59 | -68.74 | -74.03 |
| **1OH0_B** | 165.56 | -78.96 | -178.52 | -179.44 | -80.43 | 62.59 | -68.74 | -74.03 |
| **3VSY_A** | 171.44 | -79.73 | -173.63 | 167.92 | -76.91 | 58.74 | -70.79 | -71.29 |
| **3VSY_B** | 171.44 | -81.29 | -172.57 | 170.33 | -77.28 | 61.48 | -71.10 | -67.73 |
| **1E3V_A** | 169.50 | -78.01 | -172.43 | 169.03 | -77.21 | 61.70 | -64.57 | -70.17 |
| **1OGX_A** | 167.16 | -75.87 | -176.29 | 175.44 | -67.06 | 64.39 | -66.51 | -68.47 |
| **3FZW_A** | 168.05 | -82.40 | -175.78 | 175.30 | -74.50 | 58.45 | -66.79 | -77.47 |
| **3VGN_A** | 166.98 | -79.76 | -174.61 | 162.71 | -75.71 | 60.39 | -69.68 | -74.32 |
| **5D82_A** | 171.10 | -81.24 | -171.07 | 174.00 | -81.76 | 62.58 | -67.90 | -67.60 |
| **5D83_A** | 169.87 | -85.81 | -170.19 | 171.97 | -78.93 | 60.59 | -71.23 | -68.74 |
| **5KP3_A** | 164.43 | -83.31 | -171.69 | 179.06 | -85.64 | 60.86 | -69.03 | -67.85 |
| **5G2G_A** | 164.86 | -80.46 | -174.00 | 176.11 | -79.25 | 62.77 | -68.33 | -72.22 |
| **1E3V_B** | 168.49 | -77.56 | -176.53 | 169.41 | -75.66 | 62.65 | -62.98 | -67.93 |
| **1OGX_B** | 168.49 | -77.56 | -176.53 | 169.41 | -75.66 | 62.65 | -62.98 | -67.93 |
| **3FZW_B** | 167.00 | -79.08 | -173.10 | 178.34 | -74.20 | 58.74 | -66.58 | -72.10 |
| **3VGN_B** | 167.36 | -77.10 | -175.20 | 164.89 | -74.07 | 58.17 | -71.34 | -75.93 |
| **5D82_B** | 172.63 | -79.60 | -174.23 | 171.21 | -82.27 | 61.10 | -67.34 | -71.58 |
| **5D83_B** | 166.27 | -90.05 | -174.22 | 176.60 | -80.48 | 61.59 | -68.74 | -68.83 |
| **5KP3_B** | 166.01 | -80.05 | -172.54 | 178.50 | -83.11 | 61.21 | -69.81 | -71.73 |
| **5KP4_B** | 167.41 | -75.54 | -169.70 | 169.93 | -77.70 | 60.35 | -66.68 | -72.31 |
| **5G2G_B** | 170.83 | -78.43 | -167.68 | 171.79 | -78.76 | 58.98 | -66.03 | -72.78 |
| **3OWS_A** | 164.40 | -84.79 | -174.59 | 168.34 | -79.14 | 60.92 | -68.10 | -66.76 |
| **3OWS_B** | 164.39 | -83.04 | 178.64 | 165.19 | -80.10 | 62.57 | -65.45 | -69.10 |
| **3OWS_C** | 168.01 | -80.78 | 179.77 | 169.66 | -80.39 | 59.81 | -67.94 | -73.61 |
| **3OWS_D** | 166.91 | -85.77 | -179.50 | 167.76 | -80.86 | 61.59 | -71.71 | -74.66 |
| **2PZV_A** | 173.73 | -73.12 | -175.54 | 174.28 | -77.53 | 58.01 | -66.34 | -64.18 |
| **2PZV_B** | 172.17 | -77.10 | -174.88 | 167.98 | -81.25 | 59.04 | -70.18 | -69.43 |
| **2PZV_C** | 171.08 | -75.39 | -173.36 | 173.42 | -77.83 | 61.09 | -66.25 | -70.21 |
| **2PZV_D** | 174.47 | -76.64 | -170.00 | 172.49 | -78.49 | 63.39 | -69.91 | -68.21 |
| **3OXA_A** | 170.71 | -83.52 | 178.90 | 173.25 | -79.99 | 62.06 | -64.25 | -64.98 |
| **3OXA_B** | 171.42 | -82.65 | 179.49 | 172.94 | -79.08 | 61.91 | -64.06 | -68.17 |
| **3OXA_C** | 170.67 | -82.41 | 178.78 | 171.74 | -79.09 | 62.70 | -64.23 | -66.43 |
| **3OXA_D** | 167.93 | -82.38 | 179.27 | 171.78 | -80.75 | 60.59 | -63.48 | -66.44 |
| **5KP1_A** | 169.14 | -78.28 | -175.00 | -175.07 | -76.05 | 60.92 | -67.74 | -79.93 |
| **5KP1_B** | 167.79 | -79.35 | -168.98 | 170.79 | -70.80 | 62.37 | -68.22 | -69.06 |
| **5KP1_C** | 167.82 | -78.85 | -167.59 | 171.69 | -72.29 | 61.33 | -68.14 | -69.34 |
| **5KP1_D** | 168.37 | -78.74 | -175.84 | 178.53 | -76.42 | 62.40 | -67.70 | -78.31 |

**Table S54**. Side chain χ1 dihedral angles for all tyrosine and phenylalanine residues in KSI from the reduced pseudo-ensemble. χ1 angles for position 16 include Tyrosine residues only (phenylalanine substitutions have been omitted).

| | Angle OH(X) – TSA ring plane (°) | | Transformed angle | |
|---|---|---|---|---|
| **Structure** | **X=Y16** | **X=D103** | **180° - angle (Y16–TSA)** | **Angle (D103–TSA)*(-1)** |
| **1GS3** | 145.4 | -21 | 34.6 | 21 |
| **1W6Y** | 149.2 | -48 | 30.8 | 48 |
| **3CPO** | 163.7 | -52.4 | 16.3 | 52.4 |
| **5AI1** | 149.4 | 4.3 | 30.6 | -4.3 |
| **1OGX_A** | 151.3 | -55.4 | 28.7 | 55.4 |
| **1OGX_B** | 153.8 | -35.2 | 26.2 | 35.2 |
| **1OH0_A** | 143.1 | -42.6 | 36.9 | 42.6 |
| **1OH0_B** | 150 | -36.5 | 30 | 36.5 |
| **3FZW_A** | 154.8 | -45.8 | 25.2 | 45.8 |
| **3FZW_B** | 156.2 | -33.1 | 23.8 | 33.1 |
| **3OWS_A** | 155.5 | -32 | 24.5 | 32 |
| **3OWS_B** | 146.9 | -26.6 | 33.1 | 26.6 |
| **3OWS_C** | 151.7 | -27.9 | 28.3 | 27.9 |
| **3OWS_D** | 149.2 | -17.6 | 30.8 | 17.6 |
| **3OWU_A** | 149.4 | -20.9 | 30.6 | 20.9 |
| **3OWU_B** | 146.7 | -34.1 | 33.3 | 34.1 |
| **3OWU_C** | 154.9 | -31.7 | 25.1 | 31.7 |
| **3OWU_D** | 148.6 | -31.2 | 31.4 | 31.2 |
| **3VGN_A** | 135.4 | -34.4 | 44.6 | 34.4 |
| **3VGN_B** | 140 | -34.7 | 40 | 34.7 |
| **5KP3_A** | 144.2 | -33.8 | 35.8 | 33.8 |
| **5KP3_B** | 139.1 | -25.9 | 40.9 | 25.9 |
| **2PZV_A** | 162.4 | -51.8 | 17.6 | 51.8 |
| **2PZV_B** | 167.4 | -47 | 12.6 | 47 |
| **2PZV_C** | 166.9 | -47.7 | 13.1 | 47.7 |
| **2PZV_D** | 168.6 | -38 | 11.4 | 38 |
| **3OWY_A** | 153.3 | -31.3 | 26.7 | 31.3 |
| **3OWY_B** | 160 | -13 | 20 | 13 |
| **3OWY_C** | 146.9 | -54.1 | 33.1 | 54.1 |
| **3OWY_D** | 160.6 | -34.7 | 19.4 | 34.7 |
| **3OWY_E** | 147.2 | -42.8 | 32.8 | 42.8 |
| **3OWY_H** | 140.3 | -69.8 | 39.7 | 69.8 |
| **5PK1_A** | 160 | -48.7 | 20 | 48.7 |
| **5PK1_B** | 161.4 | -44.9 | 18.6 | 44.9 |
| **5PK1_C** | 161.5 | -44 | 18.5 | 44 |
| **5PK1_D** | 161.2 | -48.9 | 18.8 | 48.9 |

**Table S55**. Angles between the hydrogen bond donors Y16 and D103 and the plane of the A ring of transition state analogs bound to KSI (see Tables S1 and S2). Dihedral angle is measured between Y16 or D103 OH atom and Ox, C1, C2 atoms of the bound transition state analog (OH – Ox – C1 – C2, two columns on the left). In the two columns on the right the values have been transformed mathematically for clarity of presentation.

| Structure | Residue at position 40 (X) | Distance D40 – TSA (Å) | | | Bound Ligand |
|---|---|---|---|---|---|
| | | Xδ2 - C2 | Xδ2 - C4 | Xδ2 - C10 | |
| **1GS3** | N | 3.23 | 3.92 | 3.19 | Equ |
| **1W6Y** | D | 3.29 | 4.21 | 3.06 | Equ |
| **3CPO** | N | 3.18 | 4.1 | - | Phe |
| **5AI1** | N | 3.09 | 4.02 | 3.33 | Equ |
| **1OGX_A** | N | 3.08 | 4.17 | 3.31 | Equ |
| **1OH0_A** | D | 3.47 | 4.51 | 3.52 | Equ |
| **3FZW_A** | N | 3.28 | 4.34 | 3.68 | Equ |
| **3VGN_A** | N | 5.21 | 4.86 | - | Phe |
| **5KP3_A** | N | 3.24 | 4.37 | 3.65 | Equ |
| **1OGX_B** | N | 3.42 | 4.24 | 3.57 | Equ |
| **1OH0_B** | D | 3.31 | 4.37 | 3.67 | Equ |
| **3FZW_B** | N | 3.33 | 4.35 | 3.86 | Equ |
| **3VGN_B** | N | 5.58 | 5.27 | - | Phe |
| **5KP3_B** | N | 3.21 | 4.16 | 3.47 | Equ |
| **3OWS_A** | N | 3.19 | 4.12 | 3.65 | Equ |
| **3OWS_B** | N | 3.36 | 4.2 | 3.49 | Equ |
| **3OWS_C** | N | 3.15 | 3.98 | 3.3 | Equ |
| **3OWS_D** | N | 3.26 | 4.03 | 3.5 | Equ |
| **3OWU_A** | N | 3.45 | 3.96 | 3.8 | Equ |
| **3OWU_B** | N | 3.35 | 4 | 3.53 | Equ |
| **3OWU_C** | N | 3.35 | 4.01 | 3.8 | Equ |
| **3OWU_D** | N | 3.14 | 4.03 | 3.66 | Equ |
| **2PZV_A** | N | 3.18 | 4.27 | - | Phe |
| **2PZV_B** | N | 3.13 | 3.48 | - | Phe |
| **2PZV_C** | N | 3.25 | 4.17 | - | Phe |
| **2PZV_D** | N | 3.18 | 3.89 | - | Phe |
| **3OWY_A** | N | 3.19 | 3.89 | 3.27 | Equ |
| **3OWY_B** | N | 3.47 | 3.78 | 3.77 | Equ |
| **3OWY_C** | N | 3.34 | 4.19 | 3.42 | Equ |
| **3OWY_D** | N | 3.21 | 3.77 | 3.39 | Equ |
| **3OWY_E** | N | 3.52 | 3.89 | 3.23 | Equ |
| **3OWY_H** | N | 3.08 | 4.04 | 3.23 | Equ |
| **5KP1_A** | N | 3.24 | 4.31 | 3.52 | Equ |
| **5KP1_B** | N | 3.17 | 4.36 | 3.85 | Equ |
| **5KP1_C** | N | 3.19 | 4.35 | 3.85 | Equ |
| **5KP1_D** | N | 3.22 | 4.28 | 3.53 | Equ |

**Table S56**. Distances between the Xδ2 (Oδ2 and Nδ2 for aspartate and asparagine at position 40, respectively) and the carbon positions between which protons are shuffled in KSI reactions from KSI crystal with WT-like activity (see Table S1 and S2).

| Structure | Angle (°) | | |
|---|---|---|---|
| | Oδ1-Nε1-Cζ2 | O/Nδ2-Oδ1-Nε1-Cζ2 | O/ND2-OD1-Nε1 |
| **1C7H** | 97.0 | 130.7 | 121.7 |
| **1DMM** | 93.1 | 133.4 | 127.98 |
| **1DMN** | 97.1 | 148.4 | 130.71 |
| **1DMQ** | 99.0 | 145.8 | 130.8 |
| **1E 97** | 99.2 | 131.1 | 125.26 |
| **1EA2** | 93.5 | 140.9 | 125.55 |
| **1GS3** | 93.9 | 131.6 | 134.05 |
| **1OHO** | 89.5 | 156.7 | 119.08 |
| **1OPY** | 94.3 | 137.6 | 128.51 |
| **1W02** | 79.9 | 156.7 | 114.76 |
| **1W6Y** | 91.9 | 99.5 | 127.11 |
| **2INX** | 93.3 | 133.6 | 129.42 |
| **3CPO** | 95.9 | 133.5 | 132.51 |
| **3RGR** | 93.8 | 144.7 | 134.02 |
| **3SED** | 97.0 | 135.2 | 130.69 |
| **4K1V** | 94.9 | 135.1 | 130.2 |
| **5AI1** | 94.4 | 135.0 | 129.93 |
| **5D81** | 96.2 | 128.9 | 133.05 |
| **1CQS_A** | 86.8 | 108.8 | 137.41 |
| **1E3R_A** | 101.2 | 79.6 | 135.32 |
| **1E3V_A** | 94.7 | 134.8 | 127.23 |
| **1K41_A** | 77.8 | 144.0 | 103.11 |
| **1OGX_A** | 89.2 | 123.6 | 133.45 |
| **1OH0_A** | 98.3 | 118.4 | 137.17 |
| **1VZZ_A** | 89.9 | 145.1 | 122.16 |
| **1W00_A** | 97.3 | 124.4 | 131.8 |
| **1W01_A** | 98.1 | 105.4 | 139.96 |
| **3FZW_A** | 98.6 | 124.3 | 133.79 |
| **3VGN_A** | 100.2 | 13.2 | 121.01 |
| **3VSY_A** | 97.1 | 132.2 | 132.2 |
| **4K1U_A** | 89.3 | 135.8 | 127.79 |
| **5D82_A** | 94.5 | 134.9 | 131.17 |
| **5D83_A** | 98.8 | 125.7 | 134.37 |
| **5KP3_A** | 96.9 | 120.7 | 133.2 |
| **5KP4_A** | 99.4 | 99.7 | 148.16 |
| **5G2G_A** | 98.6 | 130.1 | 138.33 |
| **1CQS_B** | 95.4 | 97.2 | 143.14 |
| **1E3R_B** | 104.0 | 102.3 | 132.76 |
| **1E3V_B** | 91.6 | 125.2 | 124.95 |
| **1K41_B** | 90.4 | 137.0 | 136.41 |
| **1OGX_B** | 95.4 | 109.3 | 138.99 |
| **1OH0_B** | 97.4 | 121.5 | 136.22 |
| **1VZZ_B** | 92.9 | 132.1 | 134.42 |
| **1W00_B** | 90.2 | 135.8 | 127.77 |
| **1W01_B** | 89.7 | 123.9 | 127.11 |
| **3FZW_B** | 100.4 | 119.1 | 136.46 |
| **3VGN_B** | 99.1 | 3.3 | 113.37 |

| | | | |
|---|---|---|---|
| **3VSY_B** | 91.6 | 138.3 | 129.75 |
| **4K1U_B** | 87.4 | 138.5 | 127.86 |
| **5D82_B** | 98.2 | 132.4 | 133.38 |
| **5D83_B** | 95.7 | 124.1 | 137.34 |
| **5KP3_B** | 89.8 | 124.8 | 128.14 |
| **5KP4_B** | 91.5 | 136.9 | 125.38 |
| **5G2G_B** | 97.1 | 129.6 | 134.73 |
| **3OWS_A** | 89.0 | 131.4 | 125.93 |
| **3OWS_B** | 94.1 | 124.3 | 129.02 |
| **3OWS_C** | 92.8 | 128.8 | 129.5 |
| **3OWS_D** | 95.4 | 137.2 | 124.4 |
| **3OWU_A** | 90.8 | 129.3 | 135.38 |
| **3OWU_B** | 88.6 | 122.6 | 134.31 |
| **3OWU_C** | 94.1 | 127.9 | 134.83 |
| **3OWU_D** | 90.5 | 128.3 | 129.63 |
| **2PZV_A** | 94.6 | 131.3 | 137.25 |
| **2PZV_B** | 96.0 | 128.3 | 133.94 |
| **2PZV_C** | 95.2 | 124.4 | 136.87 |
| **2PZV_D** | 96.9 | 124.7 | 135.56 |
| **3IPT_A** | 98.9 | 117.5 | 140.06 |
| **3IPT_B** | 95.4 | 123.8 | 135.88 |
| **3IPT_C** | 95.1 | 126.1 | 131.62 |
| **3IPT_D** | 93.1 | 133.6 | 135.52 |
| **3OWY_A** | 85.6 | 123.5 | 116.96 |
| **3OWY_B** | 82.1 | 129.2 | 113.16 |
| **3OWY_C** | 84.6 | 124.4 | 115.63 |
| **3OWY_D** | 86.5 | 127.3 | 117.32 |
| **3OWY_E** | 88.9 | 118.3 | 123.75 |
| **3OWY_F** | 87.8 | 119.8 | 118.27 |
| **3OWY_G** | 87.6 | 131.1 | 116.78 |
| **3OWY_H** | 86.7 | 119.0 | 124.97 |
| **3OX9_A** | 95.4 | 119.7 | 128.44 |
| **3OX9_B** | 97.2 | 120.4 | 130.38 |
| **3OX9_C** | 97.3 | 119.1 | 130.47 |
| **3OX9_D** | 96.1 | 122.2 | 129.33 |
| **3OXA_A** | 94.8 | 139.1 | 130.44 |
| **3OXA_B** | 95.4 | 138.9 | 131.4 |
| **3OXA_C** | 93.6 | 142.8 | 126.57 |
| **3OXA_D** | 93.9 | 142.1 | 129.15 |
| **3T8N_A** | 105.8 | 123.3 | 141.52 |
| **3T8N_B** | 105.0 | 121.9 | 135.48 |
| **3T8N_C** | 100.4 | 112.5 | 139.09 |
| **3T8N_D** | 102.1 | 120.4 | 136.46 |
| **5KP1_A** | 100.4 | 112.1 | 135.16 |
| **5KP1_B** | 100.3 | 121.8 | 136.68 |
| **5KP1_C** | 100.9 | 120.3 | 137.55 |
| **5KP1_D** | 99.8 | 114.7 | 135.06 |

**Table S57**. Angles characterizing the general base and W120 relative sidechain orientations obtained from the full pseudo-ensemble (see Table S1 and S2).

| *Enzyme* | $k_{cat}$ s$^{-1}$ | $K_M$ μM | $k_{cat}$ rel | $K_M$ rel | [E] nM |
|---|---|---|---|---|---|
| KSI WT | 9.7[a] | 29 | (1) | (1) | - |
| KSI W120F | 33.6 | 168.1 | 3.5 | 5.8 | 15 |
| | 34.8 | 174.7 | 3.6 | 6.0 | 40 |
| KSI W120F Average | 34.2 ± 0.6 | 171.4 ± 3.3 | 3.5 | 5.9 | |
| $KSI_{homolog}$ WT | 37[a] | 28 | (1) | (1) | - |
| $KSI_{homolog}$ F120W | 4.9 | 20.4 | 0.13 | 0.7 | 40 |
| | 4.5 | 20.2 | 0.12 | 0.7 | 80 |
| $KSI_{homolog}$ F120W Average | 4.7 ± 0.2 | 20.3 ± 0.1 | 0.13 | 0.7 | |

[a] from (6)

**Table S58**. Michaelis-Menten kinetic parameters for KSI WT and W120F variant and $KSI_{homolog}$ WT and F120W obtained in this study. Kinetics have been measured with the substrate 5(10)-Estrene-3,17-dione as the chemical step for this substrate is rate limiting. Note that W120F mutation in KSI results in an enzyme which is more efficient than the natural (WT) variant.

| Enzyme | $k_{cat}$ ($s^{-1}$) | $K_M$ (μM) | $K_M^{rel}$ | Reference |
|---|---|---|---|---|
| WT | 9.9 ± 0.9 | 19 ± 4 | (1) | (7) |
| Y16A | $4.8 \pm 0.4 \times 10^{-2}$ | 30 ± 10 | 1.6 ± 0.6 | (7) |
| Y16G | $4.0 \pm 0.4 \times 10^{-2}$ | 23 ± 2 | 1.2 ± 0.3 | (7) |
| Y16S | $3.8 \pm 0.5 \times 10^{-2}$ | 18 ± 6 | 0.95 ± 0.4 | (7) |
| Y16T | $5.2 \pm 0.3 \times 10^{-2}$ | 19 ± 3 | 1 ± 0.3 | (7) |
| Y16F | $5.4 \pm 0.1 \times 10^{-4}$ | 41 ± 6 | 2.15 ± 0.6 | (7) |
| WT | 9.9 ± 0.9 | 30 ± 4 | 1 ± 0.2 | (6) |
| D103L | $5.9 \pm 0.1 \times 10^{-2}$ | 39 ± 5 | 1.3 ± 0.2 | (6) |
| D103A | $6.6 \pm 0.2 \times 10^{-2}$ | 52 ± 4 | 1.7 ± 0.3 | (6) |
| D103G | $3.1 \pm 0.3 \times 10^{-1}$ | 51 ± 13 | 1.7 ± 0.5 | (6) |
| Y16F/D103L | $8.0 \pm 2.0 \times 10^{-6}$ | 25 ± 4 | 0.8 ± 0.2 | (6) |
| Y16A/D103A | $1.1 \pm 0.1 \times 10^{-4}$ | 32 ± 1 | 1.1 ± 0.2 | (6) |
| Y16G/D103G | $1.3 \pm 1.2 \times 10^{-3}$ | 36 ± 8 | 1.2 ± 0.3 | (6) |
| D103N | $7.8 \pm 0.1 \times 10^{-1}$ | - | - | (8) |

**Table S59.** Effects of various KSI oxyanion hole mutations on $K_M$ for the substrate 10-EST.

| KSI W120F | $KSI_{homolog}$ F120W |
|---|---|
| TTTTGTTTAACTTTAAGAAGGAGATATACAT<br>ATGAACCTACCGACTGCGCAGGAAGTCCA<br>GGGCCTGATGGCCCGTTACATCGAGCTGG<br>TCGATGTCGGGGATATCGAGGCGATCGTG<br>CAGATGTACGCCGATGACGCCACGGTCGA<br>AGACCCGTTTGGCCAGCCGCCGATCCACG<br>GCCGCGAGCAGATTGCCGCGTTCTATCGC<br>CAGGGTTTGGGCGGAGGCAAGGTCCGCGC<br>CTGCCTGACCGGGCCGGTACGGGCCAGCC<br>ATAACGGCTGCGGGGCGATGCCGTTTCGC<br>GTCGAGATGGTCTGGAACGGCCAGCCCTG<br>TGCACTGGATGTCATCGATGTGATGCGCTT<br>TGATGAGCACGGCCGGATCCAGACGATGC<br>AAGCCTACTTTAGCGAGGTCAACCTCAGCG<br>TGCGCGAGCCGCAGTAGTGAAAGCTTGCG<br>GCCGCACTCGAGCACCACCACCACCACCA<br>CTGAGATCCGGCTGCTAACAAAGCCCGAAA<br>GGAAGCTGAGTTGGCTGCTGCCACCGCTG<br>AGCAATAACTAGCATAACCCCTTGGGGCCT<br>CTAAACGGGTCTTGAGGGGTTTTTTGCTGA<br>AAGGAGGAACTATATCCGGATTGGCGAATG<br>GGACGCGCCCTGTAGCGGCGCATTAAGCG<br>CGGCGGGTGTGGTGGTTACGCGCAGCGTG<br>ACCGCTACACTTGCCAGCGCCCTAGCGCC<br>CGCTCCTTTCGCTTTCTTCCCTTCCTTTCTC<br>GCCACGTTCGCCGGCTTTCCCCGTCAAGC<br>TCTAA | TTTTGTTTAACTTTAAGAAGGAGATATACAT<br>ATGAATACCCCAGAACACATGACCGCCGTG<br>GTACAGCGCTATGTGGCTGCGCTCAATGC<br>CGGCGATCTGGACGGCATCGTCGCGCTGT<br>TTGCCGATGACGCCACGGTGGAAGACCCC<br>GTGGGTTCCGAGCCCAGGTCCGGTACGGC<br>TGCGATTCGTGAGTTTTACGCCAACTCGCT<br>CAAACTGCCTTTGGCGGTGGAGCTGACGC<br>AGGAGGTACGCGCGGTCGCCAACGAAGCG<br>GCCTTCGCTTTCACCGTCAGCTTCGAGTAT<br>CAGGGCCGCAAGACCGTGGTTGCGCCCAT<br>CGATCACTTTCGCTTCAATGGCGCCGGCAA<br>GGTGGTGAGCATGCGCGCCTTGTGGGGCG<br>AGAAGAATATTCACGCTGGCGCCTGAAGCT<br>TGCGGCCGCACTCGAGCACCACCACCACC<br>ACCACTGAGATCCGGCTGCTAACAAAGCCC<br>GAAAGGAAGCTGAGTTGGCTGCTGCCACC<br>GCTGAGCAATAACTAGCATAACCCCTTGGG<br>GCCTCTAAACGGGTCTTGAGGGGTTTTTTG<br>CTGAAAGGAGGAACTATATCCGGATTGGCG<br>AATGGGACGCGCCCTGTAGCGGCGCATTA<br>AGCGCGGCGGGTGTGGTGGTTACGCGCAG<br>CGTGACCGCTACACTTGCCAGCGCCCTAG<br>CGCCCGCTCCTTTCGCTTTCTTCCCTTCCT<br>TTCTCGCCACGTTCGCCGGCTTTCCCCGTC<br>AAGCTCTAAATCGGGGGCTCCCTTTAGGGT<br>TCCGA |

**Table S60.** DNA sequencing results for KSI W120F and $KSI_{homolog}$ F120W (KSI numbering) mutants.
